# *Caenorhabditis briggsae* ancestral genomic hyper-diversity contrasts with globally distributed genome-wide haplotypes

**DOI:** 10.64898/2025.12.08.693002

**Authors:** Nicolas D. Moya, Bowen Wang, Robyn E. Tanny, Michael E.G. Sauria, Lance M. O’Connor, Ayeh Khorshidian, Ryan McKeown, Charlie Gosse, Clayton M. Dilks, Timothy A. Crombie, Gaotian Zhang, Emha Rais, Lise Frézal, Viet Dai Dang, Elkana Haryoso, Mia P. Devi, Clotilde Gimond, Daniel E. Cook, Jung-Chen Hsu, Amanda O. Shaver, Stefan Zdraljevic, Aurélien Richaud, Tongshu Wen, Aatira Mehraj, H Sharanya, Karthick Raja Arulprakasam, Emily J. Koury, Nicole M. Roberto, Etta S. Schaye, Varsha Singh, Hagus Tarno, Michael Ailion, Annalise B. Paaby, Zhongying Zhao, Asher D. Cutter, John Wang, Matthew V. Rockman, Marie-Anne Félix, Christian Braendle, Erik C. Andersen

## Abstract

Comparative genomics provides a powerful framework to uncover the molecular and evolutionary mechanisms that shape genetic diversity within and across species, revealing how shared and lineage-specific processes influence their evolutionary trajectories through time. The nematode *Caenorhabditis briggsae* is distributed world-wide and is a comparative model to *Caenorhabditis elegans* in the biology of development, cellular mechanisms, neurobiology, genetic mappings of complex traits, and genome evolution. Following massive collection efforts by the nematode research community, we present the isolation of over 2,000 wild strains and analyses of genome sequences that catalog over six million single-nucleotide and insertion-deletion variants. This genome and strain resource provide a powerful means to interrogate the causal genetic bases of phenotypic variation for diverse traits. Additionally, we describe its global population structure and discover new and genetically distinct groups within this primarily self-fertilizing species, including groups of highly related strains that were sampled across different continents. We leverage expansive genetic variation to decipher the effects of linkage and selection on the distribution of genetic diversity across the genome and across geographic regions. Within the species, we find genomic regions with extremely high levels of genetic variation similar to hyper-divergent regions found in *C. elegans* and other species. These regions harbor new genes and variation enriched for environmental sensing and pathogen responses. In comparison to the outbreeding sister species *Caenorhabditis nigoni*, we conclude that long-term balancing selection has maintained substantial functional variation since the divergence from their outbreeding ancestor, likely in response to differences in the ecological niche. Overall, this massive strain resource enables future comparative genetics and genomics studies, including genome-wide association studies between *Caenorhabditis* species.

## Introduction

Comparative population genomic analyses among closely related species are invaluable for investigating the evolutionary forces that shape genome diversity both within and between species. Such cross-species population genomic studies reveal whether genetic mechanisms act universally across taxa or reflect lineage-specific phenomena, providing insights into how demographic and selective processes can influence both neutral and functional genetic variation^1–5^. In some cases, closely related species that occupy similar niches exhibit convergent evolution in genotypes and phenotypes^1,2^. For example, the model organism *Drosophila melanogaster* and its sister species *Drosophila simulans* have independently evolved parallel clinal adaptation in genes along climatic gradients to diverse environments^1^. In other cases, shared genotypes among distinct species might be caused by the maintenance of ancestral alleles^3,4^. For example, long-term balancing selection has maintained ancient haplotypes across plant species in the *Capsella* genus, where ancestral polymorphism at immunity-related loci might have been maintained despite divergent pathogen pressures^4^. Comparative analyses also can decipher the evolutionary origins of species-specific traits. For example, comparisons between the yeasts *Saccharomyces cerevisiae* and *Saccharomyces paradoxus* revealed that thermal-tolerance loci evolved under positive selection in an ancestral population of *S. cerevisiae* before spreading across diverse regions and ecological niches^5^. In modern humans, global variation in diverse traits can be traced to adaptive introgression of alleles from closely related but now-extinct archaic Neanderthal and Denisovan populations, including skin and hair pigmentation, metabolism, and immunity^6–8^. It is unresolved to what extent evolutionary convergence, maintenance of ancestral polymorphism, adaptive divergence, and adaptive introgression each contribute to the genetic basis of biodiversity in different taxa. We must use a comparative population genomics approach to answer this question.

*Caenorhabditis* nematodes offer a strong comparative platform for these types of studies. The nematode *Caenorhabditis briggsae* is a self-fertilizing species related to the widely adopted model species *Caenorhabditis elegans*^9–11^. Both species share similar ecology, naturally proliferating in rotting plant material and often isolated in the same substrate^12,13^. Additionally, both species have independently evolved self-fertilization from outbreeding ancestors^14,15^. Extensive collections of wild *C. elegans* strains have been used to establish the genetic basis of phenotypic variation across various traits and to study the effects of self-fertilization on the distribution of genetic diversity across their genomes^16,17^. For example, *C. elegans* carries extreme genetic variation concentrated in punctuated genomic regions thought to be maintained by long-term balancing selection since the evolution of self-fertilization^18^, despite experiencing a general reduction in genetic diversity relative to its outbreeding relatives^19,20^, largely because of chromosome-scale positive selective sweeps^16^ and background selection^21^. These hyper-divergent regions are enriched for genes associated with environmental responses, suggesting that they contribute to adaptation to diverse ecological niches. Although *C. briggsae* offers a powerful comparative platform to study conserved or convergent patterns of genetic diversity with *C. elegans*, a genome-wide analysis of a comparably large *C. briggsae* strain collection has yet to be explored.

Early studies of multilocus genotypes across wild *C. briggsae* strains uncovered evidence of genetic differentiation among geographically separated groups, particularly between strains isolated from tropical and temperate latitudes^22–24^. The incorporation of strains from new geographic locations continued to support population structure across latitudinal ranges but also introduced rare, geographically restricted divergent haplotypes^25,26^. More recently, investigation of the *C. briggsae* evolutionary history using a population genomics study of 37 wild whole-genome sequences^27^ implicated a near-simultaneous split of at least six distinct “lineages” (which we will call relatedness groups in this study) approximately 200,000 generations ago preceded by a more ancient split from all other relatedness groups approximately two million generations ago. Importantly, recent efforts have generated a vastly improved *C. briggsae* reference genome using Oxford Nanopore sequencing and provided updated gene models from high-quality transcriptome data and extensive manual curation^28,29^. These advances have narrowed the genome resource gap between *C. elegans* and *C. briggsae*, priming a comprehensive analysis of *C. briggsae* collections to enable large-scale comparative analyses of *Caenorhabditis* species.

In this study, we sequenced the whole genomes of over 2,000 wild *C. briggsae* strains collected from all over the world by the *Caenorhabditis* research community, massively expanding previous *C. briggsae* population studies. To characterize natural variation across the species, we leveraged updated genomic resources to discover single-nucleotide and insertion-deletion variants to reveal extensive genetic and geographic differentiation, allowing us to classify genetically distinct strains into twelve relatedness groups that likely represent independently evolving populations. Notably, strains with near identical whole-genome haplotypes have been isolated across different continents, suggesting that *C. briggsae* has undergone recent human-assisted dispersal and expansion. The distribution of polymorphism across *C. briggsae* genomes mirrors that of *C. elegans* genomes, such that chromosomal arm domains with low gene density and high-recombination display greater diversity relative to gene-dense, low-recombination chromosomal center domains. Our investigation of the patterns of genetic variation also revealed widespread hyper-divergent regions (HDRs) across 36% of the reference genome. Much like *C. elegans*, these HDRs display remarkable gene content variation and are enriched for genes associated with environmental responses. We uncovered remarkable differences in the frequencies of polymorphism across strains from different relatedness groups associated with HDRs. Comparison of shared polymorphism between pairs of *C. briggsae* relatedness groups revealed clusters of tightly linked alleles that do not appear to be recently introgressed from its closest outbreeding relative *Caenorhabditis nigoni*, but rather share an ancestral origin and have been maintained by long-term balancing selection. Overall, the updated genomic resources, catalog of genetic diversity, population structure analysis, and characterization of HDRs for this large collection of *C. briggsae* strains will improve power to detect associations between genotype and phenotype and to inform comparative and intra-specific models of evolution.

## Results

### *C. briggsae* genome-wide variation reveals both cosmopolitan and localized phylogeographic structure

*C. briggsae* has a wide global distribution with samples collected across six continents and many oceanic islands (Fig. 1a-b, Table S1). These samples come from bacteria-laden substrates of rotting vegetal materials. We compared whole-genome sequence data from 2,018 wild *C. briggsae* strains to understand patterns and levels of genetic variation (see Methods, Table S2). Because of self-fertilization, wild *C. briggsae* strains are naturally inbred and essentially homozygous at all loci, so we classified the 2,018 strains into 715 distinct genome-wide haplotypes (referred to as isotypes) (Fig. S1, Table S1, Table S2). The majority of isotypes are composed of a single strain (n=457) or multiple strains isolated less than one kilometer away from each other (n=173). Two isotypes comprise multiple strains where all but one strain lack information of their geographic site of isolation. Additionally, 66 isotypes are composed of strains isolated in the same geographic region between 1 km and 2000 km from each other (*e.g.*, different Hawaiian Islands or distant parts of China). By contrast, 17 isotypes comprised strains sampled from over 2,000 km away from each other (referred to as cosmopolitan isotypes), spanning multiple geographic regions and even different continents (Fig. S2a). The majority of the cosmopolitan isotypes do not appear to be closely related to one another (Table S2, Fig. S2b), suggesting that their geographic spread derives from multiple independent dispersal origins. We further investigated the genotype similarity and fine-scale geographic distribution of the largest cosmopolitan isotype (NIC174). This isotype includes 260 strains from Europe, 16 strains from North America, two strains from Asia, and one strain from each Africa, Australia, and South America, as well as one strain of unknown sampling location. Within the NIC174 isotype, we compared the geographic distribution of strains against genotype identities at all segregating SNV sites and found that strains from the same locality often display increased genotype similarity (Figure S3). This result suggests long-distance dispersal followed by recent local divergence. We applied a molecular clock at four-fold degenerate sites and assumed a *C. elegans* mutation rate^30^ to estimate that approximately 6,000 generations separate the most genetically dissimilar strains in this isotype. Assuming a minimum of 10 generations per year, we estimate that this isotype diverged less than 600 years ago. This isotype is consistent with the rapid dispersal and establishment of a single genome-wide haplotype likely because of human activity in recent centuries, suggesting an unprecedented sweep even more extensive than chromosomal-level selective sweeps found in *C. elegans*^16,18^.

**Figure 1:**
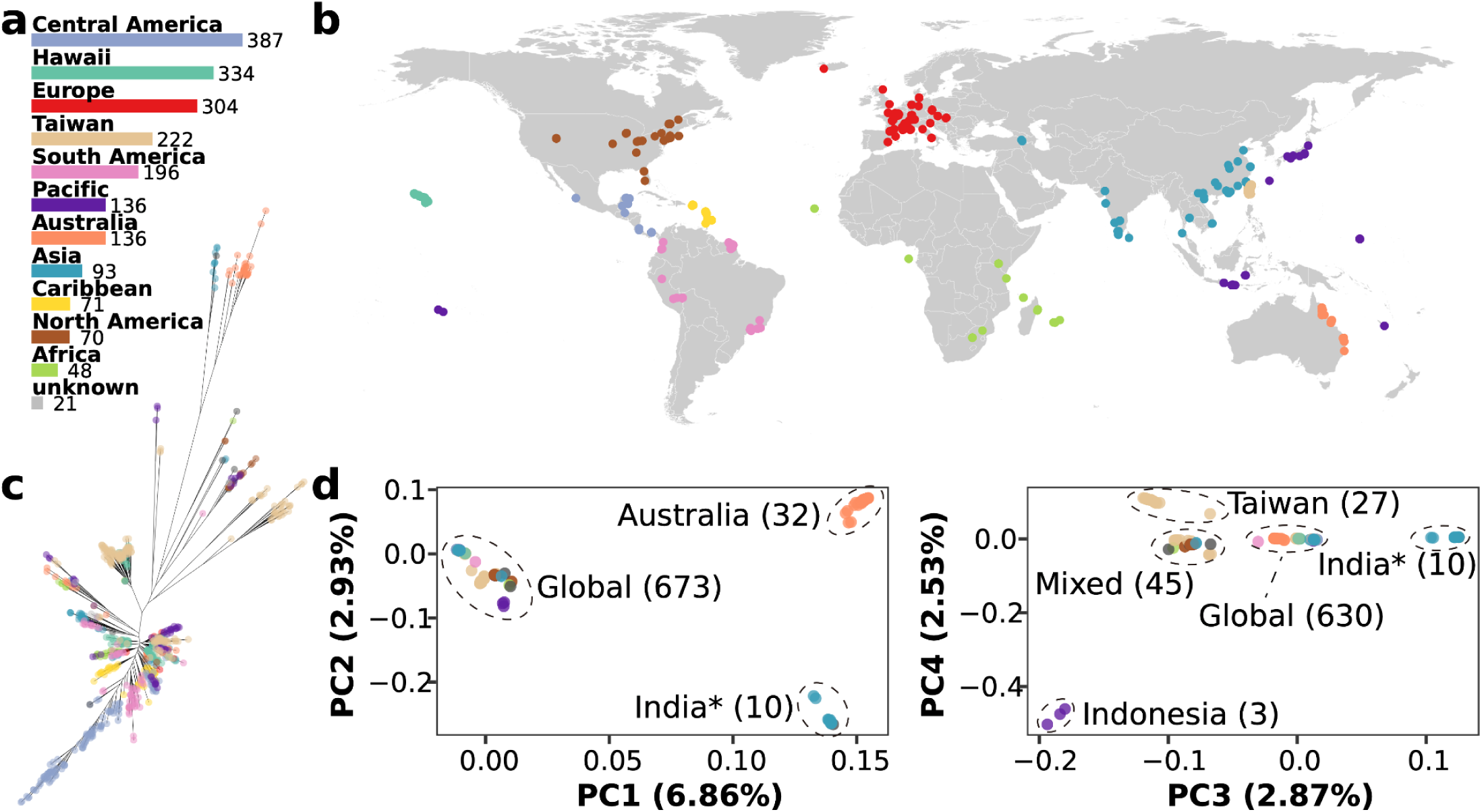
Global sampling efforts define 715 genetically distinct strains. **a,** Total number of strains in each geographic region. **b,** Global sampling map. **c,** Unrooted equal angle maximum likelihood tree of 715 *C. briggsae* isotype reference strains. The tree was generated from LD-pruned variants with r^2^ value less than 0.9 with the GTR+F+ASC+R10 maximum-likelihood substitution model (see Methods). **d,** Principal Component Analysis (PCA) of global *C. briggsae* isotype reference strains. Colors in each panel represent the sampling geographic region of each strain. The ellipses highlight differentiated isotypes. The numbers in parentheses represent the count of isotypes within each cluster, and the asterisk represents a cosmopolitan isotype (JU2801). JU2801 comprises two strains, JU2801 and JU3199, collected from Europe and Asia, respectively. The colors of the points in panels b-d correspond to their geographic regions as in panel a, except that cosmopolitan isotypes in panels c and d are shown in gray.

Across the genomes of 715 isotype reference strains, we identified approximately 5.65 million single-nucleotide variants (SNVs) and 876,200 short (<50 base pairs (bp)) insertion-deletion (indel) variants (see Data Availability, Materials and Methods). After pruning variants in strong linkage disequilibrium (r^2^ > 0.9, see Methods), we used biallelic SNVs to perform principal component analysis (PCA) (Fig. 1c-d). Nearly all isotypes fall within a single cluster in principal component (PC) space, separated from the minority of isotypes that define discrete clusters and also come from narrow geographic regions (Fig. 1d). Four PCs distinguish a cluster of 32 isotypes from the east coast of Australia, a cluster of nine isotypes from the south of India accompanied by one cosmopolitan isotype, a cluster of three isotypes from Indonesia, a cluster of 27 isotypes from central and northern Taiwan, and a Mixed cluster of 45 isotypes from various regions (Asia, Africa, Taiwan, North America, and cosmopolitan) (Fig. 1d, Fig. S4, Table S3). Chromosome-specific PCs largely recapitulate the patterns observed genome-wide (Fig. S5). We performed iterative outlier removal followed by PCA to further dissect genetic differentiation in the Global population cluster (Fig. S6) and found few distinct clusters, leading us to pursue additional approaches to explore *C. briggsae* population genetic differentiation.

We estimated mean pairwise genetic similarity by calculating the proportion of identical alleles across genome-wide SNVs between isotypes and clustered all isotypes into twelve relatedness groups (Fig. 2). Four of these twelve groups represent the geographically restricted PC clusters from Australia (referred to as Australia Divergent or AD), India (referred to as Kerala Divergent or KD), Taiwan (referred to as Taiwan Divergent 1, or TD1), and Indonesia (referred to as Indonesia Divergent or ID). We defined eight remaining relatedness groups that showed a mean within-group genetic similarity of >98% but <93% between-group genetic similarity (Fig. S7). Six relatedness groups represent subdivision of isotypes from the Mixed PC cluster and the remaining two represent subdivision of the Global PC cluster. A relatedness group of 28 isotypes isolated in locations with temperate latitudes in Asia, Europe, North America, and the Pacific was named after the “Temperate lineage” defined in previous studies^24–27^. Another relatedness group, named Nairobi-Wisconsin Divergent (NWD), includes one isotype from Africa (Nairobi), two isotypes from USA (Wisconsin), and one cosmopolitan isotype. Two groups of seven and three isotypes from Taiwan were named Taiwan Divergent 2 and 3 (or TD2 and TD3), respectively. Another group of 92 isotypes, which we define as the Taiwan-Hawaii (TH) relatedness group, includes 73 isotypes from Taiwan and eight isotypes from Hawaii as well as one isotype from each South America, Central America, and the Caribbean, six cosmopolitan isotypes, and two isotypes from other locations in Asia. The final relatedness group of 502 isotypes found across every sampled geographic region represents the circum-global “Tropical lineage” described in previous studies^25–27^. Overall, these analyses added new isotypes to previously defined relatedness groups (Tropical, Temperate, Hubei, Quebec, Kerala Divergent, and Taiwan Divergent 1) and allowed us to define many newly sampled groups (Australia Divergent, Taiwan Divergent 2 and 3, Indonesia Divergent, and Nairobi-Wisconsin Divergent). Despite extensive sampling efforts from the nematode research community, most relatedness groups remain rare and are geographically restricted, with the Tropical group dominating most sampling sites, overshadowed only by the Temperate group in colder climates.

**Figure 2:**
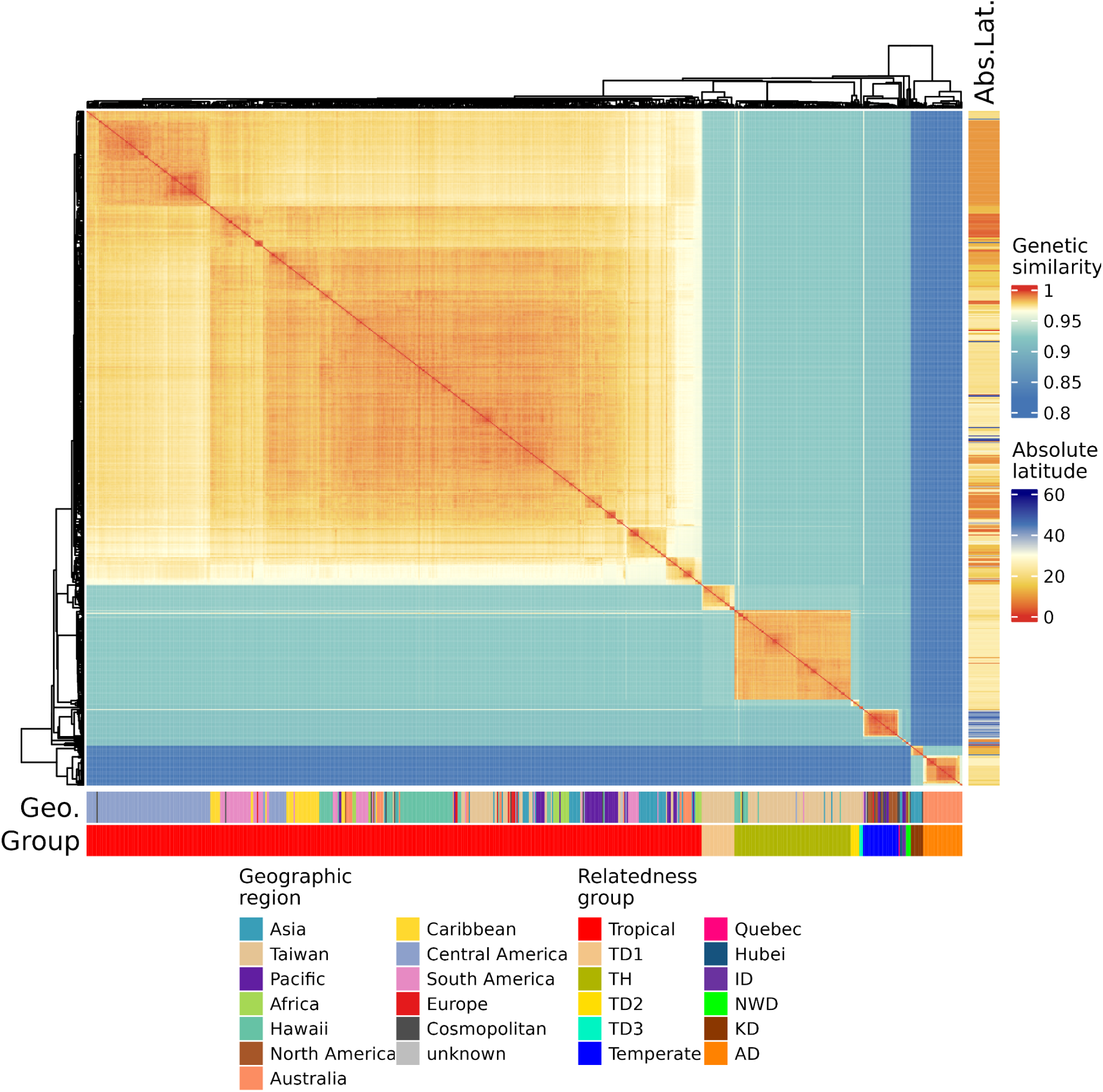
Pairwise genetic similarity defines *C. briggsae* relatedness groups. Heatmap showing pairwise genetic similarity between all 715 isotype reference strains. Pairwise genetic similarity was estimated by the proportion of identical alleles across all identified SNVs among the 715 reference strains. The genetic similarity color gradient has breaks specified at the 25th, 50th, and 75th percentile of the distribution of all pairwise estimates. The right band shows the absolute latitude of the isolation site of each isotype representative strain. The bottom bands show the geographic region of the isolation site of each isotype representative strain and the assigned relatedness group.

To investigate shared ancestry between relatedness groups, we performed admixture analysis^31^. We minimized cross-validation (CV) error by comparing CV estimates for 10 iterations at K assumed subpopulations (from K=2 to 30) and found support for at least 22 subpopulations (see Methods, Fig. S8-9). At K=22, non-admixed individuals from each subpopulation were assigned uniquely to one relatedness group, with certain relatedness groups containing several subpopulations (Fig. S10). Across ten iterations of K=22, we observed that relatedness groups with low sample sizes were sometimes assigned to the same subpopulation, although relatedness groups with many isotypes remained discrete (Fig. S11). We find admixture among Tropical relatedness group subpopulations as well as admixture of Tropical with either Temperate or TH relatedness groups (Fig. S10), suggesting that these three groups share more recent ancestry by potential microhabitat sharing that has enabled cross-breeding. By contrast, most other relatedness groups likely have been evolving independently.

Our results support a history of genetic diversification in Asia and the Pacific, although the geographic origin for *C. briggsae* remains unknown. Previous divergence time estimation dates the oldest split among *C. briggsae* relatedness groups between Kerala (KD) and all other groups^27^. The current study uncovered an additional relatedness group from Australia (AD) most closely related to KD but similarly diverged from all other groups, which offers an alternative explanation to a single ancient split in the species evolutionary history (Fig. 2). Because the updated analyses propose a Pacific geographic origin alternative to South India, it is also possible that a third unidentified site proximate to the Pacific and/or Indian Oceans could be the true origin. It is also possible that a relatedness group that could precede the most ancient detected split in this species has not been captured among the current sampled strains or is now extinct. The related outbreeding species *C. nigoni* is circum-tropical so the geographic origin of the most recent common ancestor is also unknown (Table S4) and offers no further insights into the geographic origins of *C. briggsae*.

### Genetic diversity is concentrated in punctuated hyper-divergent regions

Population genetics theory predicts self-fertilizing species exhibit lower population-wide genetic diversity than their outbreeding relatives, because frequent self-fertilization increases homozygosity and lowers the effectiveness of recombination^32–34^. As expected, our global sample of *C. briggsae* exhibits more than four-fold lower average pairwise nucleotide diversity (π) than *Caenorhabditis remanei*^20^ isolated from one location (Fig. 3, Table S5). Both π and the population mutation rate metric of polymorphism (Watterson’s θ ^35^, θ_W_) for *C. briggsae* are higher than those same metrics for *C. elegans* (Table S5). We found a 1.6-fold higher mean π and a 1.5-fold higher mean θ_W_ in arm domains relative to center domains on autosomes (Table S6), consistent with previous analyses of self-fertilizing *Caenorhabditis* nematodes^13,27^. This intra-chromosomal difference is attributable to pervasive linkage disequilibrium (LD) caused by self-fertilization, which intensifies the effects of linked selection in chromosome regions with lower recombination and higher gene density as in the chromosomal center domains^34,36^. This pattern is less pronounced on the X chromosome, where only a 1.2-fold higher mean π and 1.1-fold higher mean θ_W_ were measured in arm domains relative to the center domain. At a finer scale, π displays local peaks of nucleotide diversity within relatedness groups (example of Tropical group shown in Fig. S12), which defies the theoretical reduction of genetic diversity caused by self-fertilization. The discovery of hyper-divergent regions (HDRs) in other self-fertilizing *Caenorhabditis* nematodes, such as *C. elegans* and *C. tropicalis*, provides an attractive explanation for this observation^18,37^.

**Figure 3:**
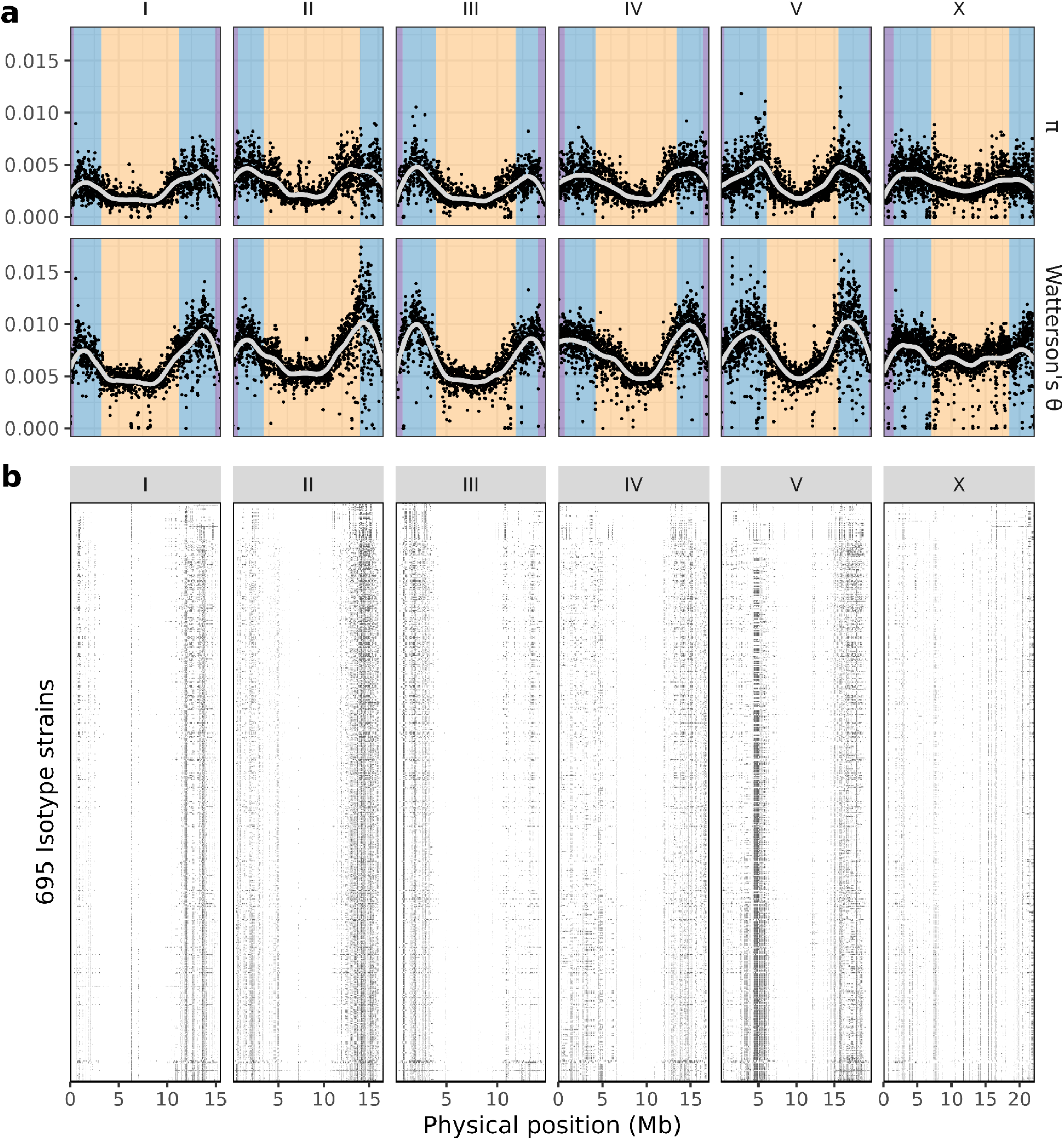
Genetic diversity is concentrated in punctuated hyper-divergent regions. Each vertical facet separates *C. briggsae* chromosomes with physical coordinates of the QX1410 reference genome on the x-axis. **a,** Plots of π and Watterson’s θ along genomic positions on each of the six chromosomes. Each point represents a 10 kb window estimate, with weighted locally estimated scatterplot smoothing (LOESS) curves where each local regression used 30% of the data. Colored backgrounds in nucleotide diversity panels delineate the chromosomal domain boundaries (purple, tip; blue, arm; yellow, center). **b,** Each row represents one of 695 isotype reference strains, with HDRs marked as black boxes. From a total of 715 isotype reference strains, 20 were excluded because they were used as references for each relatedness group, belonged to relatedness groups that lacked a reference genome, or belonged to relatedness groups that comprised fewer than three isotypes.

To identify and characterize HDRs in *C. briggsae*, we employed methods previously used in *C. elegans* with several modifications (see Methods). In summary, we used multiple reference genomes, one for each relatedness group with genomic coordinates lifted over onto a single reference coordinate system and discovered 1,010 non-overlapping HDRs with an average size of 63 kb, ranging from 5 kb to 3.7 Mb (see Methods; Fig. 3b, Fig. S13, Table S7-8). These HDRs covered approximately 60% of the QX1410 reference genome. When assessing relatedness groups individually, we find an average of 750 non-overlapping HDRs per relatedness group, ranging from a total of 402 (KD group) to 1,349 (Tropical group). Group-specific HDRs range from 5 kb to 1.3 Mb in size and cover from 16% to 38% of the reference genome, comparable to *C. elegans* HDRs (9 kb to 1.35 Mb in size, covering approximately 20% of the N2 genome^18^). The Tropical relatedness group has more HDRs compared to all other groups (Fig. S14, Table S8), because it comprises the most isotypes (501) from across all sampled geographic regions with a higher likelihood to identify rare hyper-divergent haplotypes.

As expected, HDRs are enriched for SNVs. For example, HDRs found in individual isotypes from the Tropical relatedness group contain up to 39% (mean = 31%) of the total alternative alleles across all SNVs, despite HDRs covering just 11% or less (mean = 6%) of the reference genome (Fig. S15, Table S8). Even in the relatedness groups with fewer HDRs than the Tropical group, HDRs in individual isotypes cover on average 5% to 11% of the relatedness group reference genome and contain on average 37% to 55% of the total alternative alleles across all SNVs called relative to their respective relatedness group reference genomes (Table S8). These results are consistent with HDRs in *C. elegans*, where despite covering less than 2% of the genome in individual isotypes they contribute more than 20% of the total alternative alleles across all SNVs on average. The levels of genetic variation across *C. briggsae* HDRs are likely severely underestimated considering that these regions contain many novel genes, copy-number variation, or genes that are highly divergent compared to the reference genome (Fig. S16-17), much like *C. elegans*^18^. The highly variable gene content offers clues to the possible functional roles for HDRs in *C. briggsae*.

Previous gene set enrichment analysis of *C. elegans* HDRs has shown overrepresentation of gene families involved in environmental sensing and pathogen responses, including C-type lectins, G-protein coupled receptors, nuclear hormone receptors, and E3 ubiquitin ligases (*e.g.*, F-box genes)^18^. A similar analysis in *C. briggsae* is more difficult because many of the reference genes lack gene ontology (GO) terms. To address this issue, we annotated functional protein domains and their associated GO identifiers using InterProScan^38^ (see Methods). This analysis identified functional protein domain annotations for approximately 69% (15,289 / 22,196) of all reference genes (Table S9) and approximately 53% (11,756 / 22,196) had GO identifiers. Because HDRs are primarily located on chromosome arm domains, we performed an enrichment test for genes in or outside of HDRs in these chromosomal regions (Fig. 4). We find that genes in HDRs on chromosomal arms are significantly enriched for functional protein domains associated with environmental and pathogenic responses, such as G-protein coupled receptors (60/63 genes, class h domain), superfamily domain nuclear hormone receptors (115/137 genes) or orphan *C. elegans*-like nuclear hormone receptors (58/60 genes), and E3 ubiquitin ligases (*e.g.*, F-box A domains, 66/73 genes; BTB and MATH domains, 52/57 genes; SKP1/BTB/POZ superfamily domains, 85/107 genes). Among enriched GO terms for the same gene sets, we find innate immune response genes like C-type lectins and F-box-containing genes (121/144 genes, *e.g.*, orthologs of *C. elegans clec-85*, *clec-209*, *clec-218*, *fbxa-141*, and *Y60A3A.8*) (Fig. 4b). Additionally, we find enrichment for xenobiotic metabolism (28/35 genes, Fig. 4b) and monooxygenase activity (39/50 genes, Fig. 4c) involved in xenobiotic metabolism, where many of the genes in both terms were identified as orthologs of *C. elegans* cytochrome P450 families such as *cyp-32*, *cyp-33*, *cyp-34*, and *cyp-35* genes. These results suggest that *C. briggsae* HDRs are enriched for environmental response genes likely involved in local adaptation.

**Figure 4.**
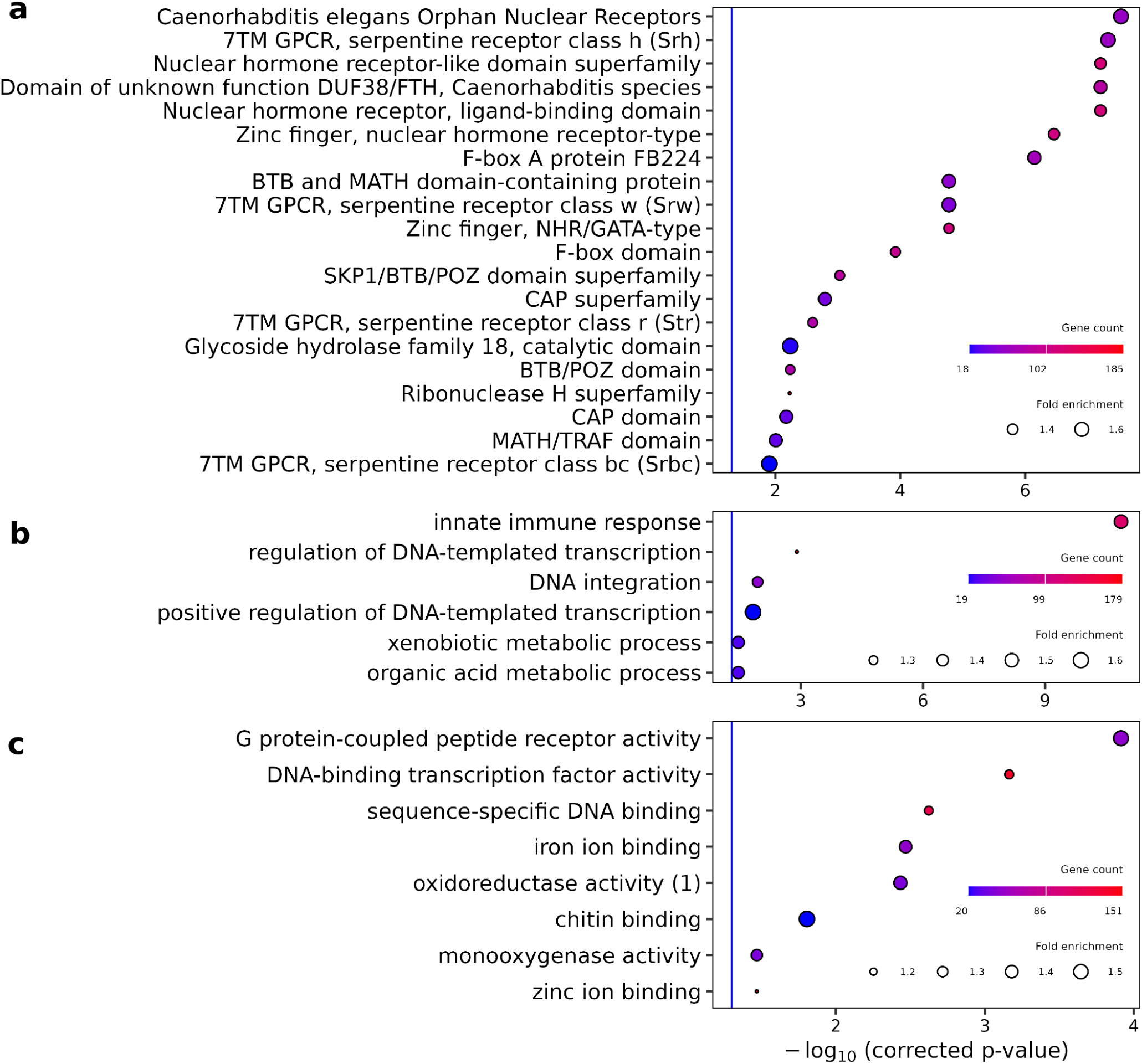
*C. briggsae* hyper-divergent regions are enriched for environmental response genes. (**a**) Significantly enriched InterProScan functional protein domains in genes found in HDR chromosome arms. Each protein domain annotation on the y-axis showed a false discovery rate-corrected p-value (q-value) of lower than 0.05 (negative log-scaled adjusted p-values shown on the x-axis) biological process Significant biological process (**b**) and molecular function (**c**) gene ontology terms from gene set enrichment analysis. Each term present in the y-axis showed a Benjamini-Hochberg adjusted p-value lower than 0.05 (negative log-scaled adjusted p-values shown in the x-axis). In each row, the circle varies in size based on the fold-enrichment associated with that term, and in shading based on the number of genes in chromosome arm HDRs associated with that term. “Oxidoreductase activity (1)” refers to the gene set term “oxidoreductase activity, acting on paired donors, with incorporation or reduction of molecular oxygen”. We performed the same tests for autosomal arm regions that were not classified as hyper-divergent and found no enriched protein domains, and found a few, non-environment related gene ontology terms. All enrichment tests shown use arm domain genes as a background gene set.

Overall, these results reveal that a substantial percentage of variation (on average, 31% to 56% across relatedness groups) is found in punctuated HDRs that cover a relatively small span of *C. briggsae* genomes (on average, up to 11% of individual isotype genomes). Reference genes within HDRs are associated with a multitude of environmental responses, such as olfaction and sensing, transcriptional regulation, and xenobiotic and pathogen stress response. Across genome assemblies of isotypes within and across relatedness groups, we observe drastic gene content variation in HDRs, including novel genes lacking orthologs in the reference genome. For this reason, a pangenome gene set analysis is required to more comprehensively investigate the composition and functions of genes found in HDRs.

### Introgression rarely explains hyper-divergent regions

Multiple mechanisms have been proposed to explain how HDRs are generated (*e.g*., by introgression) and maintained (*e.g*., by balancing selection)^18,39^. To explore the possibility that HDRs could originate from introgression between *C. briggsae* and its outbreeding sister species *C. nigoni*, we needed to search for evidence of introgression or the sharing of alleles in orthologous genes between the two species. *C. nigoni* population resources are not developed so we sequenced the genomes of eleven *C. nigoni* inbred lines using PacBio HiFi technology and five additional *C. nigoni* isofemale lines using Oxford Nanopore Technologies (ONT), assembled highly contiguous genomes, and predicted protein-coding genes (see Methods, Table S10). To compare *C. nigoni* genes to genes found in *C. briggsae* HDRs, we needed to characterize the gene content and variation across diverse *C. briggsae* isotypes, so we performed similar analyses using PacBio HiFi technology on 20 Tropical *C. briggsae* isotype reference strains. We then predicted gene models and identified orthologous groups using OrthoFinder^40^ (File S1). We identified a total of 32,423 orthologous groups, with 22,422 shared between at least one member from both species and 2,321 and 7,680 were unique to either *C. briggsae* or *C. nigoni*, respectively. From the shared orthologous groups, 7,530 represent single-copy orthologs from which 5,205 were shared across all *C. briggsae* and *C. nigoni* individuals. To enable comparisons of genes found in both species and investigate possible introgression events, we constructed gene trees using protein sequences of genes from each orthologous group (see Methods). We postulated that if HDRs originated by introgression, then gene sequences from *C. nigoni* inbred lines would be more similar to gene sequences from a group of *C. briggsae* that carry either the reference or a hyper-divergent haplotype such that possible introgression events would lack clear clustering of haplotypes by species. We found 2,449 gene trees that represented potential cases of introgression, identified by the absence of two distinct monophyletic groups (one for each species, Fig S18). We then removed any gene trees where the gene is highly conserved across all *C. briggsae* and *C. nigoni* samples or the trees lacked a reference gene (see Methods). Trees that include gene duplications or gene families with high sequence similarity that could have evolved before or after speciation pose complications when trying to assess haplotypes that cluster by species, so we focused our analysis on the remaining 850 gene trees corresponding to single-copy orthologs. Among these trees, 60 trees included reference genes that are located within HDRs in at least one *C. briggsae* isotype strain (Table S11). These 60 gene trees include subsets of *C. briggsae* gene sequences with increased sequence similarity to *C. nigoni* sequences that could be a product of introgression (File S2). However, we found only 10 gene trees where either the hyper-divergent or reference *C. briggsae* haplotype displayed increased sequence similarity to *C. nigoni* than to the other *C. briggsae* haplotype (File S2; OG0015043, OG0013045, OG0019613, OG0013391, OG0022648, OG0024532, OG0022300, OG0016700, OG0019907, OG0018191). These results indicate that, although some divergent gene sequences in *C. briggsae* HDRs could plausibly be explained by introgression from *C. nigoni*, such instances are extremely rare and cannot account for the vast majority of HDRs that pervade the *C. briggsae* genome, inconsistent with introgression as a primary source of HDRs.

### Hyper-divergent regions are likely maintained by balancing selection

The discovery of *C. elegans* HDRs led to the hypothesis that these regions contain adaptive alleles maintained by long-term balancing selection^18^, likely in populations subject to spatial or temporal environmental variation. Previous analyses used Tajima’s *D*^41^ to quantify the allele frequency skews that can suggest different signatures of selection^18^. We wanted to perform similar analyses of *C. briggsae* HDRs, but the interpretations of Tajima’s *D* are biased by the combined influences of demographic size change, direct and linked selection, population structure, and unequal sampling of subpopulations^42,43^, given the high degree of genetic variation between relatedness groups (Fig. 2). To limit these biases, we analyzed Tajima’s *D* separately for each relatedness group containing more than ten isotype reference strains (AD, KD, TH, TD1, Temperate, Tropical). Consistent with global *C. briggsae* estimates, we found lower π and θ_W_ in chromosomal centers relative to arms but to varying degrees (Fig. 5, Fig. S19-20, Table S12). Although mean estimates of *D* across chromosomal arms are slightly negative, we find abundant genomic peaks of positive *D* in every relatedness group comparison (Fig. S21). These peaks are most often found in HDRs (Fig. 5a), suggesting the presence of intermediate-frequency alleles that could be products of balancing selection. We were curious whether the alleles found in HDRs within relatedness groups significantly diverged between relatedness groups. Estimates of absolute divergence (D_xy_) enabled these comparisons where we found that HDRs, which were previously shown to carry increased π and θ_W_ (Fig. 3), display lower to no difference in absolute divergence between relatedness groups relative to non-HDRs, most evidently in arm domains (Fig. 5b, Fig. S22). The largest differences in D_xy_ between HDRs and non-HDRs are observed in more extreme pairwise comparisons (Tropical to AD or KD). These results are consistent with those regions that encode increased diversity carrying distinct allelic haplotypes that have been maintained since before the split of the relatedness groups. HDRs likely arose by the establishment of distinct allele combinations present in a hyper-diverse ancestor, with selection and self-fertilization driving allele fixation and extinction (purifying) or alternative allele maintenance (balancing) across *C. briggsae* relatedness groups.

**Figure 5.**
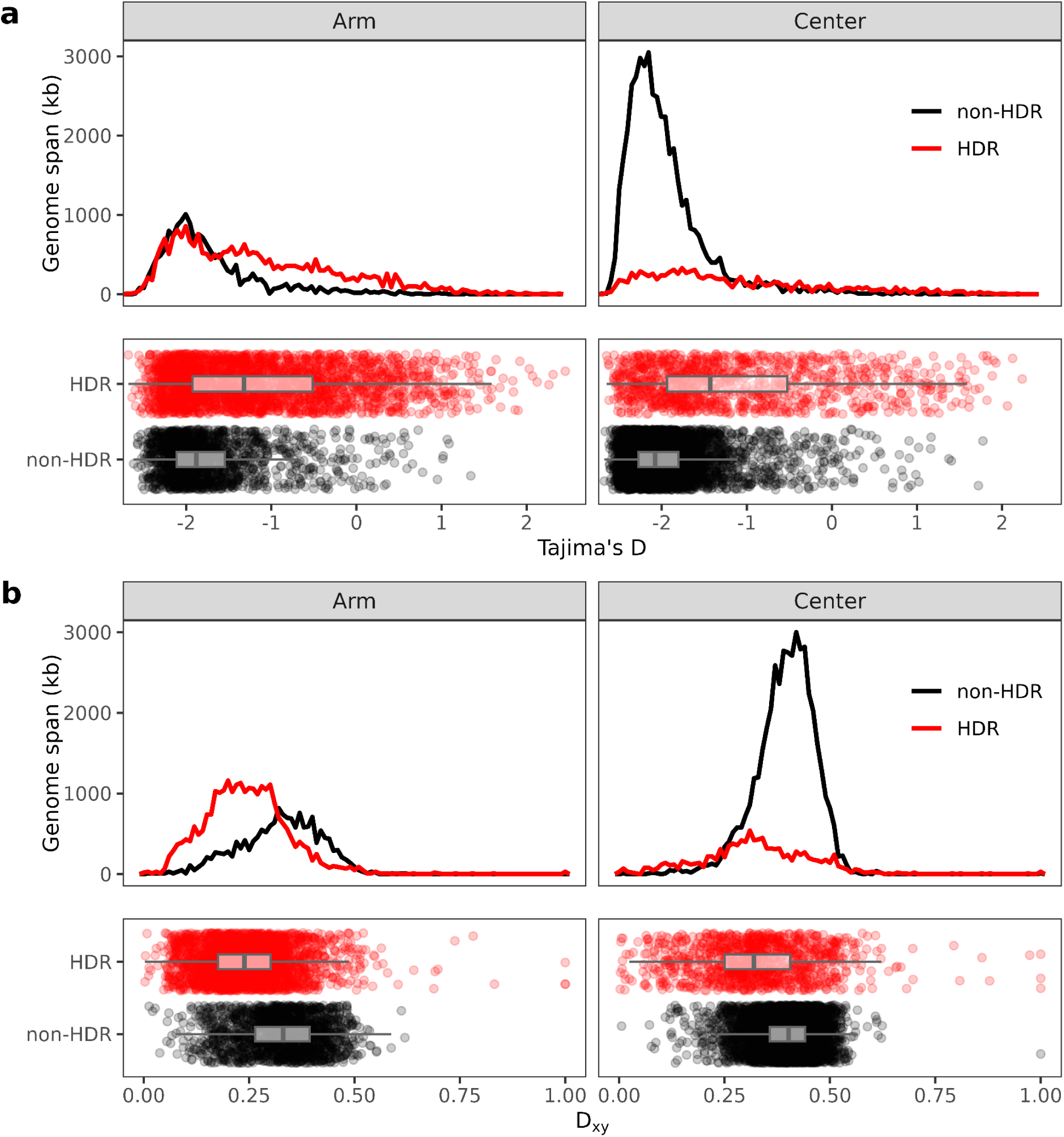
Skew towards intermediate frequency alleles and reduced absolute divergence in *C. briggsae* hyper-divergent regions. **a,** Tajima’s *D* estimates across 10 kb genome segments among Tropical group isotypes comparing hyper-divergent and non-HDRs. Center and arm estimates are shown as separate vertical facets, with the top panel of each chromosomal domain showing the genome span in the y-axis against each value of Tajima’s *D* on the x-axis (binned in increments of 0.05), and the bottom panel showing a boxplot comparing hyper-divergent and non-hyper-divergent genome segments. **b,** D_xy_ estimates across 10 kb genome segments between Tropical and KD groups comparing hyper-divergent and non-HDRs. Each point represents the average number of pairwise differences for a unique 10 kb genomic interval. Again, center and arm estimates are shown as separate vertical facets, with the top panel of each chromosomal domain showing the genome span in the y-axis against each value of D_xy_ on the x-axis (binned in increments of 0.01), and the bottom panel showing a boxplot comparing hyper-divergent and non-hyper-divergent genome segments.

## Discussion

The massive expansion of sequenced *Caenorhabditis briggsae* genomes from around the world, enabled by the efforts of the nematode research community, allows for more comprehensive characterization of genomic diversity throughout this species. Our analyses revealed millions of single-nucleotide and thousands of insertion-deletion variants across 715 distinct genome-wide haplotypes (isotypes). These isotypes differ widely in their geographic distribution and relatedness. We discovered cosmopolitan isotypes comprising large strain cohorts isolated in multiple continents, underscoring the influence of recent human activity in the dispersal of this species primarily through Europe and North America. Cosmopolitan isotypes contrast sharply with the majority of endemic isotypes showing deep genetic divergence. We defined 12 relatedness groups that capture the major axes of genetic divergence, from rare local groups to large globally distributed groups. Previous population genomic studies used highly diverged strains from the south of India (Kerala) to suggest that the earliest diversification of *C. briggsae* could have occurred near the Indian Ocean^27^, where its outbreeding sister species *C. nigoni* also is found. The discovery of an Australian relatedness group most closely related to the Kerala relatedness group and similarly diverged from all other groups widens the species origin to the Asia-Pacific region. These results are consistent with the potential geographic origins of several lineages of the Elegans supergroup^44^ and emphasize the importance of continued sampling to refine estimates of divergence time and to trace the biogeographic origins of this species.

Our analysis also corroborates chromosomal profiles of genetic diversity observed across self-fertilizing *Caenorhabditis* nematodes where most genetic variation is concentrated within or near chromosomal arm domains. This uneven distribution reflects the combined effects of linked selection and low recombination in gene-dense central regions. Self-fertilization further exacerbates these effects, producing long tracts of linkage disequilibrium and purging genetic variation when selection acts on adaptive or deleterious mutations^13,16,24,27,28,37,45,46^. At a finer scale, nucleotide diversity is condensed in punctuated genomic regions that resemble the recently discovered HDRs in *C. elegans*^18^. Much like *C. elegans*, HDRs in *C. briggsae* encode haplotypes with remarkable gene content variation and encompass large proportions (approximately 30-50%) of genetic variation within *C. briggsae* relatedness groups. Additionally, *C. briggsae* HDRs also show enrichment for environmental response genes, indicative of the potential role in adaptation for self-fertilization *Caenorhabditis* species. The characterization of HDRs in *C. briggsae*, accompanied by the contiguous genomes and protein-coding gene models that we have generated for diverse *C. briggsae* wild strains provides new avenues to explore natural variation in this species. Although we provide examples where HDRs implicate novel genes and copy-number variants that yield structurally distinct haplotypes, it remains unknown to what extent different classes of structural variants pervade these regions. Although alignment-based approaches might not be suitable to effectively investigate haplotype diversity in HDRs, the advent of pangenome-based analyses offer a potential solution. Pangenome graphs targeted to HDRs, perhaps associated with phenotypic differences as observed in *C. elegans*^47–50^, can help distinguish variation hidden to alignment-based approaches, improve genotype-phenotype associations, and narrow to causal genes.

Importantly, HDRs also provide insights into the evolutionary trajectory of *C. briggsae*. Because *C. briggsae* and its sister species *C. nigoni* can hybridize under laboratory conditions^51^, we tested whether HDRs in *C. briggsae* might have originated through recurrent introgression from *C. nigoni*. We generated high-quality genome assemblies and gene models for *C. nigoni* lines and used our diverse cohort of *C. briggsae* wild strain genomes to identify single-copy orthologs and compare their protein sequences. Although we detected hundreds of potentially introgressed genes, only a small proportion occurred within HDRs, and even fewer display divergent haplotypes with increased sequence similarity to *C. nigoni* haplotypes. Therefore, we propose that genetic hyper-diversity like that found in other outbreeding *Caenorhabditis* species was once present in the outbreeding ancestor of present-day *C. briggsae* and has since been reduced to punctuated segments surrounding hyper-divergent haplotypes. The emergence of a self-fertilizing lineage leads to rapid reductions in heterozygosity, which exposes recessive alleles that can lead to reductions in fitness or lethality (inbreeding depression) and removal of these recessive deleterious alleles and linked neutral alleles, leading to overall reductions in genetic diversity across a population^32–34^. However, genetically diverse allelic haplotypes could be retained within a self-fertilizing lineage in genomic regions where recessive deleterious alleles are absent. These (hyper-)diverse genomic regions potentially persist over long periods of time in the self-fertilizing lineage because of balancing selection, where different adaptive haplotypes become fixed in distinct niches or fluctuate in frequency because of adaptive advantages to temporally varying environmental factors.

We showed that HDRs carry intermediate-frequency alleles within *C. briggsae* relatedness groups, a canonical signature of genomic regions under balancing selection. Additionally, the juxtaposition of increased within-group diversity and decreased absolute divergence observed between relatedness groups in HDRs suggest that these regions carry allelic haplotypes that have been maintained since relatedness groups split. Although the different relatedness groups likely diverged from a single self-fertilizing lineage, it remains unclear whether hyper-divergent haplotypes originated from selection acting on a common ancestral genetic background or distinct intercrossing events with multiple genetic backgrounds of the outbreeding progenitor of *C. briggsae*. HDRs offer an opportunity to study the evolutionary origins of the species, because these regions could be used to approximate the effective population size of the outbreeding ancestor of *C. briggsae*. Such approximations can be used under a simulation framework to set expectations surrounding different models of transition to self-fertilization, from a gradual increase in self-fertilization rate with ample intercrossing with the outbreeding progenitor to a large-effect self-fertilization origin with limited intercrossing.

Overall, these newly created strain resources for *C. briggsae* enable comparative genomics studies not possible in other animal species. Within the *Caenorhabditis* genus, several species have worldwide samples comprising thousands of wild isolates, whole-genome sequences, and a variety of chromosomal assemblies. Genome-wide association studies in *C. briggsae* can be performed using specific relatedness groups (*e.g.*, Tropical) to minimize the effects of population structure, thereby reducing false discovery of quantitative trait loci^52^. By comparing quantitative variation in traits shared between *C. briggsae* and *C. elegans*, the field could identify conserved loci not possible by studying a single species in isolation. These approaches are now possible and future high-throughput assays that take advantage of large strain sets from both species should be studied to make significant impacts on our understanding of evolutionary processes.

## Methods

### Strains

A total of 2,018 *C. briggsae* strains and 16 *C. nigoni* inbred lines were included in this study (Table S2, Table S4). *C. briggsae* strains and 11 *C. nigoni* inbred lines (ECA2852, ECA2857, JU1422, JU1418, JU1419, JU1420, NIC2143, NIC2150, NIC2152, VX151, VX153) were reared at 20°C using *Escherichia coli* bacteria (strain OP50) grown on modified nematode growth medium (NGMA)^53^ containing 1% agar and 0.7% agarose to prevent animals from burrowing. All *C. briggsae* isotype reference strains used in this study have been deposited at and are available upon request from the *Caenorhabditis* Natural Diversity Resource (CaeNDR)^54^. The remaining five *C. nigoni* isofemale lines (EG5268, JU2484, JU2617, YR106, ZF1220) for ONT sequencing were cultured at 25°C on NGM plates with twice the standard agar concentration to prevent nematode burrowing. The following strains were checked for cross compatibility with AF16 using a fluorescently labeled strain (QG2801): QG4064, QG4094, QG4095, QG4131, QG4208, QG4233, JU2801, JU3202, JU3203, and JU3206. All strains gave rise to viable progeny.

### Whole-genome sequencing

DNA extraction was performed following established protocols^17^. In brief, to extract DNA, we transferred nematodes from three 10 cm NGMA plates spotted with OP50 *E. coli* into a 15 ml conical tube by washing with 10 mL of M9. We then used gravity to settle animals on the bottom of the conical tube, removed the supernatant, and added 10 mL of fresh M9. We repeated this wash method three times to serially dilute the *E. coli* in the M9 and allow the animals time to purge ingested *E. coli*. Genomic DNA was isolated from 100 to 300 µl nematode pellets using the DNAEasy Blood and Tissue isolation kit cat# 69506 (cat# 69506, QIAGEN, Valencia, CA). The DNA concentration was determined for each sample with the Qubit dsDNA Broad Range Assay Kit cat# Q32850 (cat# Q32850 Invitrogen, Carlsbad, CA). All strains were used to construct short-read libraries using NEBNext® Ultra™ II FS DNA Library Prep (cat# E6177L). These libraries were sequenced on several Illumina NovaSeq platforms (paired-end 150 bp reads) at the Duke Center for Genomic and Computational Biology, Novogene, NUSeq, the New York University core facility, and the Johns Hopkins Genetics Resources Core Facility. The raw short-read sequencing data are available from the NCBI Sequence Read Archive (Project PRJNA1149159).

For PacBio HiFi sequencing, 13 *C. briggsae* samples (BRC20341, VX34, NIC1660, ECA2670, QG4132, ECA2666, BRC20492, JU3237, ECA2617, QX1796, TWN1824, NIC1529, JU3200) were each grown on 15 10 cm plates spotted with OP50 *E. coli*. The nematodes were washed off of the plates with M9 and allowed to settle by gravity. The pellets were washed three times using M9 and then frozen at −80°C. All of these pellets were processed (DNA extraction and library generation) and sequenced at the Cold Spring Harbor Laboratory Genome Center. For the remaining 46 *C. briggsae* (Table S10) and 11 *C. nigoni* samples (ECA2852, ECA2857, JU1422, JU1418, JU1419, JU1420, NIC2143, NIC2150, NIC2152, VX151, VX153), animals were grown on 4-5 10 cm NGMA plates spotted with OP50 *E. coli*. Just prior to starvation, the animals were washed off with cold M9 into a 50 ml conical and spun at 2000 rpm for eight minutes with a break of three. The supernatant was removed and the pellet was washed three times with cold M9 + 0.01% Tween. A final wash was performed with cold M9. The majority of the supernatant was removed and the pellet was transferred to a 1.7 ml microfuge tube using a glass Pasteur pipette. The animals were spun at 2000 rpm for 3 minutes, the supernatant was removed, the pellet was flash frozen, and then stored at −80°C. The pellet should weigh no more than 60 mg. To make HMW DNA, we used the Monarch HMW DNA Extraction Kit for Tissue (cat# T3060L, NEB, Ipswich, MA). The protocol was completed with the following modifications/specifics: Prior to using the pestle, we added 50 µl of the lysis buffer to the sample and added the remaining 600 µl after homogenization; for the lysis at 56°C after homogenization, we incubated in a Thermomixer for 15 minutes at 750 rpm and then at 550 rpm for 30 minutes or longer; use cold Protein Separation Solution; leave on ice for 10 minutes after the inversion step; use three Capture beads; to elute, we added 200 µl Elution Buffer, incubated for 5 minutes at 56°C and 300 rpm, another 50 µl of Elution Buffer was added prior to separating the DNA from the beads; the final solubilization was done at 37°C for 30 minutes at 300 rpm. Library generation and sequencing were carried out by Maryland Genomics at the University of Maryland or Duke Center for Genomic and Computational Biology. For ONT sequencing of five *C. nigoni* isofemale lines (EG5268, JU2484, JU2617, YR106, ZF1220), *C. nigoni* genomic DNA of each strain was extracted from at least six starved nematodes on 90 mm plates using the MasterPure Complete DNA and RNA Purification Kit (Biosearch Technologies). R10.4.1 flow cells (FLO-MIN114) and the ligation sequencing kit (SQK-LSK114) were then used to further process genomic DNA with 4 μg as the input for each sequencing run. Sequencing and basecalling were carried out on a GridION platform using MinKNOW software v5.9.18 with default parameters, and only high-quality reads were kept for genome assembly.

### Alignments and variant calling

Adapters and low-quality sequences were removed from raw reads using *fastp* v0.20.0 and default parameters^55^. Reads shorter than 20 bp were discarded after removed adapters and low-quality sequences. Subsequently, *C. briggsae* QX1410 (project: PRJNA784955)^28^ was used as reference to perform alignment with *Burrows-Wheeler Aligner, BWA* v0.7.17^56^. *Sambamba* was used to merge and index libraries of the same strain v0.7.0^57^, where duplicate sequencing reads were flagged with *Picard* v2.21.3^58^. Strains with less than 10x coverage were excluded from subsequent analyses.

Variant calling for each strain was conducted using *HaplotypeCaller* in GATK v4.1.4.0^59^. Then, the identified variants were aggregated and jointly recalled using *GenomicsDBImport* and *GenotypeGVCFs* in *GATK* v4.1.4.0^59^. The variants were further processed for polarizing heterozygous single nucleotide variants and filtering with the Andersen Lab pipeline *wi-gatk*^60–61^. Biallelic heterozygous SNVs were adjusted to homozygous reference (REF) or alternate (ALT) if there was substantial read support for this conversion. In this step, we only looked at SNVs in nucleic DNA. Specifically, the SNV was converted if the normalized Phred-scaled likelihoods (PL) met the following criteria, where smaller PL values indicate higher confidence. Any heterozygous SNVs that failed to meet these criteria were not changed in this step. The remaining SNVs were converted to homozygous ALT if PL-ALT/PL-REF <= 0.5 and PL-ALT <= 200; likewise, they were converted to homozygous REF if PL-REF/PL-ALT <= 0.5 and PL-REF <= 200. At the site-level, variants were subjected to several filters: ensuring variant quality (QUAL) > 30 (only three sites failed in this lenient filter); normalizing variant quality by read depth (QD) > 20; mitigating strand bias of ALT calls with strand odds ratio (SOR) < 5 and fisherstrand (FS) < 100; Limiting the fraction of samples with missing genotype to < 95%; and restricting fraction of samples with heterozygous genotype after heterozygous site polarization to < 10%. At the sample-level, filtration of variants were based on read depth (DP) > 5. Non-heterozygous sites were converted to missing (./.), except for heterozygous sites on mitochondria, which were not changed. After these procedures, invariant sites were removed. The generated population-wide variant call format (VCF) file was referred to as the hard-filtered VCF file.

### Isotype reference strain characterization

Strains were grouped into the highly related groups called isotypes using pairwise genetic similarity. First, genetic similarity was estimated from the ratio of identical alleles relative to the total non-missing SNV sites (total SNVs: 5,755,781) between each pair of strains computed using an in-house script with the hard-filtered VCF file as the input. Second, strains were then hierarchical agglomerative clustered by genetic similarity using complete linkage with a cutoff of 99.97% similarity, such that all pairwise similarities between strains within a given isotype group exceeded this cutoff. The threshold of 99.97% genetic similarity was selected based on the pairwise genetic similarity distribution, which aims to remove redundant (nearly-identical) genotypes from our analyses (Fig. S23). This method is available as a Nextflow pipeline (https://github.com/andersenlab/isotype-nf).

### Divergence time approximation in the NIC174 cosmopolitan isotype

Four-fold degenerate sites were annotated with the QX1410 reference genome and protein-coding gene annotation using *degenotate* v1.3 (https://github.com/harvardinformatics/degenotate). We then subsetted the hard-filtered VCF file to four-fold degenerate sites between the two least genetically similar strains in the NIC174 isotype (JU2584 and NIC2096). Assuming a *C. elegans* mutation rate (μ) of 2.3 x 10^-09^ per site per generation^30^ and using the proportion of identified four-fold degenerate site variants (k = 198/6690038) between the two least genetically similar strains in the NIC174 isotype, we applied a molecular clock under an infinite sites model^62,63^ to approximate a divergence time (T ≈ k/2μ) of 6,370 generations.

### Relatedness of isotype reference strains

Relatedness of the *C. briggsae* isotype reference strains were analysed based on the linkage disequilibrium (LD)-pruned VCF file, which was generated from the hard-filtered VCF file after the following variants-pruning steps: Genome-wide biallelic SNPs on all chromosomes were extracted and normalized by joining separate biallelic variants into multiallelic records using *bcftools norm -m +* in bcftools v1.14. Then the biallelic-only SNPs were extracted and LD pruned with PLINK v1.90b6.21 (with --*indep-pairwise 50 10 0.9*) to remove highly correlated markers^64,65^. Finally, after removal of missing sites from the markers, the loci corresponding to the remaining markers were extracted from the normalized VCF file to generate an LD-pruned VCF containing only biallelic variants without missing sites. Hereafter, this file is referred to as the “LD-pruned VCF file”. PCA was performed with smartpca v16000 executable from EIGENSOFT v7.2^66,67^. The LD-pruned marker files (.ped, .snp, and .ind) produced by PLINK were supplied as input to smartpca to calculate Eigenstrat values and Tracy-Widom statistics. A total of 219 PCs were required to explain 80% of the genetic variance, and 144 PCs contributed substantially to explaining genetic variation in that they exhibited eigenvalues >1.

### Species-wide tree construction

We used the *vcf2phylip.py* script^68^ to convert the LD-pruned VCF files into the PHYLIP^69^ format. Then, IQ-TREE2 v2.4.0^70^ was used to construct a tree with PHYLIP format file as input. The GTR+F+ASC+R10 substitution model which was selected as the maximum-likelihood model under automatic model selection based on the Bayesian information criterion (BIC). We limited our model search to extensions of the GTR substitution model. Node support was assessed using 1,000 ultra-fast bootstrap (UFBoot2) replicates. Trees were then visualized in R v4.3.2 with the ggtree v3.10.1 R package^71^. Pairwise genetic similarity among 715 global *C. briggsae* isotype reference strains were clustered and visualized with the ComplexHeatmap v2.18.0 R package^72^.

### Assignment of the cosmopolitan isotype reference strains

All the isotypes that contain any strains with pairwise geographic distances over 2000 km among sampling locations were defined as cosmopolitan isotypes. The geographic distances were calculated with the *geosphere* v1.5-18 R package. The Haversine formula was used to calculate the distance, which assumed the shape of Earth is a perfect sphere^73^.

### Repeat Masking

To mask the repeats, we identified *C. briggase* genome-wide repetitive elements using methods described previously^28^ with minor modifications. First, repetitive sequences were identified modeled *de novo* using RepeatModeler from RepeatMasker v2.0.6^74^. Transposable elements were then identified using TransposonPSI v1.0.0 (http://transposonpsi.sourceforge.net/). Long terminal repeat (LTR) retrotransposons were identified with LTRharvest and then annotated with LTRdigest from GenomeTools v1.6.5^75,76^, with HMM profiles from the Gypsy Database v2.0^77^ and Pfam v37.4 domains^78^. Repetitive sequences were then further filtered with the gt-select tool to remove sequences lacking conserved protein domains. We then obtained all Rhabditid repeats from RepBase^79^ and Dfam^80^. The generated repeat libraries were merged into a single redundant repeat library, clustered with VSEARCH v2.22.1^81^ from QIIME2 v2024.10^82^, and then classified with the RepeatClassifier tool from RepeatModeler. Finally, the unclassified repeats that had BLASTX v2.16.0 hits to *C. elegans* or *C. briggsae* proteins were removed using RepeatMasker to eliminate genuine gene sequences that had been misidentified as repeats.

### Population genomic statistics

Genome-wide nucleotide diversity (π), Watterson’s θ (θ_W_), and Tajima’s D were all calculated with scikit-allel v1.3.5^83^ per 10 kb window using the hard-filtered VCF file as the input. Only single-nucleotide variant positions were kept, repeat regions and sites with over 80% missing genotype data were excluded. To avoid bias caused by structural variation and indels that alter the number of callable sites when calculating π, θ_W_, and Tajima’s D using scikit-allel, we masked the inaccessible sites^84^. For π and θ_W_, inaccessible sites within each window were masked using the *is_accessible* parameter so that denominators reflected the number of callable bases. For Tajima’s D, genotypes at inaccessible sites were masked before calculation. D_xy_ between relatedness groups were calculated with Pixy v1.2.11.beta1^84^ per 10 kb window using the hard-filtered VCF file as the input. Accessibility to variable sites in the denominator is handled by default settings in Pixy^84^.

### Admixture analysis

The admixture analysis was performed using ADMIXTURE v1.3.0^31^ with the PED file as the input. This PED file contains the LD-pruned sites and was exported by PLINK when generating the LD-pruned VCF file. Ten independent times of ten-fold cross-validation (CV) were used to determine the optimized number of total populations where the curve of CV error first reached a minimization plateau. The total number of tested populations ranged from 2 to 30. Non-admixed representative strains were defined by the maximum subpopulation fraction over 99.9%.

### Long-read genome assembly and protein-coding gene prediction

Sequenced PacBio HiFi genomic reads were demultiplexed and adapters were removed by the sequencing cores using using *lima* (Duke: v2.9.0, CSHL: v2.2.0, UMD: v2.12.0 (https://lima.how). Demultiplexed and adapter-trimmed samples from the same strain sequenced from different facilities were merged using *samtools* v1.20^61^. PCR duplicates were removed using *pbmarkdup* v1.1.1 (https://github.com/PacificBiosciences/pbmarkdup). Primary assemblies were generated from deduplicated HiFi FASTAs using *hifiasm* v0.24.0-r702^85^ with the initial bloom filtered disabled (*-f0*, recommended for small genomes) and haplotig purging disabled (*-l0*, recommended for homozygous genomes). Genome assemblies generated from merged samples from the same strain were compared against assemblies generated from each individual sample, and the assembly with the highest N50 was kept. After assembly, contigs were taxonomically classified using DIAMOND 2.1.13^86^ with parameters *--faster* and an e-value of 1e-10 against the UniProt database (Release 2025_03)^87^. Contigs that were annotated as non-*Nematoda* were purged using BlobToolKit v4.4.5^88^ with *--param bestsumorder_phylum--Inv=Nematoda*. For assembly of ONT genomic reads, raw reads were first corrected using Canu v2.2^89^ with the parameter *-correct genomeSize=130m*. The corrected reads were assembled with Flye v2.9.3^90^ under the *-nano-raw* setting optimized for nanopore data. Misassemblies in the contigs were then corrected using the RagTag *--correct* module, which was executed with the ragtag.py correct command and MUMmer v4.0.0^91^ as the aligner. The alignment was performed with the NUCmer tool from the MUMmer package using the parameters *--mum --mincluster 100 --maxgap 300*. Subsequently, the --*scaffold* module in RagTag was used to anchor the corrected contigs to the *C. nigoni* reference genome JU1421. To remove contaminants, bacterial scaffolds were removed based on BLAST v2.11.0 searches against the NCBI nt database with E-value threshold of over 1e-09. The remaining scaffolds were then underwent five rounds of polishing with Pilon v1.24^92^ by mapping Illumina short reads to the assemblies using BWA v2.2.1 to improve sequence accuracy. For each assembled genome, protein-coding gene models were generated with BRAKER3 v3.0.8^93^ using a custom library of protein sequences from *C. elegans* N2 (WormBase, WS283) and *C. briggsae* QX1410 (PRJNA784955). Genome and gene model biological completeness was estimated using BUSCO v5.0.4^94^ with the *nematoda_odb10* database.

### Selection of relatedness group reference genomes for group-specific HDR calling

Detection of HDRs in *C. elegans* relied on clustering one kilobase (kb) partitions of the genome that exceeded empirically determined thresholds of single-nucleotide variant count and relative coverage depth using short-read sequencing alignments between wild strains and the laboratory strain (N2) reference genome^18^. Using a single reference genome to call HDRs species-wide in *C. elegans* was likely only possible because of minimal population structure driven by large-scale selective sweeps^16^. Considering previous reports of comparisons in *C. briggsae* between strains from the Tropical and Kerala relatedness groups that displayed a lack of marked differences in absolute sequence divergence among chromosomal domains^27^, we expected that the increased density of single-nucleotide variation expected in HDRs would be masked by standing genetic variation stemming from divergence between relatedness groups. To overcome this limitation, we called HDRs in each *C. briggsae* relatedness group using group-specific reference genomes. Relatedness groups composed of less than three isotypes or that lacked a reference genome were excluded. A total of seven relatedness groups reference genomes were selected (TD2, BRC20492; TD1, BRC20530; AD, ECA2670; KD, JU1348; Temperate, JU2536; TH, NIC1660; Tropical, QX1410) (Fig. S13).

### Identification of HDRs

A schematic of the HDR calling workflow described below is available (Fig. S24). To classify genomic regions as hyper-divergent, we first aligned 15 long-read genome assemblies from strains in the Tropical relatedness group (ECA1657, ECA176, EG6267, JU3207, JU3237, NIC1529, QG1005, QG2641, QG2665, QG2892, QG2902, QG2964, QG789, QG791, and QX1796) against the Tropical reference genome (QX1410) using *nucmer* v3.1 (custom parameters, --maxgap 500 --mincluster 100)^95^. Alignments smaller than 1 kb were discarded. The QX1410 reference genome was then partitioned into 1 kb bins using *bedtools* v2.31.1 *makewindows*^96^. In each strain, the alignment identity of every bin was estimated from the average identity of the alignment covering the bin. When multiple alignments of variable length and identity covered a bin, the longest alignment was used to estimate the bin identity. The percentage of bases covered by at least one alignment in each bin was also estimated. Bins under 96% identity or 60% percent bases covered were classified as hyper-divergent (Fig. S25). To further refine region boundaries, non-hyper-divergent bins that were flanked by hyper-divergent bins on both sides were re-classified as hyper-divergent (gap re-classification step). Contiguous hyper-divergent bins were clustered into HDRs (bin clustering step).

Next, we filtered the hard-filtered VCF containing variant calls from all 715 isotype reference strains down to the 501 isotype reference strains in the Tropical relatedness group using *bcftools* v1.21^61^. Only the variant sites where at least one of the 501 isotype reference strains in the Tropical relatedness group had an ALT allele were kept after subsetting. For each strain, we estimated the SNV count in each reference bin using *bcftools* v1.21^61^ and *bedtools coverage* v2.31.1^96^, and the percentage of bases covered in every reference bin using *mosdepth* v0.3.10^97^. We tested a wide range of threshold pairs, including raw variant count (5 to 30 SNVs per kb, 1 SNV step) paired with percent bases covered thresholds (5% to 90%, 5% step), to classify and cluster bins into HDRs using short-read alignment data for the 15 strains in the Tropical relatedness group where long-read-based HDRs were previously called. We selected the optimal variant count and percent bases covered thresholds for short-read HDR calls by concordance between short- and long-read HDR call boundaries. We estimated the concordance of short- and long-read HDR call boundaries by measuring the overlap fraction (proportion of a long-read HDRs that overlapped a short-read HDRs) and the excess fraction (proportion of a short-read call that exceeded the boundary of an overlapping long-read call). For each strain, we estimated the mean overlap fraction and mean excess fraction across every overlap between the short-and long-read HDR calls at every threshold pair (Fig. S26-27). We also estimated the completeness and accuracy of the short-read hyper-divergent calls by calculating recall (proportion of long-read calls that had any overlap with short-read calls), and precision (proportion of short-read calls that had any overlap with long-read calls) (Fig. S28-29). Recall and precision were used to estimate an F1 score for each threshold pair (Fig. S30). We selected the top N (starting at N=1) threshold pairs based on the F1 score for each strain, and increased the value of N by 1 until we identified a threshold pair that was present among all of the seven strains in this comparison (referred to as ‘consensus optimal’). The consensus optimal threshold pair was reached at N=10 (top 2.2% of all threshold pairs), where the optimal thresholds selected were a variant count of 11 paired with 90% bases covered (Fig. S31). We classified bins with estimated variant count or percent bases covered under these optimal thresholds as hyper-divergent, genome-wide, across all 501 wild strains in the Tropical relatedness group. We then performed the same gap re-classification, bin clustering steps described for long-read based HDRs. HDRs separated by less than 5 kb were merged (region clustering step), and HDRs smaller than 5 kb were discarded (size filtering step). For each of the remaining relatedness groups, we aligned short-read DNA sequences from each isotype reference strain against their respective relatedness group reference genome, called variants using GATK (see *Alignment and variant calling* section above), and called HDRs using the same methods and thresholds described for the Tropical relatedness group. We mapped the positions of HDRs called in non-Tropical relatedness groups to the QX1410 reference genome using genome alignments generated with *nucmer* v3.1 (Fig. S32). When genome alignments did not span one or both boundaries of a HDR, we extended the region to the most proximal alignment (up to 50 kb distance) outside of the region boundary or dropped the region when no proximal alignment was found.

### Functional enrichment analysis

InterProScan v5.75^38^ was used to identify functional protein domains and their associated Gene Ontology IDs that are enriched in genes found in HDR arm domains and non HDR arm domains. InterProScan was run with options --goterms, --iprlookup, and --disable-precalc on the longest-isoform proteome of QX1410 created using *agat_sp_keep_longest_isoform.pl* from AGAT v1.4.1^98^ and gffread v0.12.7^99^. A total of 15,289 QX1410 proteins were annotated with InterProScan functional protein domains, and 11,756 proteins were annotated with Gene ontology IDs. Of the 15,289 proteins with domain annotations, genes found on chromosome arms were used as the background set for enrichment analysis, resulting in 5,301 genes being used. Of the 11,756 proteins with Gene Ontology ID annotations, 3,905 genes found on chromosome arms were used as the background set for Gene Ontology enrichment analysis. Of the reference genes found within HDRs on chromosomal arms, approximately 58% (3,087 / 5,301) received InterProScan functional protein domain annotations and approximately 40% (2,105 / 5,301) received gene ontology identifiers. All QX1410 genes found on chromosome arms genes were separated based on their presence or absence in HDRs identified among all 501 *C. briggsae* strains belonging to the Tropical relatedness group. For InterProScan domain enrichment analysis, a one-sided hypergeometric test was conducted in R, with a false discovery rate adjusted p-value using the R package *stats* v.4.2.1. InterProScan functional protein domains were classified as significantly enriched if they had an adjusted p-value less than 0.05. Gene Ontology enrichment analysis was performed using the R package clusterProfiler v4.6.2^100–103^. Gene Ontology terms were classified as significantly enriched if they met the criteria of having both a Benjamini-Hochberg adjusted p-value and a q-value of less than 0.05. To identify QX1410 orthologs of N2, OrthoFinder v3.1.0^40^ was run using proteomes translated from the reference protein-coding gene annotations of QX1410^29^ and N2 (WormBase, WS283) using only the longest isoforms for each protein-coding gene model.

## Data availability

The raw short-read sequencing reads for the wild *C. briggsae* strains are publicly available from the NCBI Sequence Read Archive (project PRJNA1149159). The raw PacBio long-read data are publicly available from the NCBI Sequence Read Archive (project PRJNA1291792). Strain information and short-read genomic variation data are available from the CaeNDR (www.caendr.org)^54^. Scripts and related raw and processed data are available at GitHub (https://github.com/AndersenLab/CB_PopGen_HDR_MS/)

## Funding

This work was funded by NSF capacity grant 2224885, HFSP grant RGP0001/2019, and the National Institutes of Health NIEHS grant R01ES029930 (to E.C.A). C.B. acknowledges support by the Centre National de la Recherche Scientifique (CNRS), the Institut national de la santé et de la recherche médicale (Inserm), and Université Côte d’Azur. This work was also funded by Centre National de la Recherche Scientifique, and a grant of the Indo-French Centre for the Promotion of Advanced Research Ref. 6503-4 (to V.S. and M.A.F.). NIH National Institute of General Medical Sciences (NIGMS) grants R35 GM141906 (to M.R.). Biodiversity Research Center (Academia Sinica, Taiwan) internal funds, National Science and Technology Council (Taiwan) grant NSC 100-2311-B-001-015-MY3, and Academia Sinica grant 103-CDA-L01 (to J.W.). NSF CAREER grants MCB-1552101 (to M.A.). Hong Kong Environment Conservation Fund, ECF-160/2023 and General Research Grants (GRF), 12101522, 12100024, and 12101323 (to Z.Z.). NSF Building Research Capacity of New Faculty in Biology (BRC-BIO) grants 2218079 (to Jessica N. Sowa). National Institutes of Health NIEHS grant R50ES037948 (to M.E.G.S.). National Institutes of Health grant F32AI181342 (to A.O.S.)

## Author contributions

N.D.M. and E.C.A. conceived and designed the study.

N.D.M., B.W., L.M.O. and E.C.A. analyzed the data and wrote the manuscript.

R.E.T., M.E.G.S., A.O.S., R.M., M.A., A.B.P., Z.Z., A.D.C., M.V.R., and M.-A.F. edited the manuscript.

R.E.T., A.K, and N.M.R performed whole-genome sequencing for 2,018 *C. briggsae* wild isolates.

R.E.T. and A.K. performed long-read sequencing for 59 *C. briggsae* wild isolates and 11 *C. nigoni* inbred lines.

R.E.T., C.G., C.M.D., T.A.C., G.Z., E.R., L.F., V.D.D., E.H., M.D., C.G., D.E.C, J.C.H., A.S., S.Z., A.R., T.W., A.M., S.H., K.R.A., E.J.K., N.M.R., E.S.S., V.S., H.T., M.A., A.B.P., Z.Z., A.D.C., J.W., M.V.R., M.-A.F., C.B., and E.C.A. performed sample collection and isolation of 2,018 *C. briggsae* strains.

M.E.G.S. performed isotype characterization for 2,018 *C. briggsae* wild strains.

## Acknowledgements

We thank members of the Andersen Lab for providing comments on this manuscript. We especially thank Loraina A. Stinson, Claire Buchanan, Kristen M. Laricchia, Shannon C. Brady, Lauren Seeley, Kreena Patel, Scott E. Baird, Nausicaa Poullet, Julien Dumont, Deborah Bourc’his, Romain Salle, Gunther Hollopeter, Hinrich Schulenburg, Carola Peterson, Yen-Ping Hsueh, Ching-Ting Yang, Chia-Yi Kao, Jessica N. Sowa, Jay Ni, Carolynn M. O’Donnell, Fabien Duveau, Amhed Vargas Velazquez, Isabelle Nuez, Jean-Baptiste Pénigault, Gautier Brésard, Alexandre Peluffo, Fabrice Besnard, Henrique Teotonio, Amir Yassin, Christopher Nelson, V. Robert, H. Baylis, A. Akay, M. Volovitch, R. Ferrière, M. Murmu, Renaud de Rosa, L. Lokmane, M. Joron, V. Debat, N. Gidaszewski, L. Barre, A. Evin, C. Yujin Kim, D. Peigne, I. Felix, Bradford Heights Elementary School and Francis S. Grandinetti Elementary School Nematode Hunters, and citizen scientists for contributing wild *C. briggsae* strains to CaeNDR.

## Notes

### Competing Interest Statement

The authors have declared no competing interest.

### Summary of Updates

No changes to the manuscript or supplement were made; Author list updated

