## Supplementary Figures for "*Caenorhabditis briggsae* ancestral genomic hyper-diversity contrasts with globally distributed genome-wide haplotypes"

Figure S1: Geographic distribution of 715 *C. briggsae* isotype reference strains collected worldwide.

Figure S2: Summary of 17 cosmopolitan isotypes.

Figure S3: The genetic similarity of the strains within the largest cosmopolitan isotype, NIC174 isotype, and its most closely related isotype SOW22.

Figure S4: Maximum-likelihood tree of 715 *C. briggsae* isotype reference strains with detailed label of strain names.

Figure S5: Genotypes Principal Component Analysis (PCA) by each chromosome.

Figure S6: Genotypes Principal Component Analysis (PCA) after different iterations of outlier removal.

Figure S7: Mean genetic similarity within and between relatedness groups.

Figure S8: Minimization of cross-validation error across ADMIXTURE runs.

Figure S9: Summary of ADMIXTURE analysis.

Figure S10: Population structure of 715 global *C. briggsae* isotype reference strains.

Figure S11: The non-admixed isotypes by relatedness groups across 10 ADMIXTURE runs, displaying only the five small (<10 isotypes) relatedness groups (TD2, TD3, ID, NWD, Hubei).

Figure S12: Nucleotide diversity is concentrated in punctuated genomic regions.

Figure S13: Classification of 715 *C. briggsae* isotype reference strains into relatedness groups.

Figure S14: Hyper-divergent region counts and extent between isotypes from the Tropical and other relatedness groups.

Figure S15: Estimates of percent genome span covered by hyper-divergent regions relative to the percent of variants in hyper-divergent regions.

Figure S16: Gene content diversity across *C. briggsae* genome assemblies in a 120 kb hyper-divergent region on chromosome I.

Figure S17: Gene content diversity across *C. briggsae* genome assemblies in hyper-divergent regions on chromosomes I, III, and V.

Figure S18: Comparison of gene tree topologies against a consensus tree of all single-copy orthologs.

Figure S19: Average pairwise nucleotide diversity (π) across different geographic subsets of the 715 *C. briggsae* isotypes.

Figure S20: Population mutation rate (Watterson's θ) across different geographic subsets of the 715 *C. briggsae* isotypes.

Figure S21: Intermediate-frequency alleles near and at chromosomal arms across relatedness groups.

Figure S22: Hyper-divergent regions display lower or no difference in absolute divergence across different relatedness group comparisons.

Figure S23: Genetic similarity score distribution and cutoff.

Figure S24: Hyper-divergent region parameter optimization and calling workflows

Figure S25: Size and identity features of long-read based hyper-divergent regions at variable identity thresholds.

Figure S26: Mean overlap fraction between short- and long-read HDR calls.

Figure S27: Mean excess fraction between short- and long-read HDR calls.

Figure S28: Recall of long-read HDR calls.

Figure S29: Precision of short-read HDR calls.

Figure S30: F1 of short-read HDR calls.

Figure S31: Strain and consensus optimal threshold pairs for HDR calls.

Figure S32: Mapping hyper-divergent region calls across relatedness group reference genomes.

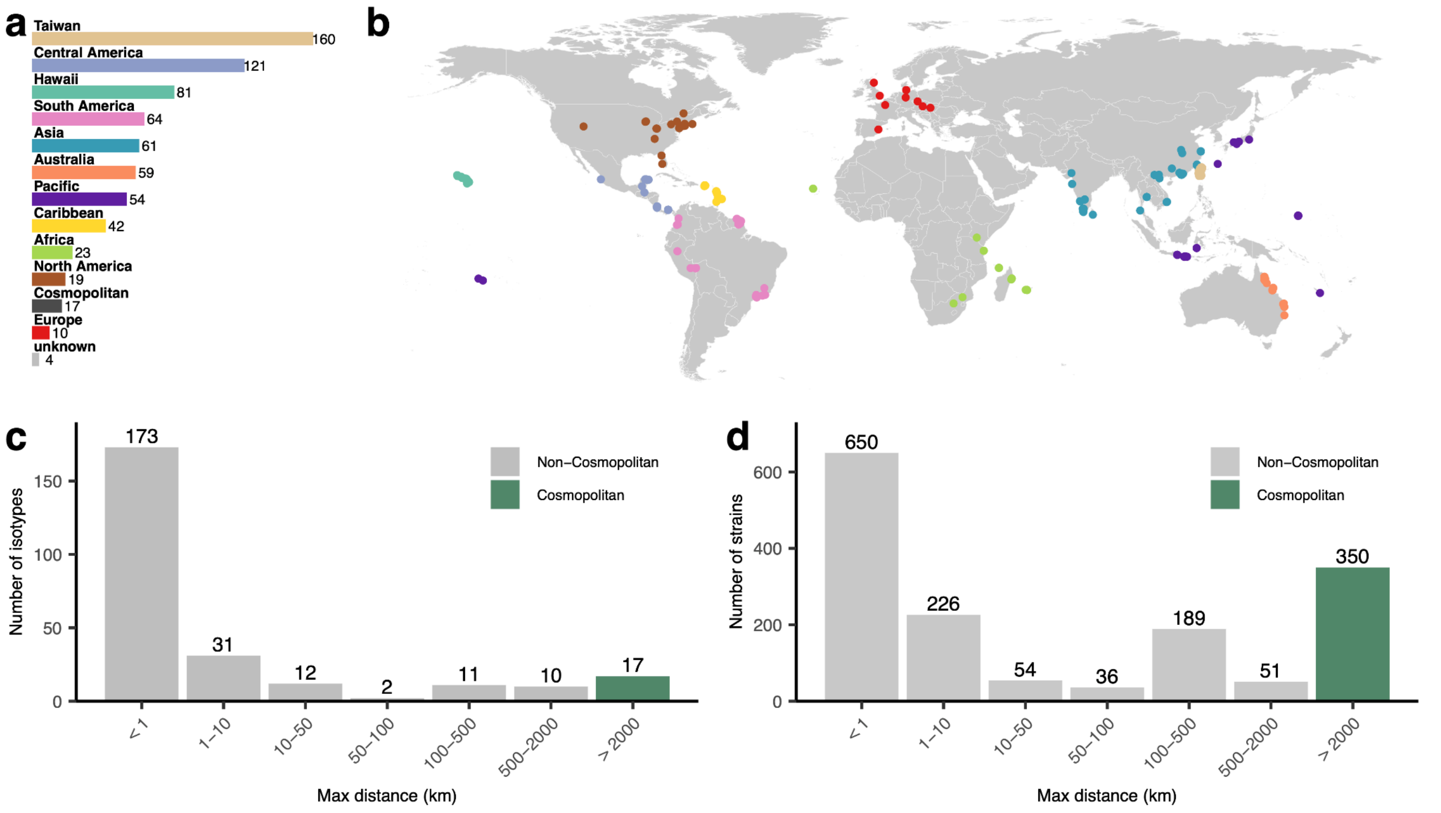


**Figure S1: Geographic distribution of 715 *C. briggsae* isotype reference strains collected worldwide. a,** Total number of isotypes in each geographic region. **b,** Map showing the isolation location of isotype reference strains. **c,** Number of isotypes across bins of maximum geographic distance between strains within each isotype, with isotypes labeled as cosmopolitan or non-cosmopolitan. **d,** Number of strains corresponding to isotypes in panel c in each maximum geographic distance bin, with strains labeled as cosmopolitan or non-cosmopolitan. 457 single-strain isotypes and two isotypes (that comprise 5 strains) where all but one strain lack isolation site information are omitted in panels c and d.

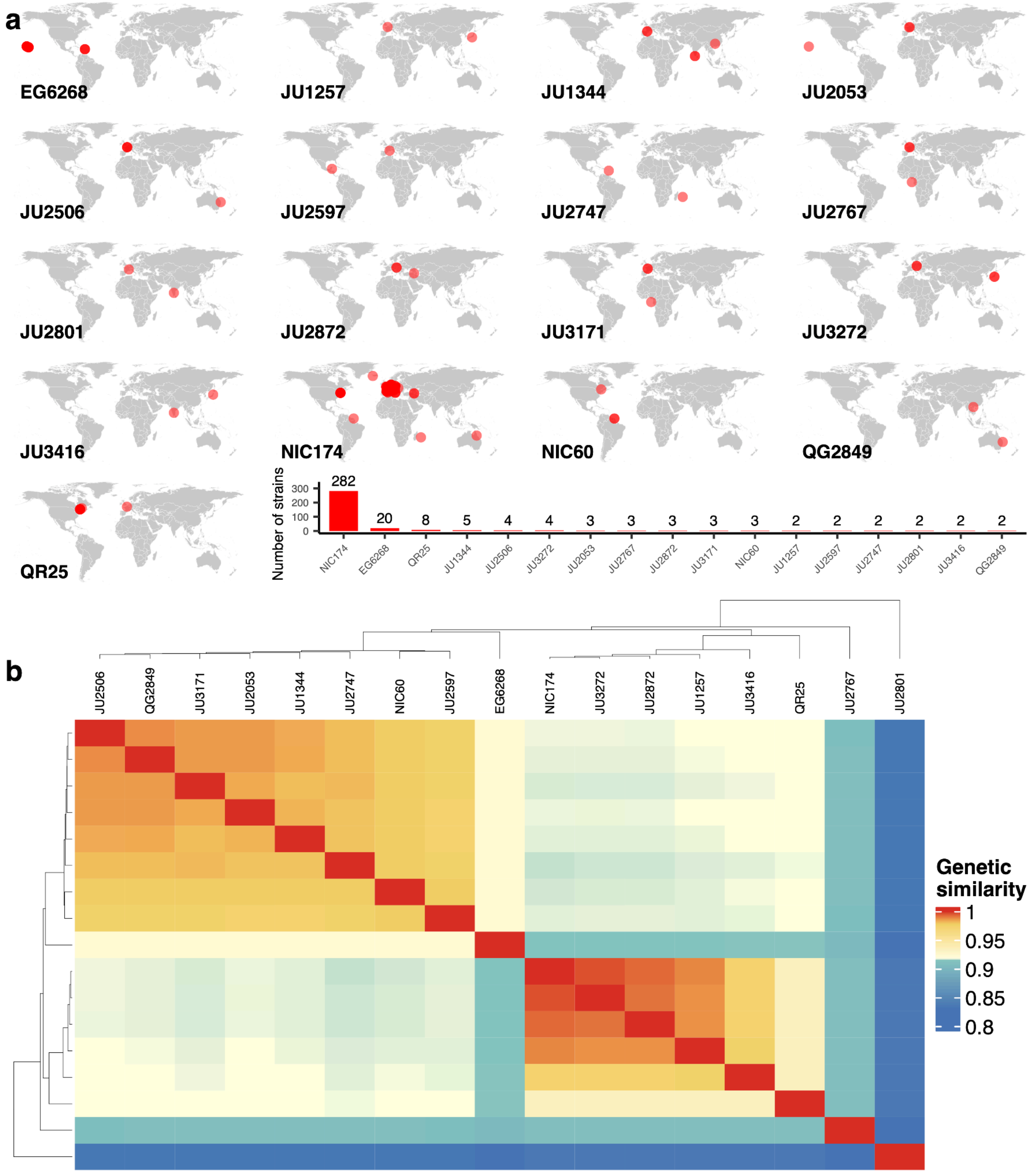


**Figure S2: Summary of 17 cosmopolitan isotypes. a,** Geographic distribution of each strain within the 17 cosmopolitan isotypes and the number of strains per isotype. Each point represents the collection location of a strain within each cosmopolitan isotype. The bar plot on the bottom right shows the number of strains in each cosmopolitan isotype. **b,** Heatmap showing pairwise genetic similarity between all 17 cosmopolitan isotype reference strains. Pairwise genetic similarity was estimated by the proportion of identical alleles across all identified SNVs. The genetic similarity color gradient has breaks specified at the 25th, 50th, and 75th percentile of its distribution.

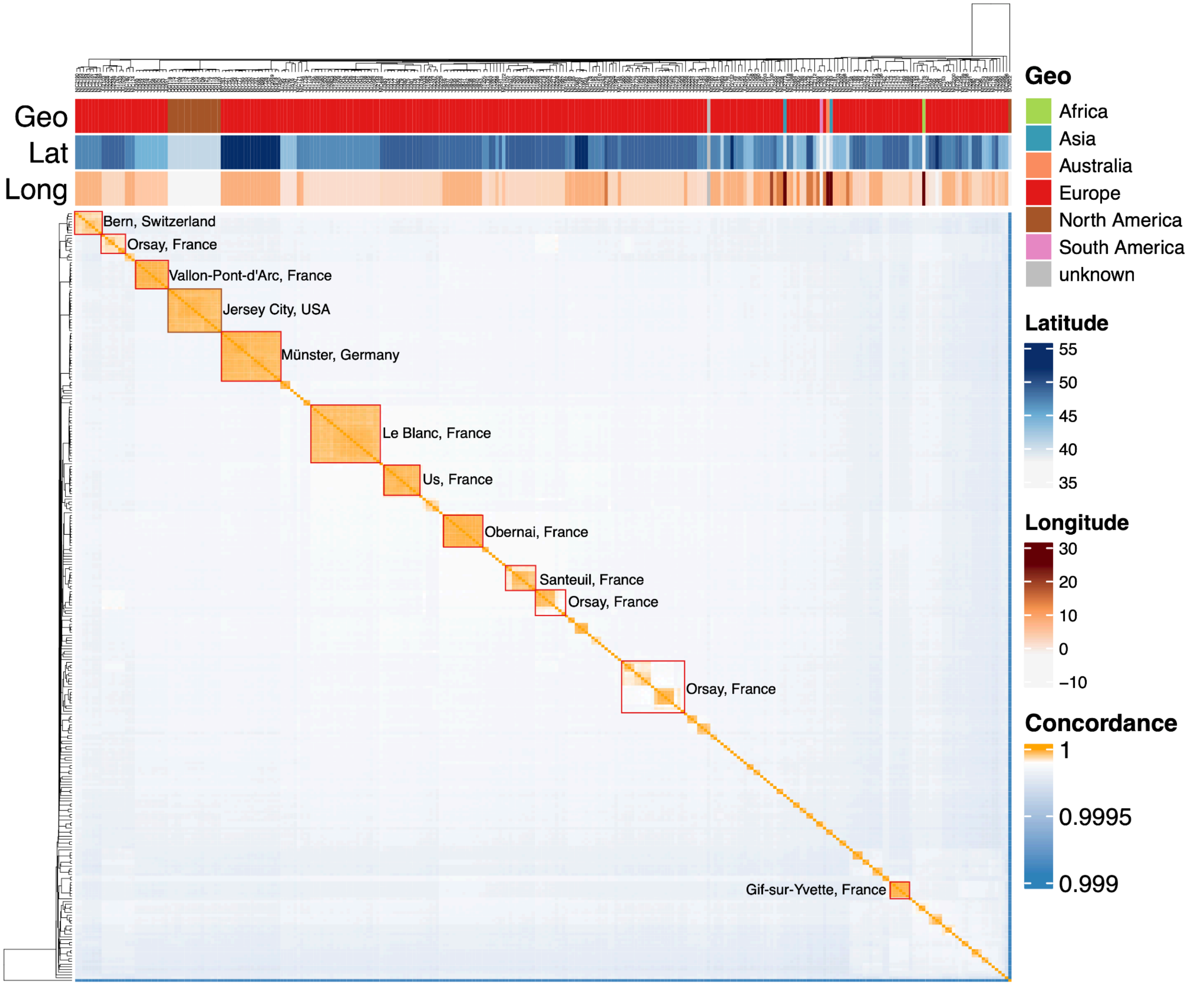


**Figure S3: The genetic similarity of the strains within the largest cosmopolitan isotype, NIC174 isotype, and its most closely related isotype SOW22.** Pairwise genetic similarity was estimated by the proportion of identical alleles across all identified SNVs. The genetic similarity color gradient has breaks specified at 0.9999. In order from top to bottom, bands show the geographic region, latitude, and longitude of the isolation site of each strain. Each red box and label marks the sampling site shared by strains within the same block.

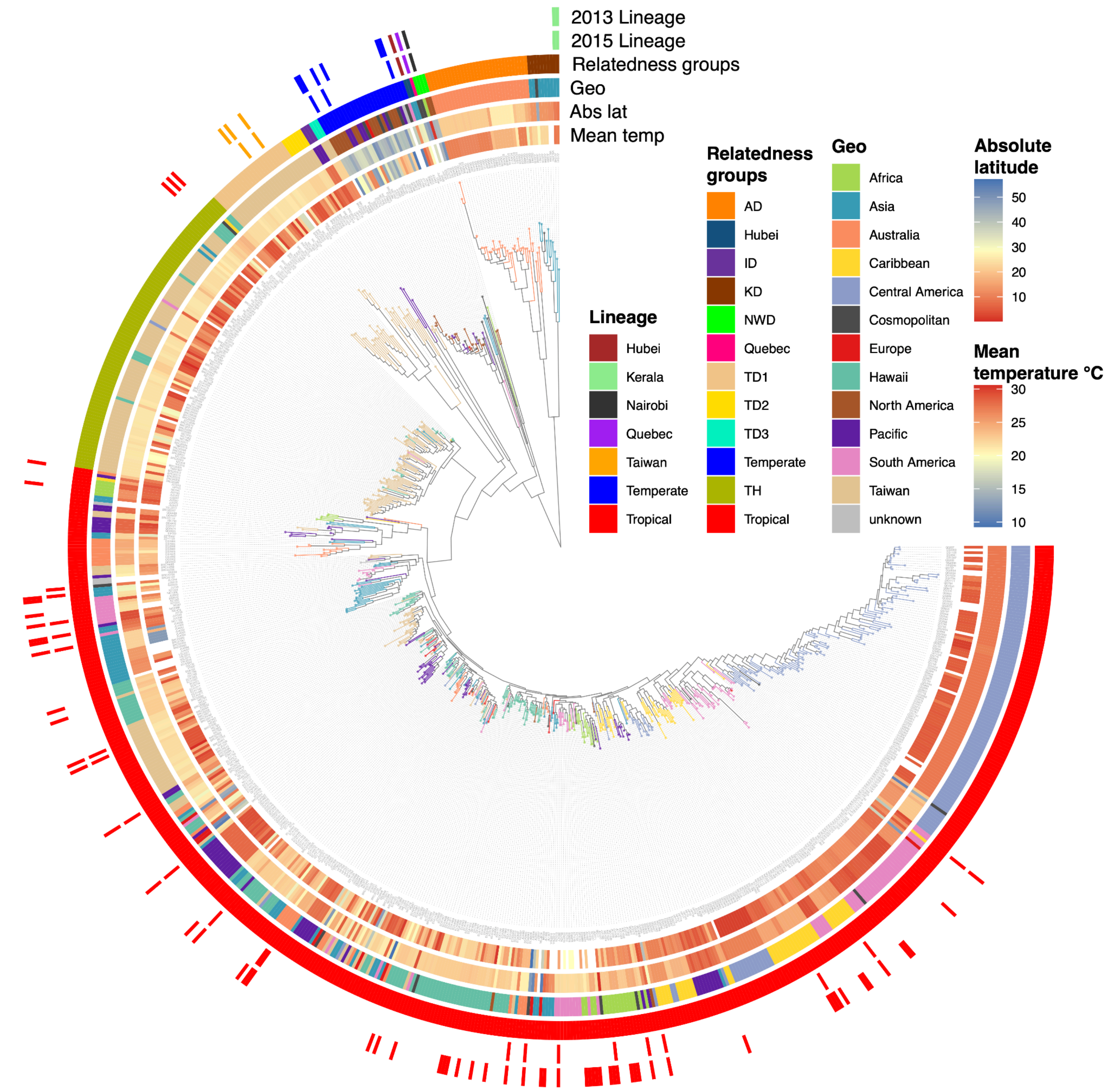


**Figure S4: Maximum-likelihood tree of 715 *C. briggsae* isotype reference strains with detailed label of strain names.** The tree was generated from LD-pruned variants with $r^2$ value less than 0.9 using the GTR+F+ASC+R10 maximum-likelihood substitution model. The color of each leaf branch corresponds to the geographic region of isolation of each strain. The tree is rooted at the split between Kerala/Australia divergent strains and all other isotypes, based on Thomas et al. 2015. From innermost to outermost, each ring is colored by average temperature, absolute latitude, geographic region, relatedness groups defined in this study, and the lineages defined by the two previous studies in 2015 and 2013 (Félix et al. 2013; Thomas et al. 2015)

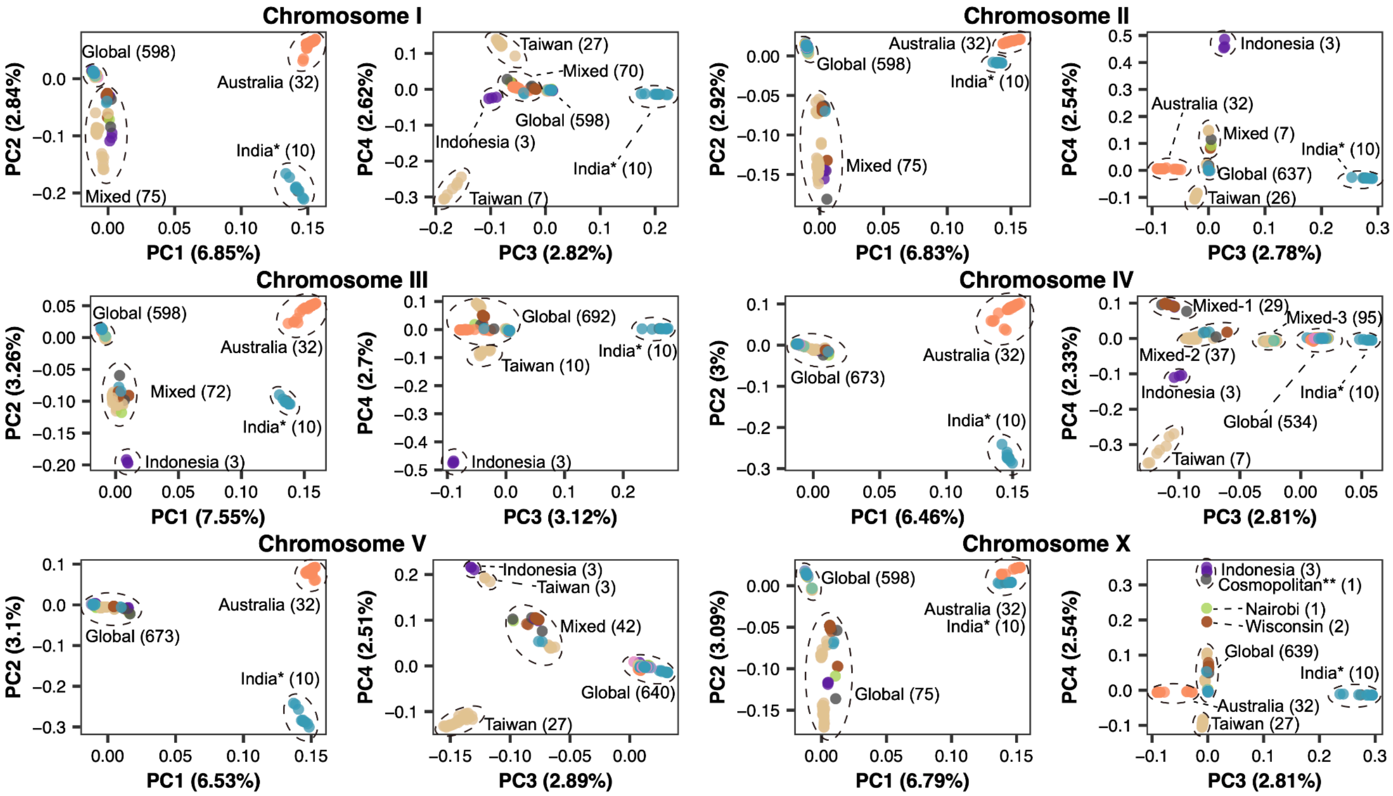


**Figure S5: Genotypes Principal Component Analysis (PCA) by each chromosome.** The ellipses highlight differentiated isotype clusters. The numbers in parentheses show the count of isotypes within each cluster. The asterisk represents that one isotype from the Asia cluster, JU2801, was a cosmopolitan isotype. JU2801 comprises two strains, JU2801 and JU3199, collected from Europe and Asia, respectively. The double asterisk represents that one cosmopolitan isotype JU2767 comprises three strains from different geographic regions. JU2814 and JU2767 collected from Europe, and JU3168 collected from Africa, respectively.

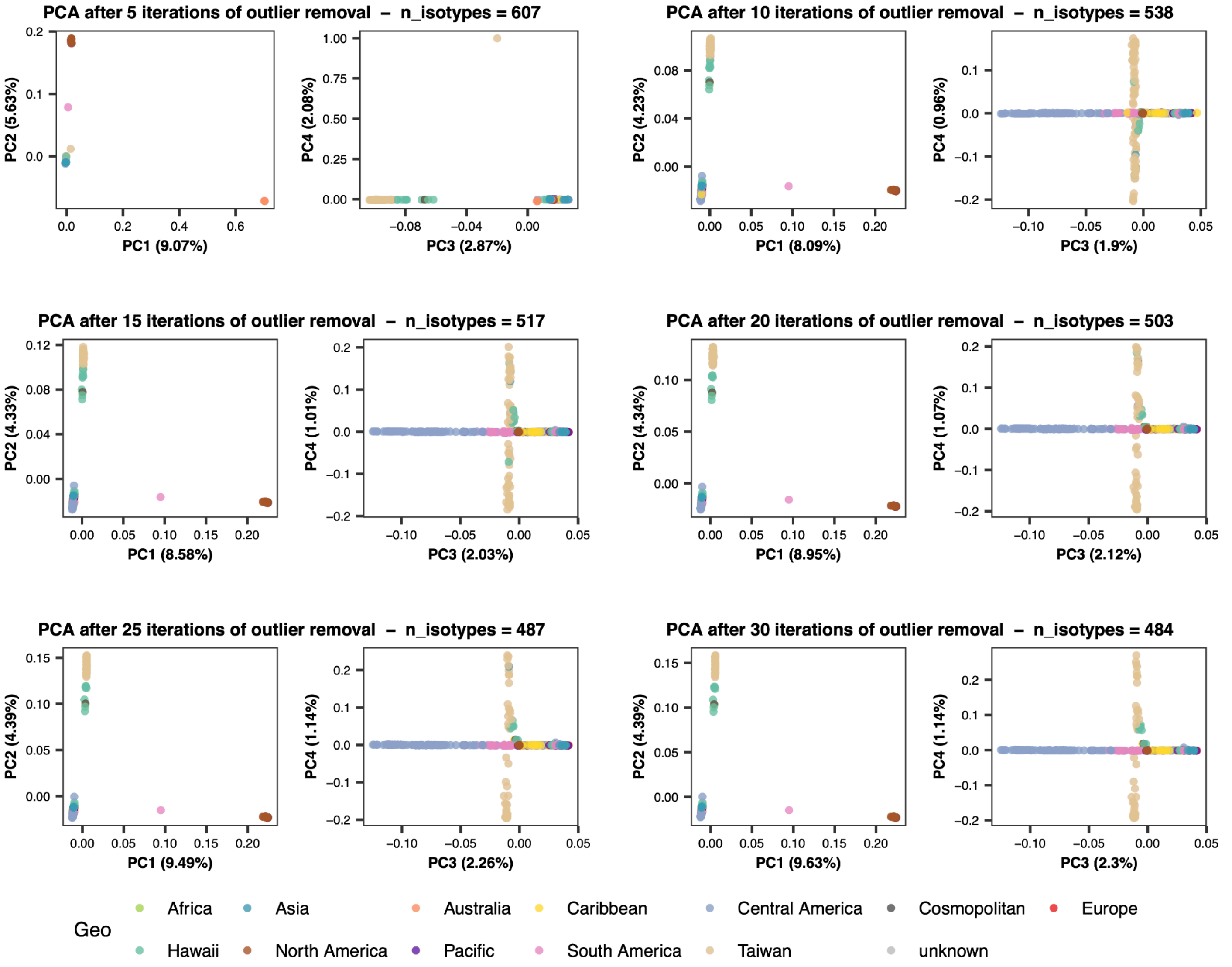


**Figure S6: Genotypes Principal Component Analysis (PCA) after different iterations of outlier removal.** Colors in each panel represent the sampling geographic region of each strain. The n_isotypes in the title of each panel represents the number of remaining isotype reference strains after certain iterations of outlier removal and adding more iterations no longer changes the n_isotypes after 30.

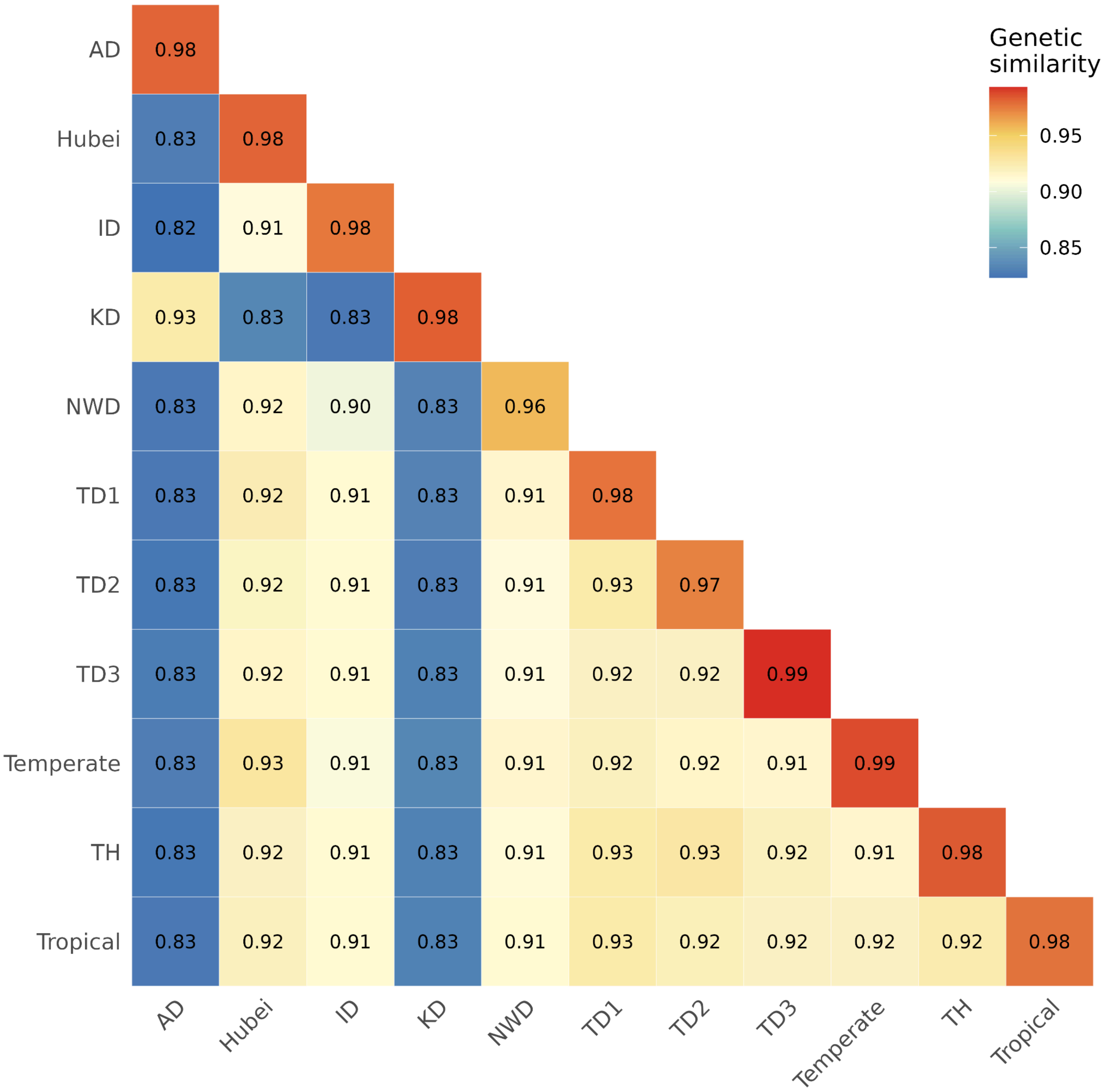


**Figure S7: Mean genetic similarity within and between relatedness groups.** Relatedness groups are shown on the x- and y-axis. Mean estimates of pairwise genetic similarity across all isotypes of any given relatedness group comparison are shown. Genetic similarity is estimated by the proportion of identical alleles at all identified SNV among the 715 isotype strains in this study.

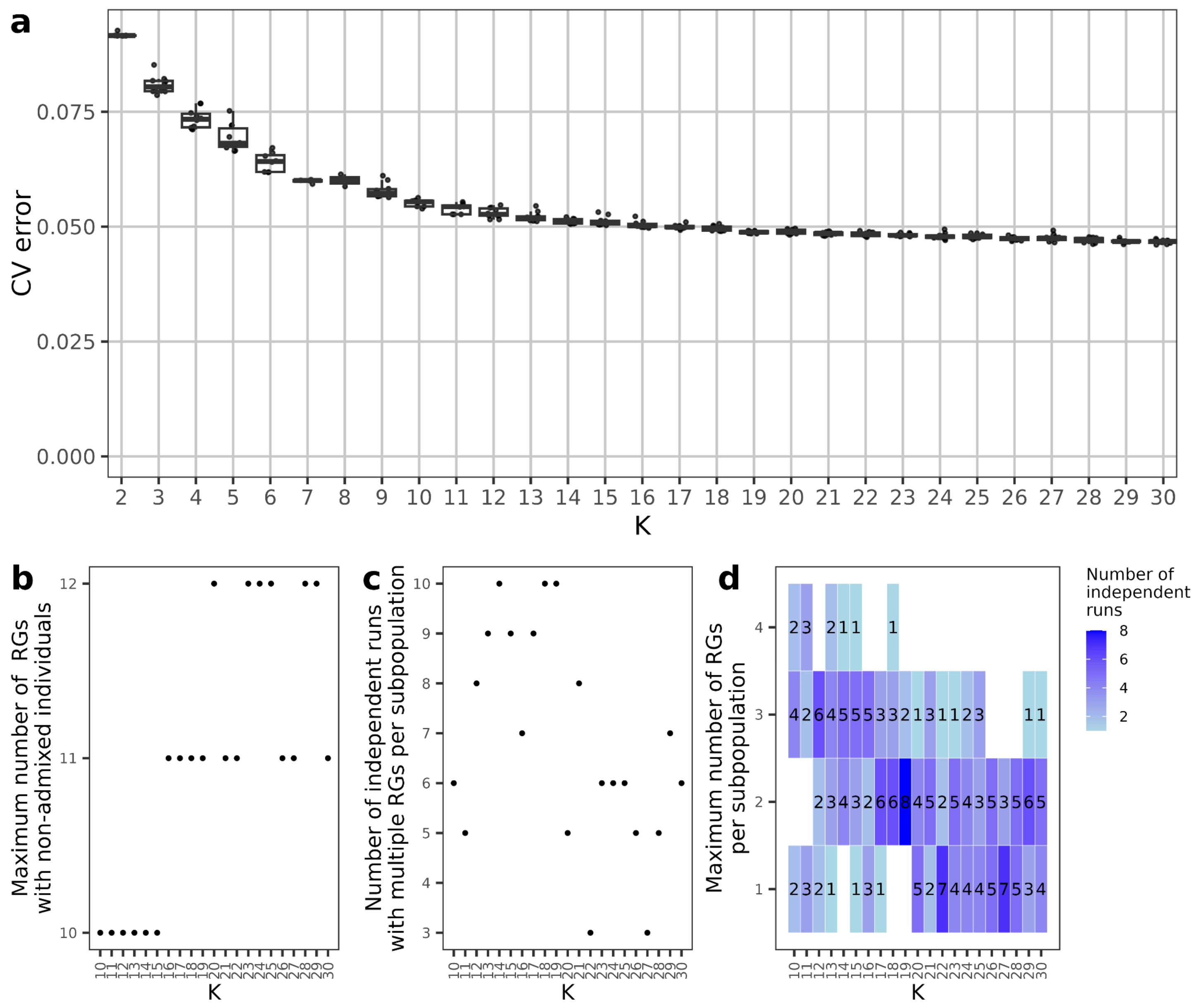


**Figure S8: Minimization of cross-validation error across ADMIXTURE runs. a,** Cross-validation error (y-axis) of ten independent ADMIXTURE runs across K assumed subpopulations. **b,** number of relatedness groups with non-admixed isotypes across K=10 to K=30. **c,** number of ADMIXTURE runs where multiple relatedness groups had individuals assigned to a single subpopulation across K=10 to K=30. **d,** Maximum number of relatedness groups assigned to a single subpopulation across K=10 to K=30.

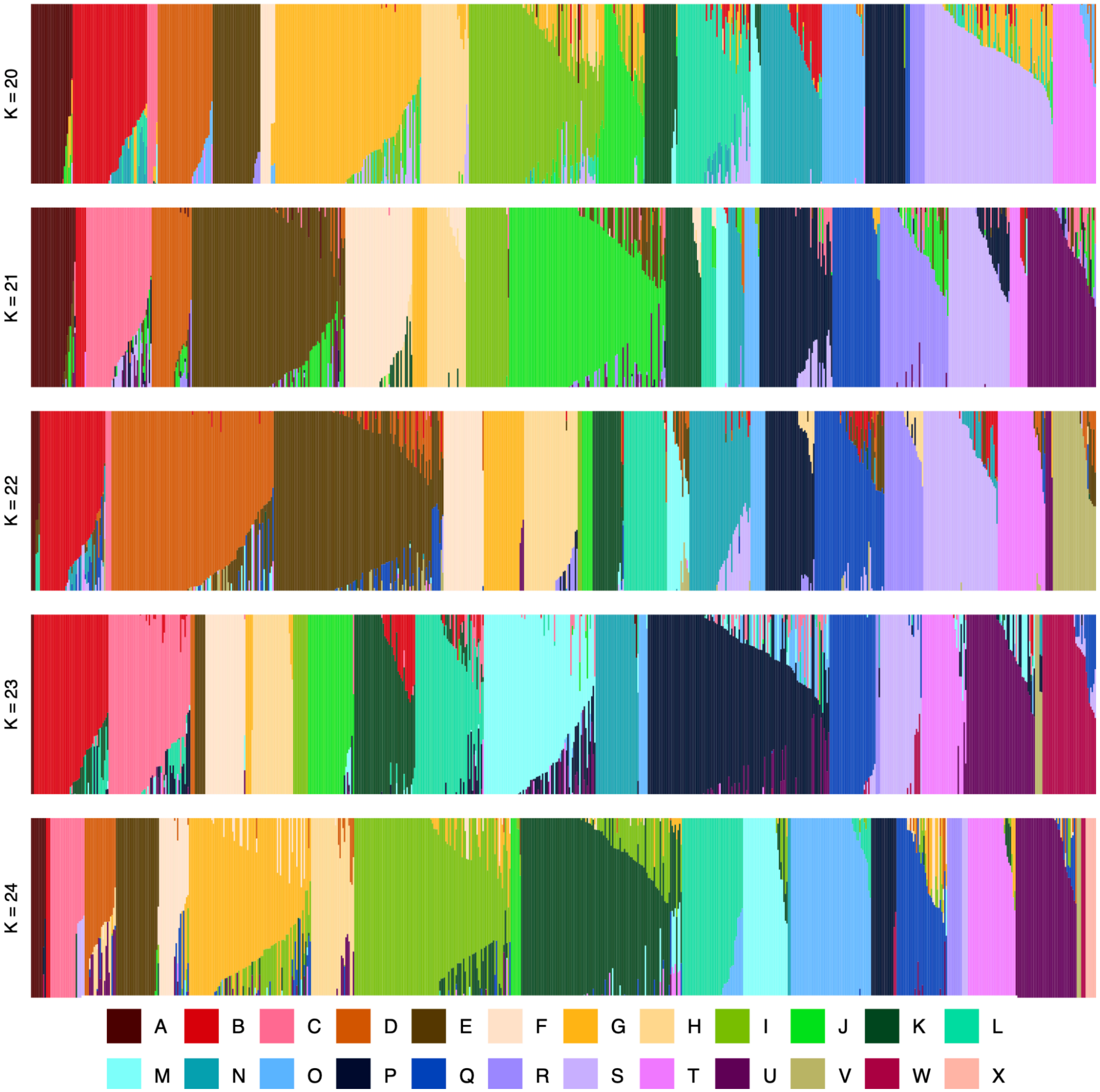


**Figure S9: Summary of ADMIXTURE analysis.** Subpopulation fractions estimated by ADMIXTURE across K ranges ± 2 of best K, 22. Each vertical line represents an isotype, which is partitioned into colored segments that represent the membership fractions for the overall subpopulations.

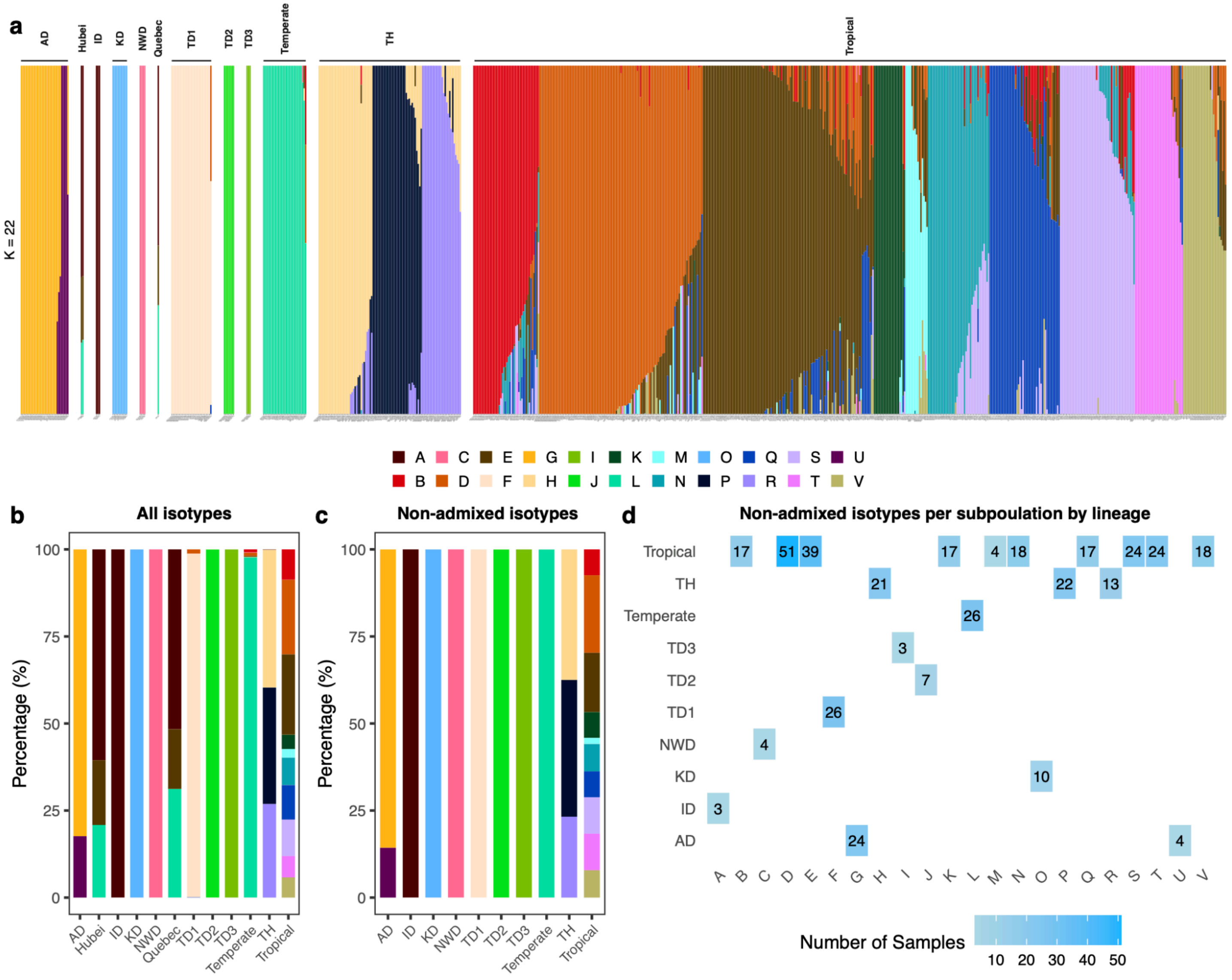


**Figure S10: Population structure of 715 global *C. briggsae* isotype reference strains. a,** Admixture bar graph of population structure in global *C. briggsae*. Each column represents an isotype reference strain. The labels above represent the relatedness groups. **b,** Population proportions among all isotype reference strains in each relatedness group. **c,** Population proportions of non-admixed representative isotype reference strains in each relatedness group. **d,** Heatmap for non-admixed representative isotype reference strains per population in each relatedness group. A non-admixed representative isotype reference strain was defined as a strain with a maximum population fraction over 99.9%. The smallest fraction (0.00001) automatically assigned by the software to each subpopulation was removed when plotting the stacked fractions in panel c. For panel d, only the largest fraction within each non-admixed isotype was used to calculate the stacked bars.

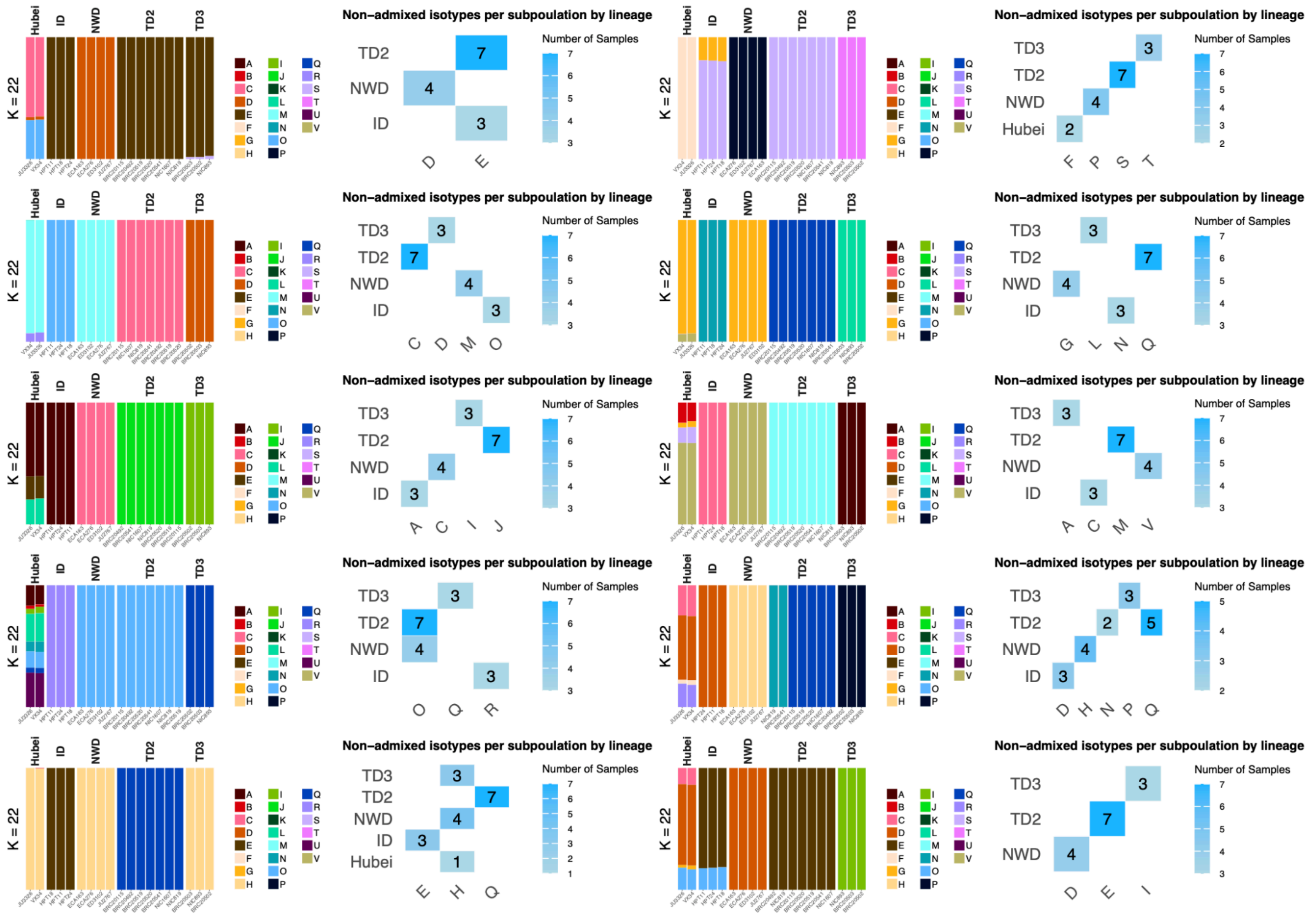

**Figure S11: The non-admixed isotypes by relatedness groups across 10 ADMIXTURE runs, displaying only the five small (<10 isotypes) relatedness groups (TD2, TD3, ID, NWD, Hubei).** Occasionally, the same subpopulation might be assigned to multiple different relatedness groups among these five.

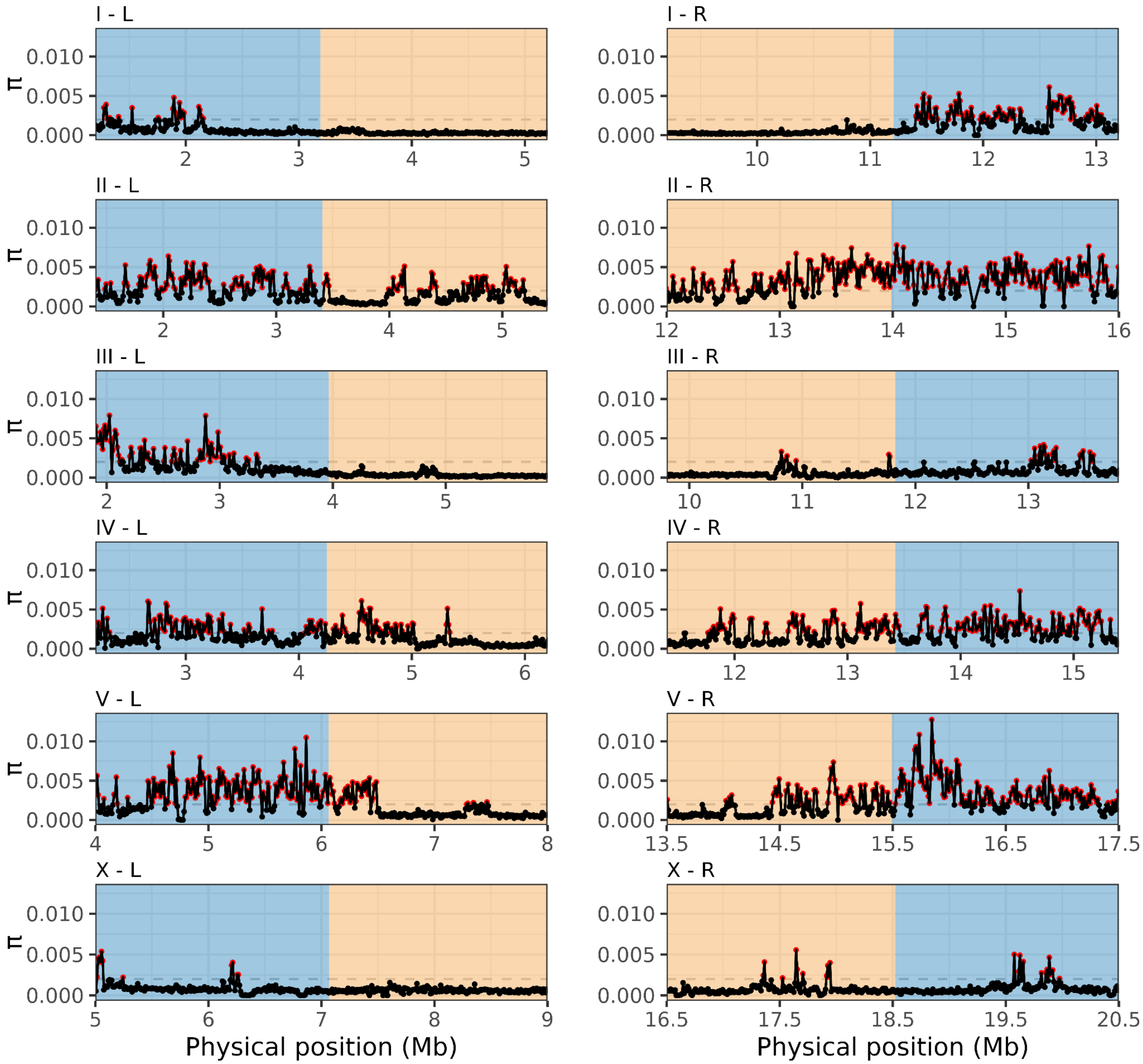


**Figure S12: Nucleotide diversity is concentrated in punctuated genomic regions.** Twelve panels showing estimates of nucleotide diversity (π) in 10 kb windows across approximately 4 Mb of each arm-center chromosomal domain boundary among isotypes of the Tropical relatedness group. Blue denotes the arm domain; yellow denotes the center domain. Peaks of elevated π (>0.002) are shown as red points.

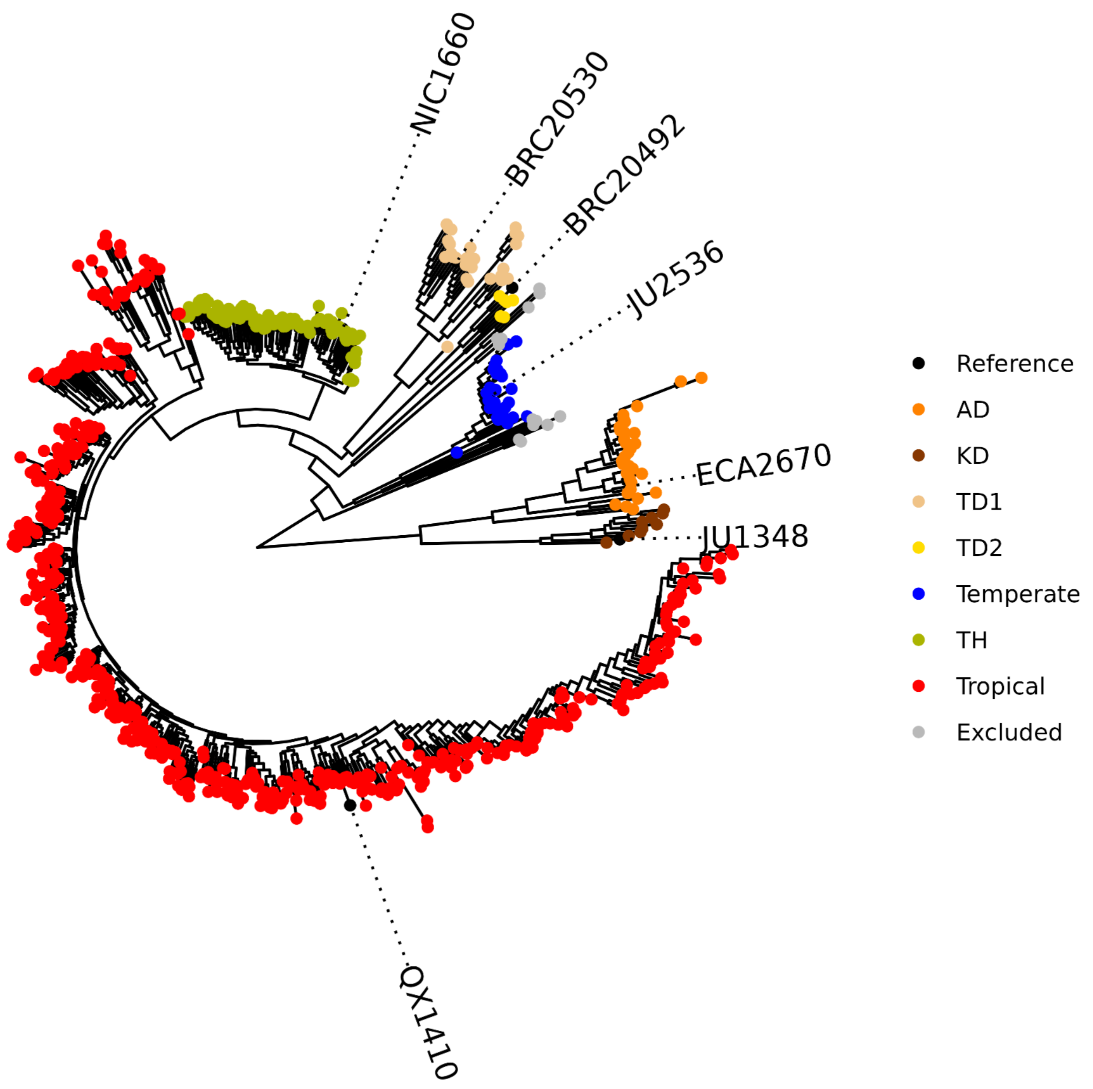


**Figure S13: Classification of 715 *C. briggsae* isotype reference strains into relatedness groups.** Tree generated from LD-pruned variants with $r^2$ value less than 0.9 using the GTR+F+ASC+R10 maximum-likelihood substitution model, with tip nodes colored by relatedness group. A total of seven reference genomes, one for each relatedness group, were selected to call hyper-divergent regions in each group. Relatedness groups (comprising 20 strains) with less than 3 isotypes or that lacked a sequenced reference genome were excluded.

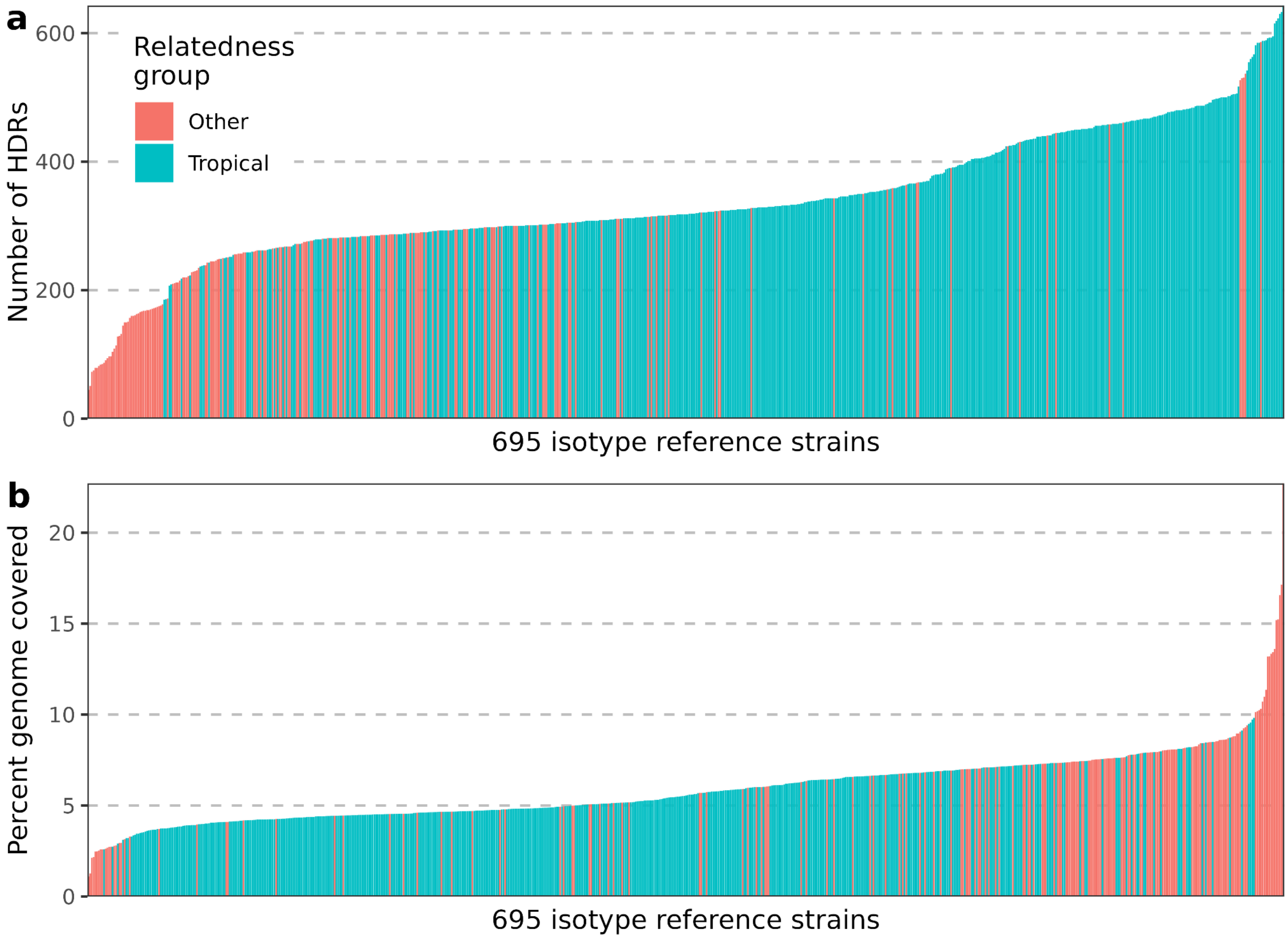


**Figure S14: Hyper-divergent region counts and extent between isotypes from the Tropical and other relatedness groups.** Each bar in the x-axis shows an isotype reference strain colored by whether they are in the Tropical or other relatedness group with the **a,** total number of hyper-divergent regions or **b,** the percentage of the reference genome covered by hyper-divergent regions on the y-axis.

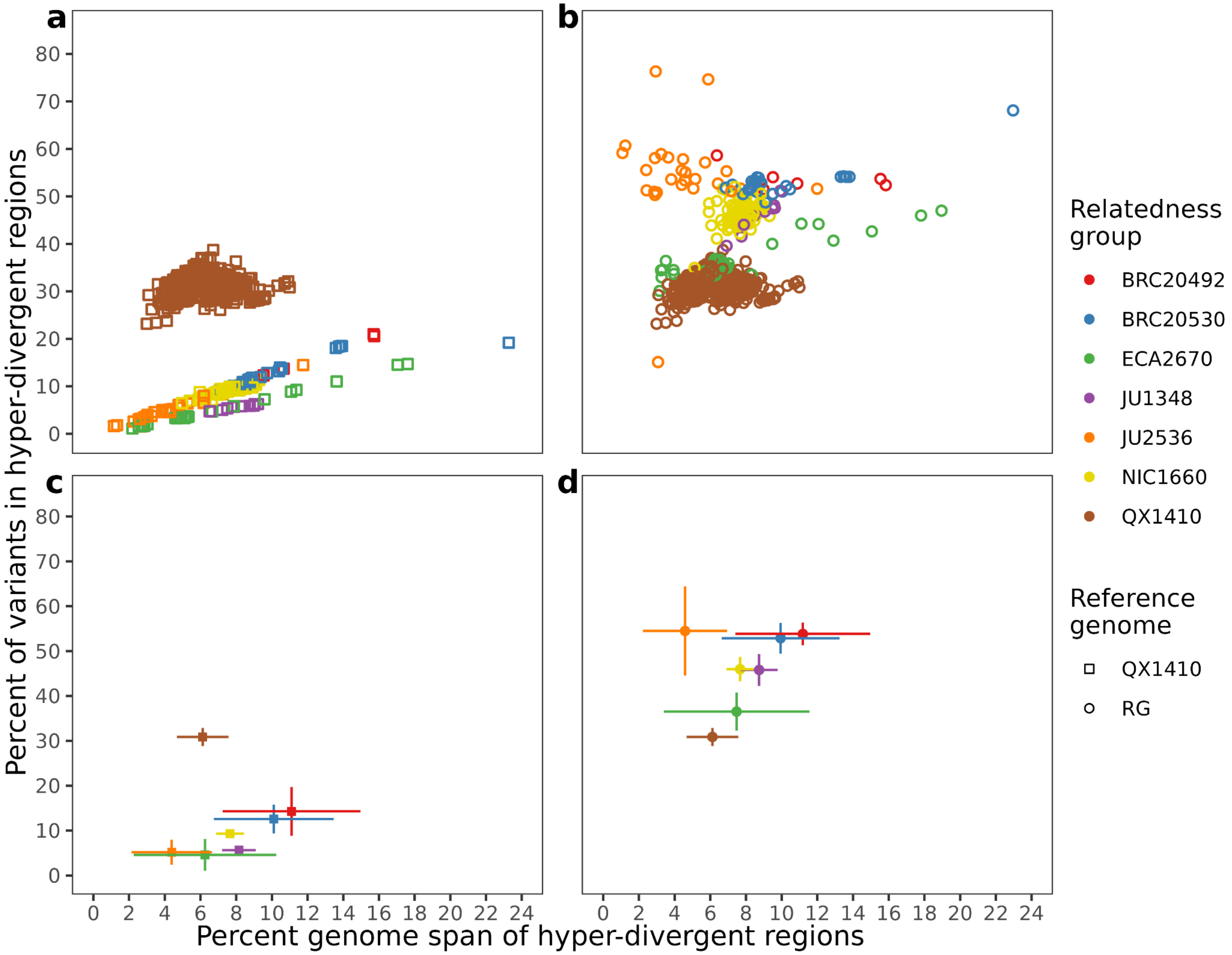


**Figure S15: Estimates of percent genome span covered by hyper-divergent regions relative to the percent of variants in hyper-divergent regions.** Hyper-divergent regions were used to estimate the percent of variants within and percent genome spanned by hyper-divergent regions **a,** in each isotype, relative to the QX1410 reference genome, shown as squares colored by relatedness group **b,** in each isotype, relative to the respective reference genome of each relatedness group, shown as circles colored by relatedness group **c,** averaged for each relatedness group, relative to the QX1410 reference genome, shown as squares colored by relatedness group, with standard deviation as lines on both axes **d,** averaged for each relatedness group, relative to the respective reference genome of each relatedness group, shown as circles colored by relatedness group, with standard deviation as lines on both axes.

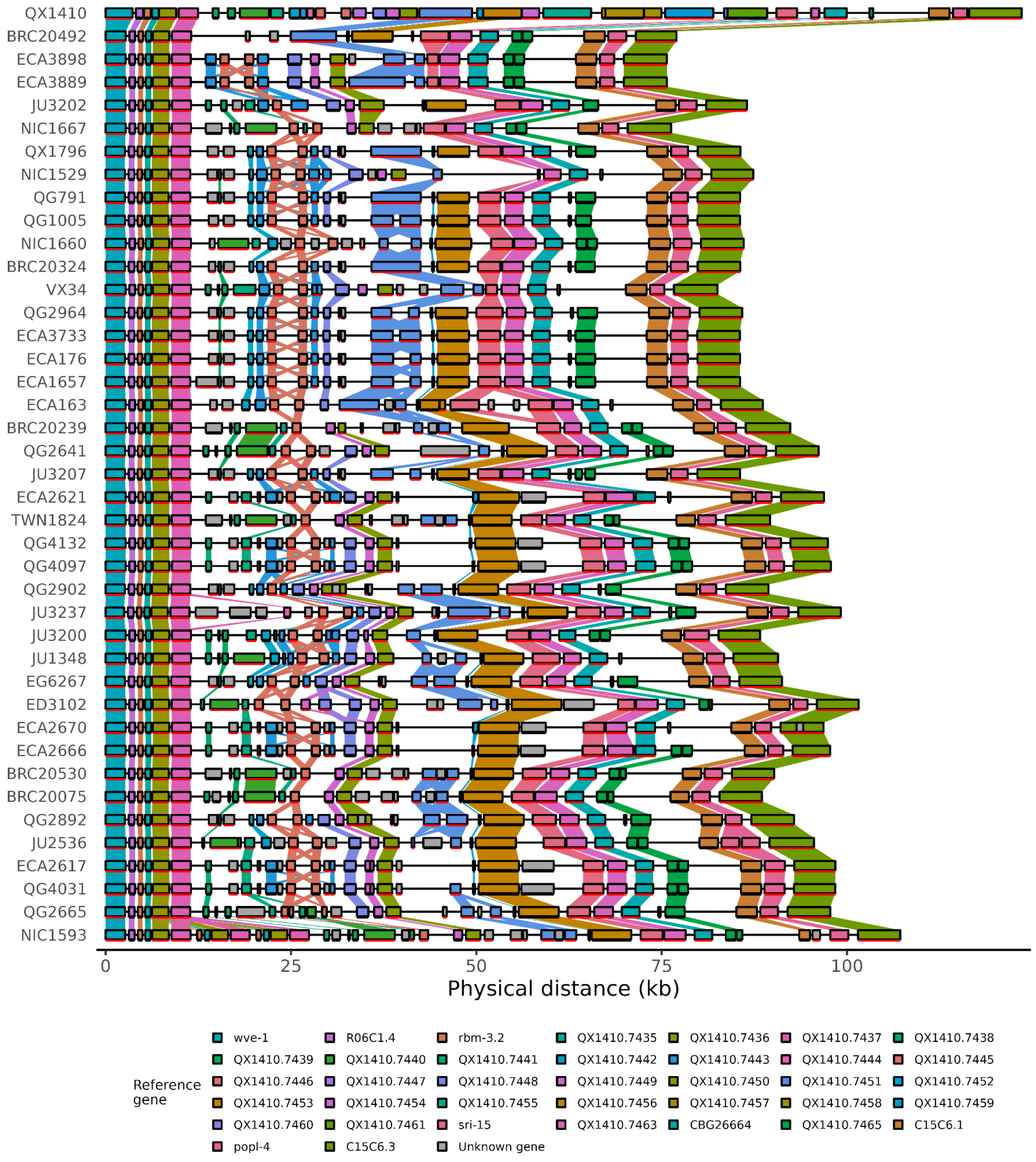


**Figure S16: Gene content diversity across *C. briggsae* genome assemblies in a 120 kb hyper-divergent region on chromosome I.** Each row represents the genome of a *C. briggsae* strain aligned to a 120 kb segment in I:X-X of the QX1410 reference genome (shown in the first row). Each rectangle represents a gene colored after their ortholog in the reference genome, or colored in grey if it's not observed in the reference genome. Colored trapeziums connect orthologs across rows.

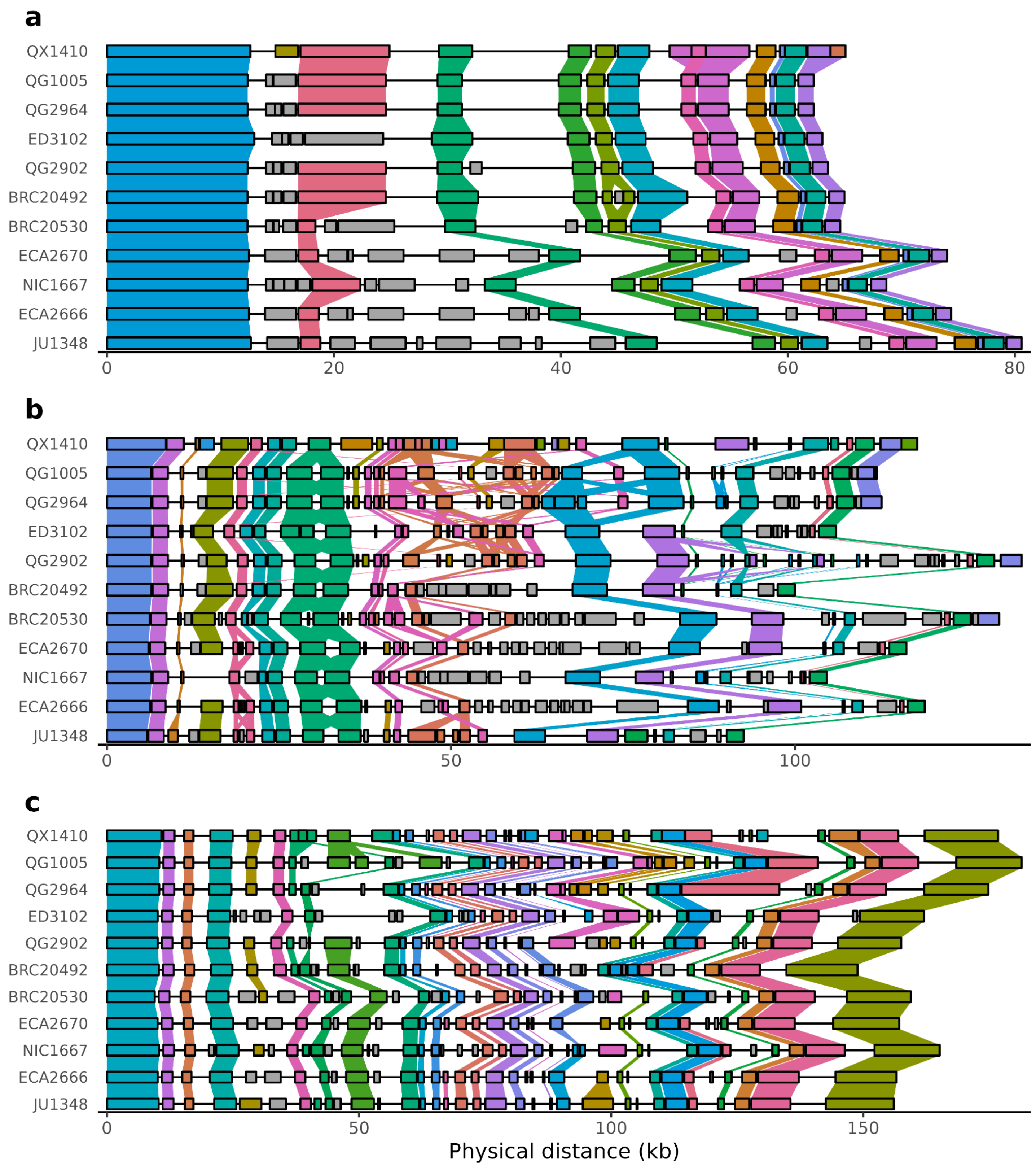


**Figure S17: Gene content diversity across *C. briggsae* genome assemblies in hyper-divergent regions on chromosomes I, II, and V.** Each row represents the genome of a *C. briggsae* strain aligned to **a,** a 56 kb hyper-divergent region in V:838000-894000, **b,** a 110 kb hyper-divergent region in II:12500000-12610000, and **c,** a 175 kb hyper-divergent region in I:12469000-12644000 of the QX1410 reference genome (shown in the first row of each panel). The genome boundaries of the plot are extended to fully include genes that overlap with the start and end positions of the aligned reference region. Each rectangle represents a gene colored after their ortholog in the reference genome, or colored in grey if it's not observed in the reference genome. Colored trapeziums connect orthologs across rows.

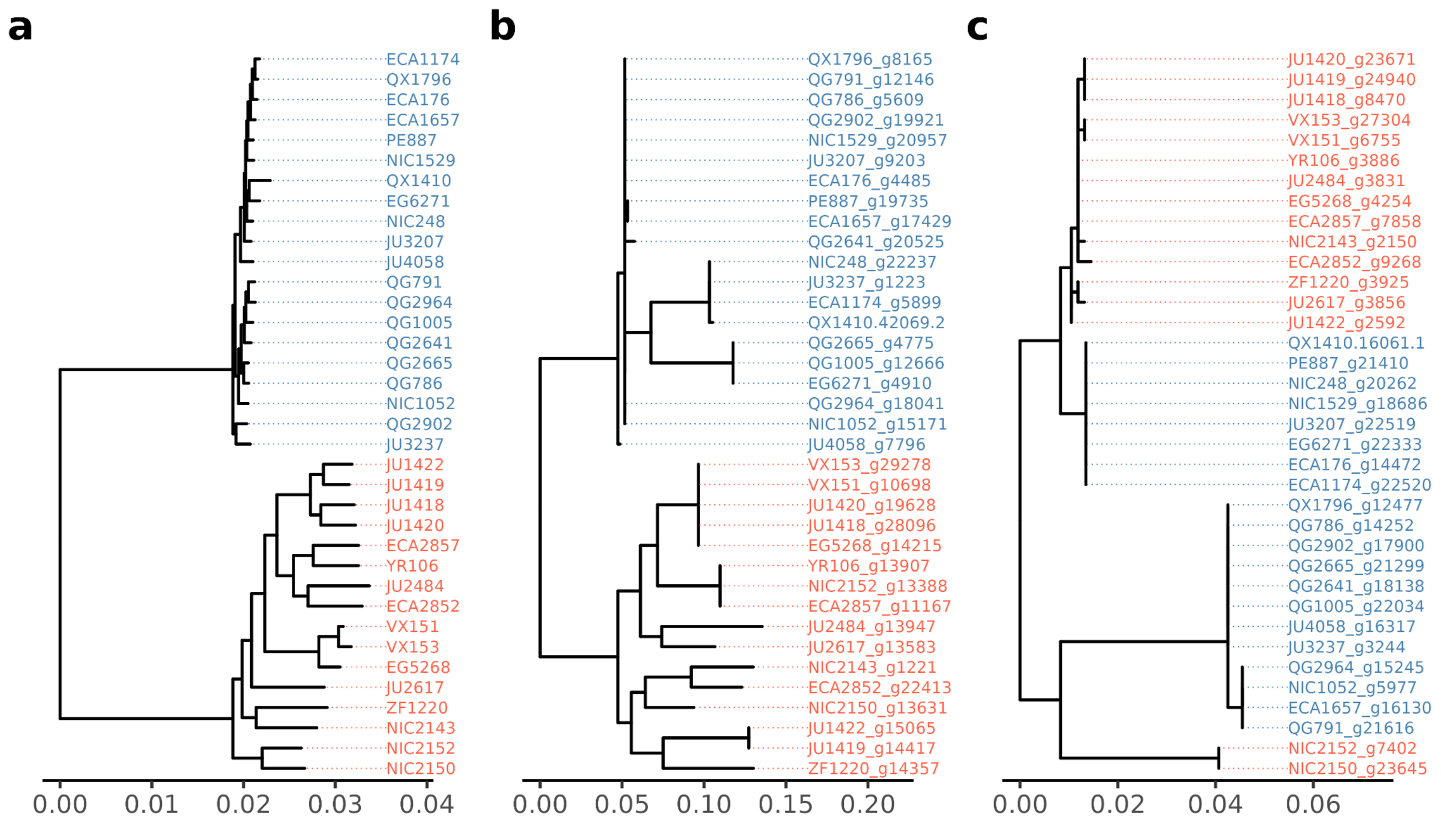


**Figure S18: Comparison of gene tree topologies against a consensus tree of all single-copy orthologs. a,** consensus tree modeled after concatenated alignments protein sequences from single-copy orthologs between *C. nigoni* (red) and *C. briggsae* (blue) compared against **b,** an example gene tree (OG0012935 in File S1) concordant with the consensus tree (with two distinct monophyletic groups, one for each species) and **c,** an example gene tree (OG0018191 in File S1) discordant with the consensus tree (with at least one polyphyletic group) that can potentially be a product of introgression.

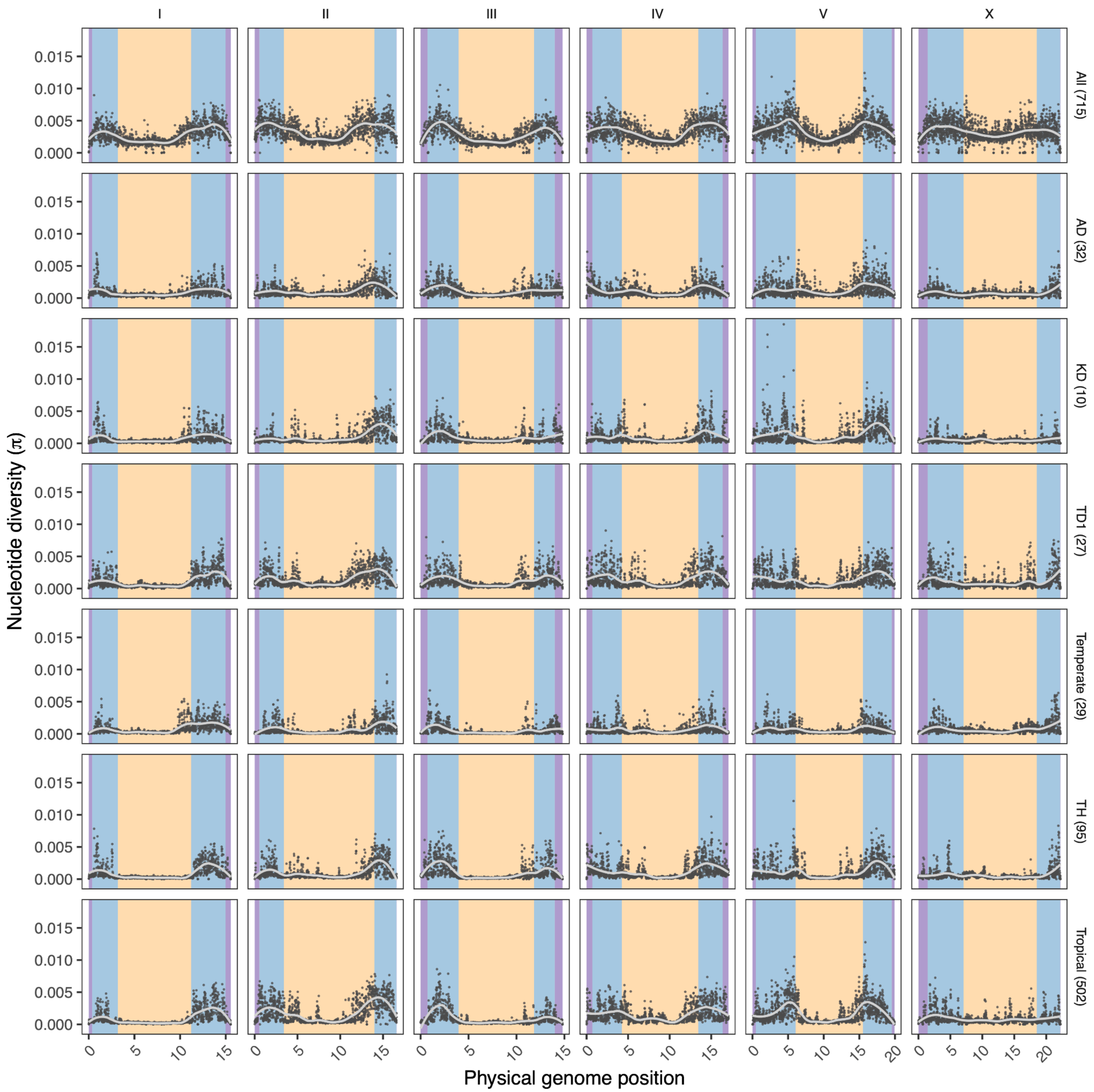


**Figure S19: Average pairwise nucleotide diversity (π) across different relatedness group subsets of the 715 *C. briggsae* isotypes.** Scatter plots showing π estimates in 10-kb windows across the genome. Vertical facets represent each *C. briggsae* chromosome, and horizontal facets represent different subsets of the *C. briggsae* isotypes. Colored backgrounds in nucleotide diversity panels delineate the chromosomal domain boundaries (purple, tip; blue, arm; yellow, center) center). Numbers in parentheses after each subset label show sample size.

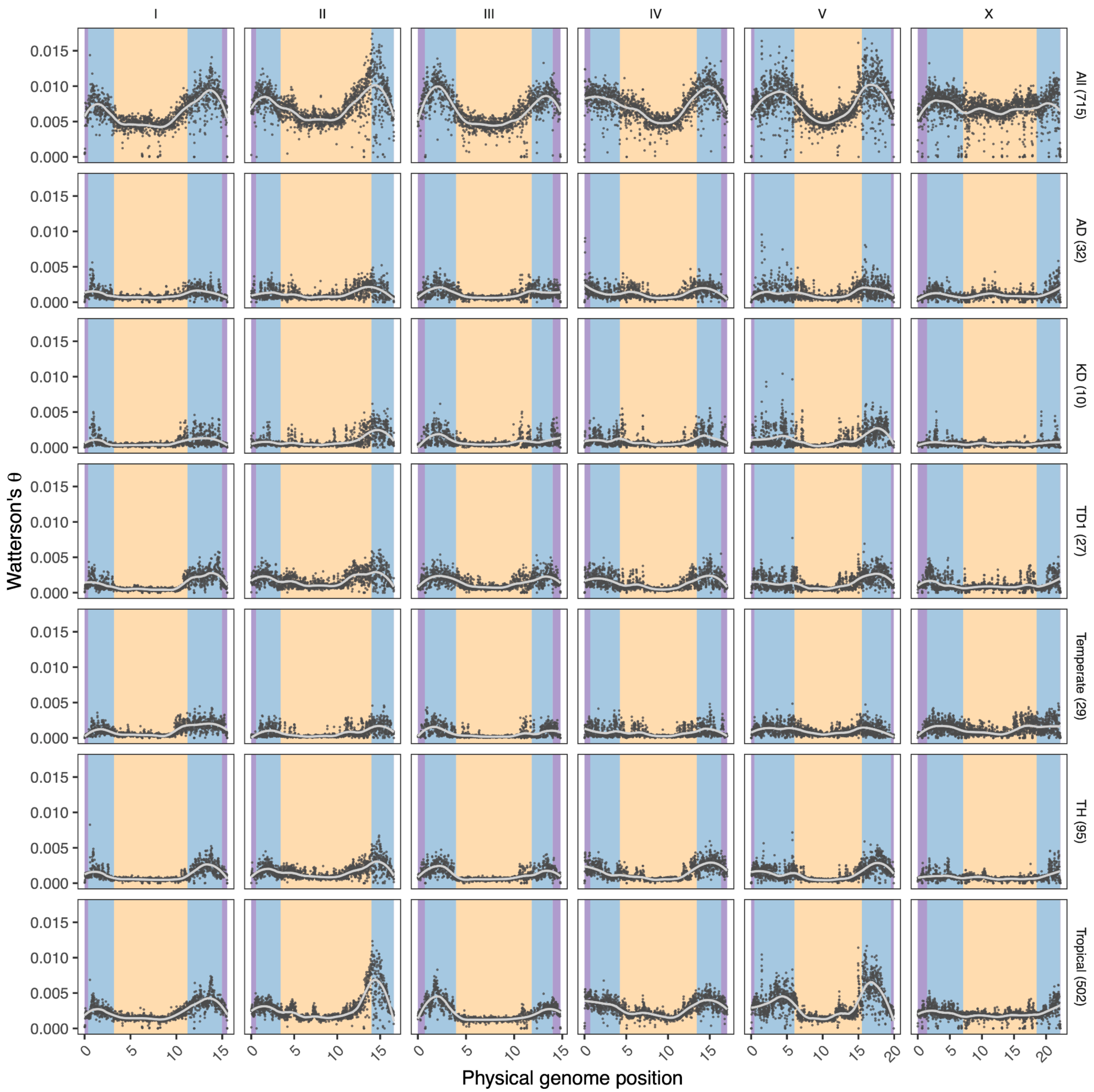


**Figure S20: Population mutation rate (Watterson's θ) across different relatedness group subsets of the 715 *C. briggsae* isotypes.** Scatter plots showing $\theta_W$ estimates in 10-kb windows across the genome. Vertical facets represent each *C. briggsae* chromosome, and horizontal facets represent different subsets of the *C. briggsae* isotypes. Colored backgrounds in nucleotide diversity panels delineate the chromosomal domain boundaries (purple, tip; blue, arm; yellow, center). Numbers in parentheses after each subset label show sample size.

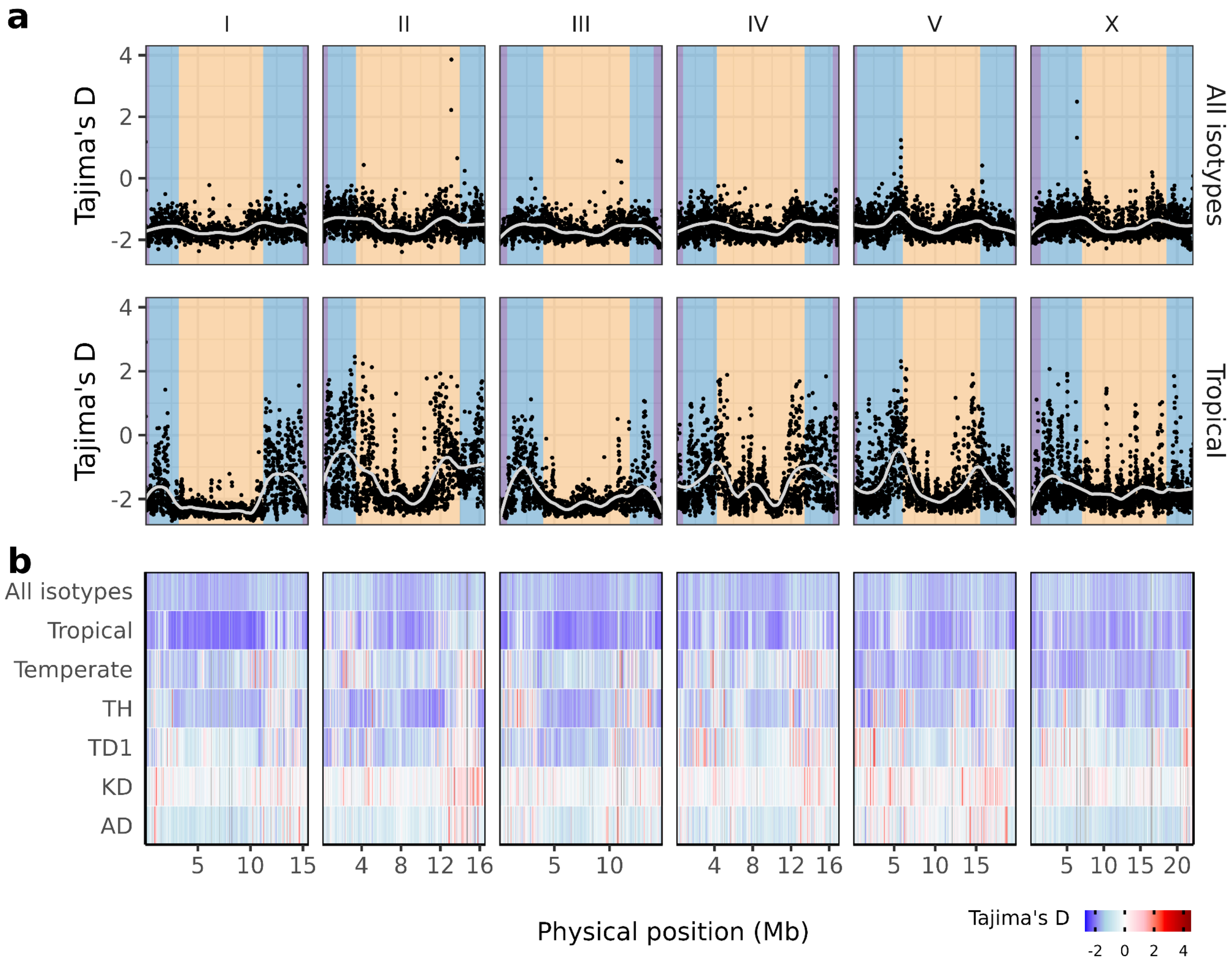


**Figure S21: Intermediate-frequency alleles near and at chromosomal arms across relatedness groups.** Estimates of Tajima's *D* in 10 kb bins shown as **a,** points across the genome (x-axis) for all isotypes and the Tropical relatedness group, and **b,** as heatmaps for all isotypes and each relatedness group with more than 10 isotype strains. Abbreviations: AD, Australia Divergent; KD, Kerala Divergent; TH, Taiwan-Hawaii; TD1, Taiwan Divergent 1.

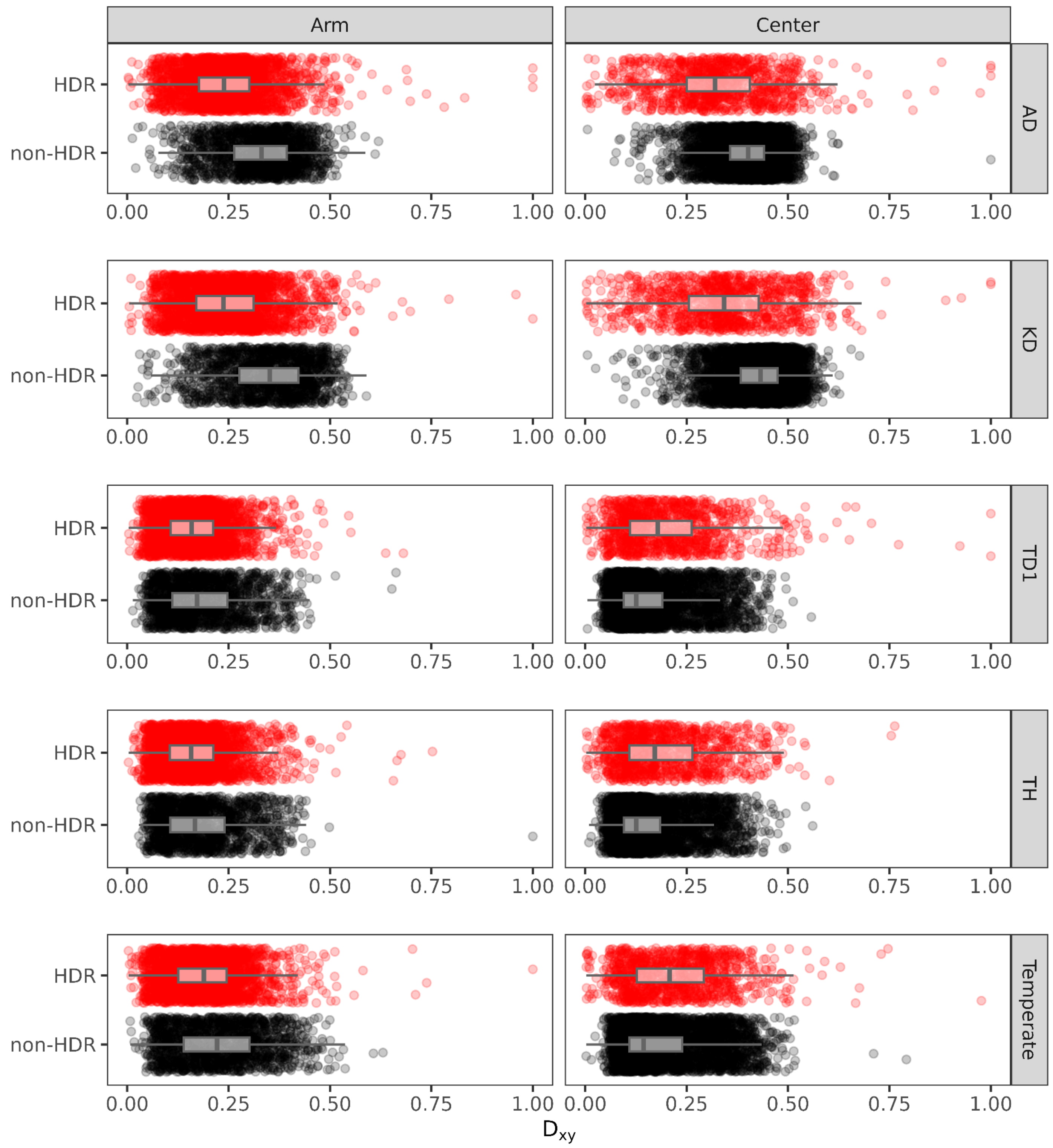


**Figure S22: Hyper-divergent regions display lower or no difference in absolute divergence across different relatedness group comparisons.** Boxplots of $D_{xy}$ estimates across 10 kb genome segments between Tropical and other relatedness groups comparing hyper-divergent (red) and non-hyper-divergent (black) regions. Each pairwise relatedness group comparison is shown as a separate facet.

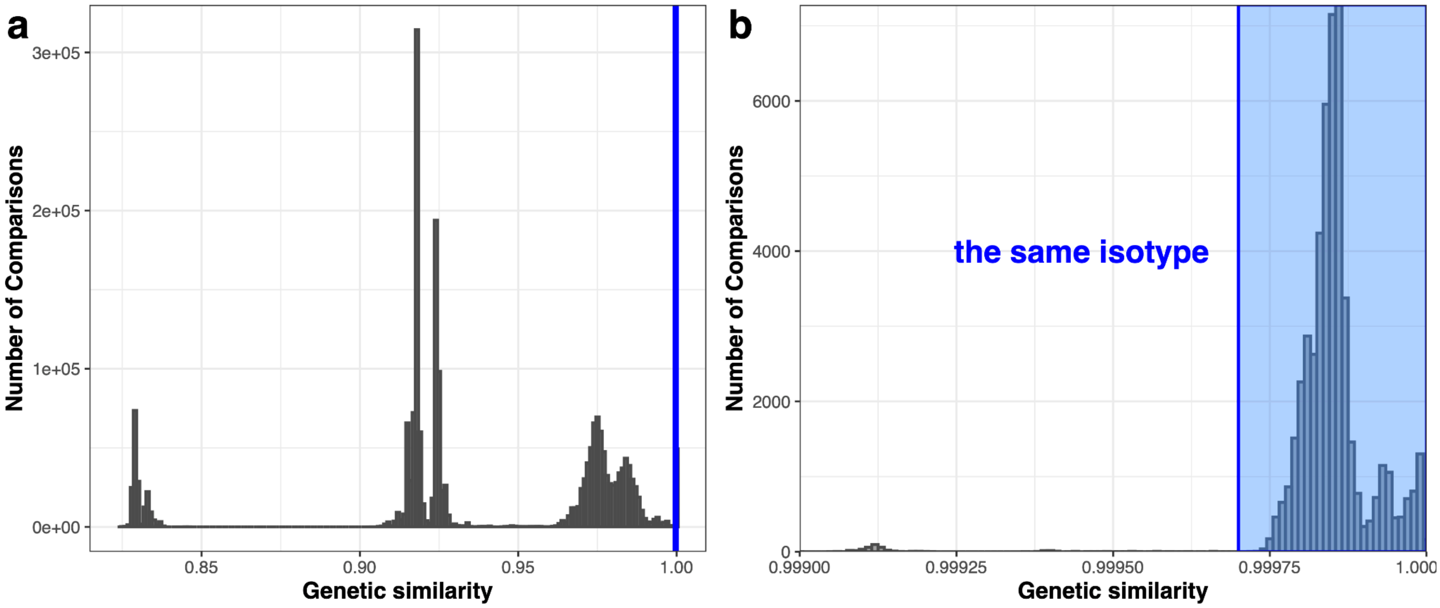


**Figure S23: Genetic similarity score distribution and isotype cutoff**. **a.** Histogram of genetic similarity for all pairwise comparisons. The vertical blue line indicates the threshold (99.97%) used to define isotype identity. **b,** Zoomed-in histogram of the high similarity range (>99.9%). The shaded blue area highlights comparisons classified as the same isotype.

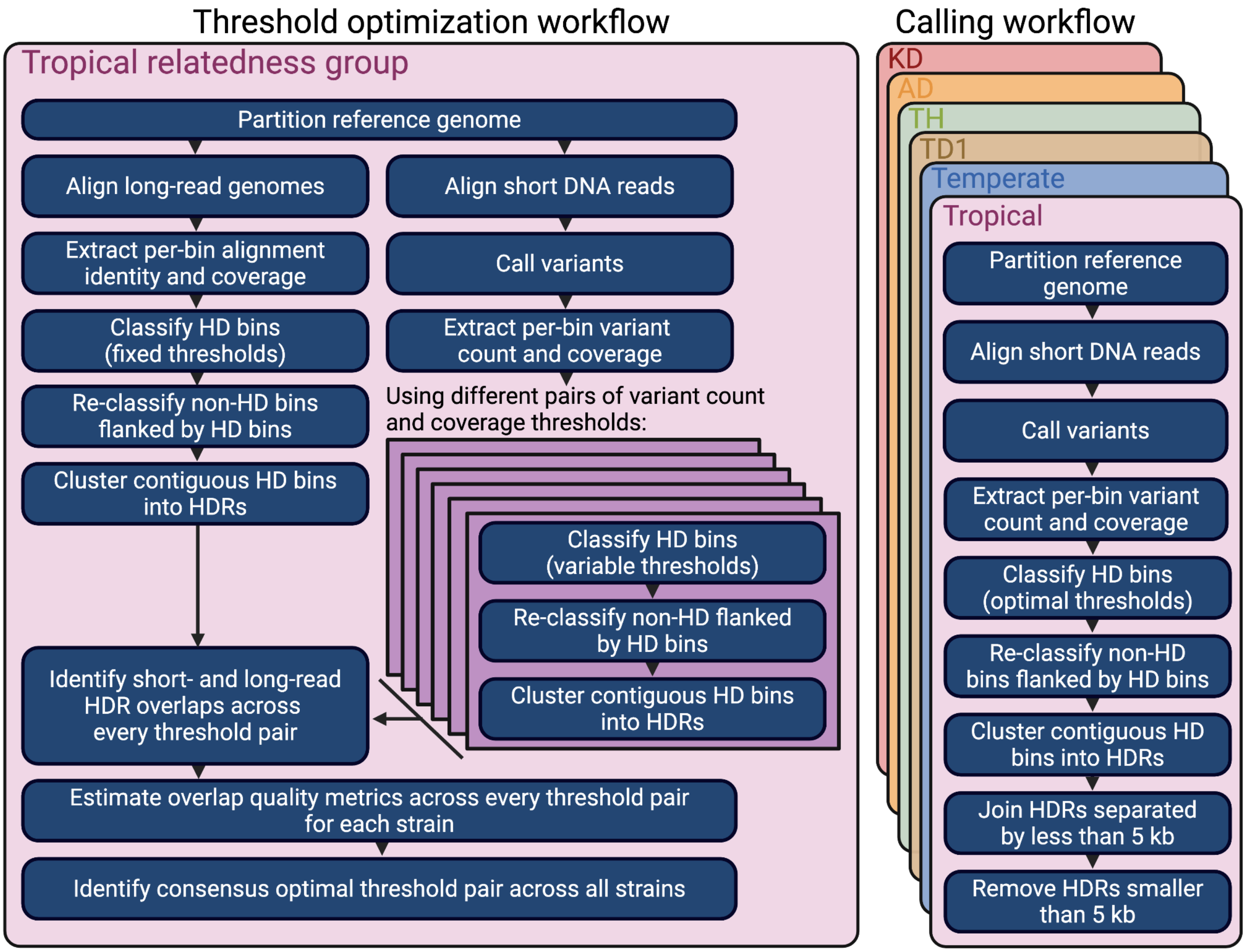


**Figure S24: Hyper-divergent region parameter optimization and calling workflows.** A schematic showing discrete steps in the workflow used to call hyper-divergent regions (HDRs). Variant count and coverage thresholds used to call HDRs species-wide are selected under the "Threshold optimization workflow" by comparing regions called using both long- and short-read genome alignments for 15 strains in the Tropical relatedness group. These optimized thresholds are then used in the "Calling workflow" to identify HDRs across all strains in each relatedness group using group-specific reference genomes, short-read genome alignments, and variant calls.

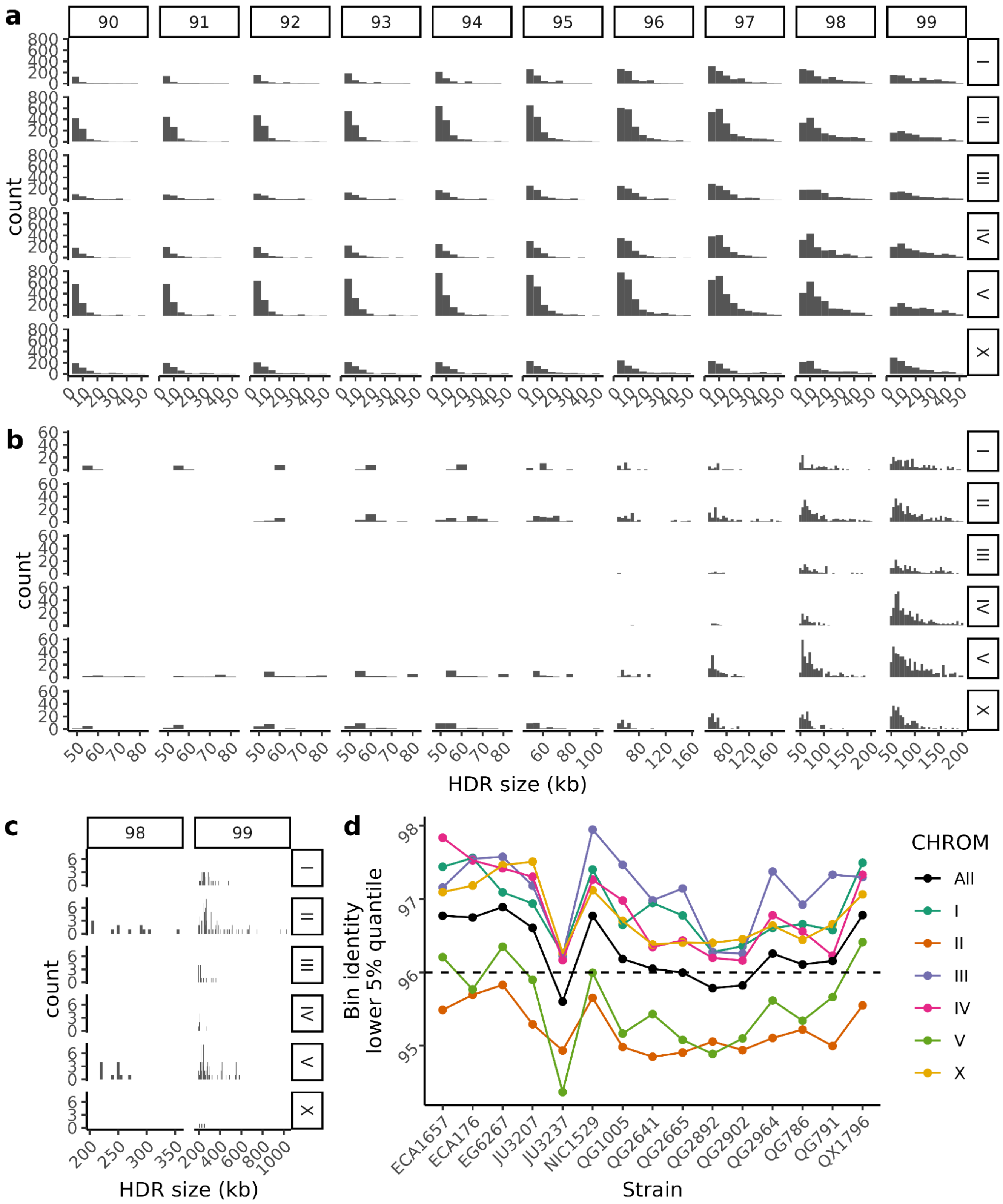


**Figure S25: Size and identity features of long-read based hyper-divergent regions at variable identity thresholds.** Counts of hyper-divergent regions across different identity thresholds (90-99%, represented on each vertical facet) across every chromosome (represented on each horizontal facet) for **a,** regions under 50 kb, **b,** regions over 50 kb and under 200 kb, and **c,** regions over 200 kb (only observed at identity thresholds 98% and 99%). **d,** for each reference Tropical strain (x-axis) and all chromosomes (black series) or each chromosome (each colored series), we estimated the lower 5% quantile of identity across all chromosome 1 kb bins (y-axis) and compared it against our selected identity threshold of 96% (dashed line).

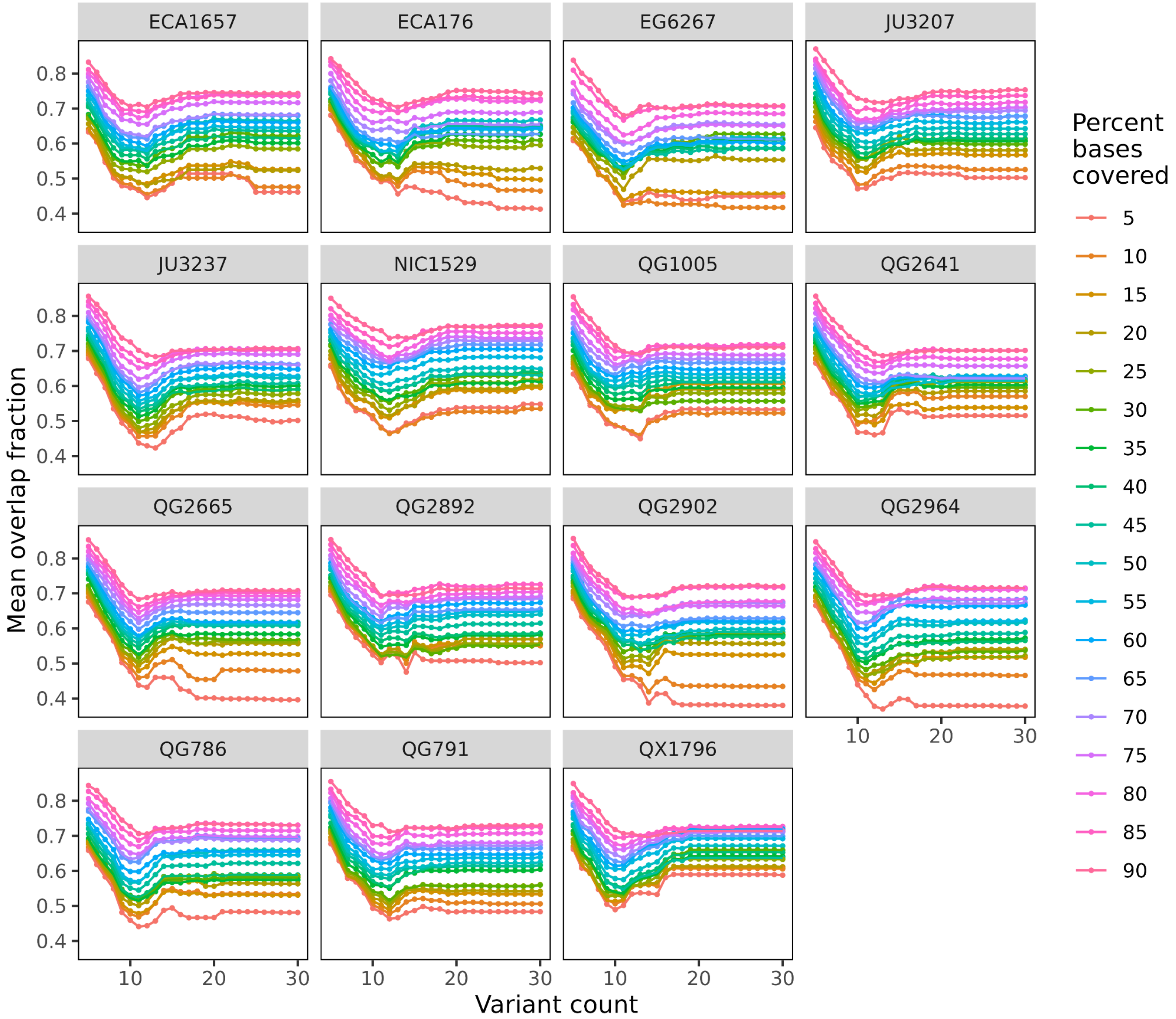


**Figure S26: Mean overlap fraction between short- and long-read HDR calls.** Estimates of hyper-divergent-region mean overlap fraction across 15 *C. briggsae* strains for every pair of variant count and percent bases covered thresholds. Overlap fraction is estimated by quantifying the extent of the overlaps between short- and long-read based hyper-divergent region calls, summarized into a mean overlap fraction estimate for each strain at each threshold pair. General trends indicate that mean overlap fraction improves at lower thresholds of variant count and higher thresholds of percent bases covered.

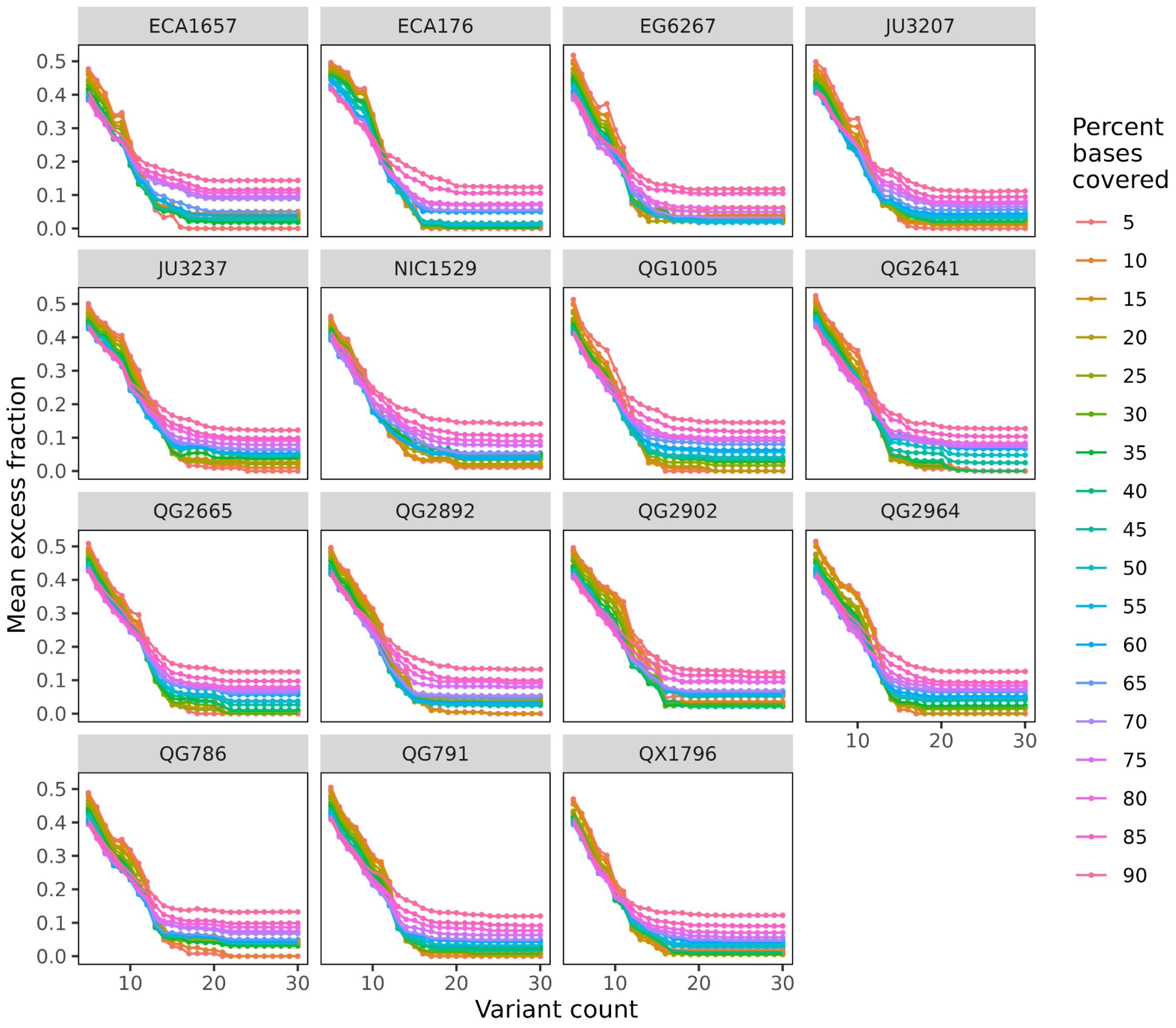


**Figure S27: Mean excess fraction between short- and long-read HDR calls.** Estimates of hyper-divergent region mean excess fraction across 15 *C. briggsae* strains for every pair of variant count and percent bases covered thresholds. Excess fraction is estimated by quantifying the extent of the short-read based hyper-divergent call that exceeds the boundaries of an overlapping long-read hyper-divergent region calls, summarized into a mean excess fraction estimate for each strain at each threshold pair. General trends indicate that mean excess fraction drastically improves at higher thresholds of variant count, with a more subtle improvement at higher thresholds of percent bases covered.

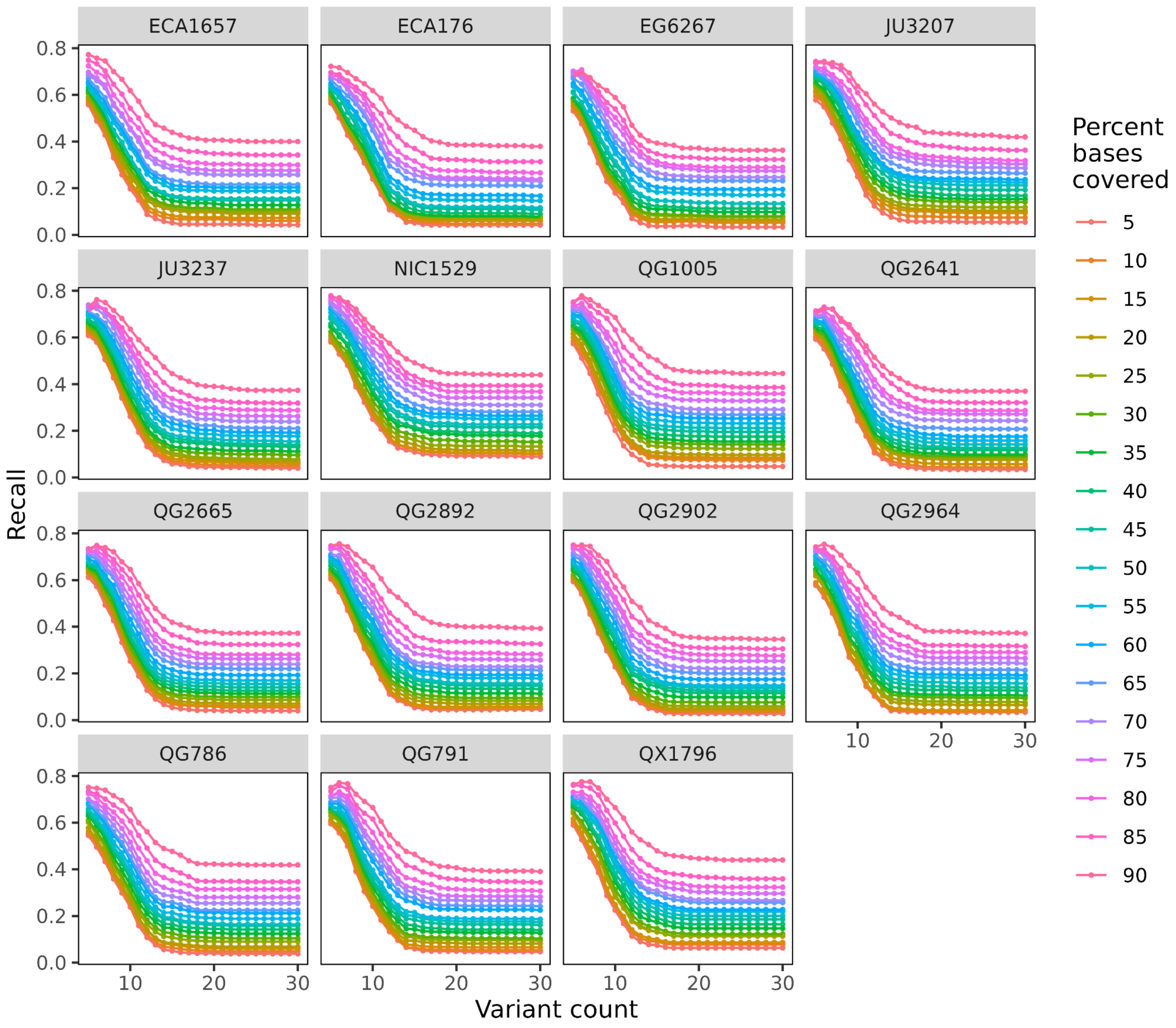


**Figure S28: Recall of long-read HDR calls.** Estimates of hyper-divergent region recall across 15 *C. briggsae* strains for every pair of variant count and percent bases covered thresholds. Recall is estimated by the proportion between the number of long-read based hyper-divergent region calls that have an overlap with short-read based hyper-divergent region calls relative to the total number of long-read based hyper-divergent region calls. Similar to mean overlap fraction, general trends indicate that recall improves at lower thresholds of variant count and higher thresholds of percent bases covered.

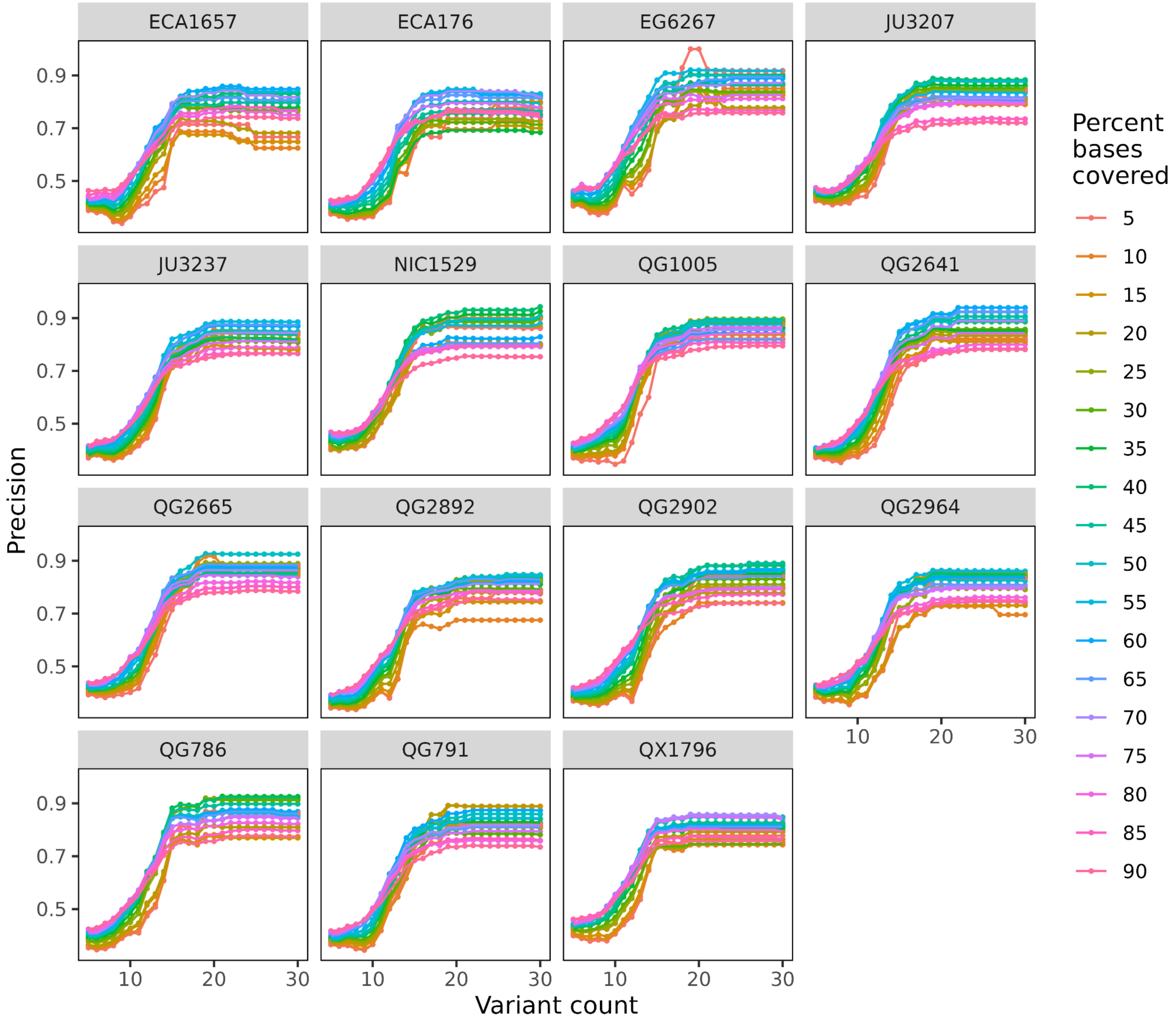


**Figure S29: Precision of short-read HDR calls.** Estimates of hyper-divergent region precision across 15 *C. briggsae* strains for every pair of variant count and percent bases covered thresholds. Precision is estimated by the proportion between the number of short-read based hyper-divergent region calls that have an overlap with long-read based hyper-divergent region calls relative to the total number of short-read based hyper-divergent region calls. Similar to mean excess fraction, general trends indicate that precision drastically improves at higher thresholds of variant count, with a more subtle improvement at higher thresholds of percent bases covered.

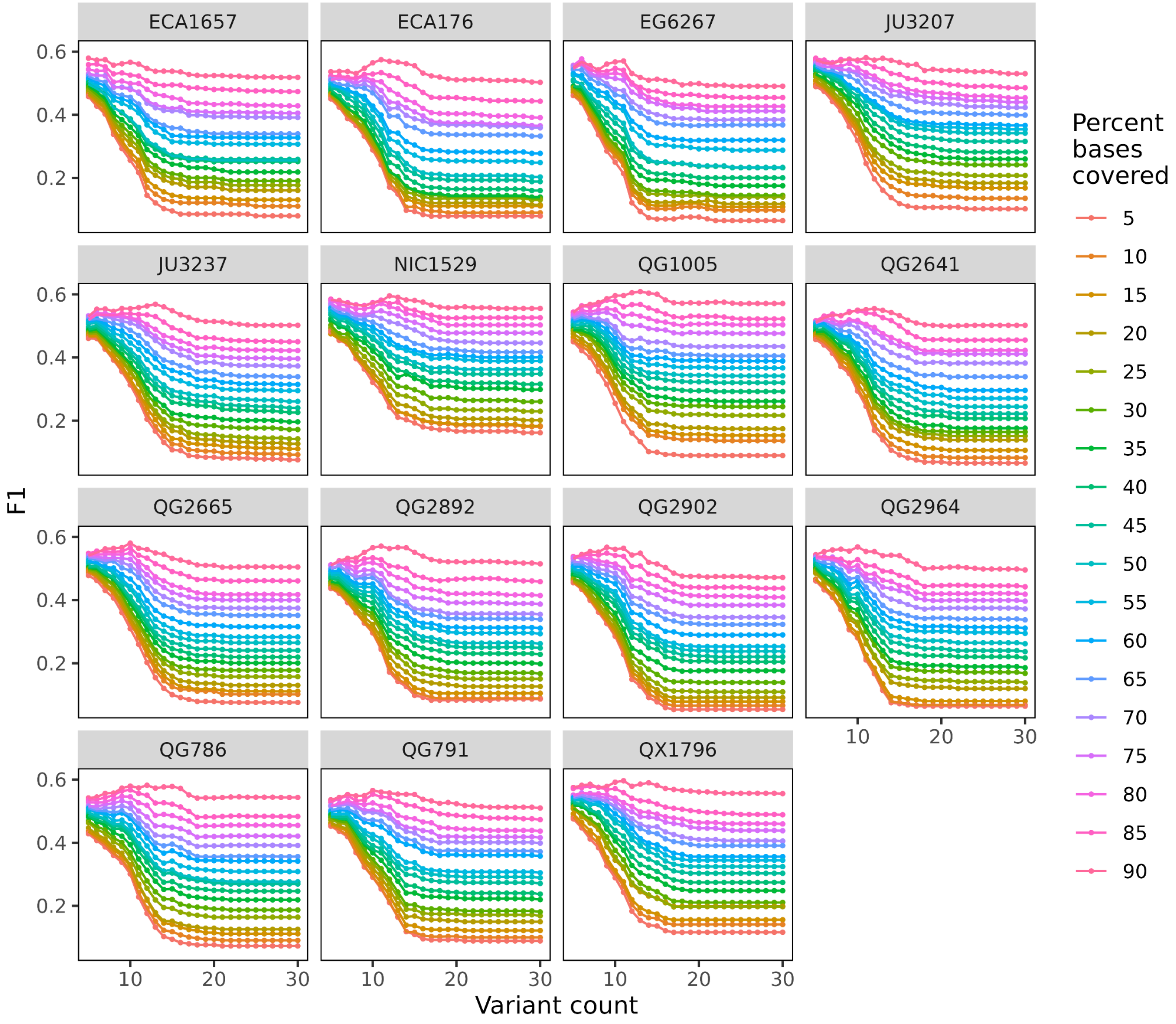


**Figure S30: F1 of short-read HDR calls.** Estimates of hyper-divergent region F1 across 15 *C. briggsae* strains for every pair of variant count and percent bases covered thresholds. F1 is estimated from the harmonic mean between recall and precision at each parameter pair.

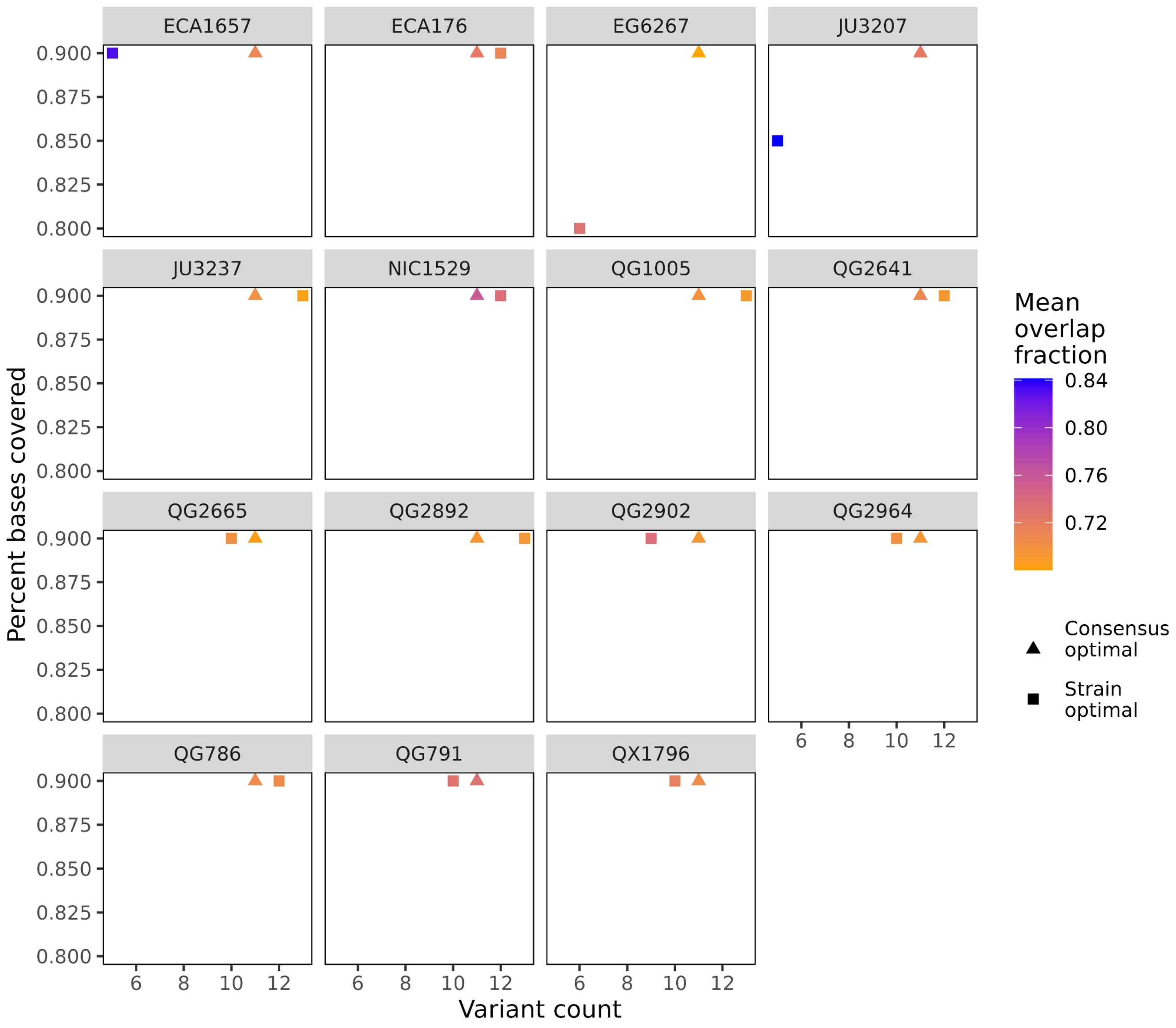


**Figure S31: Strain and consensus optimal threshold pairs for HDR calls.** A limited scatter plot showing the best threshold pair optimized by F1 for each individual strain (displayed as squares, referred to as 'strain optimal'). By selecting the 10 threshold pairs with the highest F1 in each strain, we identified a common threshold pair across all 15 strains (displayed as triangles, referred to as 'consensus optimal'). A gradient color scale describing the mean overlap fraction was applied to each threshold pair. The consensus optimal threshold pair (11 SNV, 90% bases covered) was used to call hyper-divergent regions species-wide.

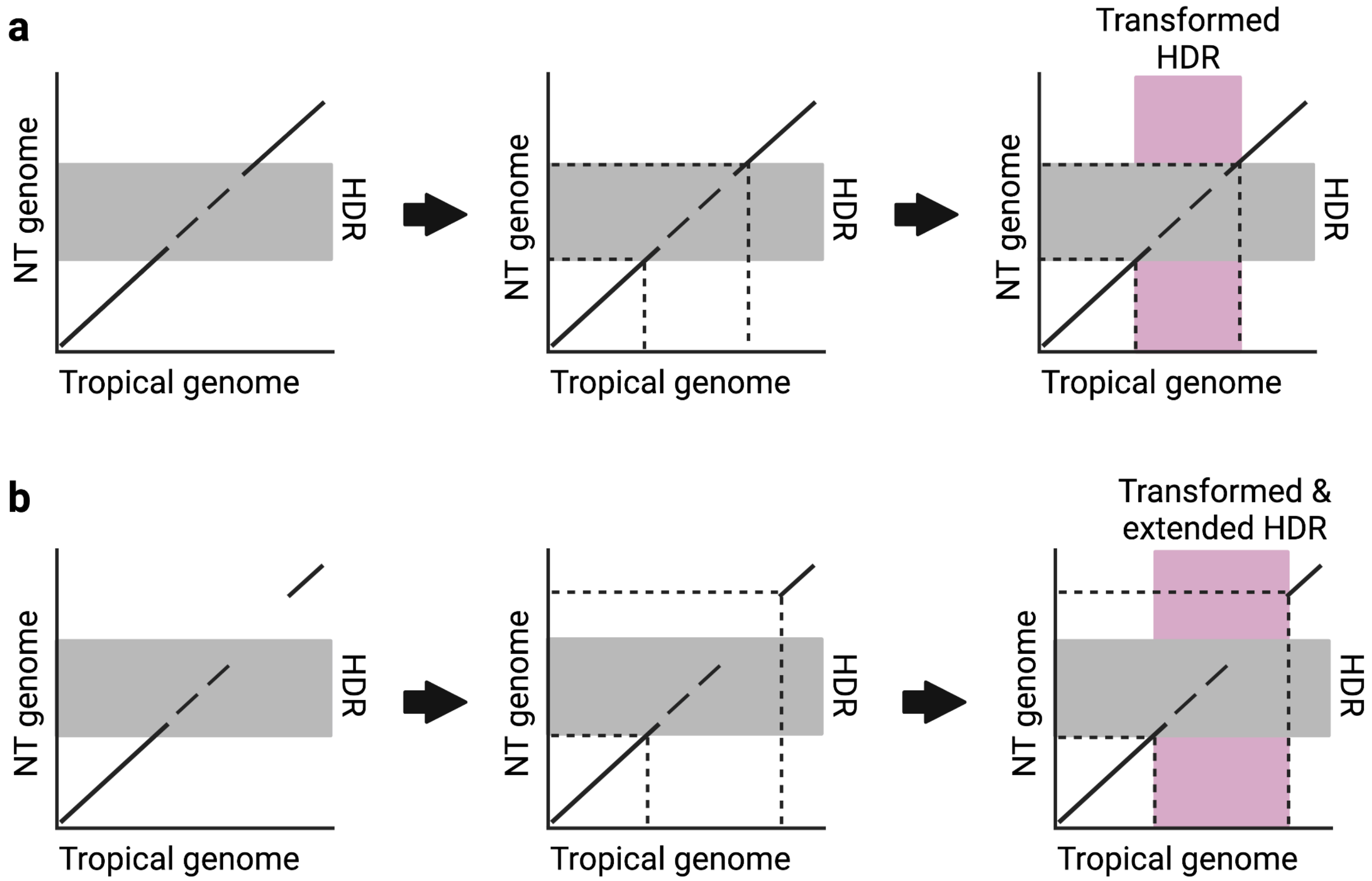


**Figure S32: Mapping hyper-divergent region calls across relatedness group reference genomes.** Illustration of steps to map coordinates of hyper-divergent regions (HDRs) between non-Tropical (NT) and Tropical relatedness group reference genomes using genome alignments. The y-axis in each plot represents the genome coordinates of a NT relatedness group reference genome, and the x-axis represents the genome coordinates of the Tropical relatedness group reference genome. Alignments between Tropical and NT relatedness group reference genomes are shown as diagonal lines. HDRs for each NT strain are called relative to their respective NT reference genome, and a hypothetical HDR is shown as a gray box. The boundaries of the HDR mapped to the Tropical relatedness group reference genome are estimated directly from **a,** single alignments spanning the NT hyper-divergent region boundaries or **b,** extended to single alignments proximal (<50kb) to the NT HDR boundaries when spanning alignments are not available because of large structural differences between the Tropical and NT genomes.
