## Supplementary File 2 for "*Caenorhabditis briggsae* ancestral genomic hyper-diversity contrasts with globally distributed genome-wide haplotypes"

**Potential introgression events in hyper-divergent regions between *C. briggsae* and *C. nigoni*.**

Each figure displays an approximately-maximum-likelihood tree modeled from protein sequence alignments of single-copy orthologs between Tropical *C. briggsae* strains and all sequenced *C. nigoni* lines that lack two distinct monophyletic groups (one for each species). Protein sequences were aligned with *famsa*[1] v2.4.1-45c9b2b and modeled using *fasttree*[2] v2.1.11 within the OrthoFinder[3] v3.1.0 pipeline using default parameters. Nodes are labeled after their respective strain and colored by species (*C. nigoni*, orange; *C. briggsae*, blue). The reference gene (from the QX1410 strain) for each tree is found within a hyper-divergent region. *C. briggsae* nodes have distinct shapes depending on whether they are found within a hyper-divergent haplotype (squares) or the reference haplotype (circles). *C. nigoni* nodes are shown as diamonds (unknown), indicating that the haplotype status has not been assessed. The trees are accompanied by amino acid identity heatmaps with matching strain labels and clustered by amino acid similarity. The top band at the bottom of each heatmap indicates whether a *C. briggsae* sequence is found within a hyper-divergent haplotype (red) or the reference haplotype (black); not applicable (grey) for *C. nigoni* sequences. The bottom band at the bottom of each heatmap indicates the species of each sequence. A total of 60 orthologous groups are shown. The orthologous group ID, gene alias, and coordinates of the reference gene are shown at the top of each figure. Orthogroup and gene IDs can be accessed in Supplemental File 1.

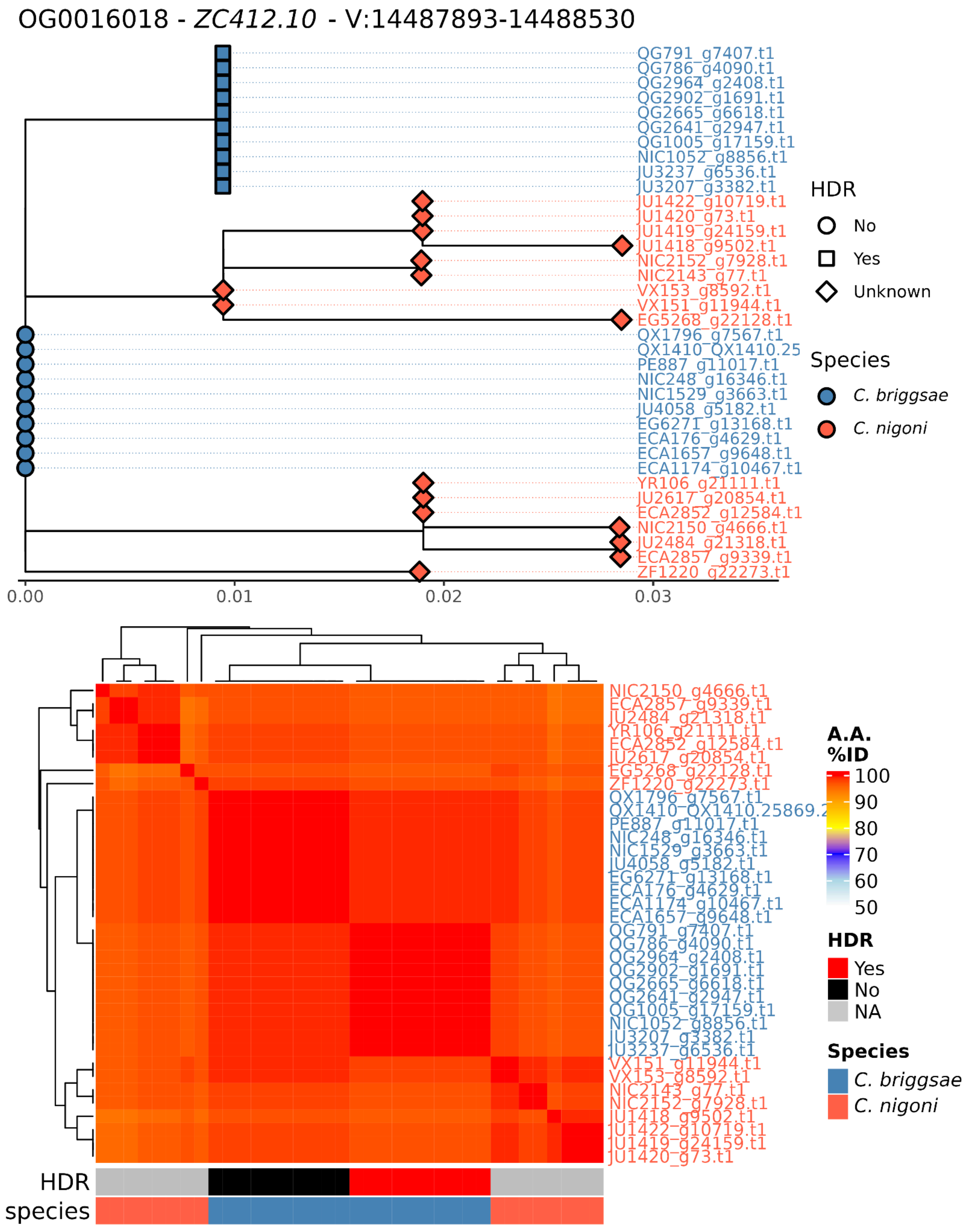
OG0016018 - *ZC412.10* - V:14487893-14488530
QG791_g7407.t1
QG786_g4090.t1
QG2964_g2408.t1
QG2902_g1691.t1
QG2665_g6618.t1
QG2641_g2947.t1
QG1005_g17159.t1
NIC1052_g8856.t1
JU3237_g6536.t1
JU3207_g3382.t1
JU1422_g10719.t1
JU1420_g73.t1
JU1419_g24159.t1
JU1418_g9502.t1
NIC2152_g7928.t1
NIC2143_g77.t1
VX153_g8592.t1
VX151_g11944.t1
EG5268_g22128.t1
QX1796_g7567.t1
QX1410_QX1410.25
PE887_g11017.t1
NIC248_g16346.t1
NIC1529_g3663.t1
JU4058_g5182.t1
EG6271_g13168.t1
ECA176_g4629.t1
ECA1657_g9648.t1
ECA1174_g10467.t1
YR106_g21111.t1
JU2617_g20854.t1
ECA2852_g12584.t1
NIC2150_g4666.t1
JU2484_g21318.t1
ECA2857_g9339.t1
ZF1220_g22273.t1
0.00
0.01
0.02
0.03
HDR
No
Yes
Unknown
Species
*C. briggsae*
*C. nigoni*
NIC2150_g4666.t1
ECA2857_g9339.t1
JU2484_g21318.t1
YR106_g21111.t1
ECA2852_g12584.t1
JU2617_g20854.t1
EG5268_g22128.t1
ZF1220_g22273.t1
QX1796_g7567.t1
QX1410_QX1410.25869.2
PE887_g11017.t1
NIC248_g16346.t1
NIC1529_g3663.t1
JU4058_g5182.t1
EG6271_g13168.t1
ECA176_g4629.t1
ECA1174_g10467.t1
ECA1657_g9648.t1
QG791_g7407.t1
QG786_g4090.t1
QG2964_g2408.t1
QG2902_g1691.t1
QG2665_g6618.t1
QG2641_g2947.t1
QG1005_g17159.t1
NIC1052_g8856.t1
JU3207_g3382.t1
JU3237_g6536.t1
VX151_g11944.t1
VX153_g8592.t1
NIC2143_g77.t1
NIC2152_g7928.t1
JU1418_g9502.t1
JU1422_g10719.t1
JU1419_g24159.t1
JU1420_g73.t1
**A.A. %ID**
100
90
80
70
60
50
**HDR**
Yes
No
NA
**Species**
*C. briggsae*
*C. nigoni*
HDR
species

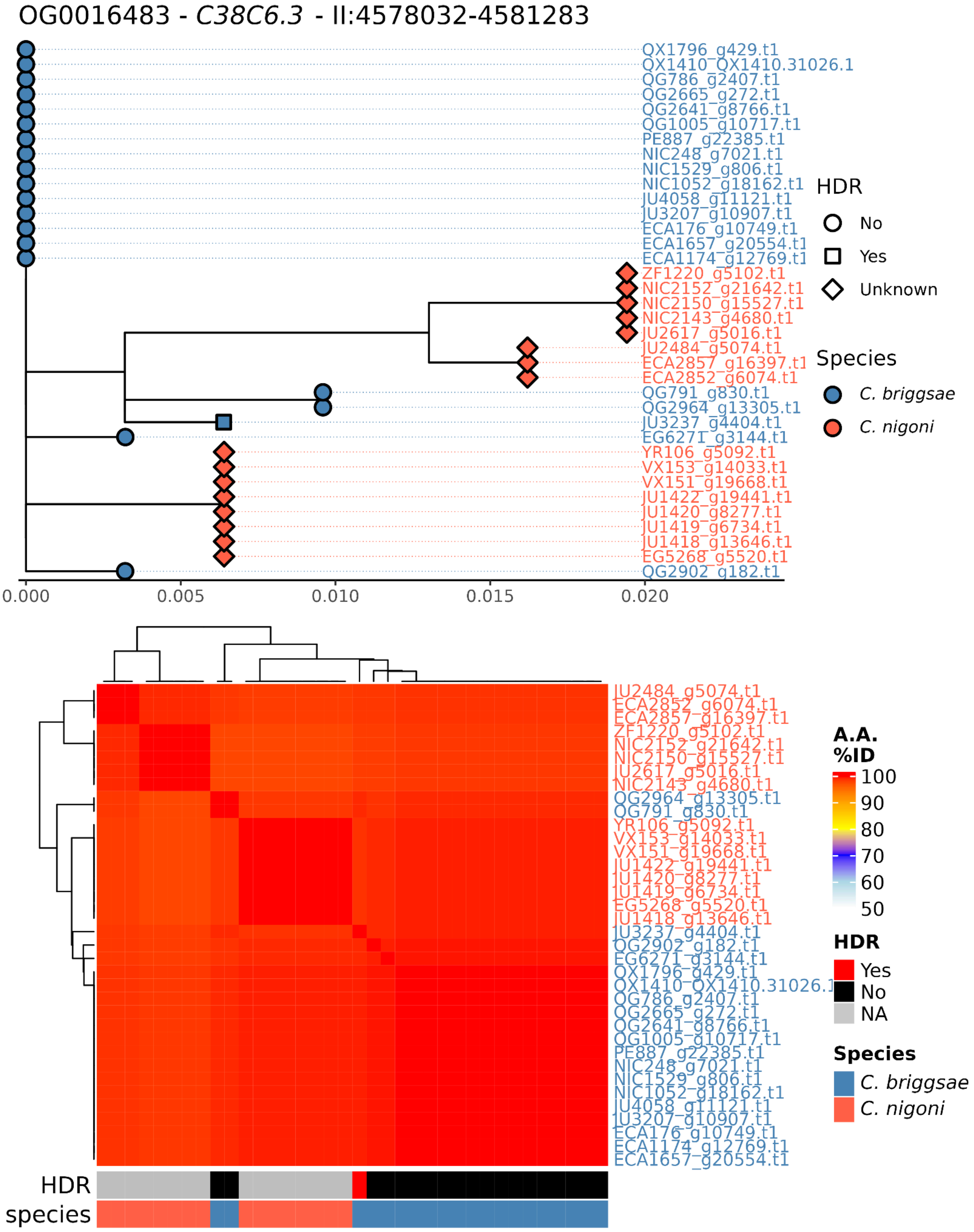

OG0016483 - *C38C6.3* - II:4578032-4581283
QX1796_g429.t1
QX1410_QX1410.31026.1
QG786_g2407.t1
QG2665_g272.t1
QG2641_g8766.t1
QG1005_g10717.t1
PE887_g22385.t1
NIC248_g7021.t1
NIC1529_g806.t1
NIC1052_g18162.t1
JU4058_g11121.t1
JU3207_g10907.t1
ECA176_g10749.t1
ECA1657_g20554.t1
ECA1174_g12769.t1
ZF1220_g5102.t1
NIC2152_g21642.t1
NIC2150_g15527.t1
NIC2143_g4680.t1
JU2617_g5016.t1
JU2484_g5074.t1
ECA2857_g16397.t1
ECA2852_g6074.t1
QG791_g830.t1
QG2964_g13305.t1
JU3237_g4404.t1
EG6271_g3144.t1
YR106_g5092.t1
VX153_g14033.t1
VX151_g19668.t1
JU1422_g19441.t1
JU1420_g8277.t1
JU1419_g6734.t1
JU1418_g13646.t1
EG5268_g5520.t1
QG2902_g182.t1
0.000
0.005
0.010
0.015
0.020
HDR
No
Yes
Unknown
Species
C. briggsae
C. nigoni
A.A. %ID
100
90
80
70
60
50
HDR
Yes
No
NA
Species
C. briggsae
C. nigoni
HDR
species

OG0020710 - *QX1410.33333* - II:14522471-14523231

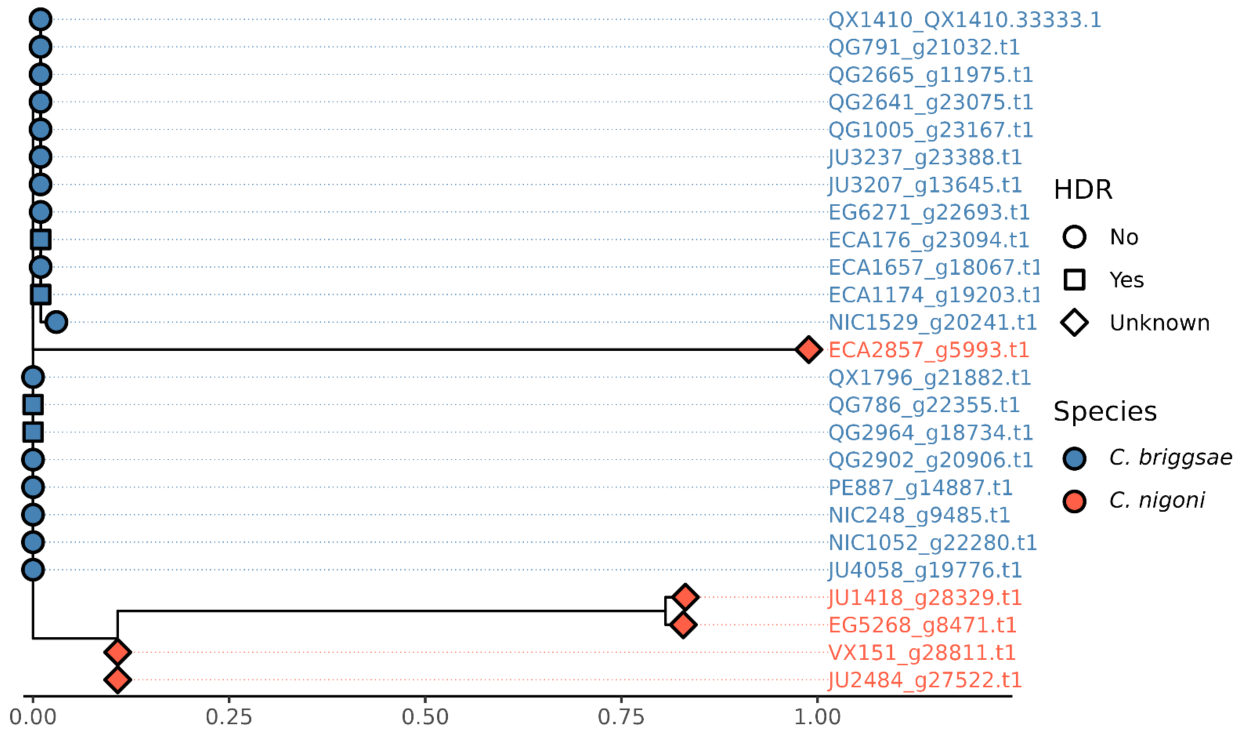


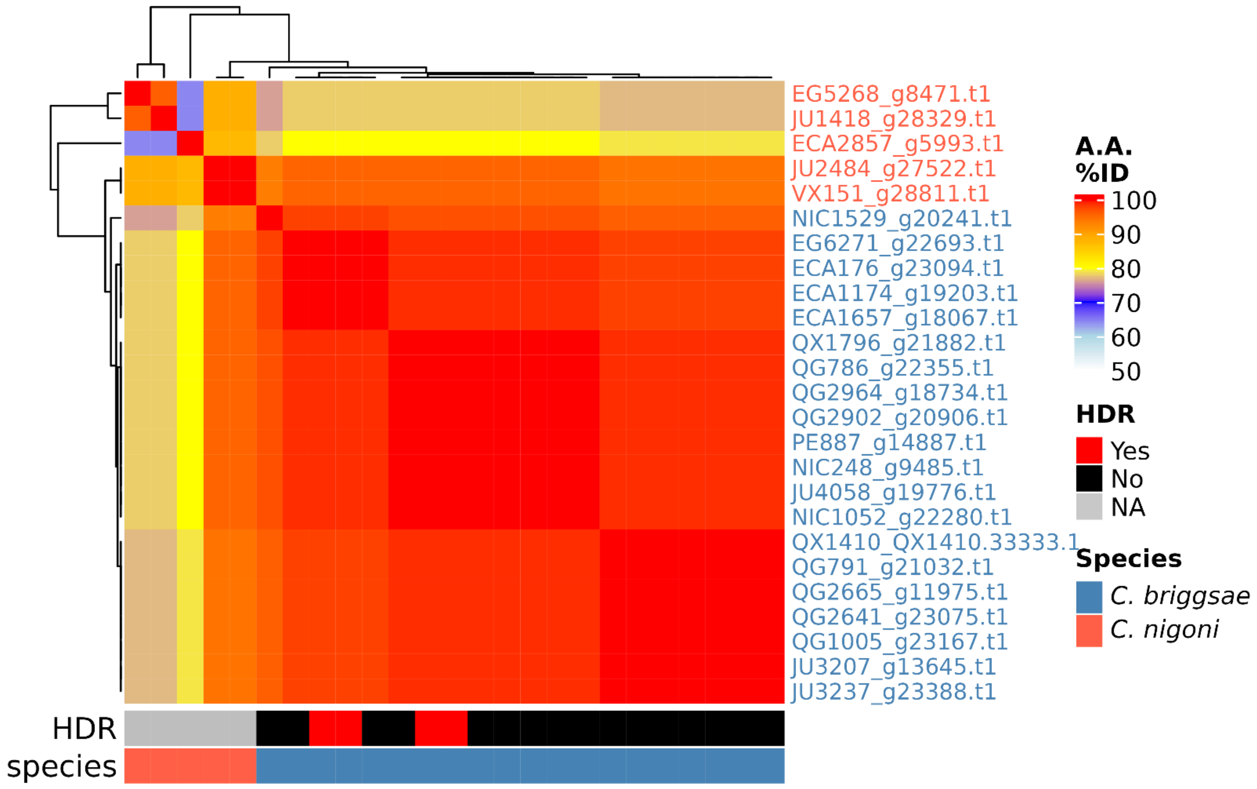

OG0021359 - *CBG29897* - I:13178058-13179269

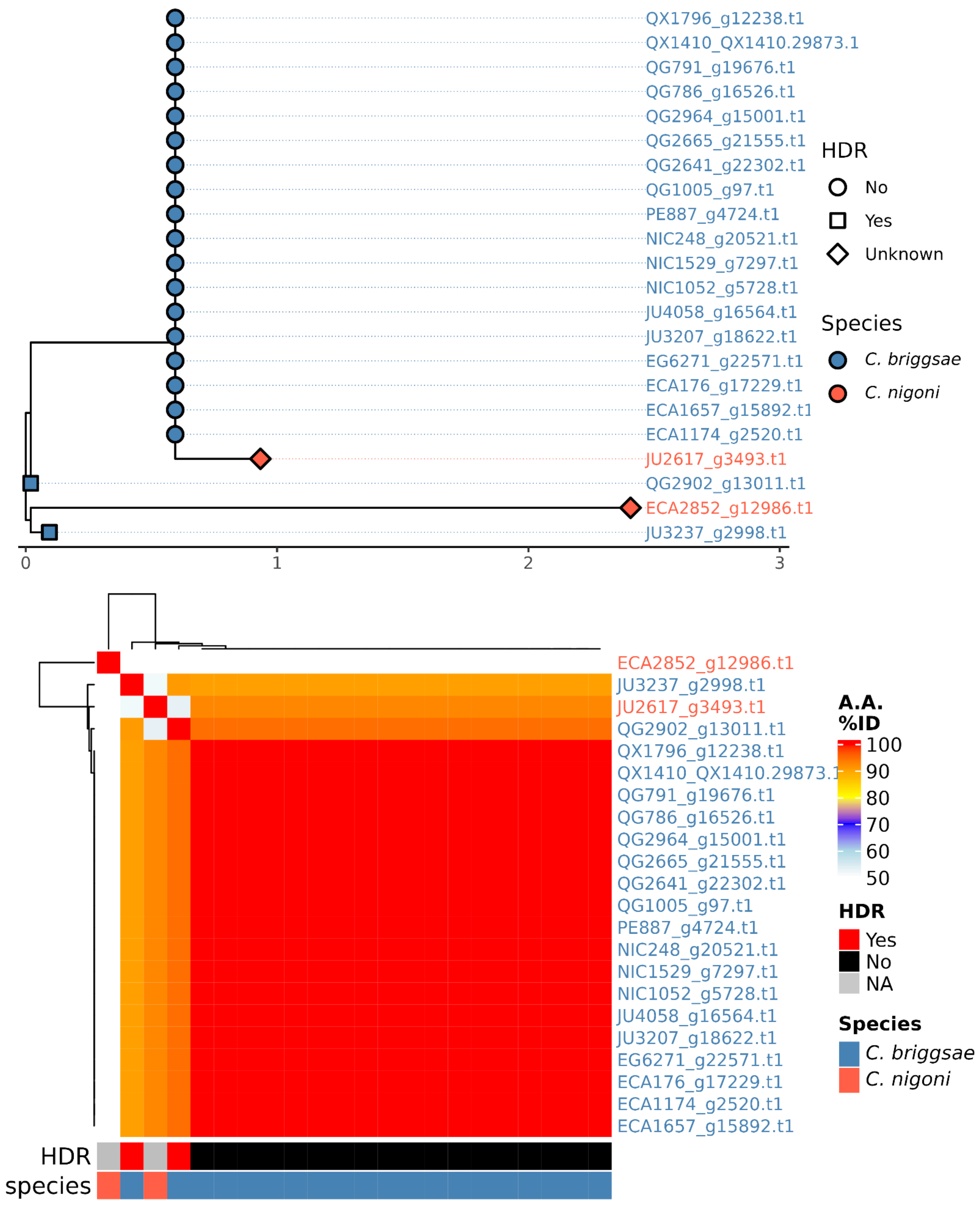

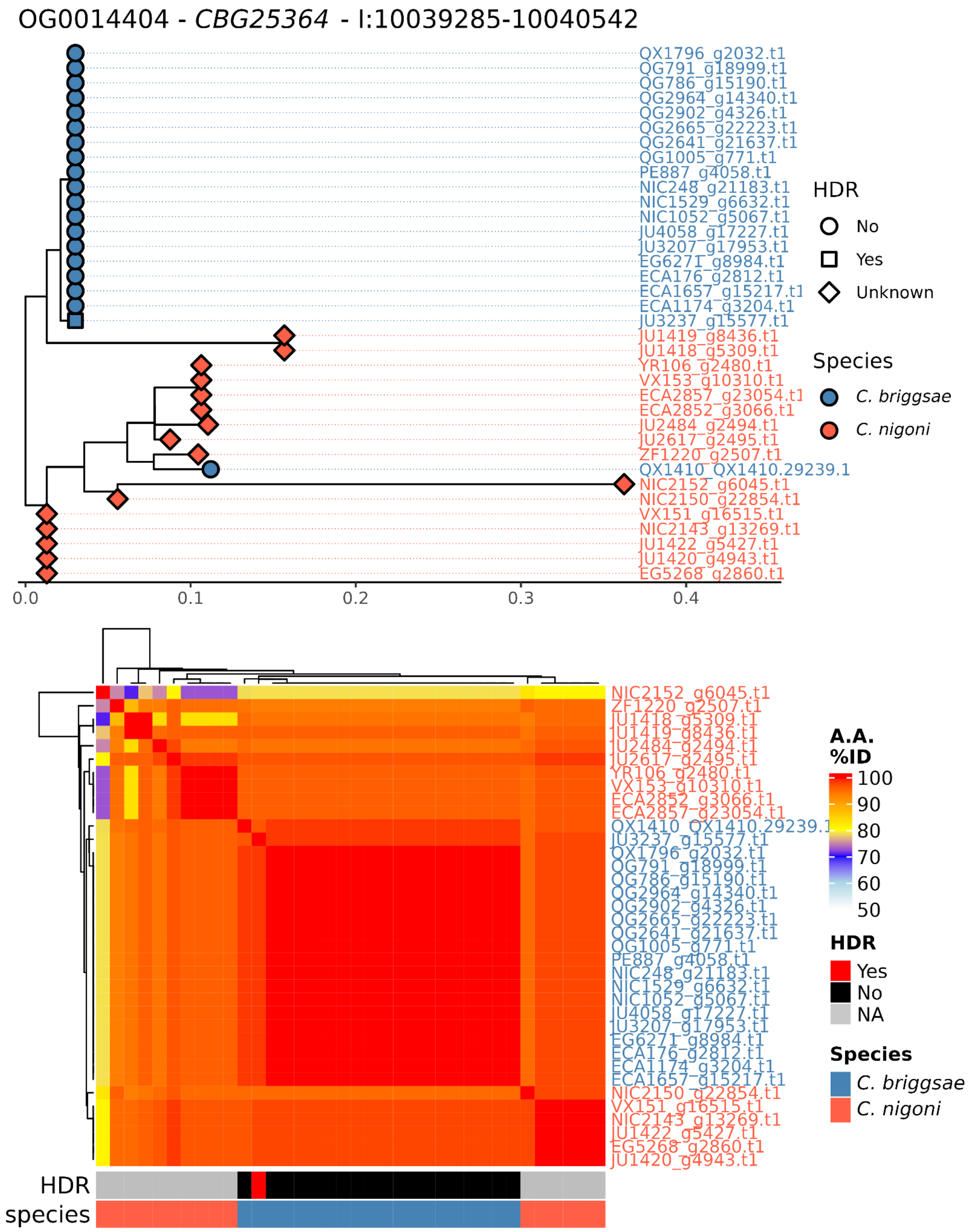

OG0014404 - *CBG25364* - I:10039285-10040542
QX1796_g2032.t1
QG791_g18999.t1
QG786_g15190.t1
QG2964_g14340.t1
QG2902_g4326.t1
QG2665_g22223.t1
QG2641_g21637.t1
QG1005_g771.t1
PE887_g4058.t1
NIC248_g21183.t1
NIC1529_g6632.t1
NIC1052_g5067.t1
JU4058_g17227.t1
JU3207_g17953.t1
EG6271_g8984.t1
ECA176_g2812.t1
ECA1657_g15217.t1
ECA1174_g3204.t1
JU3237_g15577.t1
JU1419_g8436.t1
JU1418_g5309.t1
YR106_g2480.t1
VX153_g10310.t1
ECA2857_g23054.t1
ECA2852_g3066.t1
JU2484_g2494.t1
JU2617_g2495.t1
ZF1220_g2507.t1
QX1410_QX1410.29239.1
NIC2152_g6045.t1
NIC2150_g22854.t1
VX151_g16515.t1
NIC2143_g13269.t1
JU1422_g5427.t1
JU1420_g4943.t1
EG5268_g2860.t1
0.0
0.1
0.2
0.3
0.4
HDR
No
Yes
Unknown
Species
C. briggsae
C. nigoni
NIC2152_g6045.t1
ZF1220_g2507.t1
JU1418_g5309.t1
JU1419_g8436.t1
JU2484_g2494.t1
JU2617_g2495.t1
YR106_g2480.t1
VX153_g10310.t1
ECA2852_g3066.t1
ECA2857_g23054.t1
QX1410_QX1410.29239.1
JU3237_g15577.t1
QX1796_g2032.t1
QG791_g18999.t1
QG786_g15190.t1
QG2964_g14340.t1
QG2902_g4326.t1
QG2665_g22223.t1
QG2641_g21637.t1
QG1005_g771.t1
PE887_g4058.t1
NIC248_g21183.t1
NIC1529_g6632.t1
NIC1052_g5067.t1
JU4058_g17227.t1
JU3207_g17953.t1
EG6271_g8984.t1
ECA176_g2812.t1
ECA1174_g3204.t1
ECA1657_g15217.t1
NIC2150_g22854.t1
VX151_g16515.t1
NIC2143_g13269.t1
JU1422_g5427.t1
EG5268_g2860.t1
JU1420_g4943.t1
A.A. %ID
100
90
80
70
60
50
HDR
Yes
No
NA
Species
C. briggsae
C. nigoni
HDR
species

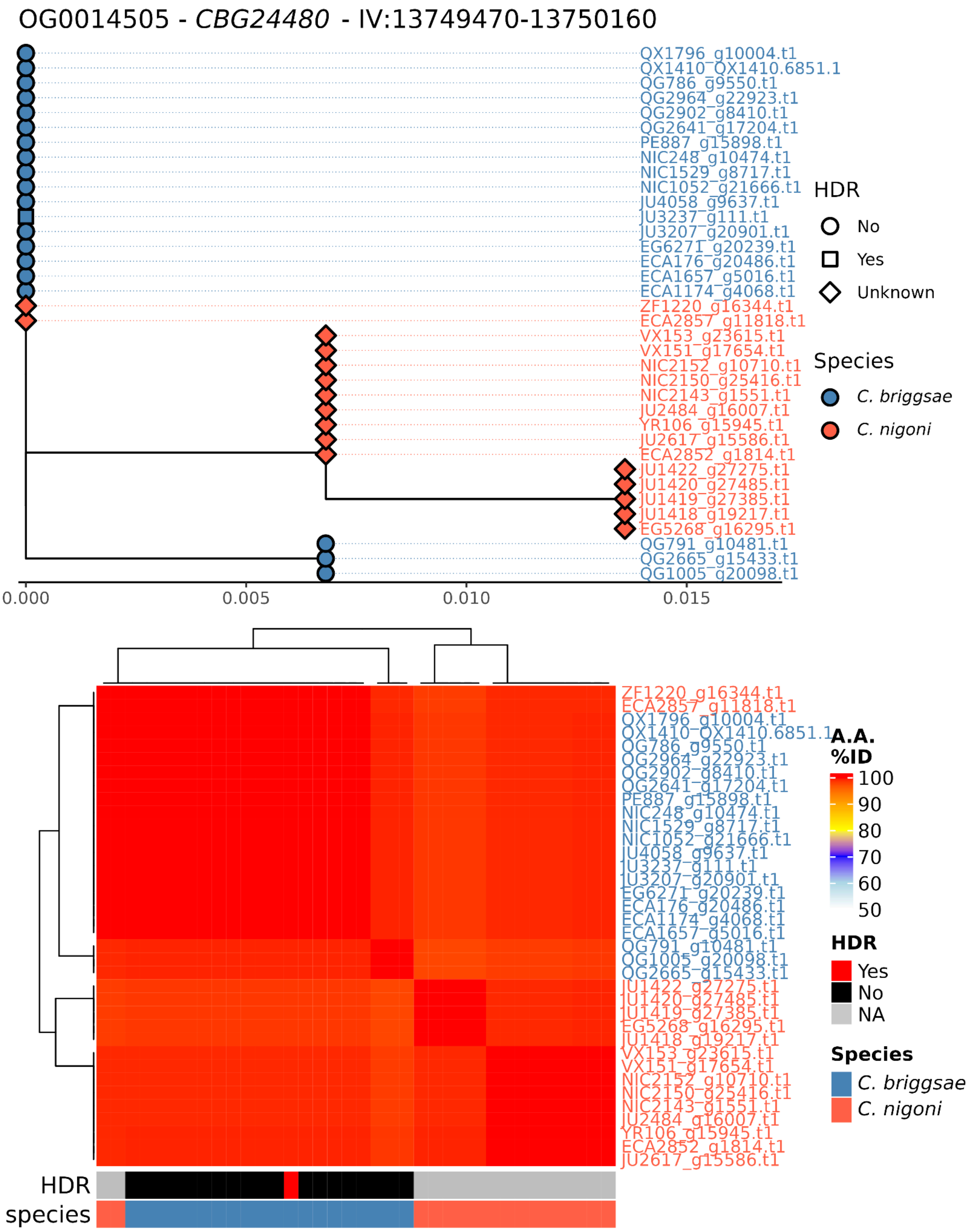
OG0014505 - CBG24480 - IV:13749470-13750160
QX1796_g10004.t1
QX1410_QX1410.6851.1
QG786_g9550.t1
QG2964_g22923.t1
QG2902_g8410.t1
QG2641_g17204.t1
PE887_g15898.t1
NIC248_g10474.t1
NIC1529_g8717.t1
NIC1052_g21666.t1
JU4058_g9637.t1
JU3237_g111.t1
JU3207_g20901.t1
EG6271_g20239.t1
ECA176_g20486.t1
ECA1657_g5016.t1
ECA1174_g4068.t1
ZF1220_g16344.t1
ECA2857_g11818.t1
VX153_g23615.t1
VX151_g17654.t1
NIC2152_g10710.t1
NIC2150_g25416.t1
NIC2143_g1551.t1
JU2484_g16007.t1
YR106_g15945.t1
JU2617_g15586.t1
ECA2852_g1814.t1
JU1422_g27275.t1
JU1420_g27485.t1
JU1419_g27385.t1
JU1418_g19217.t1
EG5268_g16295.t1
QG791_g10481.t1
QG2665_g15433.t1
QG1005_g20098.t1
0.000
0.005
0.010
0.015
HDR
No
Yes
Unknown
Species
C. briggsae
C. nigoni
A.A. %ID
100
90
80
70
60
50
HDR
Yes
No
NA
Species
C. briggsae
C. nigoni
HDR
species

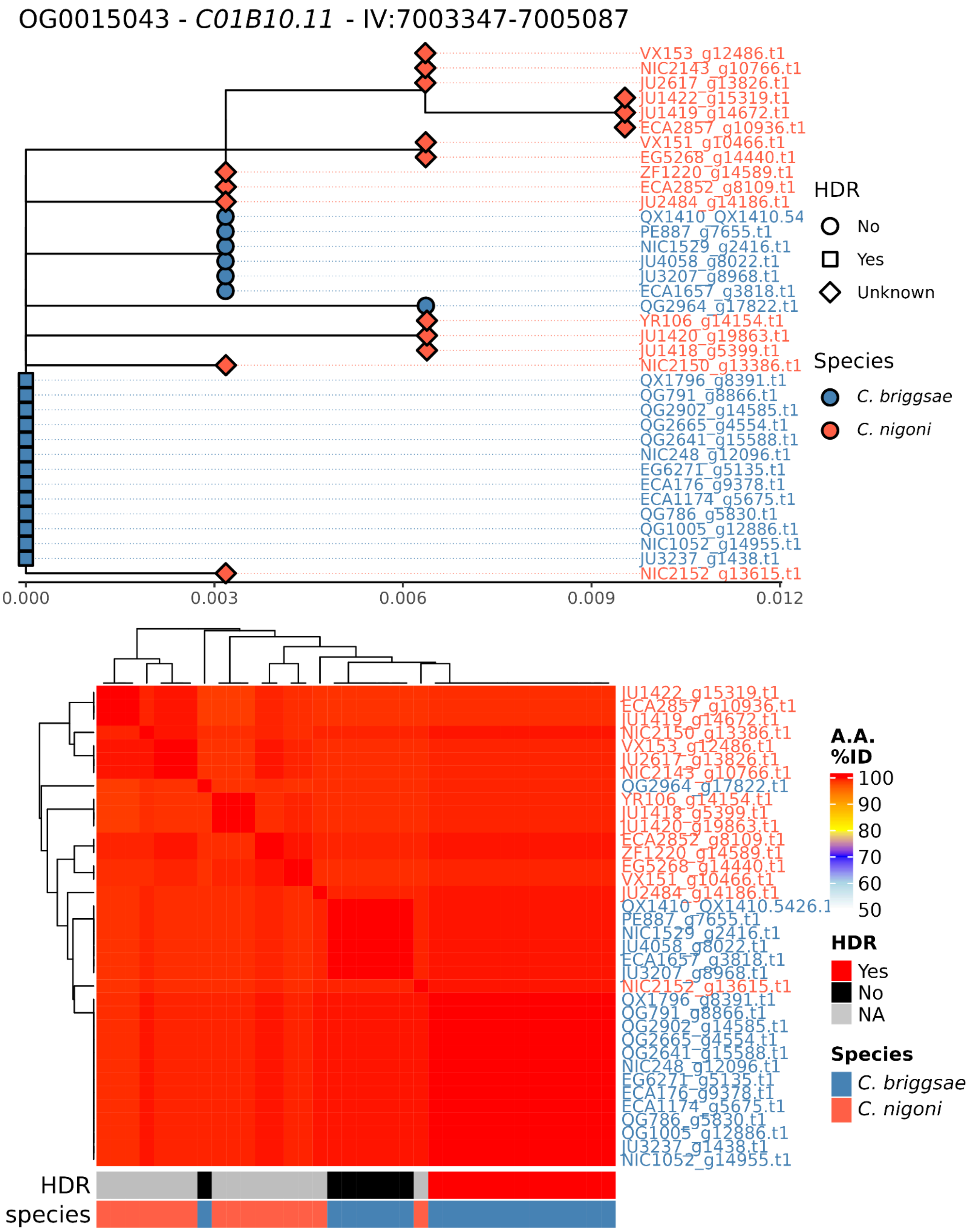
OG0015043 - C01B10.11 - IV:7003347-7005087
VX153_g12486.t1
NIC2143_g10766.t1
JU2617_g13826.t1
JU1422_g15319.t1
JU1419_g14672.t1
ECA2857_g10936.t1
VX151_g10466.t1
EG5268_g14440.t1
ZF1220_g14589.t1
ECA2852_g8109.t1
JU2484_g14186.t1
QX1410_QX1410.54
PE887_g7655.t1
NIC1529_g2416.t1
JU4058_g8022.t1
JU3207_g8968.t1
ECA1657_g3818.t1
QG2964_g17822.t1
YR106_g14154.t1
JU1420_g19863.t1
JU1418_g5399.t1
NIC2150_g13386.t1
QX1796_g8391.t1
QG791_g8866.t1
QG2902_g14585.t1
QG2665_g4554.t1
QG2641_g15588.t1
NIC248_g12096.t1
EG6271_g5135.t1
ECA176_g9378.t1
ECA1174_g5675.t1
QG786_g5830.t1
QG1005_g12886.t1
NIC1052_g14955.t1
JU3237_g1438.t1
NIC2152_g13615.t1
0.000
0.003
0.006
0.009
0.012
HDR
No
Yes
Unknown
Species
C. briggsae
C. nigoni
JU1422_g15319.t1
ECA2857_g10936.t1
JU1419_g14672.t1
NIC2150_g13386.t1
VX153_g12486.t1
JU2617_g13826.t1
NIC2143_g10766.t1
QG2964_g17822.t1
YR106_g14154.t1
JU1418_g5399.t1
JU1420_g19863.t1
ECA2852_g8109.t1
ZF1220_g14589.t1
EG5268_g14440.t1
VX151_g10466.t1
JU2484_g14186.t1
QX1410_QX1410.5426.1
PE887_g7655.t1
NIC1529_g2416.t1
JU4058_g8022.t1
ECA1657_g3818.t1
JU3207_g8968.t1
NIC2152_g13615.t1
QX1796_g8391.t1
QG791_g8866.t1
QG2902_g14585.t1
QG2665_g4554.t1
QG2641_g15588.t1
NIC248_g12096.t1
EG6271_g5135.t1
ECA176_g9378.t1
ECA1174_g5675.t1
QG786_g5830.t1
QG1005_g12886.t1
JU3237_g1438.t1
NIC1052_g14955.t1
HDR
species
A.A.
%ID
100
90
80
70
60
50
HDR
Yes
No
NA
Species
C. briggsae
C. nigoni

OG0016494 - *Y54G11A.17* - II:4356559-4357276

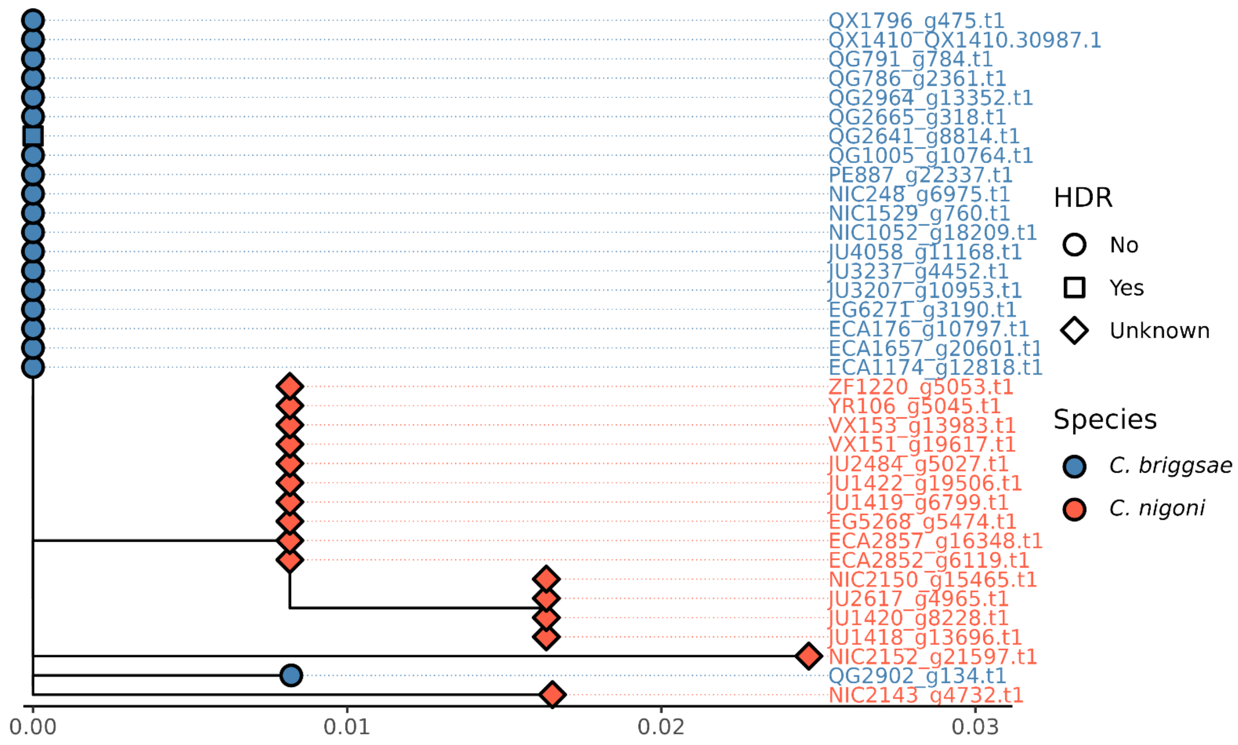


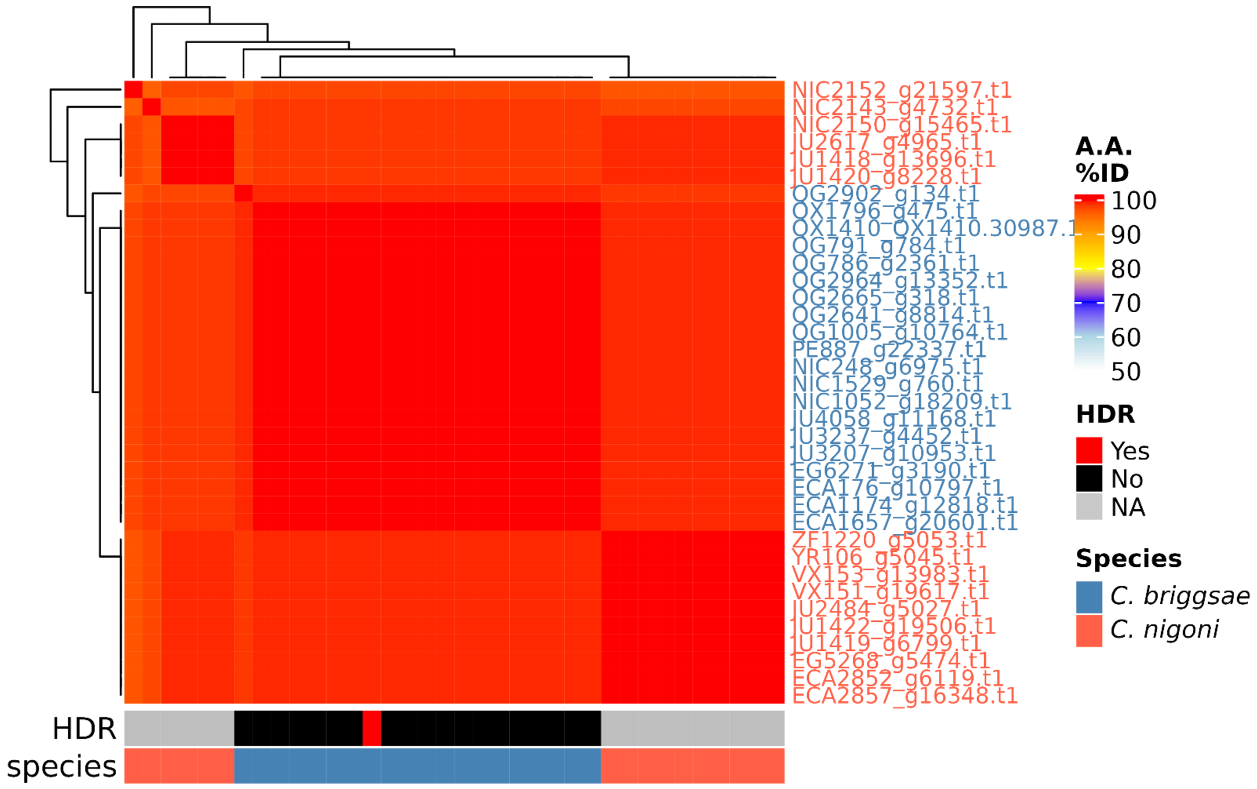

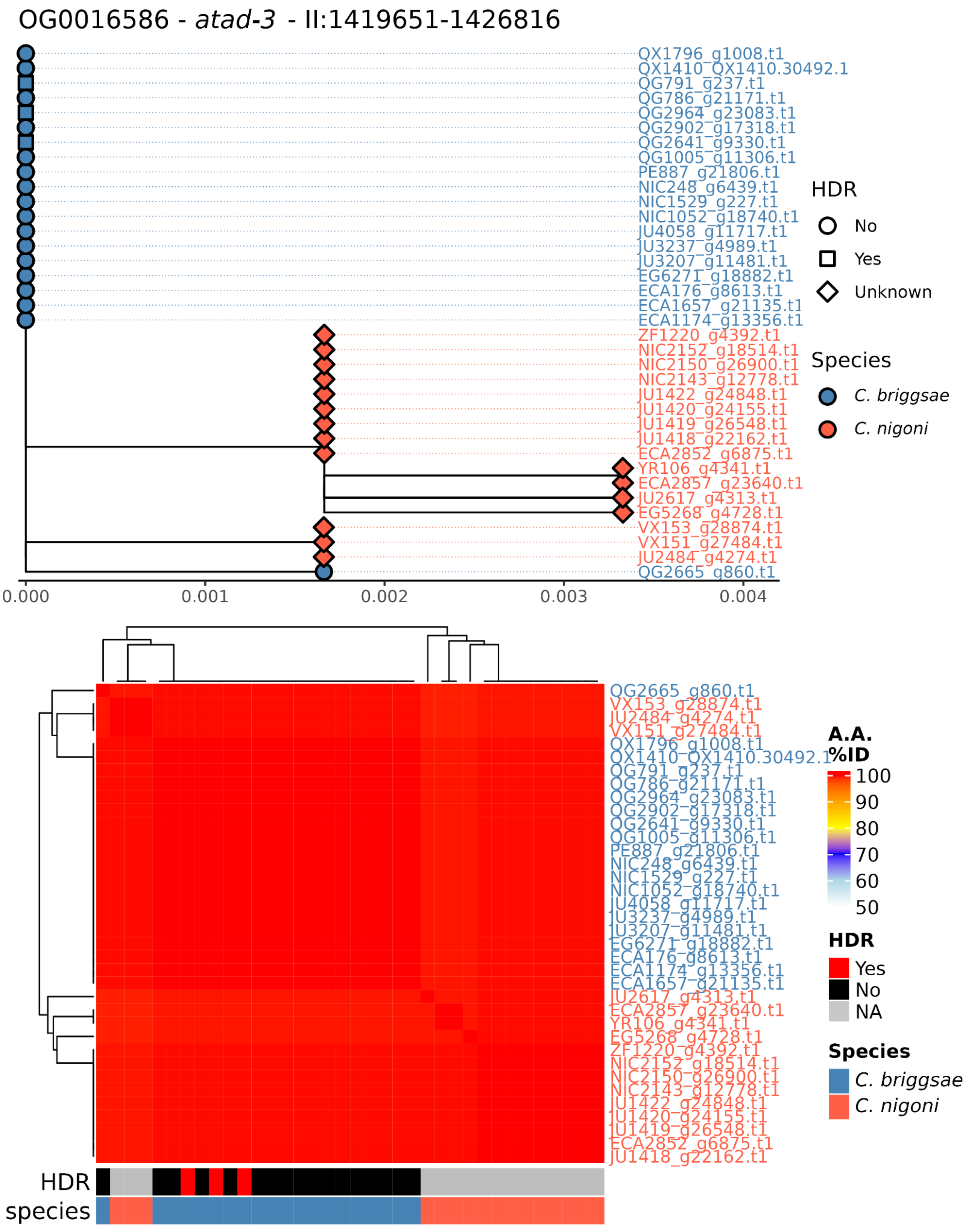
OG0016586 - atad-3 - II:1419651-1426816
QX1796_g1008.t1
QX1410_QX1410.30492.1
QG791_g237.t1
QG786_g21171.t1
QG2964_g23083.t1
QG2902_g17318.t1
QG2641_g9330.t1
QG1005_g11306.t1
PE887_g21806.t1
NIC248_g6439.t1
NIC1529_g227.t1
NIC1052_g18740.t1
JU4058_g11717.t1
JU3237_g4989.t1
JU3207_g11481.t1
EG6271_g18882.t1
ECA176_g8613.t1
ECA1657_g21135.t1
ECA1174_g13356.t1
ZF1220_g4392.t1
NIC2152_g18514.t1
NIC2150_g26900.t1
NIC2143_g12778.t1
JU1422_g24848.t1
JU1420_g24155.t1
JU1419_g26548.t1
JU1418_g22162.t1
ECA2852_g6875.t1
YR106_g4341.t1
ECA2857_g23640.t1
JU2617_g4313.t1
EG5268_g4728.t1
VX153_g28874.t1
VX151_g27484.t1
JU2484_g4274.t1
QG2665_g860.t1
0.000
0.001
0.002
0.003
0.004
HDR
No
Yes
Unknown
Species
C. briggsae
C. nigoni
A.A. %ID
100
90
80
70
60
50
HDR
Yes
No
NA
Species
C. briggsae
C. nigoni
HDR
species

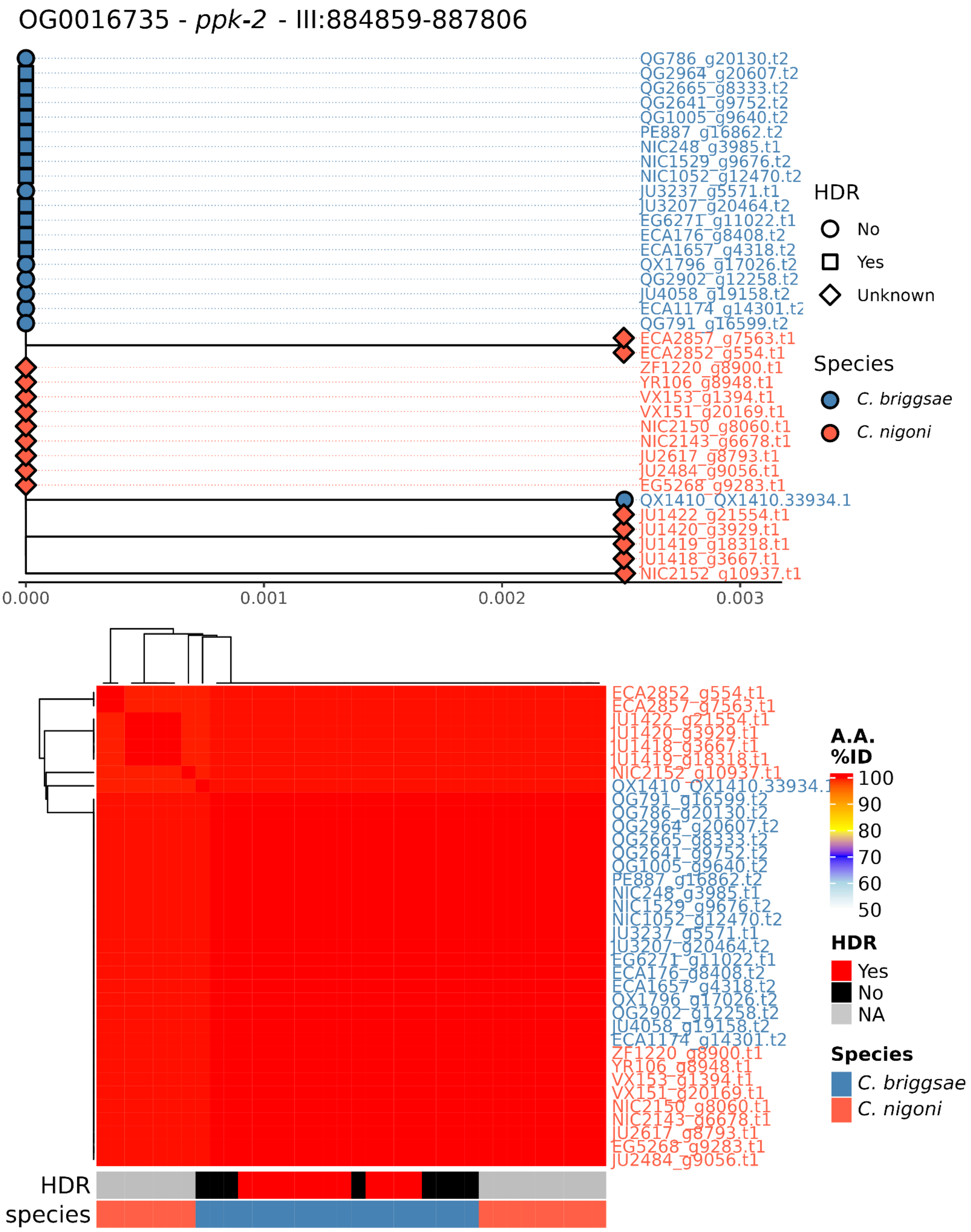
OG0016735 - ppk-2 - III:884859-887806
QG786_g20130.t2
QG2964_g20607.t2
QG2665_g8333.t2
QG2641_g9752.t2
QG1005_g9640.t2
PE887_g16862.t2
NIC248_g3985.t1
NIC1529_g9676.t2
NIC1052_g12470.t2
JU3237_g5571.t1
JU3207_g20464.t2
EG6271_g11022.t1
ECA176_g8408.t2
ECA1657_g4318.t2
QX1796_g17026.t2
QG2902_g12258.t2
JU4058_g19158.t2
ECA1174_g14301.t2
QG791_g16599.t2
ECA2857_g7563.t1
ECA2852_g554.t1
ZF1220_g8900.t1
YR106_g8948.t1
VX153_g1394.t1
VX151_g20169.t1
NIC2150_g8060.t1
NIC2143_g6678.t1
JU2617_g8793.t1
JU2484_g9056.t1
EG5268_g9283.t1
QX1410_QX1410.33934.1
JU1422_g21554.t1
JU1420_g3929.t1
JU1419_g18318.t1
JU1418_g3667.t1
NIC2152_g10937.t1
0.000
0.001
0.002
0.003
HDR
No
Yes
Unknown
Species
C. briggsae
C. nigoni
ECA2852_g554.t1
ECA2857_g7563.t1
JU1422_g21554.t1
JU1420_g3929.t1
JU1418_g3667.t1
JU1419_g18318.t1
NIC2152_g10937.t1
QX1410_QX1410.33934.1
QG791_g16599.t2
QG786_g20130.t2
QG2964_g20607.t2
QG2665_g8333.t2
QG2641_g9752.t2
QG1005_g9640.t2
PE887_g16862.t2
NIC248_g3985.t1
NIC1529_g9676.t2
NIC1052_g12470.t2
JU3237_g5571.t1
JU3207_g20464.t2
EG6271_g11022.t1
ECA176_g8408.t2
ECA1657_g4318.t2
QX1796_g17026.t2
QG2902_g12258.t2
JU4058_g19158.t2
ECA1174_g14301.t2
ZF1220_g8900.t1
YR106_g8948.t1
VX153_g1394.t1
VX151_g20169.t1
NIC2150_g8060.t1
NIC2143_g6678.t1
JU2617_g8793.t1
EG5268_g9283.t1
JU2484_g9056.t1
A.A. %ID
100
90
80
70
60
50
HDR
Yes
No
NA
Species
C. briggsae
C. nigoni
HDR
species

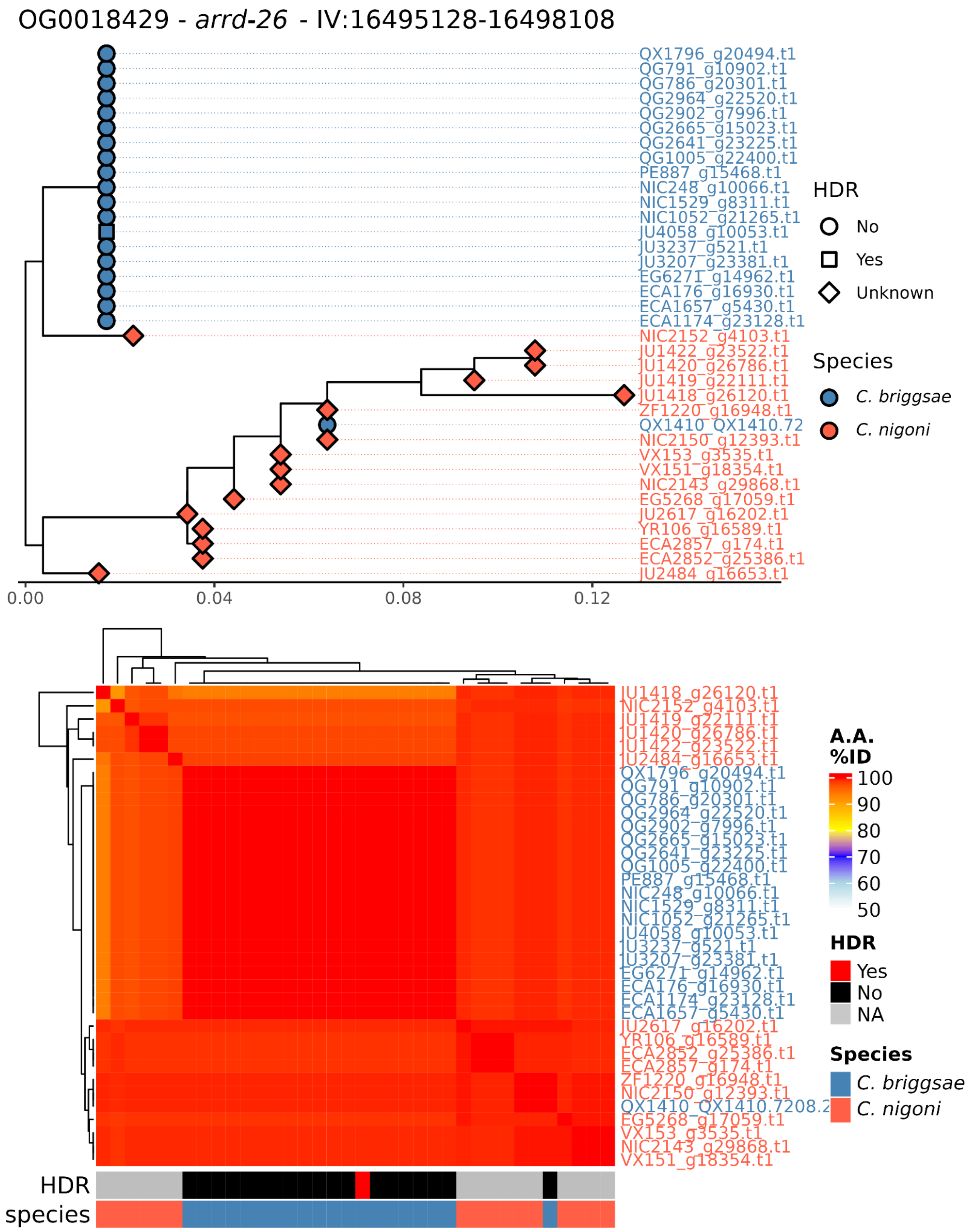
OG0018429 - *arrd-26* - IV:16495128-16498108
QX1796_g20494.t1
QG791_g10902.t1
QG786_g20301.t1
QG2964_g22520.t1
QG2902_g7996.t1
QG2665_g15023.t1
QG2641_g23225.t1
QG1005_g22400.t1
PE887_g15468.t1
NIC248_g10066.t1
NIC1529_g8311.t1
NIC1052_g21265.t1
JU4058_g10053.t1
JU3237_g521.t1
JU3207_g23381.t1
EG6271_g14962.t1
ECA176_g16930.t1
ECA1657_g5430.t1
ECA1174_g23128.t1
NIC2152_g4103.t1
JU1422_g23522.t1
JU1420_g26786.t1
JU1419_g22111.t1
JU1418_g26120.t1
ZF1220_g16948.t1
QX1410_QX1410.72
NIC2150_g12393.t1
VX153_g3535.t1
VX151_g18354.t1
NIC2143_g29868.t1
EG5268_g17059.t1
JU2617_g16202.t1
YR106_g16589.t1
ECA2857_g174.t1
ECA2852_g25386.t1
JU2484_g16653.t1
0.00
0.04
0.08
0.12
HDR
No
Yes
Unknown
Species
*C. briggsae*
*C. nigoni*
JU1418_g26120.t1
NIC2152_g4103.t1
JU1419_g22111.t1
JU1420_g26786.t1
JU1422_g23522.t1
JU2484_g16653.t1
QX1796_g20494.t1
QG791_g10902.t1
QG786_g20301.t1
QG2964_g22520.t1
QG2902_g7996.t1
QG2665_g15023.t1
QG2641_g23225.t1
QG1005_g22400.t1
PE887_g15468.t1
NIC248_g10066.t1
NIC1529_g8311.t1
NIC1052_g21265.t1
JU4058_g10053.t1
JU3237_g521.t1
JU3207_g23381.t1
EG6271_g14962.t1
ECA176_g16930.t1
ECA1174_g23128.t1
ECA1657_g5430.t1
JU2617_g16202.t1
YR106_g16589.t1
ECA2852_g25386.t1
ECA2857_g174.t1
ZF1220_g16948.t1
NIC2150_g12393.t1
QX1410_QX1410.7208.2
EG5268_g17059.t1
VX153_g3535.t1
NIC2143_g29868.t1
VX151_g18354.t1
A.A. %ID
100
90
80
70
60
50
HDR
Yes
No
NA
Species
*C. briggsae*
*C. nigoni*
HDR
species

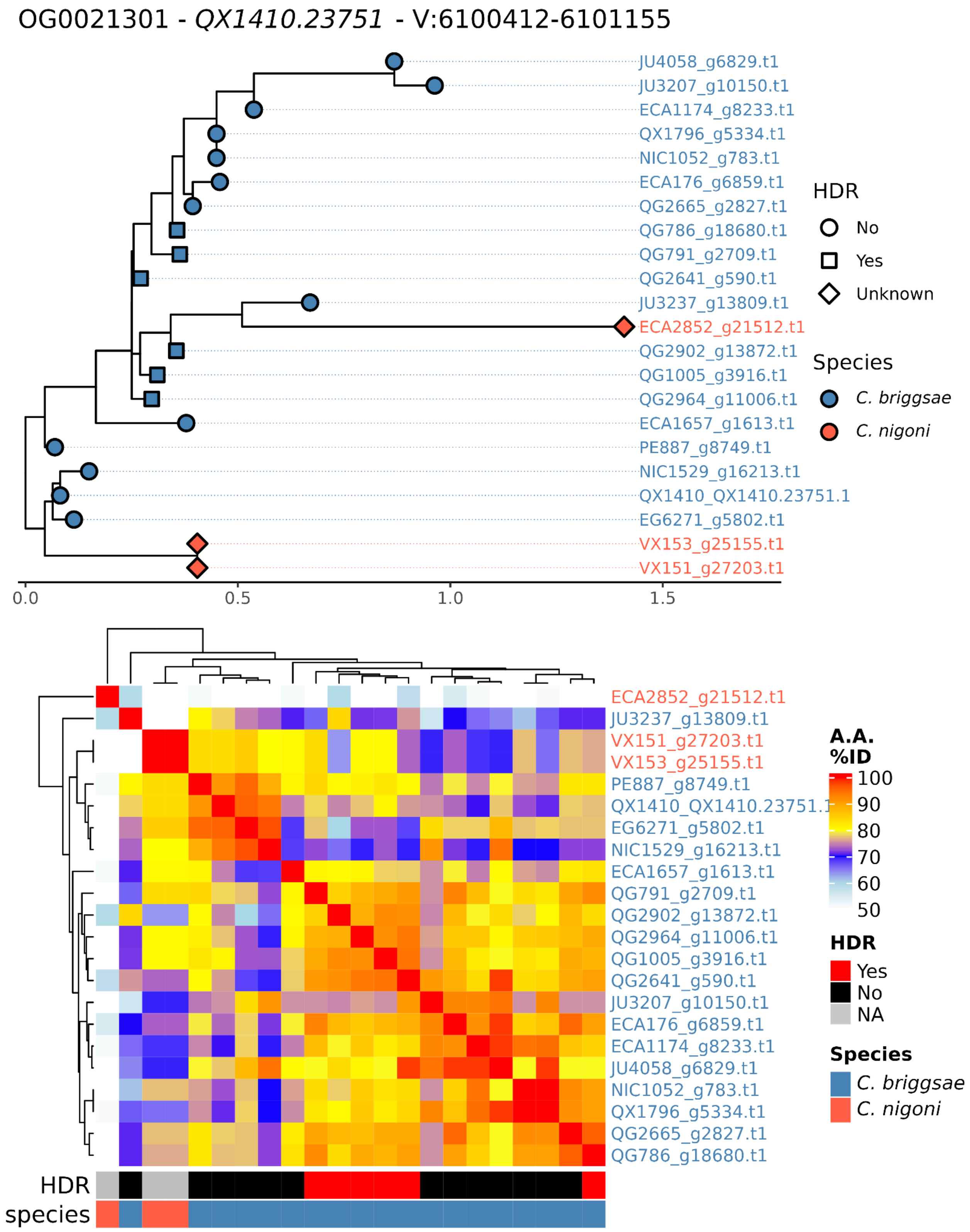
OG0021301 - QX1410.23751 - V:6100412-6101155
JU4058_g6829.t1
JU3207_g10150.t1
ECA1174_g8233.t1
QX1796_g5334.t1
NIC1052_g783.t1
ECA176_g6859.t1
QG2665_g2827.t1
QG786_g18680.t1
QG791_g2709.t1
QG2641_g590.t1
JU3237_g13809.t1
ECA2852_g21512.t1
QG2902_g13872.t1
QG1005_g3916.t1
QG2964_g11006.t1
ECA1657_g1613.t1
PE887_g8749.t1
NIC1529_g16213.t1
QX1410_QX1410.23751.1
EG6271_g5802.t1
VX153_g25155.t1
VX151_g27203.t1
0.0
0.5
1.0
1.5
HDR
No
Yes
Unknown
Species
C. briggsae
C. nigoni
ECA2852_g21512.t1
JU3237_g13809.t1
VX151_g27203.t1
VX153_g25155.t1
PE887_g8749.t1
QX1410_QX1410.23751.1
EG6271_g5802.t1
NIC1529_g16213.t1
ECA1657_g1613.t1
QG791_g2709.t1
QG2902_g13872.t1
QG2964_g11006.t1
QG1005_g3916.t1
QG2641_g590.t1
JU3207_g10150.t1
ECA176_g6859.t1
ECA1174_g8233.t1
JU4058_g6829.t1
NIC1052_g783.t1
QX1796_g5334.t1
QG2665_g2827.t1
QG786_g18680.t1
A.A. %ID
100
90
80
70
60
50
HDR
Yes
No
NA
Species
C. briggsae
C. nigoni
HDR
species

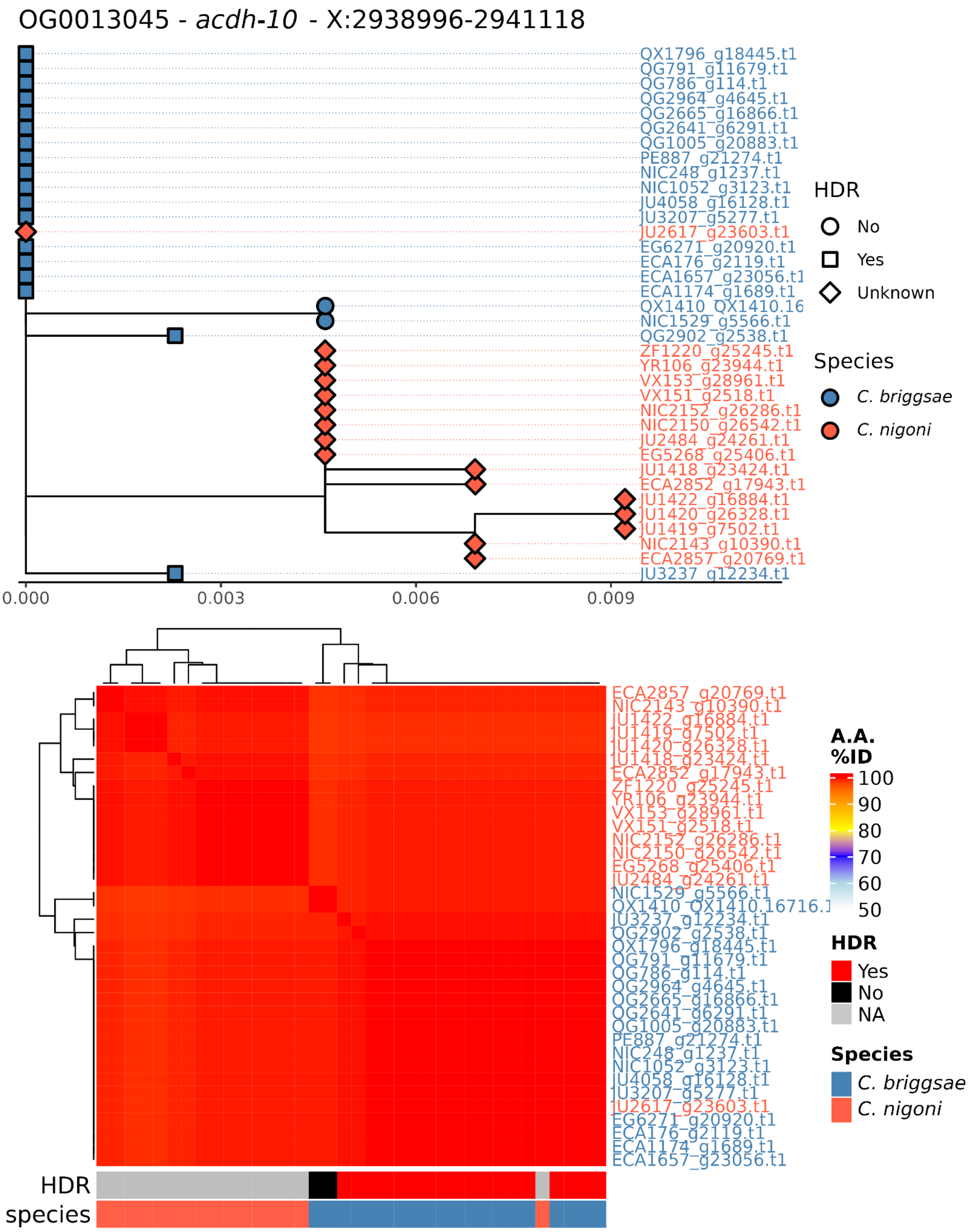
OG0013045 - acdh-10 - X:2938996-2941118
QX1796_g18445.t1
QG791_g11679.t1
QG786_g114.t1
QG2964_g4645.t1
QG2665_g16866.t1
QG2641_g6291.t1
QG1005_g20883.t1
PE887_g21274.t1
NIC248_g1237.t1
NIC1052_g3123.t1
JU4058_g16128.t1
JU3207_g5277.t1
JU2617_g23603.t1
EG6271_g20920.t1
ECA176_g2119.t1
ECA1657_g23056.t1
ECA1174_g1689.t1
QX1410_QX1410.16
NIC1529_g5566.t1
QG2902_g2538.t1
ZF1220_g25245.t1
YR106_g23944.t1
VX153_g28961.t1
VX151_g2518.t1
NIC2152_g26286.t1
NIC2150_g26542.t1
JU2484_g24261.t1
EG5268_g25406.t1
JU1418_g23424.t1
ECA2852_g17943.t1
JU1422_g16884.t1
JU1420_g26328.t1
JU1419_g7502.t1
NIC2143_g10390.t1
ECA2857_g20769.t1
JU3237_g12234.t1
0.000
0.003
0.006
0.009
HDR
No
Yes
Unknown
Species
C. briggsae
C. nigoni
ECA2857_g20769.t1
NIC2143_g10390.t1
JU1422_g16884.t1
JU1419_g7502.t1
JU1420_g26328.t1
JU1418_g23424.t1
ECA2852_g17943.t1
ZF1220_g25245.t1
YR106_g23944.t1
VX153_g28961.t1
VX151_g2518.t1
NIC2152_g26286.t1
NIC2150_g26542.t1
EG5268_g25406.t1
JU2484_g24261.t1
NIC1529_g5566.t1
QX1410_QX1410.16716.1
JU3237_g12234.t1
QG2902_g2538.t1
QX1796_g18445.t1
QG791_g11679.t1
QG786_g114.t1
QG2964_g4645.t1
QG2665_g16866.t1
QG2641_g6291.t1
QG1005_g20883.t1
PE887_g21274.t1
NIC248_g1237.t1
NIC1052_g3123.t1
JU4058_g16128.t1
JU3207_g5277.t1
JU2617_g23603.t1
EG6271_g20920.t1
ECA176_g2119.t1
ECA1174_g1689.t1
ECA1657_g23056.t1
A.A. %ID
100
90
80
70
60
50
HDR
Yes
No
NA
Species
C. briggsae
C. nigoni
HDR
species

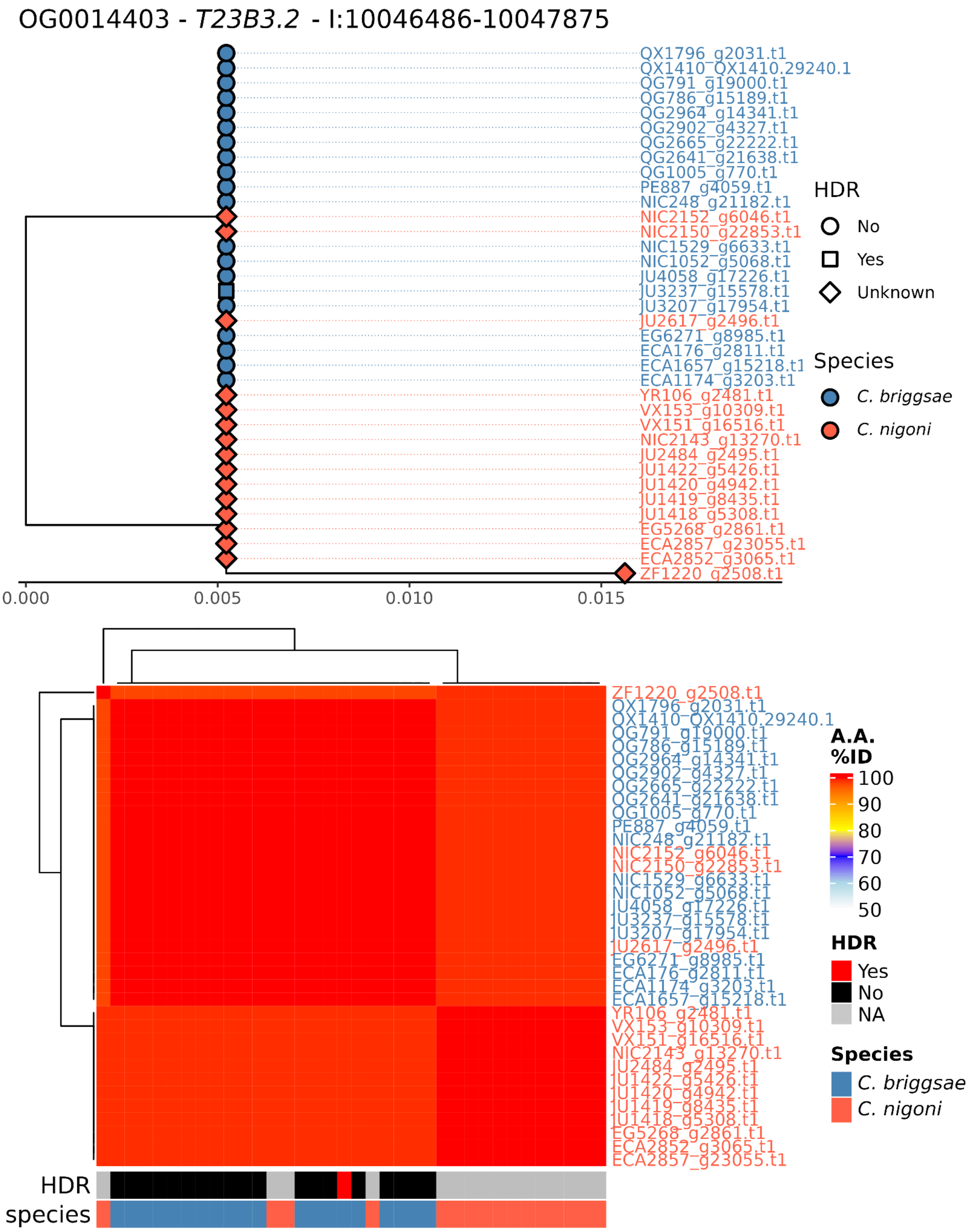
OG0014403 - T23B3.2 - I:10046486-10047875
QX1796_g2031.t1
QX1410_QX1410.29240.1
QG791_g19000.t1
QG786_g15189.t1
QG2964_g14341.t1
QG2902_g4327.t1
QG2665_g22222.t1
QG2641_g21638.t1
QG1005_g770.t1
PE887_g4059.t1
NIC248_g21182.t1
NIC2152_g6046.t1
NIC2150_g22853.t1
NIC1529_g6633.t1
NIC1052_g5068.t1
JU4058_g17226.t1
JU3237_g15578.t1
JU3207_g17954.t1
JU2617_g2496.t1
EG6271_g8985.t1
ECA176_g2811.t1
ECA1657_g15218.t1
ECA1174_g3203.t1
YR106_g2481.t1
VX153_g10309.t1
VX151_g16516.t1
NIC2143_g13270.t1
JU2484_g2495.t1
JU1422_g5426.t1
JU1420_g4942.t1
JU1419_g8435.t1
JU1418_g5308.t1
EG5268_g2861.t1
ECA2857_g23055.t1
ECA2852_g3065.t1
ZF1220_g2508.t1
0.000
0.005
0.010
0.015
HDR
No
Yes
Unknown
Species
C. briggsae
C. nigoni
A.A. %ID
100
90
80
70
60
50
HDR
Yes
No
NA
Species
C. briggsae
C. nigoni
HDR
species

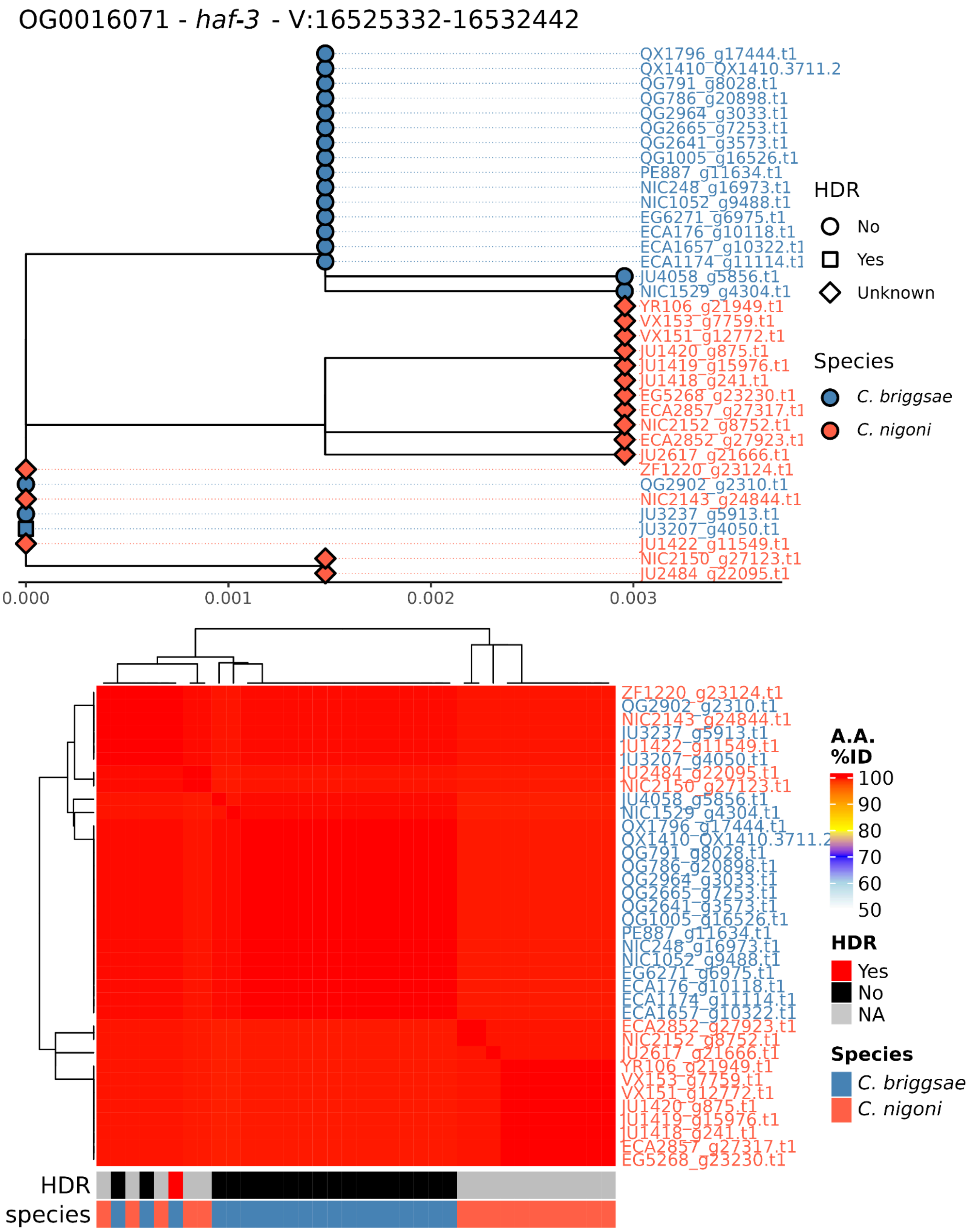
OG0016071 - *haf-3* - V:16525332-16532442
QX1796_g17444.t1
QX1410_QX1410.3711.2
QG791_g8028.t1
QG786_g20898.t1
QG2964_g3033.t1
QG2665_g7253.t1
QG2641_g3573.t1
QG1005_g16526.t1
PE887_g11634.t1
NIC248_g16973.t1
NIC1052_g9488.t1
EG6271_g6975.t1
ECA176_g10118.t1
ECA1657_g10322.t1
ECA1174_g11114.t1
JU4058_g5856.t1
NIC1529_g4304.t1
YR106_g21949.t1
VX153_g7759.t1
VX151_g12772.t1
JU1420_g875.t1
JU1419_g15976.t1
JU1418_g241.t1
EG5268_g23230.t1
ECA2857_g27317.t1
NIC2152_g8752.t1
ECA2852_g27923.t1
JU2617_g21666.t1
ZF1220_g23124.t1
QG2902_g2310.t1
NIC2143_g24844.t1
JU3237_g5913.t1
JU3207_g4050.t1
JU1422_g11549.t1
NIC2150_g27123.t1
JU2484_g22095.t1
0.000
0.001
0.002
0.003
HDR
No
Yes
Unknown
Species
*C. briggsae*
*C. nigoni*
A.A. %ID
100
90
80
70
60
50
HDR
Yes
No
NA
Species
*C. briggsae*
*C. nigoni*
HDR
species

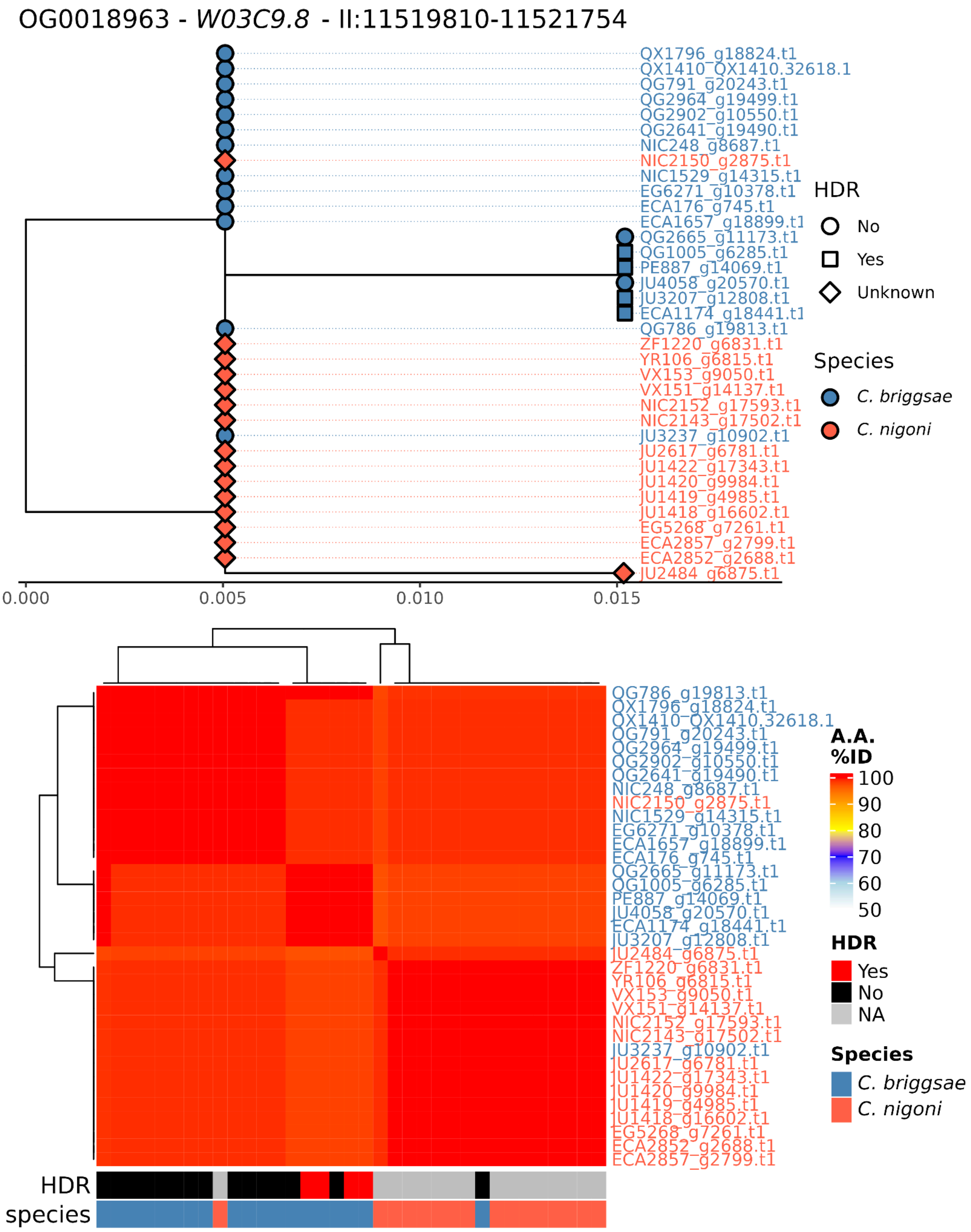

OG0018963 - W03C9.8 - II:11519810-11521754
QX1796_g18824.t1
QX1410_QX1410.32618.1
QG791_g20243.t1
QG2964_g19499.t1
QG2902_g10550.t1
QG2641_g19490.t1
NIC248_g8687.t1
NIC2150_g2875.t1
NIC1529_g14315.t1
EG6271_g10378.t1
ECA176_g745.t1
ECA1657_g18899.t1
QG2665_g11173.t1
QG1005_g6285.t1
PE887_g14069.t1
JU4058_g20570.t1
JU3207_g12808.t1
ECA1174_g18441.t1
QG786_g19813.t1
ZF1220_g6831.t1
YR106_g6815.t1
VX153_g9050.t1
VX151_g14137.t1
NIC2152_g17593.t1
NIC2143_g17502.t1
JU3237_g10902.t1
JU2617_g6781.t1
JU1422_g17343.t1
JU1420_g9984.t1
JU1419_g4985.t1
JU1418_g16602.t1
EG5268_g7261.t1
ECA2857_g2799.t1
ECA2852_g2688.t1
JU2484_g6875.t1
0.000
0.005
0.010
0.015
HDR
No
Yes
Unknown
Species
C. briggsae
C. nigoni
A.A. %ID
100
90
80
70
60
50
HDR
Yes
No
NA
Species
C. briggsae
C. nigoni
HDR
species

OG0019613 - *C08E8.10* - V:18098522-18099604

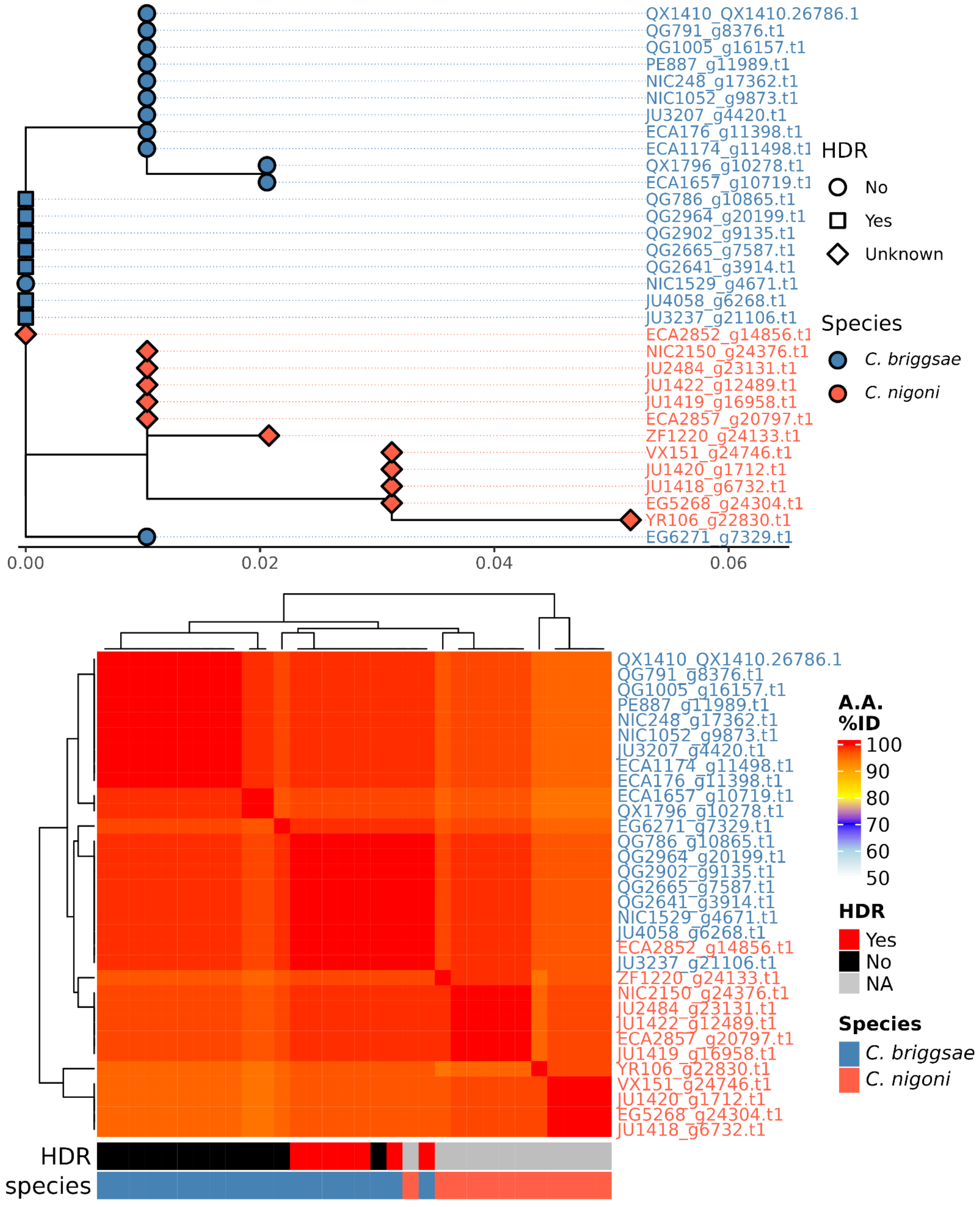

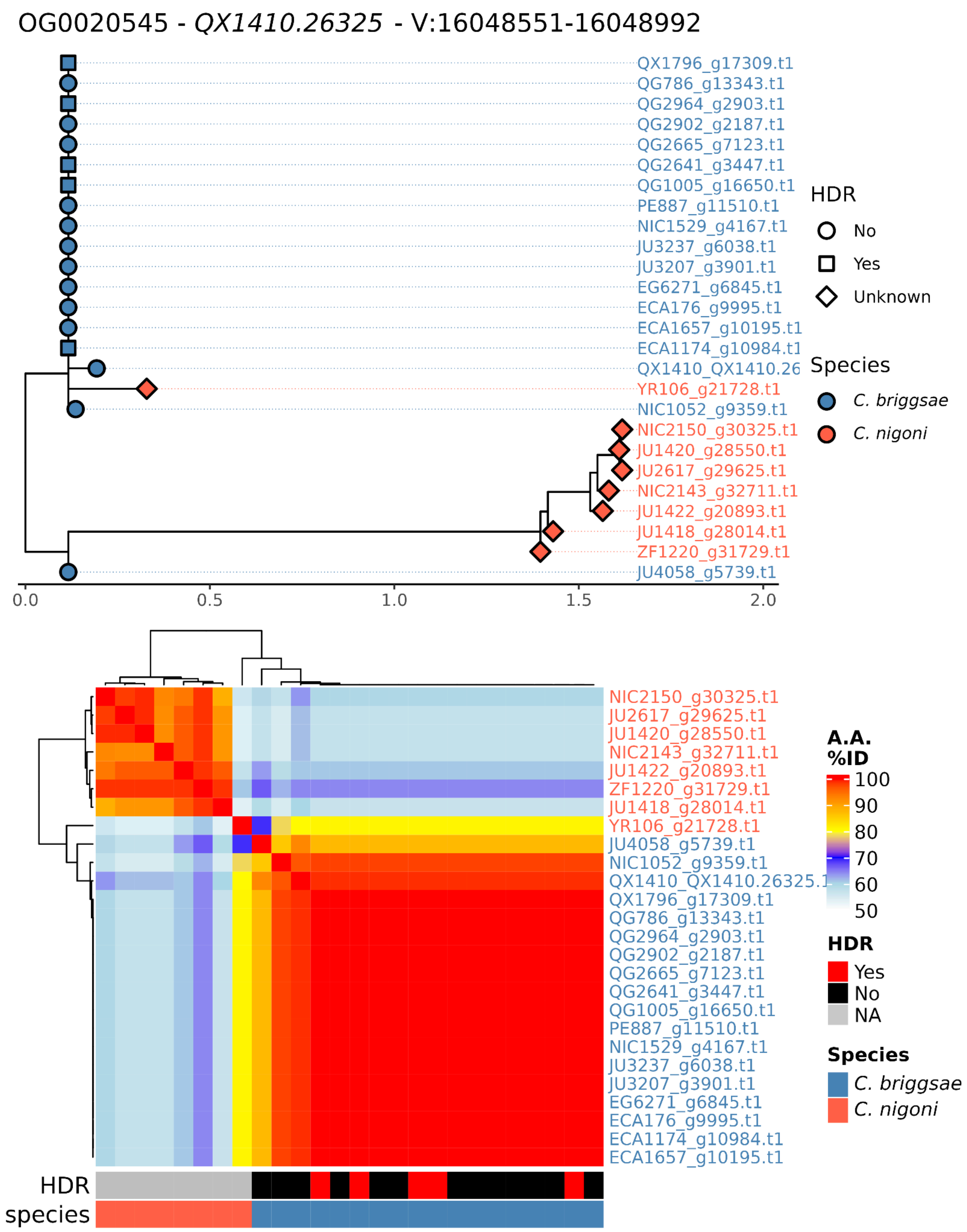
OG0020545 - QX1410.26325 - V:16048551-16048992
QX1796_g17309.t1
QG786_g13343.t1
QG2964_g2903.t1
QG2902_g2187.t1
QG2665_g7123.t1
QG2641_g3447.t1
QG1005_g16650.t1
PE887_g11510.t1
NIC1529_g4167.t1
JU3237_g6038.t1
JU3207_g3901.t1
EG6271_g6845.t1
ECA176_g9995.t1
ECA1657_g10195.t1
ECA1174_g10984.t1
QX1410_QX1410.26
YR106_g21728.t1
NIC1052_g9359.t1
NIC2150_g30325.t1
JU1420_g28550.t1
JU2617_g29625.t1
NIC2143_g32711.t1
JU1422_g20893.t1
JU1418_g28014.t1
ZF1220_g31729.t1
JU4058_g5739.t1
0.0
0.5
1.0
1.5
2.0
HDR
No
Yes
Unknown
Species
C. briggsae
C. nigoni
NIC2150_g30325.t1
JU2617_g29625.t1
JU1420_g28550.t1
NIC2143_g32711.t1
JU1422_g20893.t1
ZF1220_g31729.t1
JU1418_g28014.t1
YR106_g21728.t1
JU4058_g5739.t1
NIC1052_g9359.t1
QX1410_QX1410.26325.1
QX1796_g17309.t1
QG786_g13343.t1
QG2964_g2903.t1
QG2902_g2187.t1
QG2665_g7123.t1
QG2641_g3447.t1
QG1005_g16650.t1
PE887_g11510.t1
NIC1529_g4167.t1
JU3237_g6038.t1
JU3207_g3901.t1
EG6271_g6845.t1
ECA176_g9995.t1
ECA1174_g10984.t1
ECA1657_g10195.t1
HDR
species
A.A. %ID
100
90
80
70
60
50
HDR
Yes
No
NA
Species
C. briggsae
C. nigoni

OG0021407 - *QX1410.37633* - X:4211196-4221211

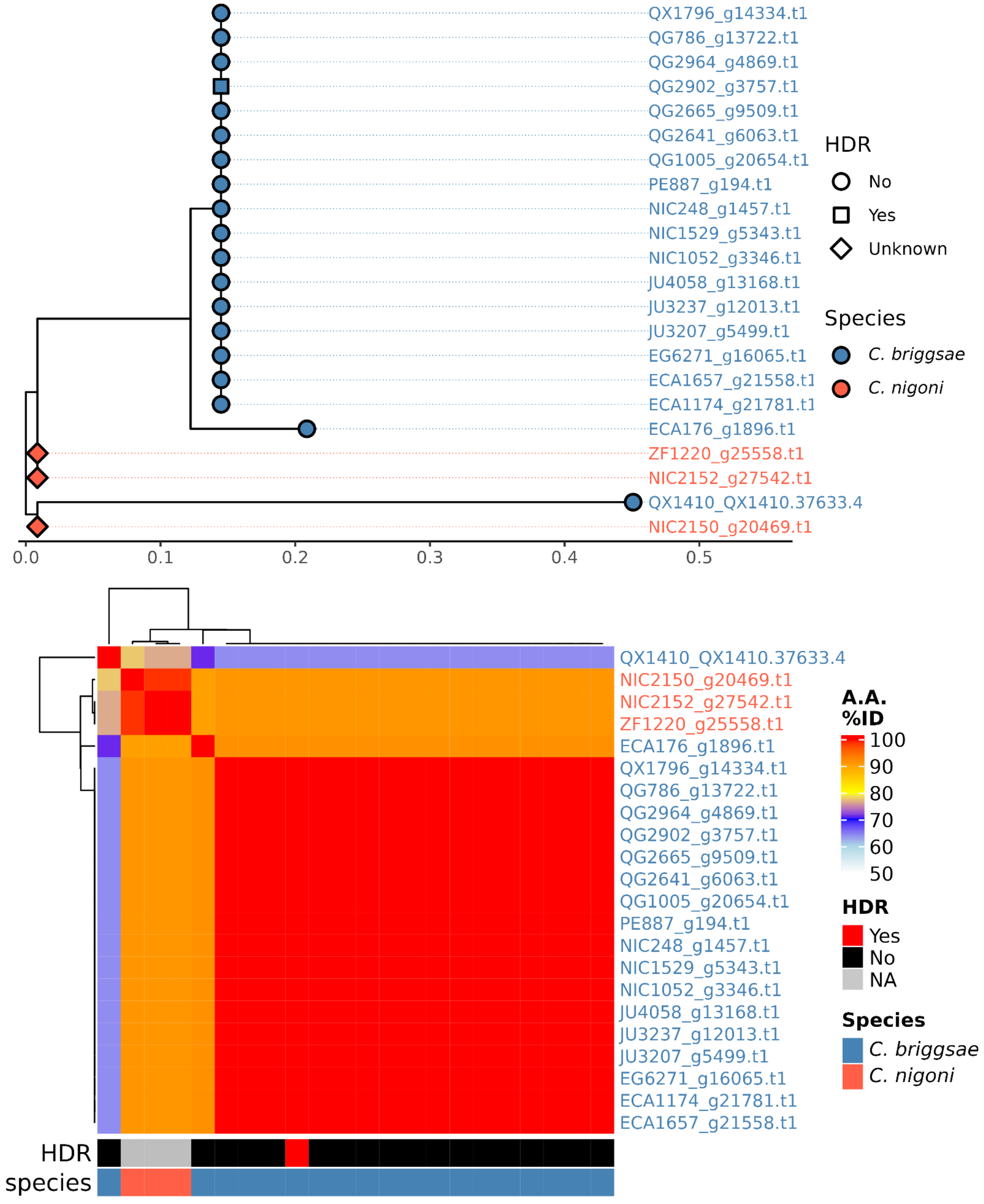

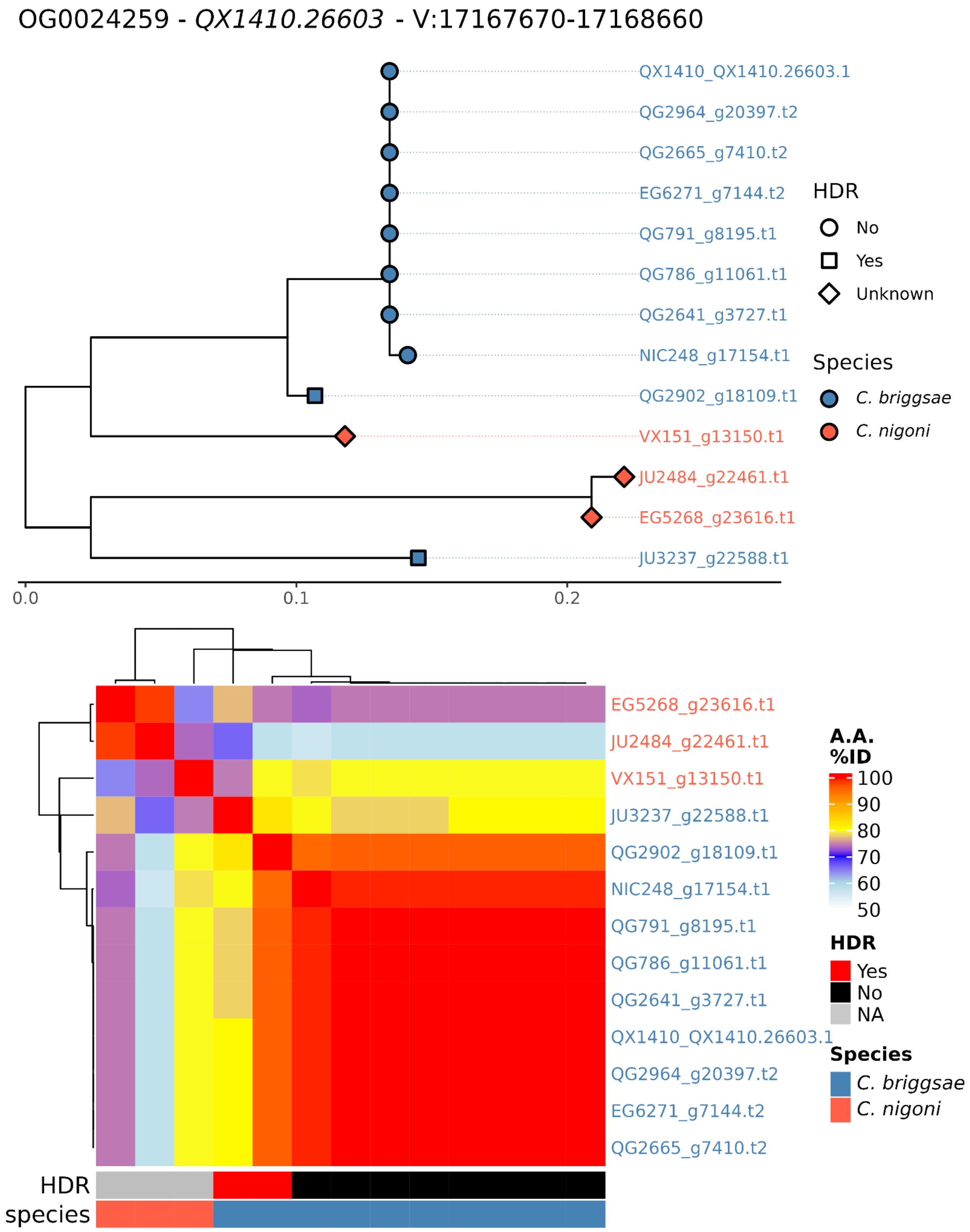

OG0024259 - QX1410.26603 - V:17167670-17168660
QX1410_QX1410.26603.1
QG2964_g20397.t2
QG2665_g7410.t2
EG6271_g7144.t2
QG791_g8195.t1
QG786_g11061.t1
QG2641_g3727.t1
NIC248_g17154.t1
QG2902_g18109.t1
VX151_g13150.t1
JU2484_g22461.t1
EG5268_g23616.t1
JU3237_g22588.t1
0.0
0.1
0.2
HDR
No
Yes
Unknown
Species
C. briggsae
C. nigoni
EG5268_g23616.t1
JU2484_g22461.t1
VX151_g13150.t1
JU3237_g22588.t1
QG2902_g18109.t1
NIC248_g17154.t1
QG791_g8195.t1
QG786_g11061.t1
QG2641_g3727.t1
QX1410_QX1410.26603.1
QG2964_g20397.t2
EG6271_g7144.t2
QG2665_g7410.t2
HDR
species
A.A. %ID
100
90
80
70
60
50
HDR
Yes
No
NA
Species
C. briggsae
C. nigoni

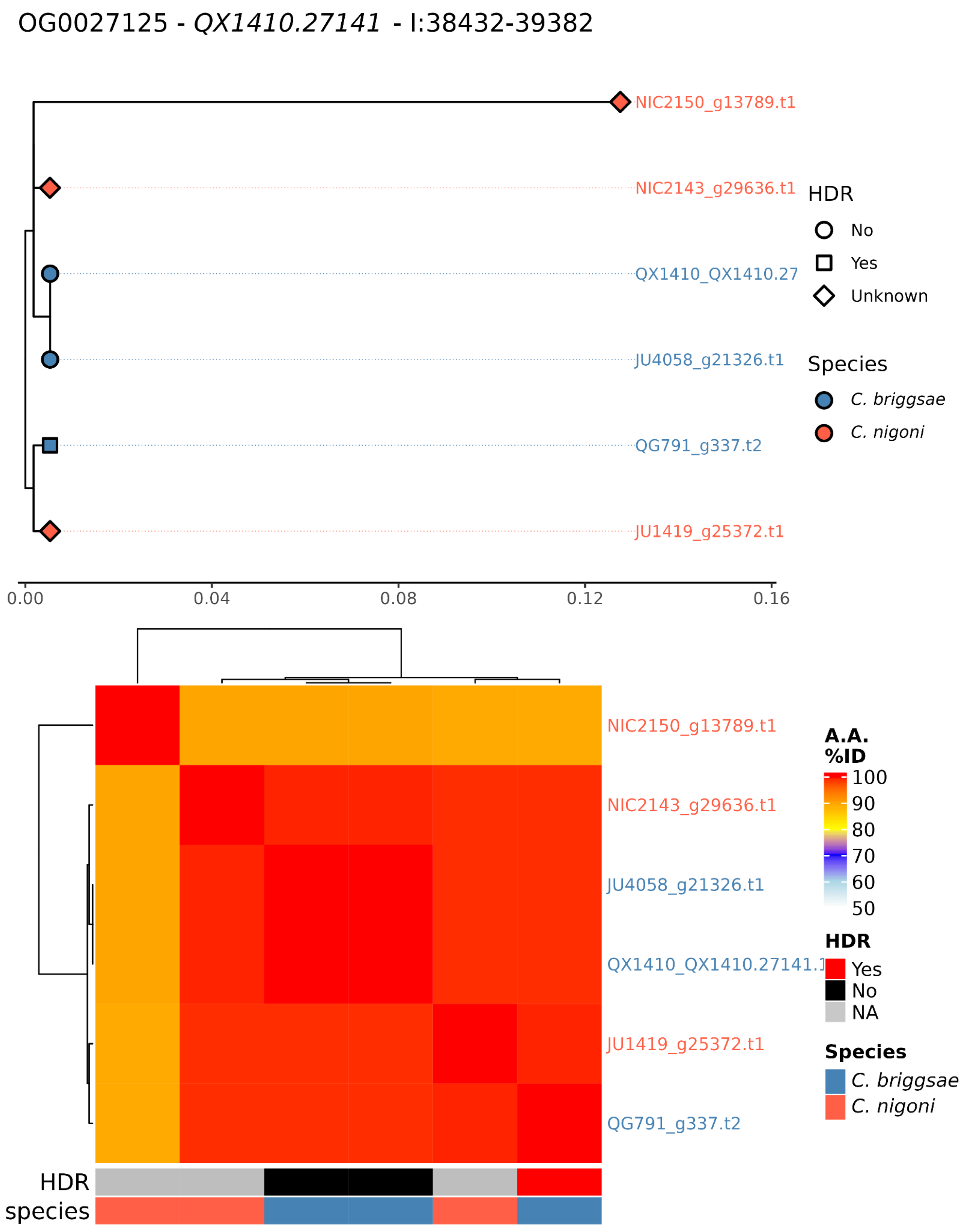
OG0027125 - QX1410.27141 - I:38432-39382
NIC2150_g13789.t1
NIC2143_g29636.t1
QX1410_QX1410.27
JU4058_g21326.t1
QG791_g337.t2
JU1419_g25372.t1
HDR
No
Yes
Unknown
Species
C. briggsae
C. nigoni
0.00
0.04
0.08
0.12
0.16
NIC2150_g13789.t1
NIC2143_g29636.t1
JU4058_g21326.t1
QX1410_QX1410.27141.1
JU1419_g25372.t1
QG791_g337.t2
A.A. %ID
100
90
80
70
60
50
HDR
Yes
No
NA
Species
C. briggsae
C. nigoni
HDR
species

OG0013391 - *arv-1* - II:835343-837443

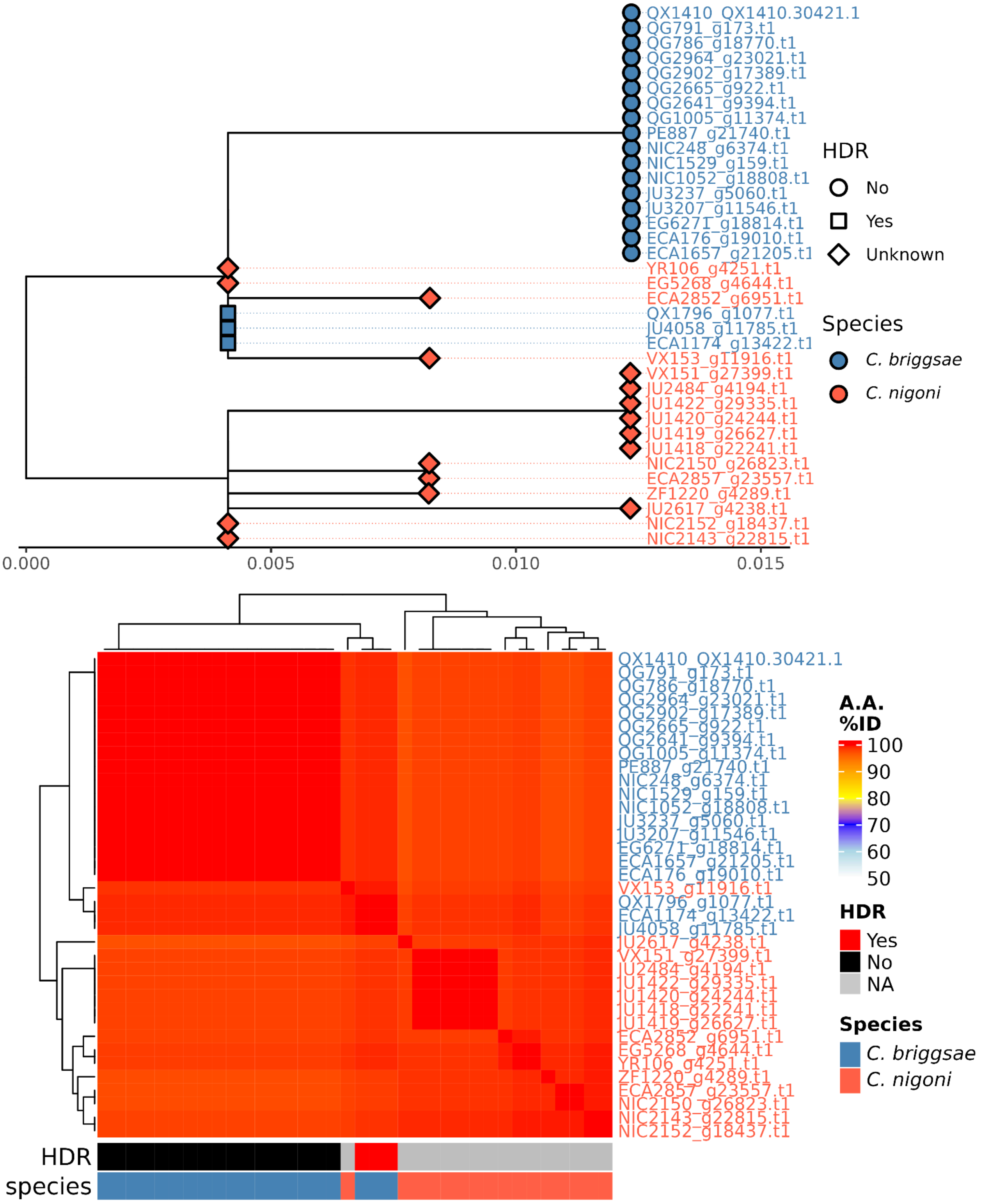

OG0014471 - *Y94H6A.12* - IV:15614265-15614759

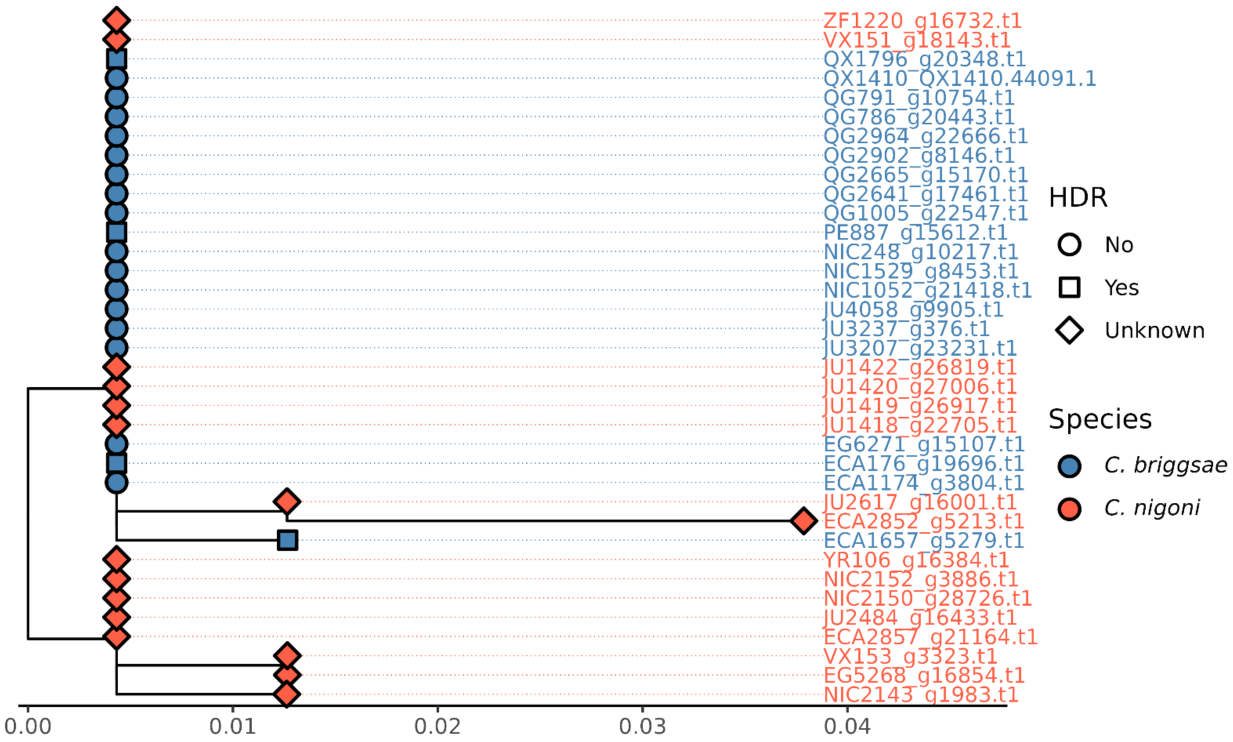


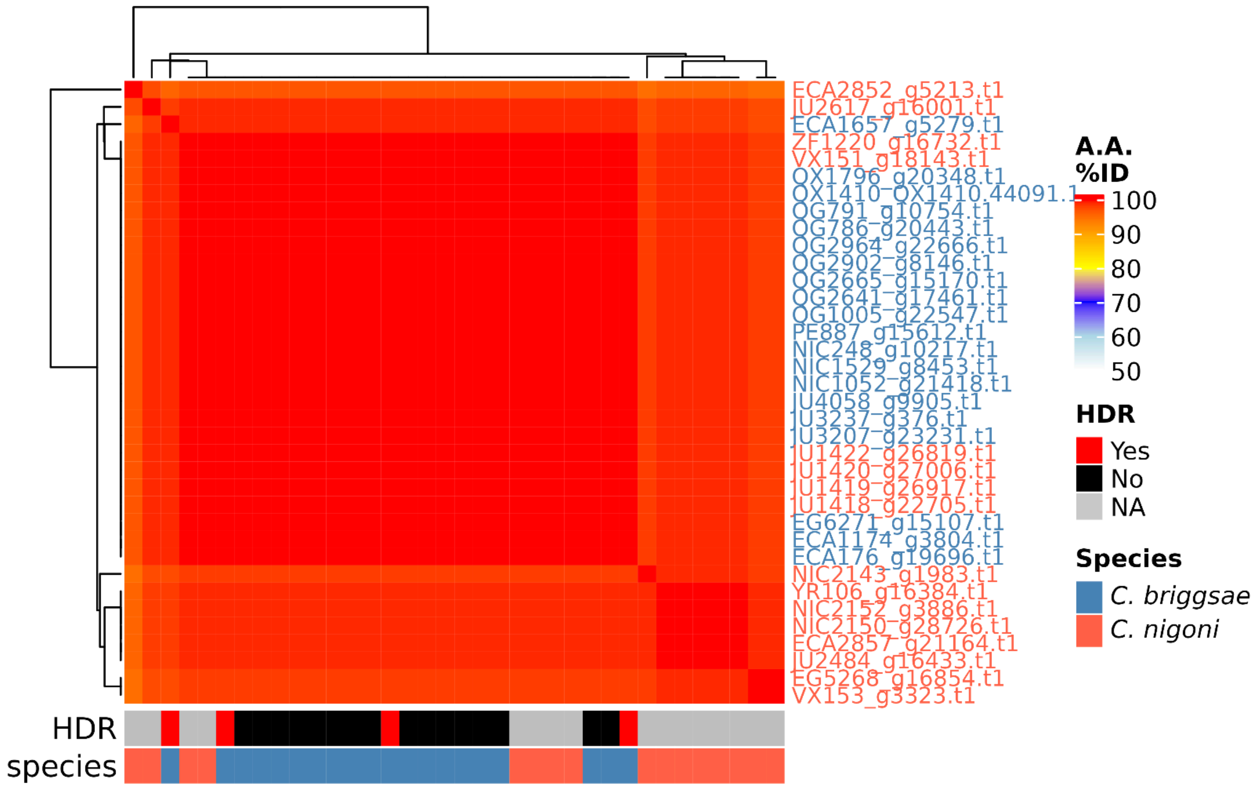

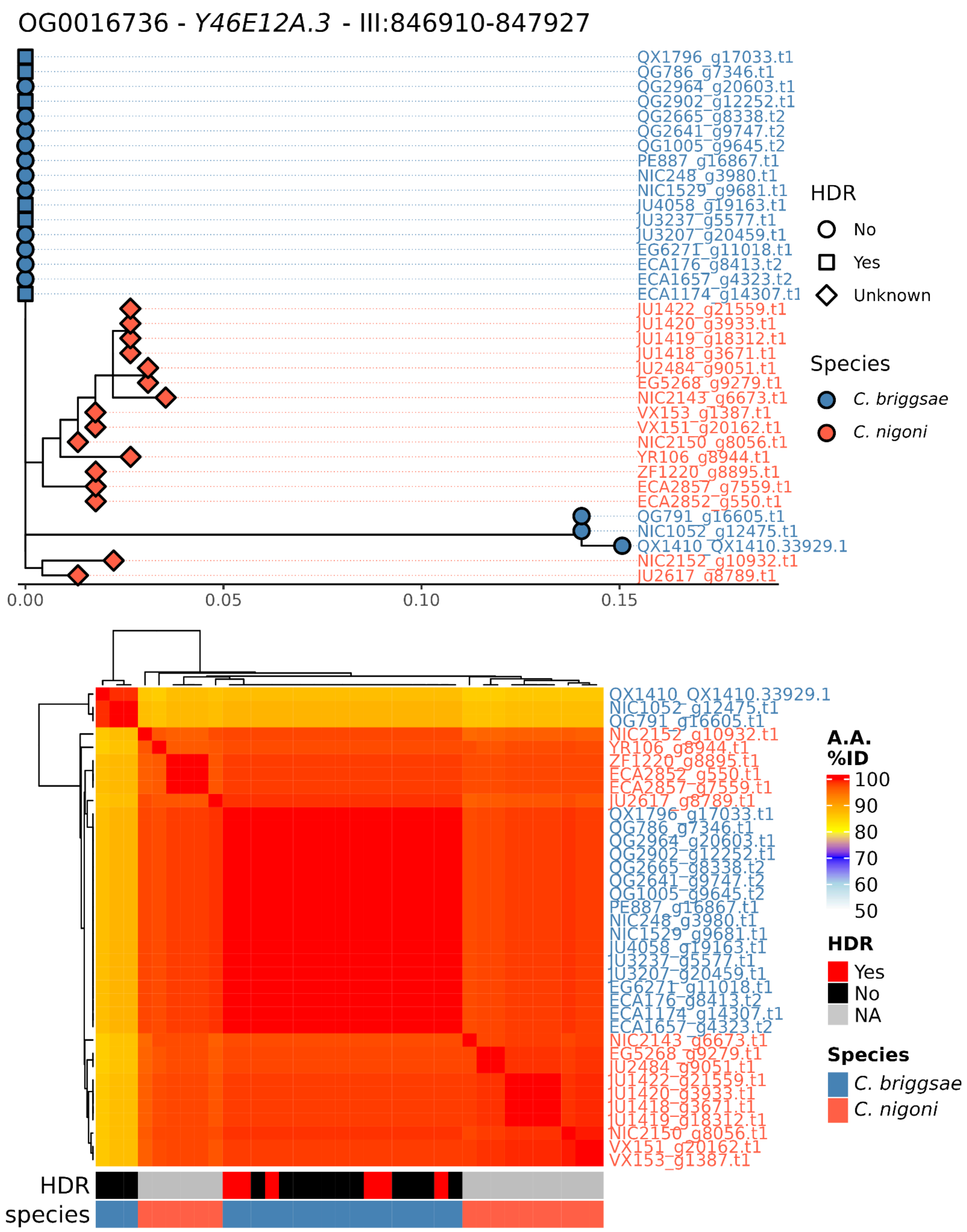
OG0016736 - Y46E12A.3 - III:846910-847927
QX1796_g17033.t1
QG786_g7346.t1
QG2964_g20603.t1
QG2902_g12252.t1
QG2665_g8338.t2
QG2641_g9747.t2
QG1005_g9645.t2
PE887_g16867.t1
NIC248_g3980.t1
NIC1529_g9681.t1
JU4058_g19163.t1
JU3237_g5577.t1
JU3207_g20459.t1
EG6271_g11018.t1
ECA176_g8413.t2
ECA1657_g4323.t2
ECA1174_g14307.t1
JU1422_g21559.t1
JU1420_g3933.t1
JU1419_g18312.t1
JU1418_g3671.t1
JU2484_g9051.t1
EG5268_g9279.t1
NIC2143_g6673.t1
VX153_g1387.t1
VX151_g20162.t1
NIC2150_g8056.t1
YR106_g8944.t1
ZF1220_g8895.t1
ECA2857_g7559.t1
ECA2852_g550.t1
QG791_g16605.t1
NIC1052_g12475.t1
QX1410_QX1410.33929.1
NIC2152_g10932.t1
JU2617_g8789.t1
0.00
0.05
0.10
0.15
HDR
No
Yes
Unknown
Species
C. briggsae
C. nigoni
QX1410_QX1410.33929.1
NIC1052_g12475.t1
QG791_g16605.t1
NIC2152_g10932.t1
YR106_g8944.t1
ZF1220_g8895.t1
ECA2852_g550.t1
ECA2857_g7559.t1
JU2617_g8789.t1
QX1796_g17033.t1
QG786_g7346.t1
QG2964_g20603.t1
QG2902_g12252.t1
QG2665_g8338.t2
QG2641_g9747.t2
QG1005_g9645.t2
PE887_g16867.t1
NIC248_g3980.t1
NIC1529_g9681.t1
JU4058_g19163.t1
JU3237_g5577.t1
JU3207_g20459.t1
EG6271_g11018.t1
ECA176_g8413.t2
ECA1174_g14307.t1
ECA1657_g4323.t2
NIC2143_g6673.t1
EG5268_g9279.t1
JU2484_g9051.t1
JU1422_g21559.t1
JU1420_g3933.t1
JU1418_g3671.t1
JU1419_g18312.t1
NIC2150_g8056.t1
VX151_g20162.t1
VX153_g1387.t1
A.A. %ID
100
90
80
70
60
50
HDR
Yes
No
NA
Species
C. briggsae
C. nigoni
HDR
species

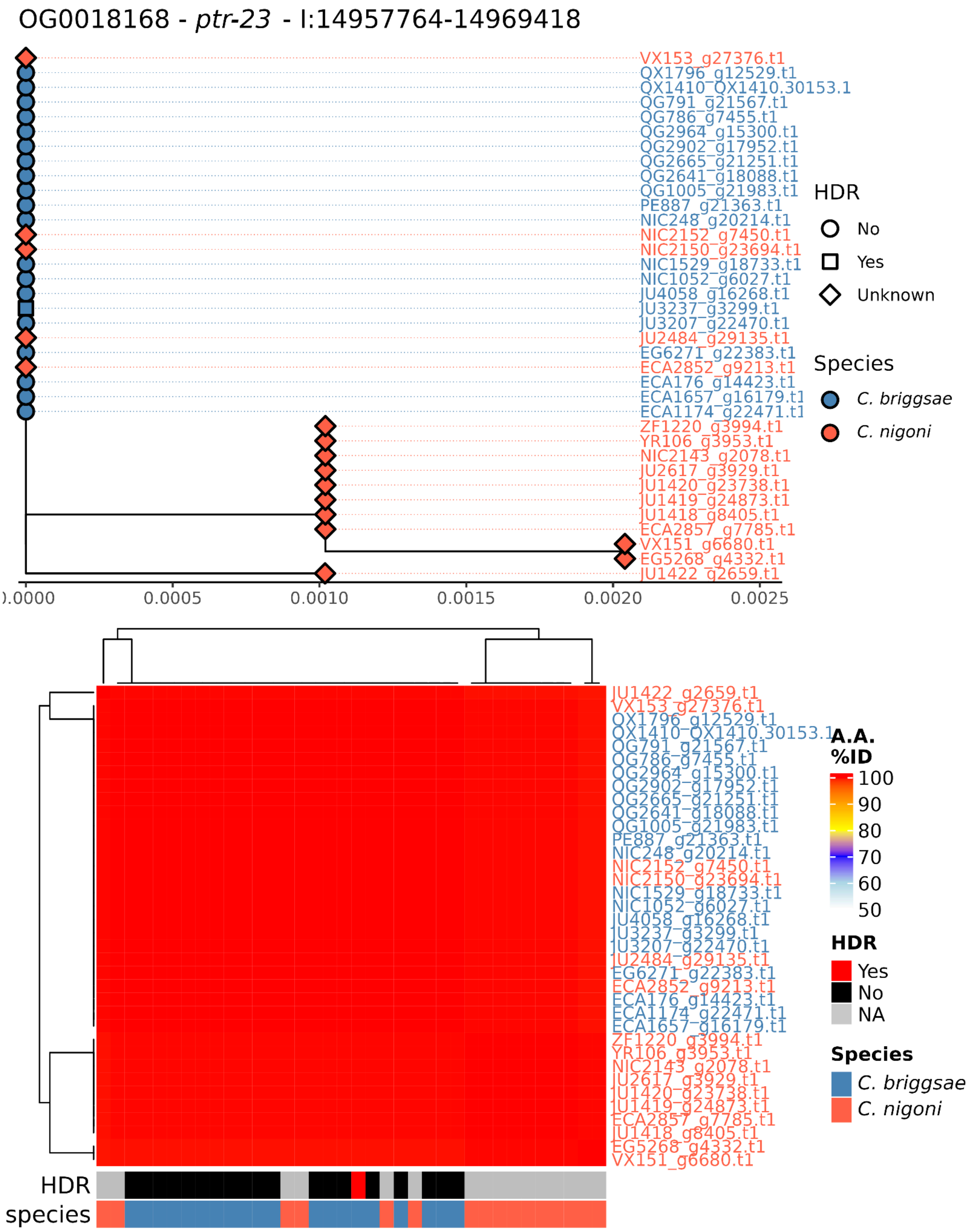
OG0018168 - ptr-23 - I:14957764-14969418
VX153_g27376.t1
QX1796_g12529.t1
QX1410_QX1410.30153.1
QG791_g21567.t1
QG786_g7455.t1
QG2964_g15300.t1
QG2902_g17952.t1
QG2665_g21251.t1
QG2641_g18088.t1
QG1005_g21983.t1
PE887_g21363.t1
NIC248_g20214.t1
NIC2152_g7450.t1
NIC2150_g23694.t1
NIC1529_g18733.t1
NIC1052_g6027.t1
JU4058_g16268.t1
JU3237_g3299.t1
JU3207_g22470.t1
JU2484_g29135.t1
EG6271_g22383.t1
ECA2852_g9213.t1
ECA176_g14423.t1
ECA1657_g16179.t1
ECA1174_g22471.t1
ZF1220_g3994.t1
YR106_g3953.t1
NIC2143_g2078.t1
JU2617_g3929.t1
JU1420_g23738.t1
JU1419_g24873.t1
JU1418_g8405.t1
ECA2857_g7785.t1
VX151_g6680.t1
EG5268_g4332.t1
JU1422_g2659.t1
0.0000
0.0005
0.0010
0.0015
0.0020
0.0025
HDR
No
Yes
Unknown
Species
C. briggsae
C. nigoni
JU1422_g2659.t1
VX153_g27376.t1
QX1796_g12529.t1
QX1410_QX1410.30153.1
QG791_g21567.t1
QG786_g7455.t1
QG2964_g15300.t1
QG2902_g17952.t1
QG2665_g21251.t1
QG2641_g18088.t1
QG1005_g21983.t1
PE887_g21363.t1
NIC248_g20214.t1
NIC2152_g7450.t1
NIC2150_g23694.t1
NIC1529_g18733.t1
NIC1052_g6027.t1
JU4058_g16268.t1
JU3237_g3299.t1
JU3207_g22470.t1
JU2484_g29135.t1
EG6271_g22383.t1
ECA2852_g9213.t1
ECA176_g14423.t1
ECA1174_g22471.t1
ECA1657_g16179.t1
ZF1220_g3994.t1
YR106_g3953.t1
NIC2143_g2078.t1
JU2617_g3929.t1
JU1420_g23738.t1
JU1419_g24873.t1
ECA2857_g7785.t1
JU1418_g8405.t1
EG5268_g4332.t1
VX151_g6680.t1
A.A.
%ID
100
90
80
70
60
50
HDR
Yes
No
NA
Species
C. briggsae
C. nigoni
HDR
species

OG0024289 - *QX1410.26842* - V:18415681-18416930

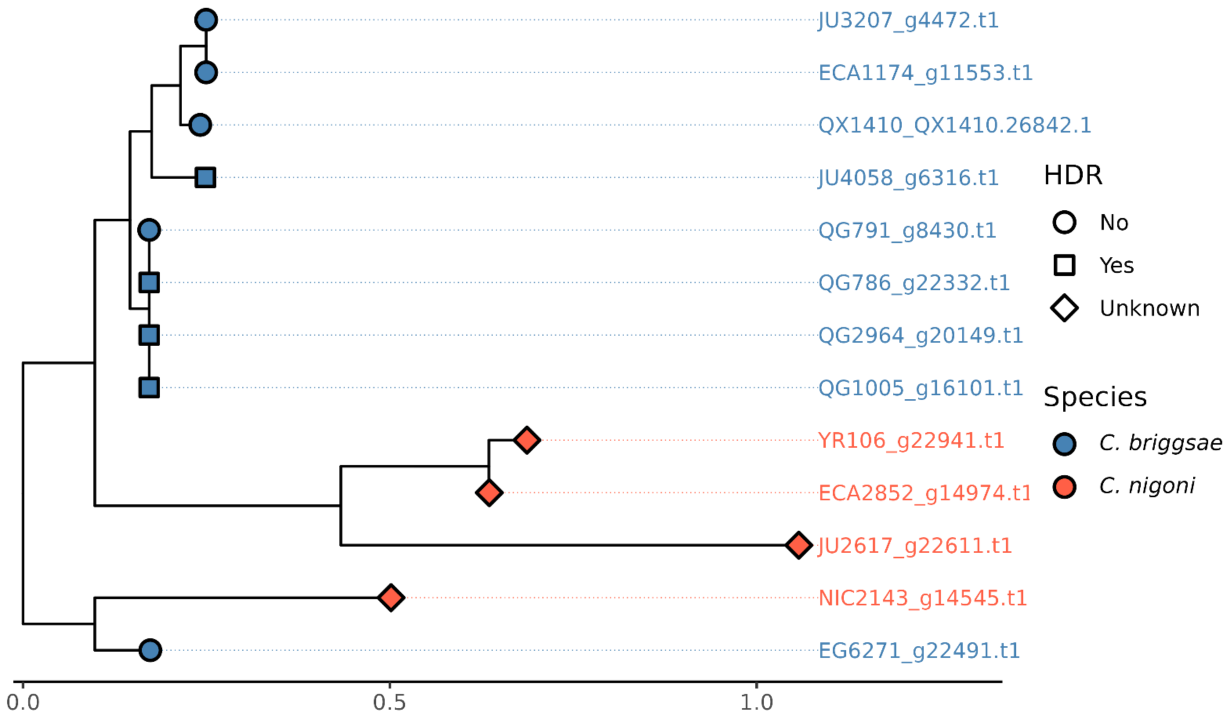


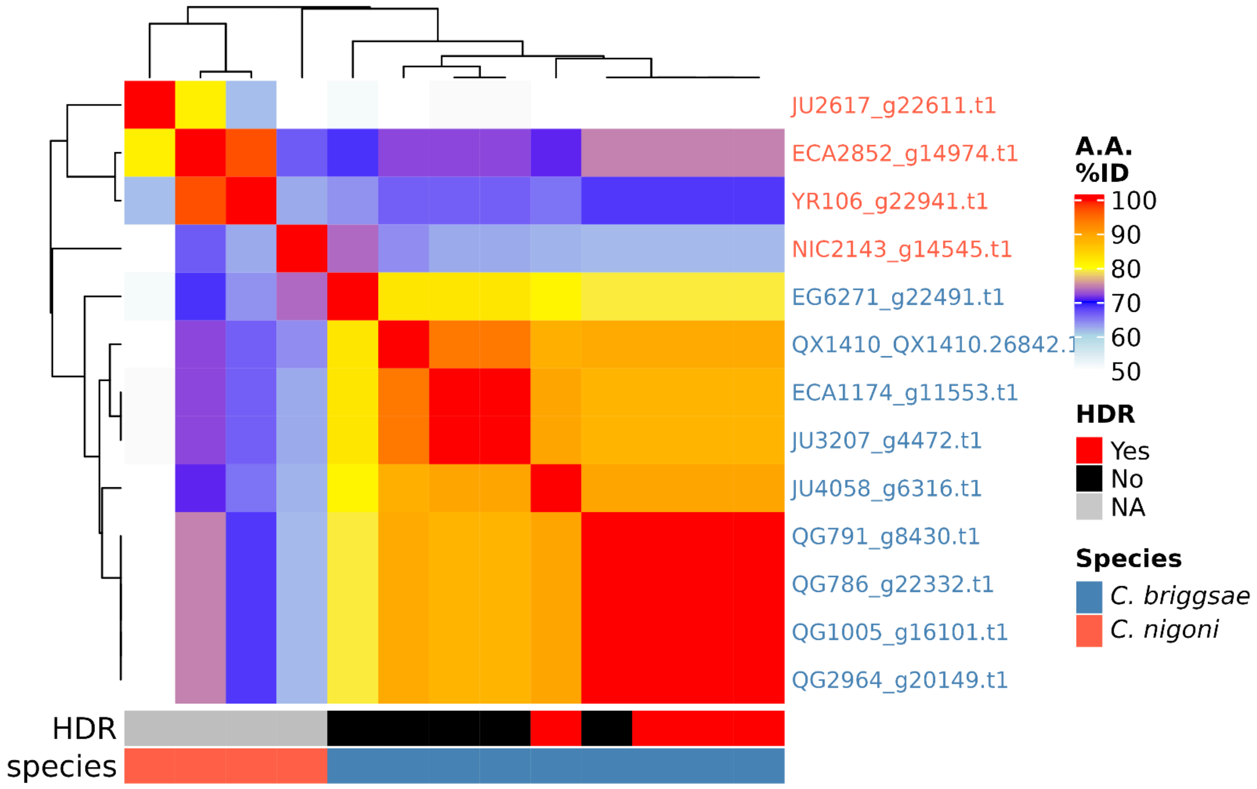

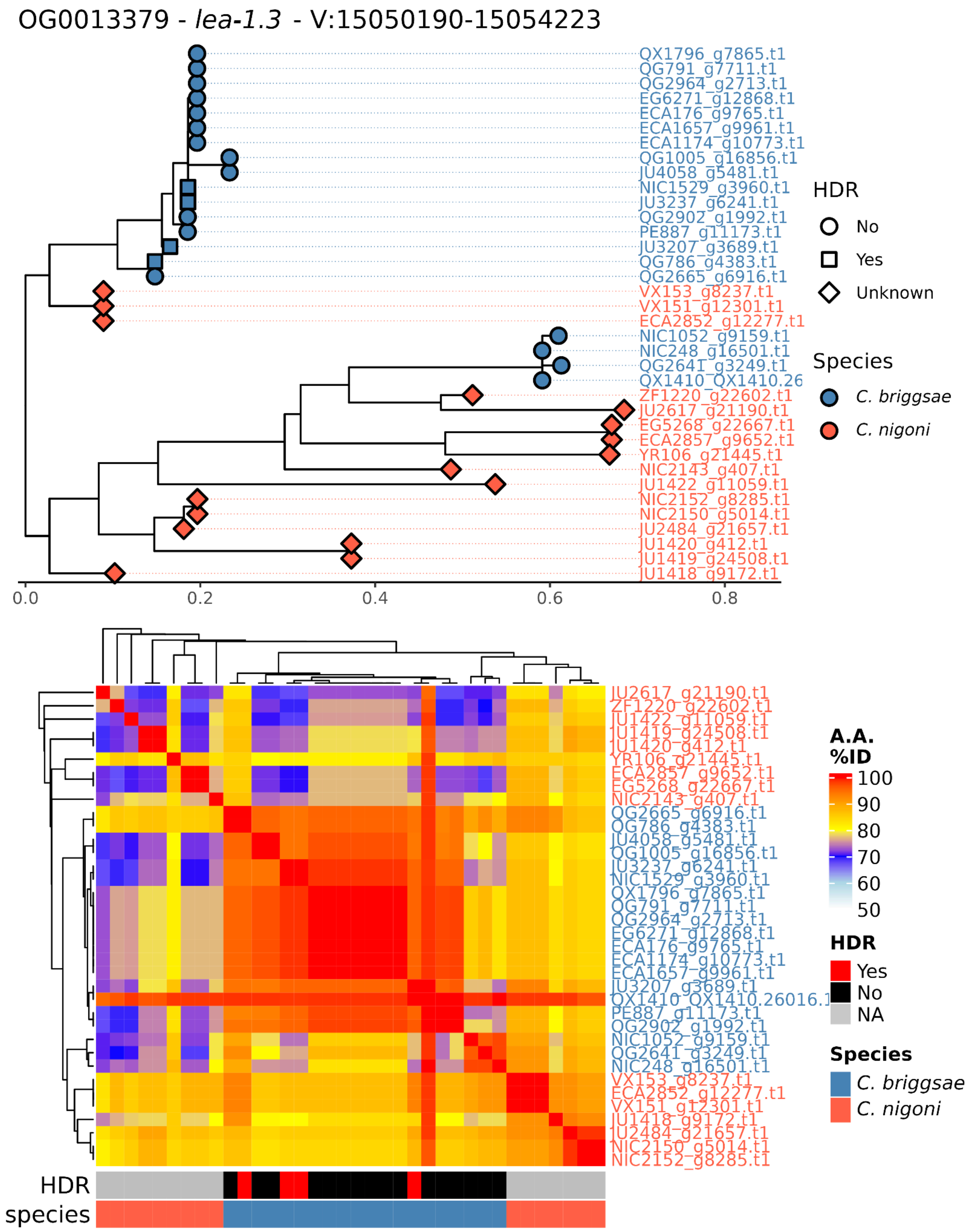
OG0013379 - lea-1.3 - V:15050190-15054223
QX1796_g7865.t1
QG791_g7711.t1
QG2964_g2713.t1
EG6271_g12868.t1
ECA176_g9765.t1
ECA1657_g9961.t1
ECA1174_g10773.t1
QG1005_g16856.t1
JU4058_g5481.t1
NIC1529_g3960.t1
JU3237_g6241.t1
QG2902_g1992.t1
PE887_g11173.t1
JU3207_g3689.t1
QG786_g4383.t1
QG2665_g6916.t1
VX153_g8237.t1
VX151_g12301.t1
ECA2852_g12277.t1
NIC1052_g9159.t1
NIC248_g16501.t1
QG2641_g3249.t1
QX1410_QX1410.26
ZF1220_g22602.t1
JU2617_g21190.t1
EG5268_g22667.t1
ECA2857_g9652.t1
YR106_g21445.t1
NIC2143_g407.t1
JU1422_g11059.t1
NIC2152_g8285.t1
NIC2150_g5014.t1
JU2484_g21657.t1
JU1420_g412.t1
JU1419_g24508.t1
JU1418_g9172.t1
0.0
0.2
0.4
0.6
0.8
HDR
No
Yes
Unknown
Species
C. briggsae
C. nigoni
JU2617_g21190.t1
ZF1220_g22602.t1
JU1422_g11059.t1
JU1419_g24508.t1
JU1420_g412.t1
YR106_g21445.t1
ECA2857_g9652.t1
EG5268_g22667.t1
NIC2143_g407.t1
QG2665_g6916.t1
QG786_g4383.t1
JU4058_g5481.t1
QG1005_g16856.t1
JU3237_g6241.t1
NIC1529_g3960.t1
QX1796_g7865.t1
QG791_g7711.t1
QG2964_g2713.t1
EG6271_g12868.t1
ECA176_g9765.t1
ECA1174_g10773.t1
ECA1657_g9961.t1
JU3207_g3689.t1
QX1410_QX1410.26016.t1
PE887_g11173.t1
QG2902_g1992.t1
NIC1052_g9159.t1
QG2641_g3249.t1
NIC248_g16501.t1
VX153_g8237.t1
ECA2852_g12277.t1
VX151_g12301.t1
JU1418_g9172.t1
JU2484_g21657.t1
NIC2150_g5014.t1
NIC2152_g8285.t1
A.A. %ID
100
90
80
70
60
50
HDR
Yes
No
NA
Species
C. briggsae
C. nigoni
HDR
species

OG0016702 - *Y82E9BR.22* - III:2327443-2327870

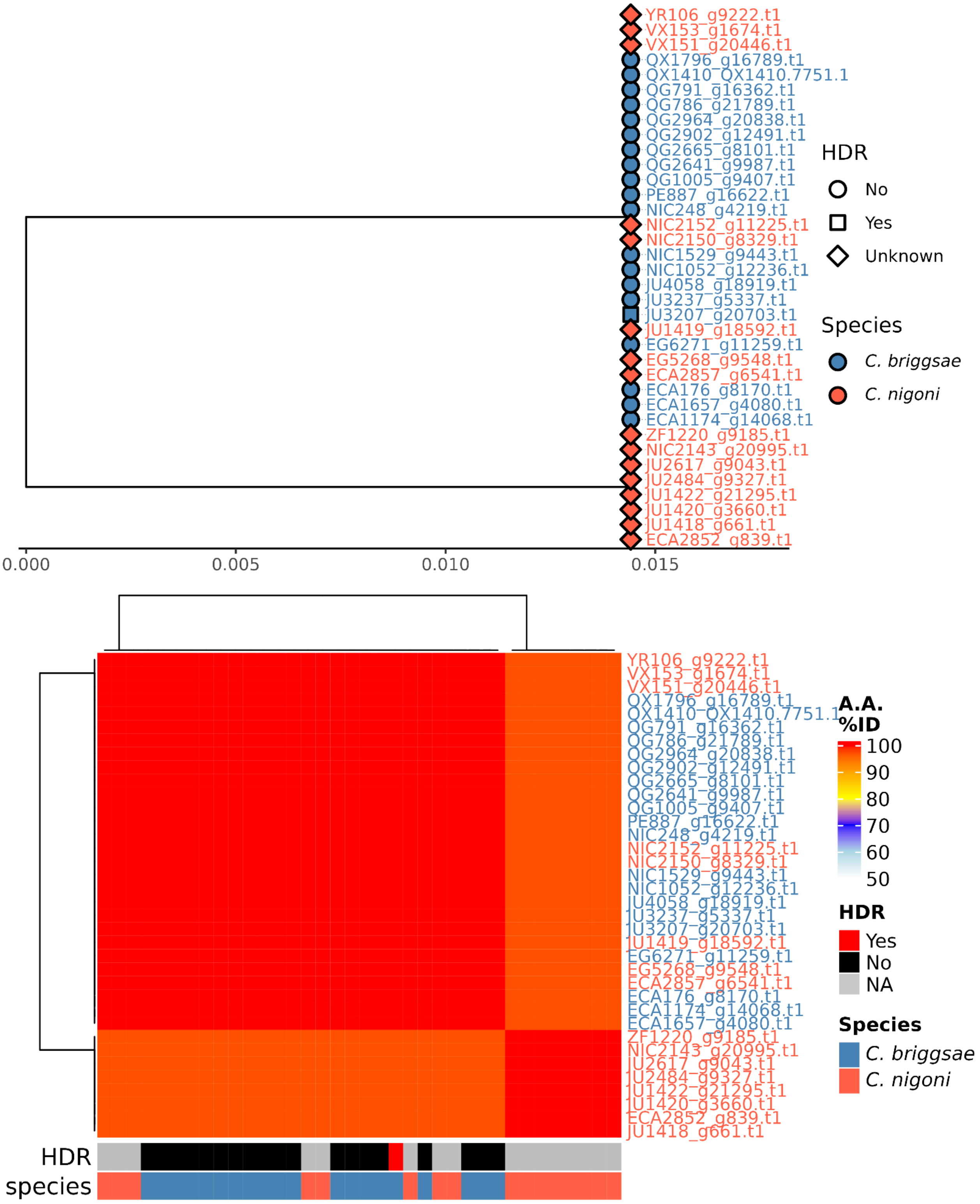

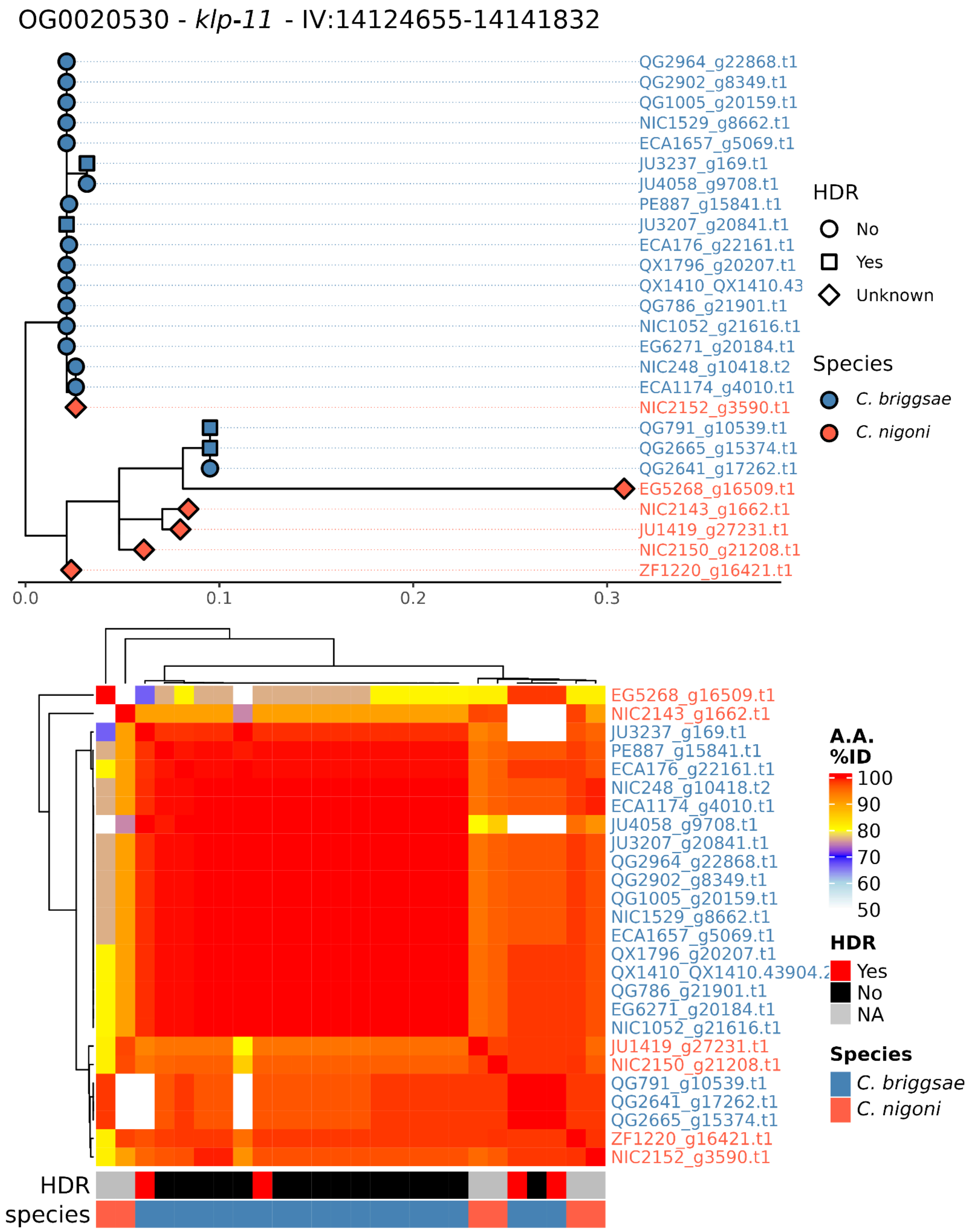
OG0020530 - klp-11 - IV:14124655-14141832
QG2964_g22868.t1
QG2902_g8349.t1
QG1005_g20159.t1
NIC1529_g8662.t1
ECA1657_g5069.t1
JU3237_g169.t1
JU4058_g9708.t1
PE887_g15841.t1
JU3207_g20841.t1
ECA176_g22161.t1
QX1796_g20207.t1
QX1410_QX1410.43
QG786_g21901.t1
NIC1052_g21616.t1
EG6271_g20184.t1
NIC248_g10418.t2
ECA1174_g4010.t1
NIC2152_g3590.t1
QG791_g10539.t1
QG2665_g15374.t1
QG2641_g17262.t1
EG5268_g16509.t1
NIC2143_g1662.t1
JU1419_g27231.t1
NIC2150_g21208.t1
ZF1220_g16421.t1
0.0
0.1
0.2
0.3
HDR
No
Yes
Unknown
Species
C. briggsae
C. nigoni
EG5268_g16509.t1
NIC2143_g1662.t1
JU3237_g169.t1
PE887_g15841.t1
ECA176_g22161.t1
NIC248_g10418.t2
ECA1174_g4010.t1
JU4058_g9708.t1
JU3207_g20841.t1
QG2964_g22868.t1
QG2902_g8349.t1
QG1005_g20159.t1
NIC1529_g8662.t1
ECA1657_g5069.t1
QX1796_g20207.t1
QX1410_QX1410.43904.2
QG786_g21901.t1
EG6271_g20184.t1
NIC1052_g21616.t1
JU1419_g27231.t1
NIC2150_g21208.t1
QG791_g10539.t1
QG2641_g17262.t1
QG2665_g15374.t1
ZF1220_g16421.t1
NIC2152_g3590.t1
HDR
species
A.A. %ID
100
90
80
70
60
50
HDR
Yes
No
NA
Species
C. briggsae
C. nigoni

OG0023405 - *QX1410.37579* - X:3840253-3841474

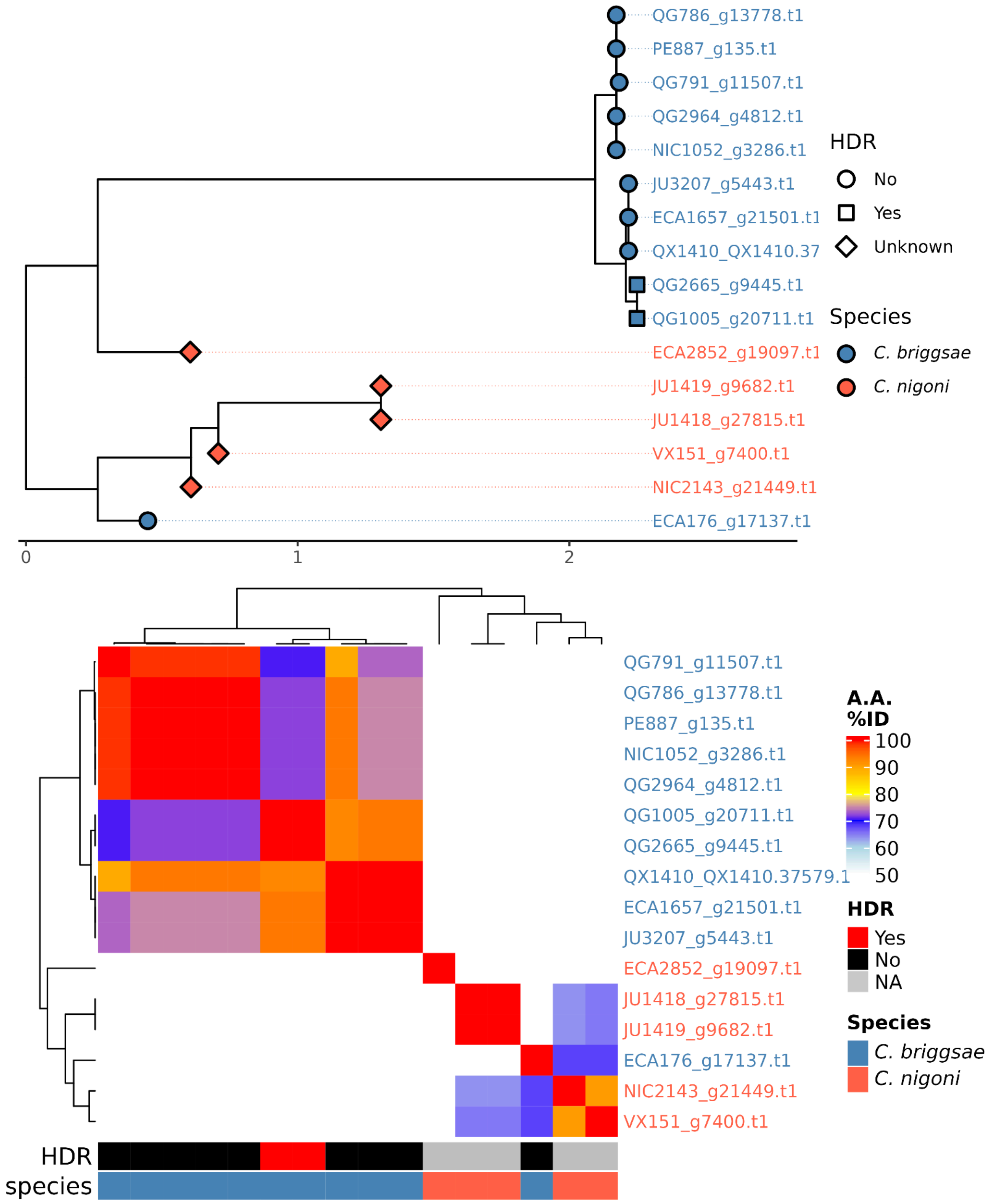

OG0013086 - *rpt-6* - III:1968729-1971601

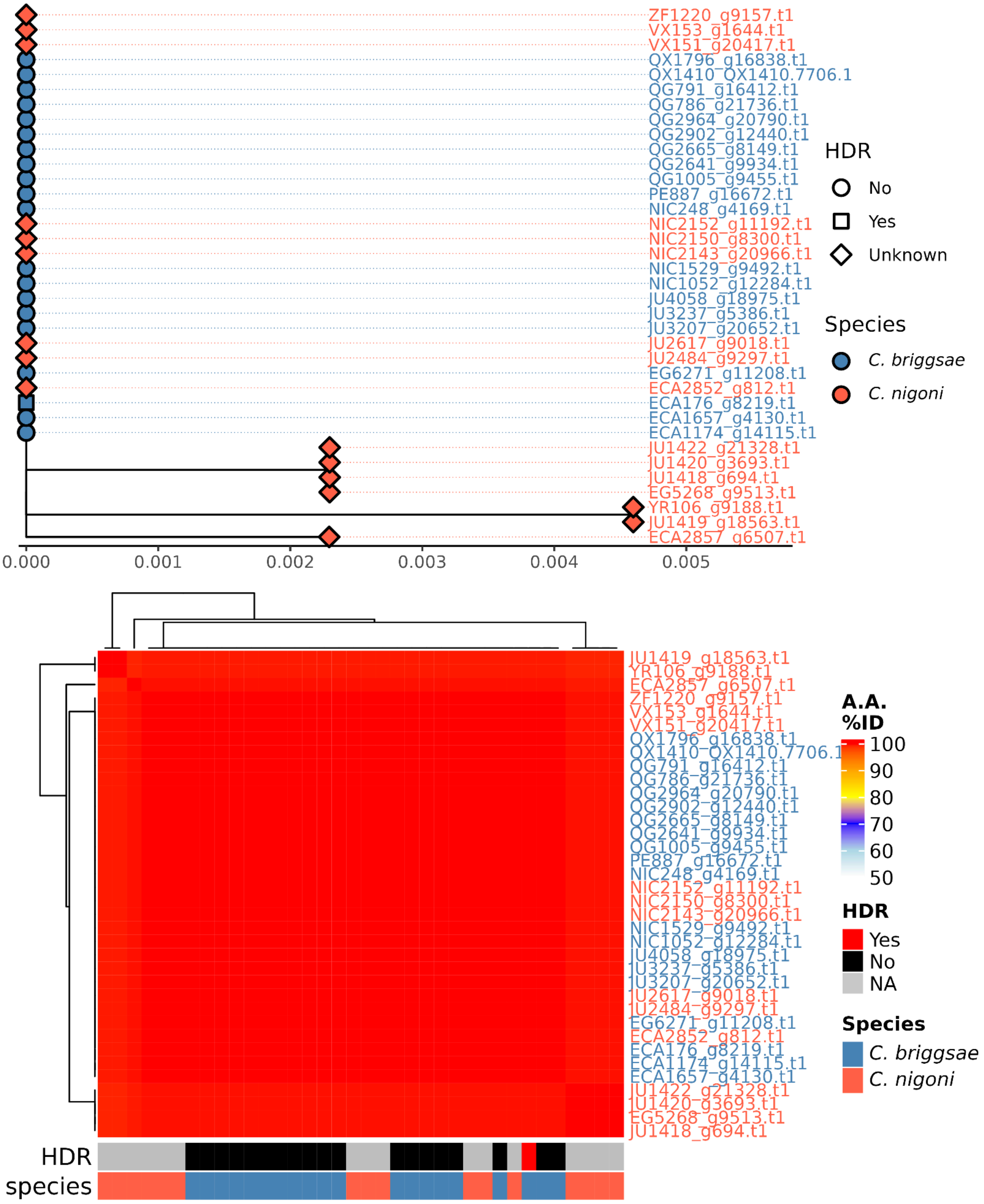

OG0018447 - *QX1410.44194* - IV:16307095-16308164

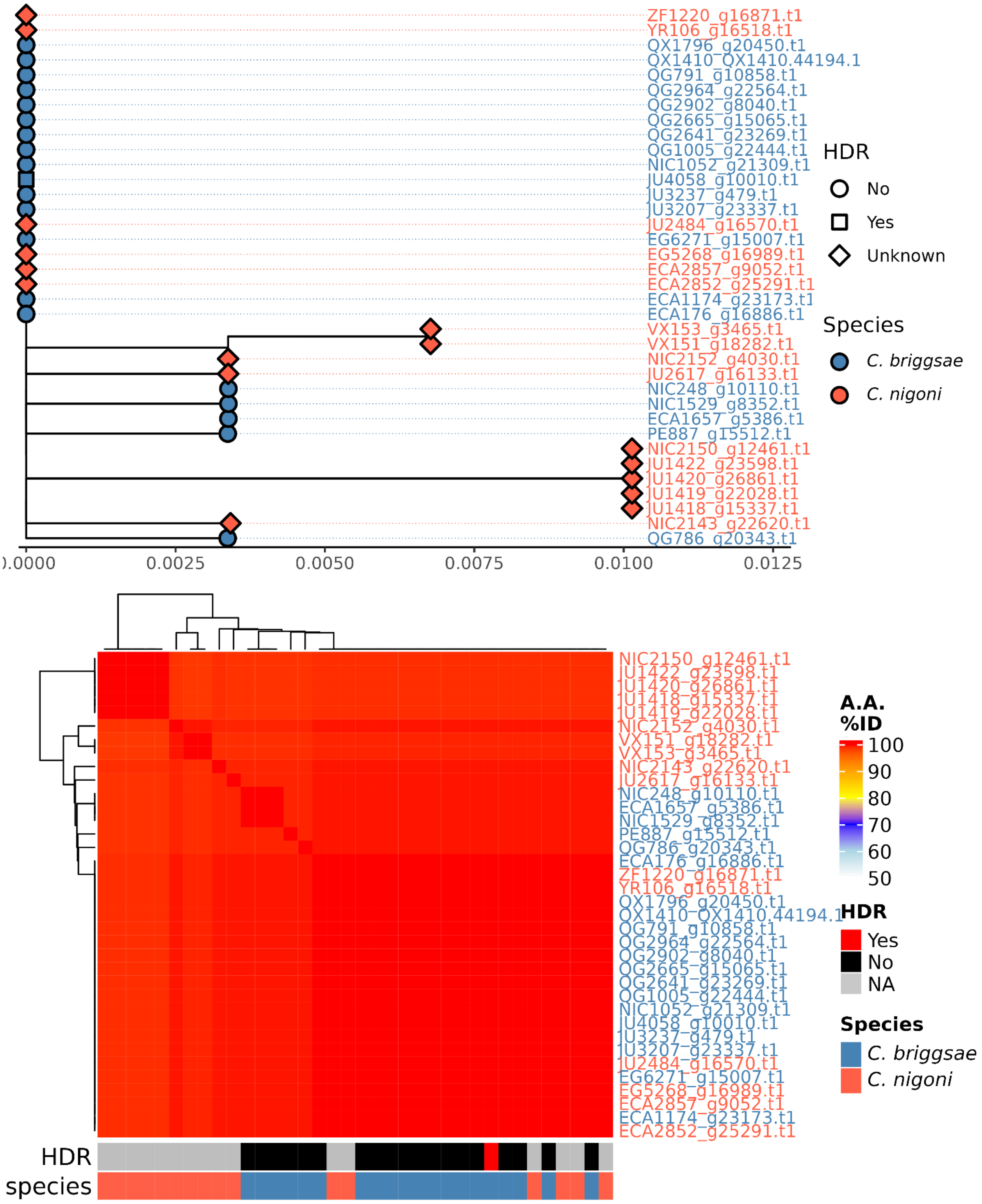

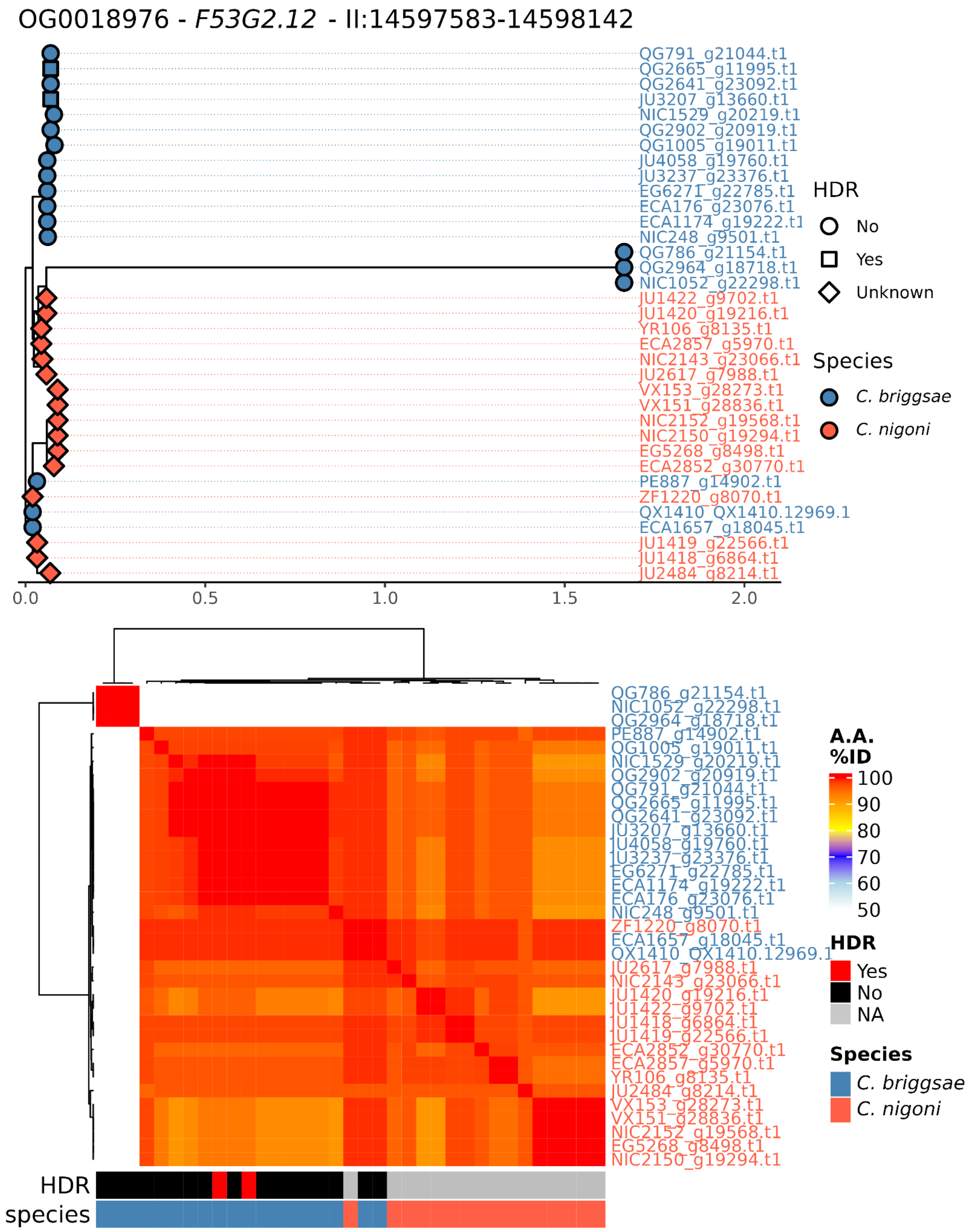

OG0018976 - F53G2.12 - II:14597583-14598142
QG791_g21044.t1
QG2665_g11995.t1
QG2641_g23092.t1
JU3207_g13660.t1
NIC1529_g20219.t1
QG2902_g20919.t1
QG1005_g19011.t1
JU4058_g19760.t1
JU3237_g23376.t1
EG6271_g22785.t1
ECA176_g23076.t1
ECA1174_g19222.t1
NIC248_g9501.t1
QG786_g21154.t1
QG2964_g18718.t1
NIC1052_g22298.t1
JU1422_g9702.t1
JU1420_g19216.t1
YR106_g8135.t1
ECA2857_g5970.t1
NIC2143_g23066.t1
JU2617_g7988.t1
VX153_g28273.t1
VX151_g28836.t1
NIC2152_g19568.t1
NIC2150_g19294.t1
EG5268_g8498.t1
ECA2852_g30770.t1
PE887_g14902.t1
ZF1220_g8070.t1
QX1410_QX1410.12969.1
ECA1657_g18045.t1
JU1419_g22566.t1
JU1418_g6864.t1
JU2484_g8214.t1
0.0
0.5
1.0
1.5
2.0
HDR
No
Yes
Unknown
Species
C. briggsae
C. nigoni
A.A. %ID
100
90
80
70
60
50
HDR
Yes
No
NA
Species
C. briggsae
C. nigoni
HDR
species

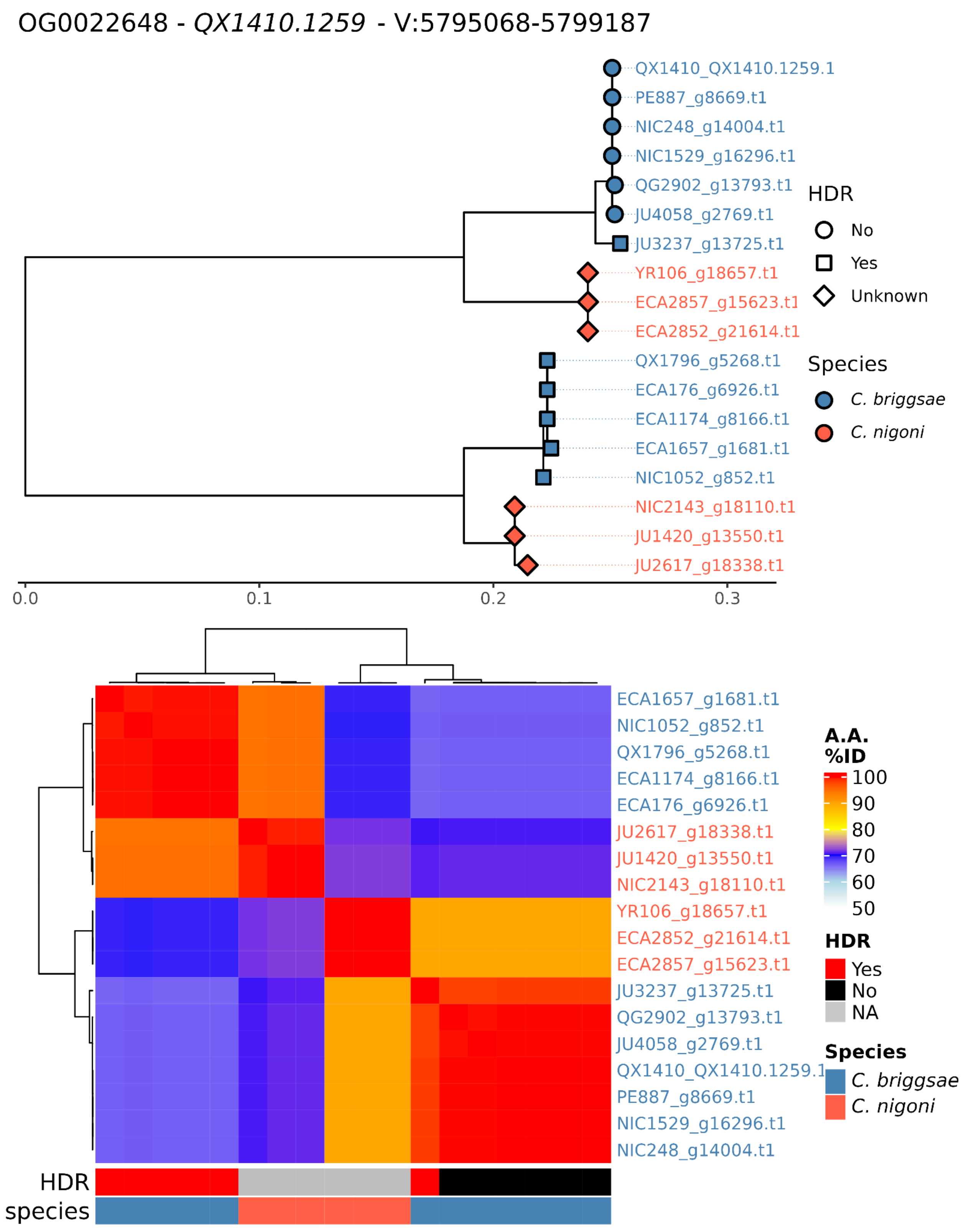
OG0022648 - QX1410.1259 - V:5795068-5799187
QX1410_QX1410.1259.1
PE887_g8669.t1
NIC248_g14004.t1
NIC1529_g16296.t1
QG2902_g13793.t1
JU4058_g2769.t1
JU3237_g13725.t1
YR106_g18657.t1
ECA2857_g15623.t1
ECA2852_g21614.t1
QX1796_g5268.t1
ECA176_g6926.t1
ECA1174_g8166.t1
ECA1657_g1681.t1
NIC1052_g852.t1
NIC2143_g18110.t1
JU1420_g13550.t1
JU2617_g18338.t1
0.0
0.1
0.2
0.3
HDR
No
Yes
Unknown
Species
C. briggsae
C. nigoni
A.A. %ID
100
90
80
70
60
50
HDR
Yes
No
NA
Species
C. briggsae
C. nigoni
HDR
species

OG0024532 - *nlp-82* - II:656478-658736

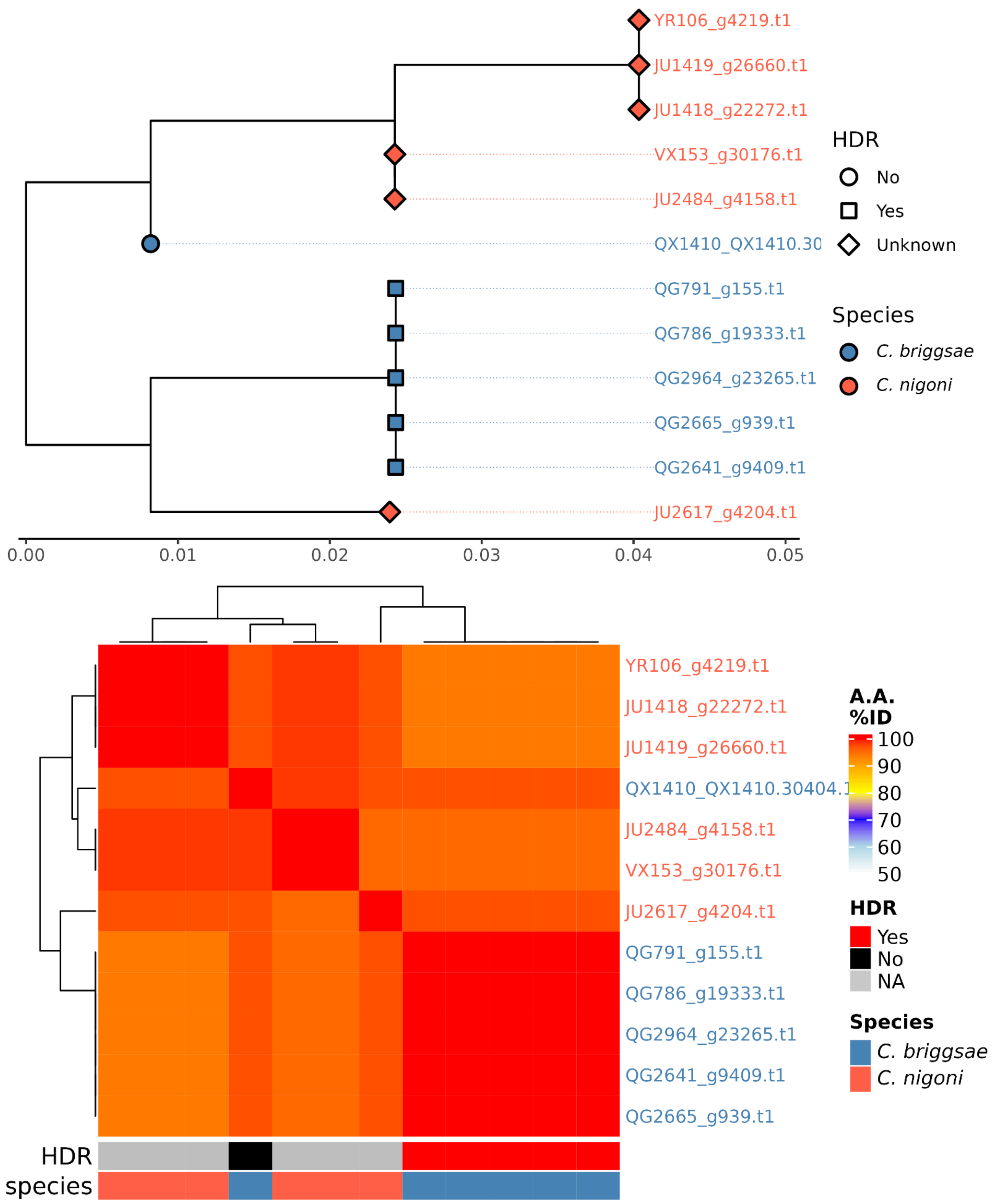

OG0014032 - *C09G1.5* - X:2688312-2690719

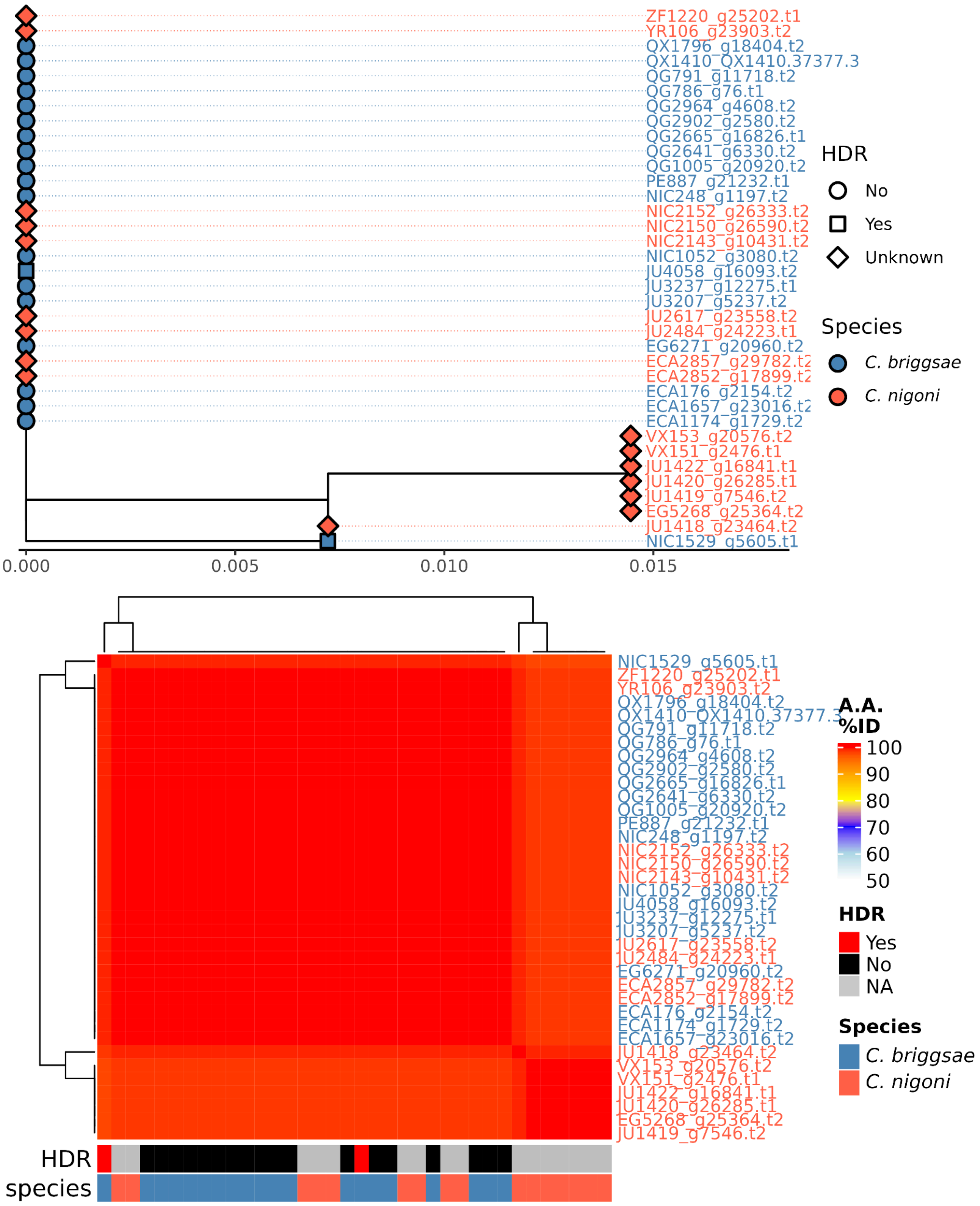

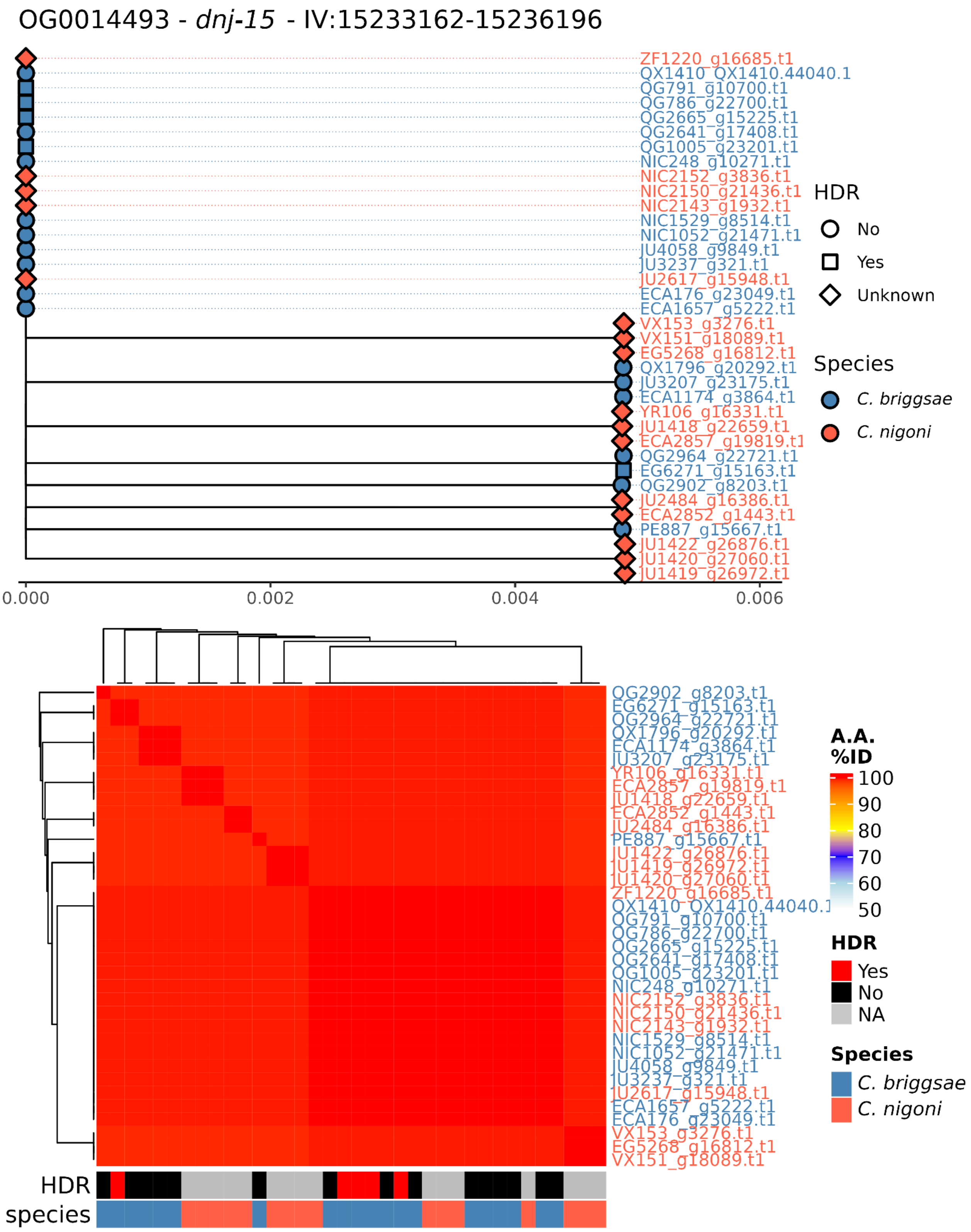
OG0014493 - dnj-15 - IV:15233162-15236196
ZF1220_g16685.t1
QX1410_QX1410.44040.1
QG791_g10700.t1
QG786_g22700.t1
QG2665_g15225.t1
QG2641_g17408.t1
QG1005_g23201.t1
NIC248_g10271.t1
NIC2152_g3836.t1
NIC2150_g21436.t1
NIC2143_g1932.t1
NIC1529_g8514.t1
NIC1052_g21471.t1
JU4058_g9849.t1
JU3237_g321.t1
JU2617_g15948.t1
ECA176_g23049.t1
ECA1657_g5222.t1
VX153_g3276.t1
VX151_g18089.t1
EG5268_g16812.t1
QX1796_g20292.t1
JU3207_g23175.t1
ECA1174_g3864.t1
YR106_g16331.t1
JU1418_g22659.t1
ECA2857_g19819.t1
QG2964_g22721.t1
EG6271_g15163.t1
QG2902_g8203.t1
JU2484_g16386.t1
ECA2852_g1443.t1
PE887_g15667.t1
JU1422_g26876.t1
JU1420_g27060.t1
JU1419_g26972.t1
0.000
0.002
0.004
0.006
HDR
No
Yes
Unknown
Species
C. briggsae
C. nigoni
QG2902_g8203.t1
EG6271_g15163.t1
QG2964_g22721.t1
QX1796_g20292.t1
ECA1174_g3864.t1
JU3207_g23175.t1
YR106_g16331.t1
ECA2857_g19819.t1
JU1418_g22659.t1
ECA2852_g1443.t1
JU2484_g16386.t1
PE887_g15667.t1
JU1422_g26876.t1
JU1419_g26972.t1
JU1420_g27060.t1
ZF1220_g16685.t1
QX1410_QX1410.44040.1
QG791_g10700.t1
QG786_g22700.t1
QG2665_g15225.t1
QG2641_g17408.t1
QG1005_g23201.t1
NIC248_g10271.t1
NIC2152_g3836.t1
NIC2150_g21436.t1
NIC2143_g1932.t1
NIC1529_g8514.t1
NIC1052_g21471.t1
JU4058_g9849.t1
JU3237_g321.t1
JU2617_g15948.t1
ECA1657_g5222.t1
ECA176_g23049.t1
VX153_g3276.t1
EG5268_g16812.t1
VX151_g18089.t1
HDR
species
A.A. %ID
100
90
80
70
60
50
HDR
Yes
No
NA
Species
C. briggsae
C. nigoni

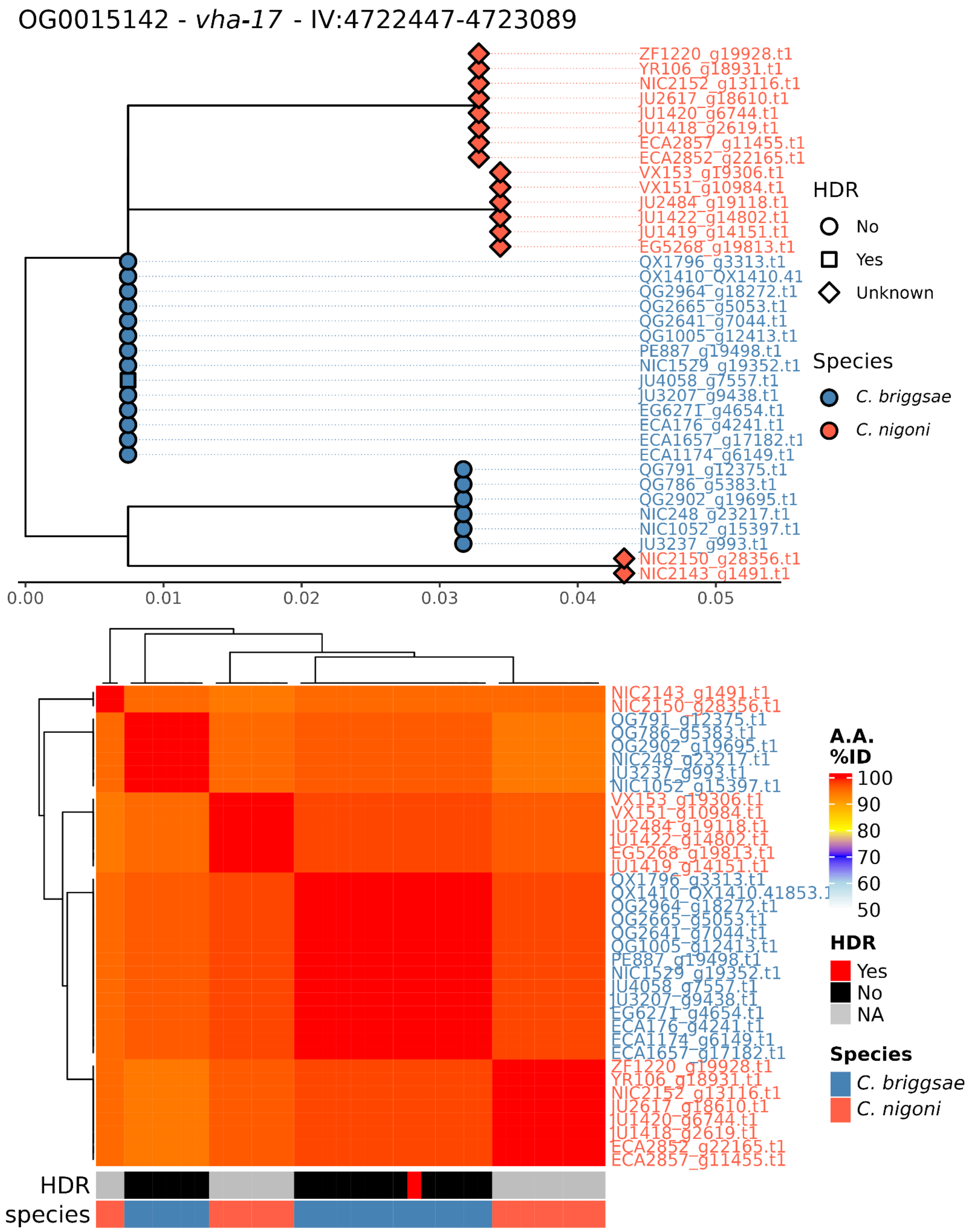
OG0015142 - vha-17 - IV:4722447-4723089
ZF1220_g19928.t1
YR106_g18931.t1
NIC2152_g13116.t1
JU2617_g18610.t1
JU1420_g6744.t1
JU1418_g2619.t1
ECA2857_g11455.t1
ECA2852_g22165.t1
VX153_g19306.t1
VX151_g10984.t1
JU2484_g19118.t1
JU1422_g14802.t1
JU1419_g14151.t1
EG5268_g19813.t1
QX1796_g3313.t1
QX1410_QX1410.41
QG2964_g18272.t1
QG2665_g5053.t1
QG2641_g7044.t1
QG1005_g12413.t1
PE887_g19498.t1
NIC1529_g19352.t1
JU4058_g7557.t1
JU3207_g9438.t1
EG6271_g4654.t1
ECA176_g4241.t1
ECA1657_g17182.t1
ECA1174_g6149.t1
QG791_g12375.t1
QG786_g5383.t1
QG2902_g19695.t1
NIC248_g23217.t1
NIC1052_g15397.t1
JU3237_g993.t1
NIC2150_g28356.t1
NIC2143_g1491.t1
0.00
0.01
0.02
0.03
0.04
0.05
HDR
No
Yes
Unknown
Species
C. briggsae
C. nigoni
NIC2143_g1491.t1
NIC2150_g28356.t1
QG791_g12375.t1
QG786_g5383.t1
QG2902_g19695.t1
NIC248_g23217.t1
JU3237_g993.t1
NIC1052_g15397.t1
VX153_g19306.t1
VX151_g10984.t1
JU2484_g19118.t1
JU1422_g14802.t1
EG5268_g19813.t1
JU1419_g14151.t1
QX1796_g3313.t1
QX1410_QX1410.41853.1
QG2964_g18272.t1
QG2665_g5053.t1
QG2641_g7044.t1
QG1005_g12413.t1
PE887_g19498.t1
NIC1529_g19352.t1
JU4058_g7557.t1
JU3207_g9438.t1
EG6271_g4654.t1
ECA176_g4241.t1
ECA1174_g6149.t1
ECA1657_g17182.t1
ZF1220_g19928.t1
YR106_g18931.t1
NIC2152_g13116.t1
JU2617_g18610.t1
JU1420_g6744.t1
JU1418_g2619.t1
ECA2852_g22165.t1
ECA2857_g11455.t1
A.A. %ID
100
90
80
70
60
50
HDR
Yes
No
NA
Species
C. briggsae
C. nigoni
HDR
species

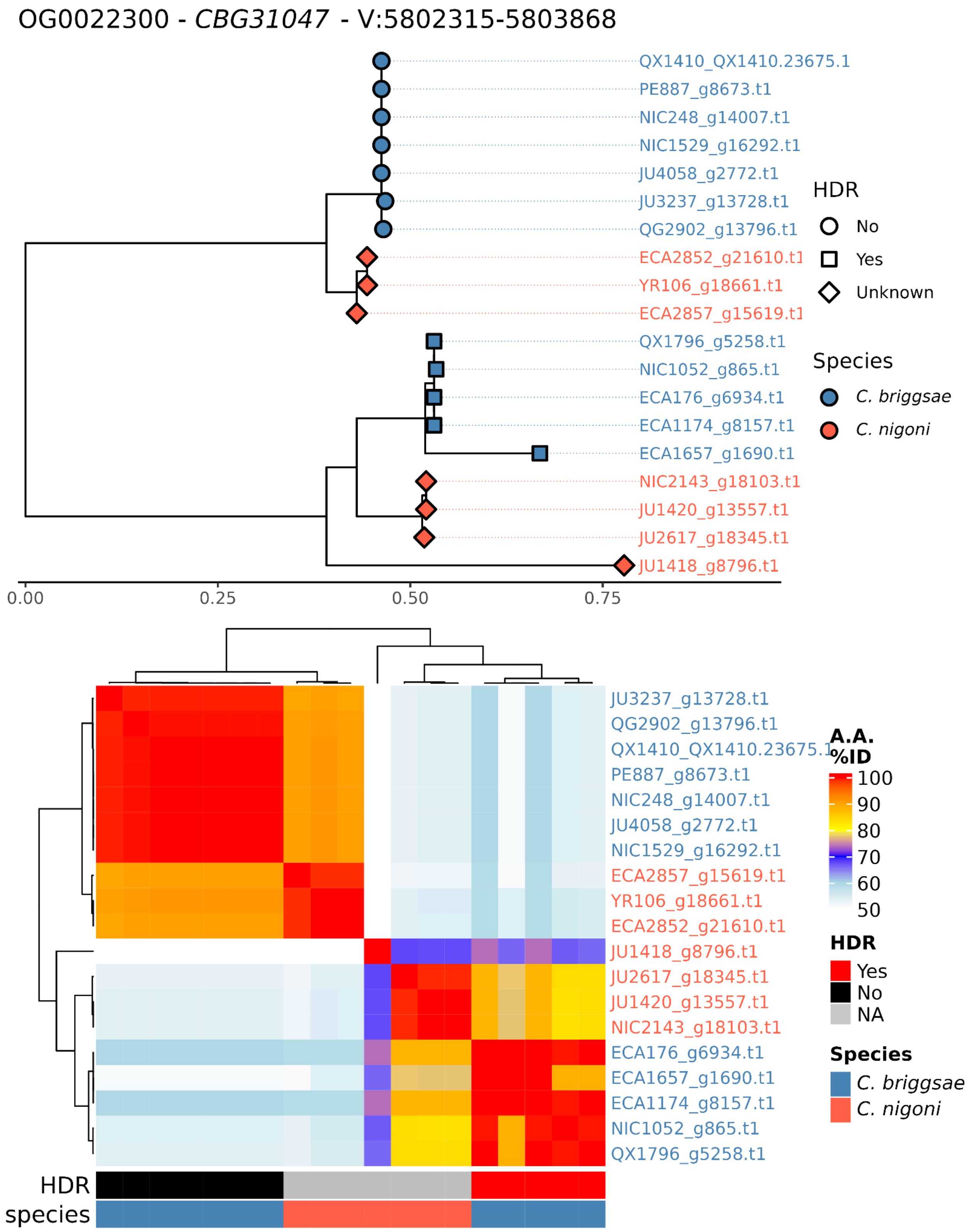
OG0022300 - CBG31047 - V:5802315-5803868
QX1410_QX1410.23675.1
PE887_g8673.t1
NIC248_g14007.t1
NIC1529_g16292.t1
JU4058_g2772.t1
JU3237_g13728.t1
QG2902_g13796.t1
ECA2852_g21610.t1
YR106_g18661.t1
ECA2857_g15619.t1
QX1796_g5258.t1
NIC1052_g865.t1
ECA176_g6934.t1
ECA1174_g8157.t1
ECA1657_g1690.t1
NIC2143_g18103.t1
JU1420_g13557.t1
JU2617_g18345.t1
JU1418_g8796.t1
HDR
No
Yes
Unknown
Species
C. briggsae
C. nigoni
0.00
0.25
0.50
0.75
JU3237_g13728.t1
QG2902_g13796.t1
QX1410_QX1410.23675.1
PE887_g8673.t1
NIC248_g14007.t1
JU4058_g2772.t1
NIC1529_g16292.t1
ECA2857_g15619.t1
YR106_g18661.t1
ECA2852_g21610.t1
JU1418_g8796.t1
JU2617_g18345.t1
JU1420_g13557.t1
NIC2143_g18103.t1
ECA176_g6934.t1
ECA1657_g1690.t1
ECA1174_g8157.t1
NIC1052_g865.t1
QX1796_g5258.t1
A.A. %ID
100
90
80
70
60
50
HDR
Yes
No
NA
Species
C. briggsae
C. nigoni
HDR
species

OG0014054 - *gpx-3* - X:2393194-2394378

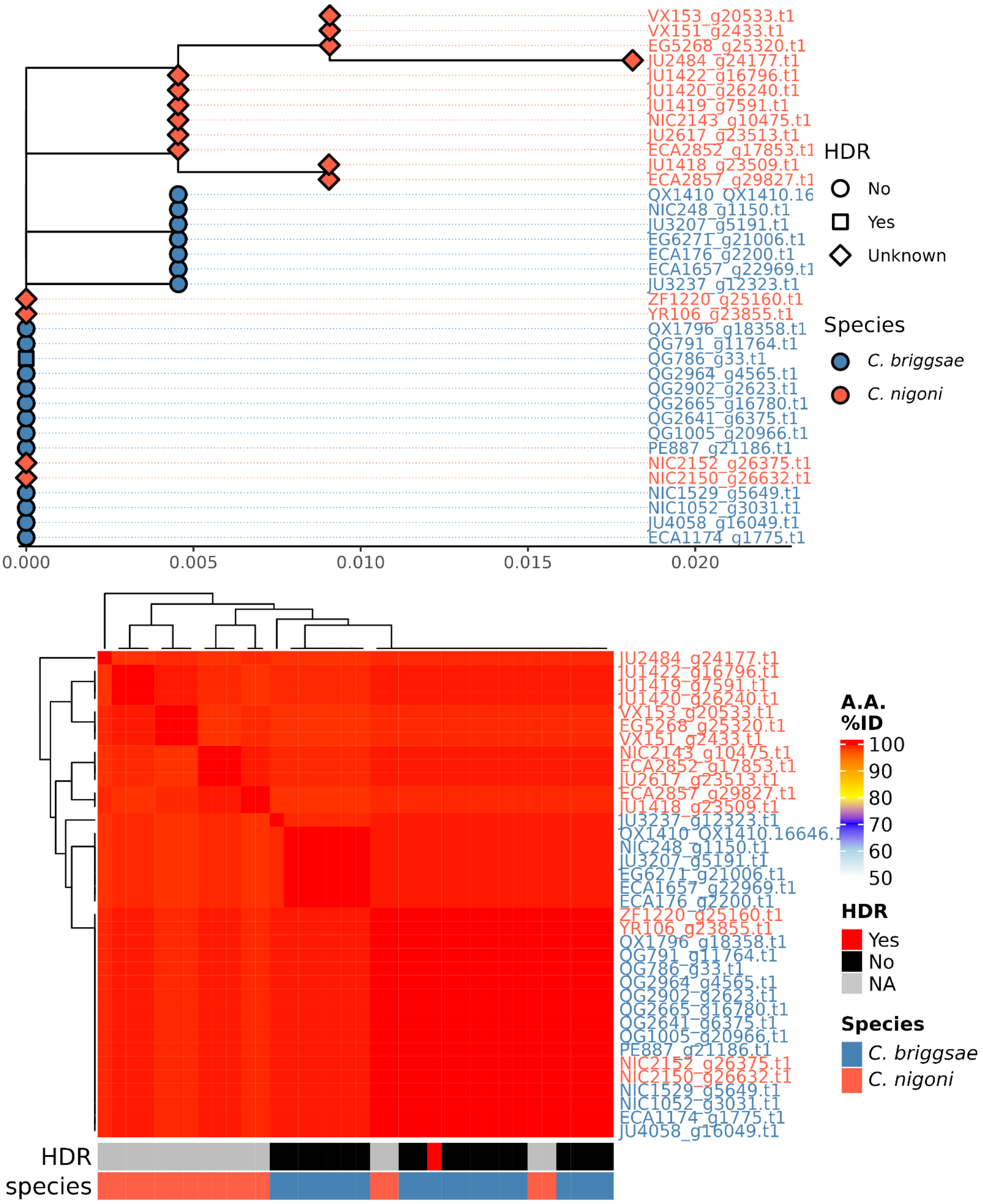

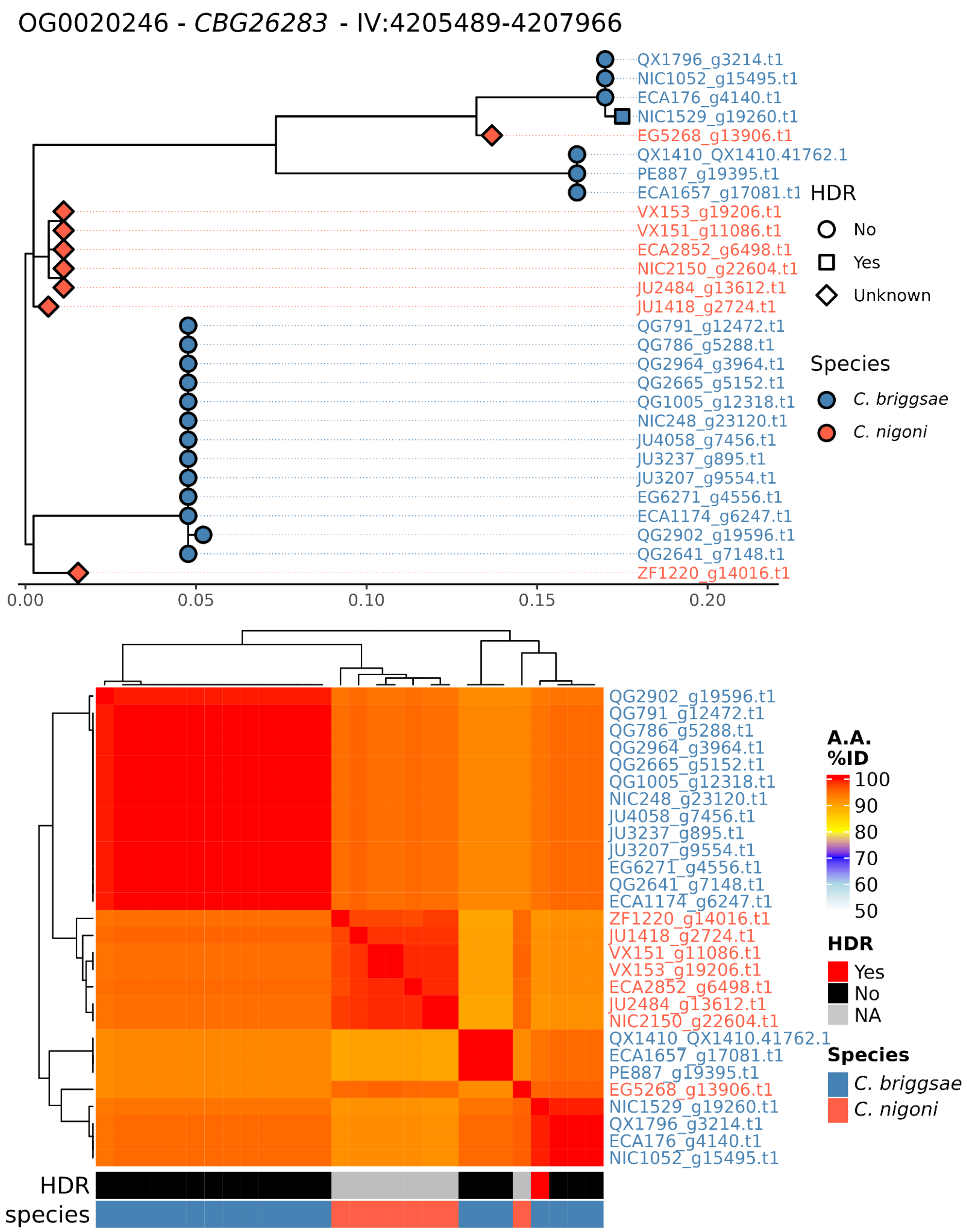

OG0020246 - *CBG26283* - IV:4205489-4207966
QX1796_g3214.t1
NIC1052_g15495.t1
ECA176_g4140.t1
NIC1529_g19260.t1
EG5268_g13906.t1
QX1410_QX1410.41762.1
PE887_g19395.t1
ECA1657_g17081.t1
VX153_g19206.t1
VX151_g11086.t1
ECA2852_g6498.t1
NIC2150_g22604.t1
JU2484_g13612.t1
JU1418_g2724.t1
QG791_g12472.t1
QG786_g5288.t1
QG2964_g3964.t1
QG2665_g5152.t1
QG1005_g12318.t1
NIC248_g23120.t1
JU4058_g7456.t1
JU3237_g895.t1
JU3207_g9554.t1
EG6271_g4556.t1
ECA1174_g6247.t1
QG2902_g19596.t1
QG2641_g7148.t1
ZF1220_g14016.t1
HDR
No
Yes
Unknown
Species
*C. briggsae*
*C. nigoni*
0.00
0.05
0.10
0.15
0.20
QG2902_g19596.t1
QG791_g12472.t1
QG786_g5288.t1
QG2964_g3964.t1
QG2665_g5152.t1
QG1005_g12318.t1
NIC248_g23120.t1
JU4058_g7456.t1
JU3237_g895.t1
JU3207_g9554.t1
EG6271_g4556.t1
QG2641_g7148.t1
ECA1174_g6247.t1
ZF1220_g14016.t1
JU1418_g2724.t1
VX151_g11086.t1
VX153_g19206.t1
ECA2852_g6498.t1
JU2484_g13612.t1
NIC2150_g22604.t1
QX1410_QX1410.41762.1
ECA1657_g17081.t1
PE887_g19395.t1
EG5268_g13906.t1
NIC1529_g19260.t1
QX1796_g3214.t1
ECA176_g4140.t1
NIC1052_g15495.t1
A.A. %ID
100
90
80
70
60
50
HDR
Yes
No
NA
Species
*C. briggsae*
*C. nigoni*
HDR
species

OG0013418 - *mnat-1* - II:14598184-14605129

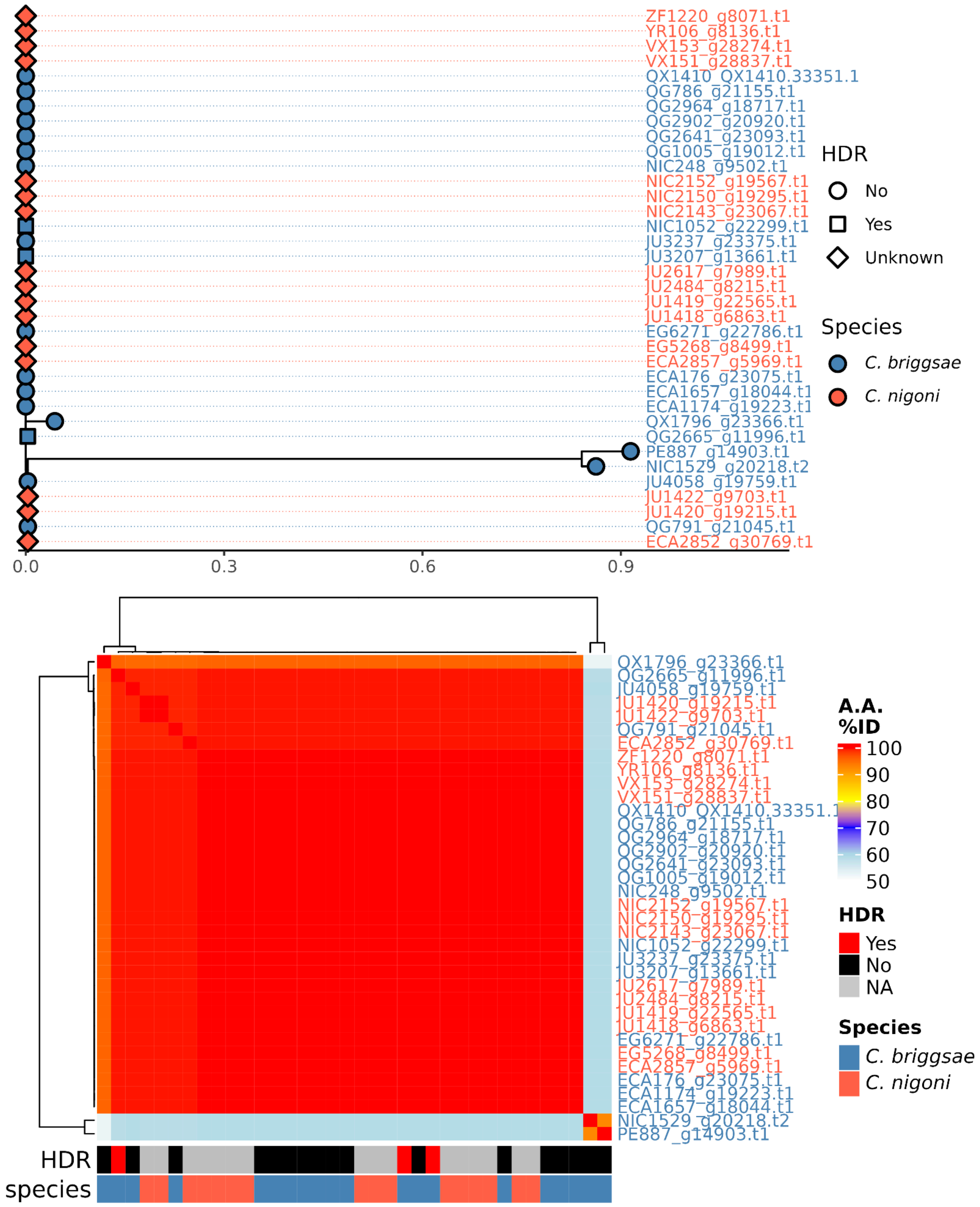

OG0015238 - *Y7A9D.1* - IV:568769-569746

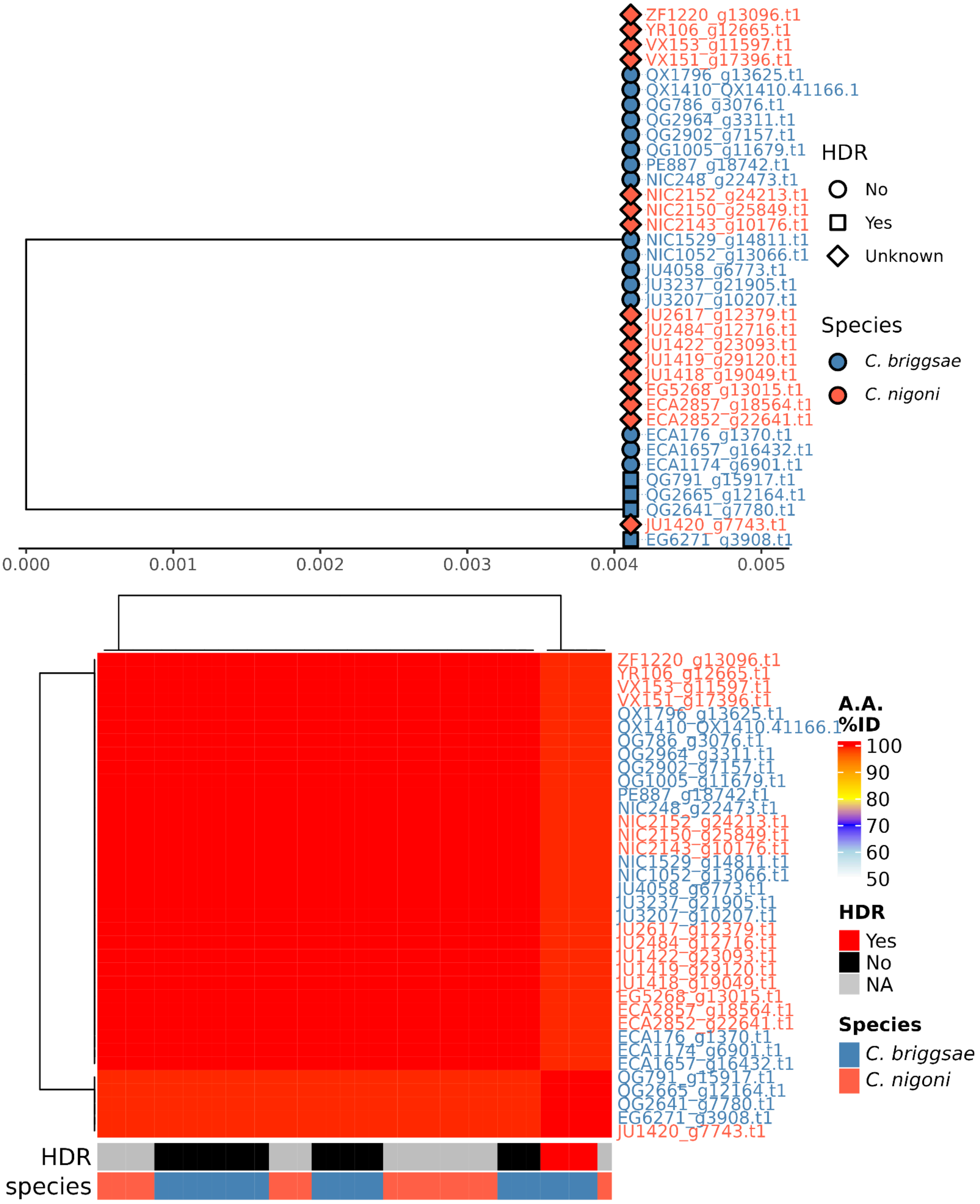

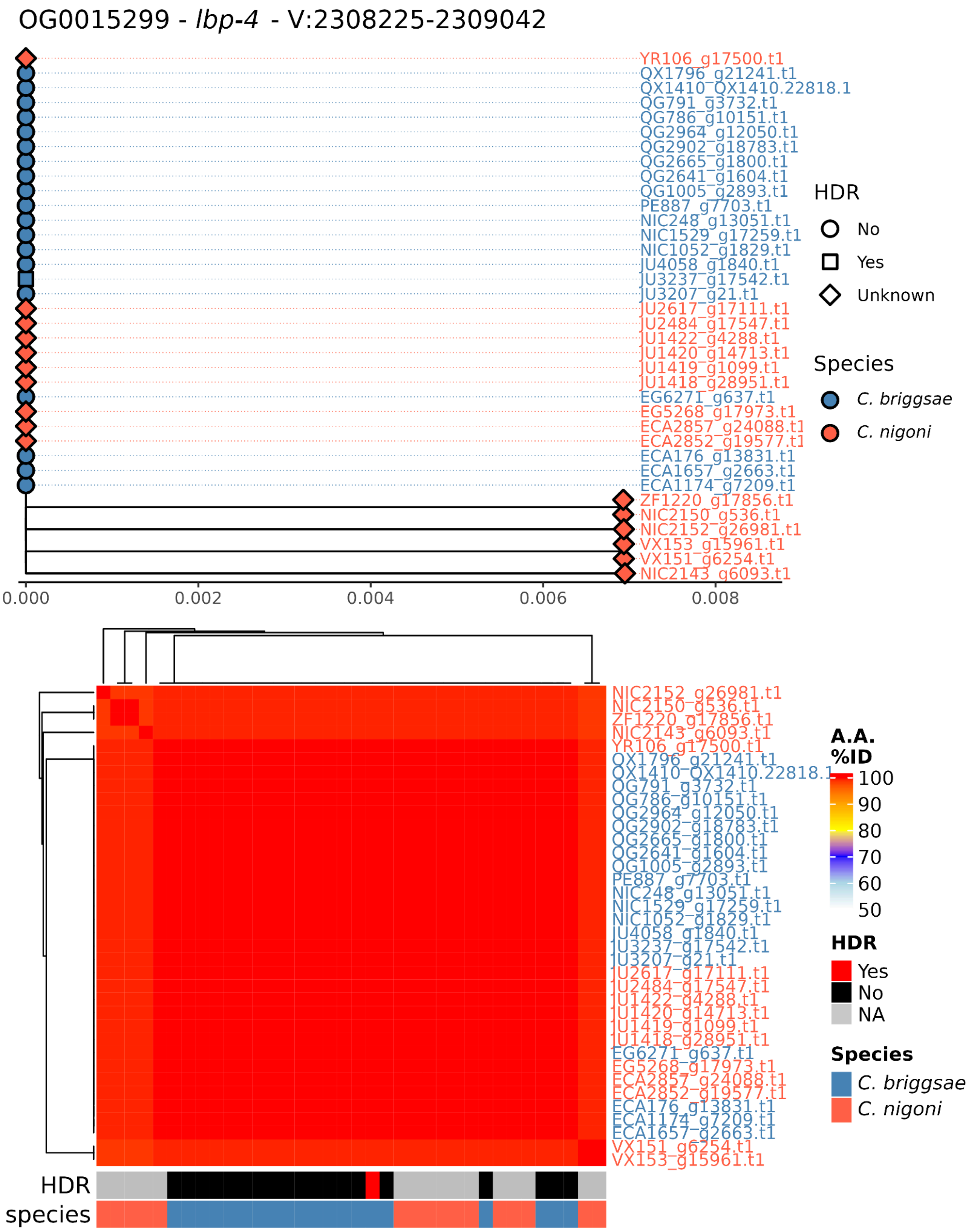
OG0015299 - lbp-4 - V:2308225-2309042
YR106_g17500.t1
QX1796_g21241.t1
QX1410_QX1410.22818.1
QG791_g3732.t1
QG786_g10151.t1
QG2964_g12050.t1
QG2902_g18783.t1
QG2665_g1800.t1
QG2641_g1604.t1
QG1005_g2893.t1
PE887_g7703.t1
NIC248_g13051.t1
NIC1529_g17259.t1
NIC1052_g1829.t1
JU4058_g1840.t1
JU3237_g17542.t1
JU3207_g21.t1
JU2617_g17111.t1
JU2484_g17547.t1
JU1422_g4288.t1
JU1420_g14713.t1
JU1419_g1099.t1
JU1418_g28951.t1
EG6271_g637.t1
EG5268_g17973.t1
ECA2857_g24088.t1
ECA2852_g19577.t1
ECA176_g13831.t1
ECA1657_g2663.t1
ECA1174_g7209.t1
ZF1220_g17856.t1
NIC2150_g536.t1
NIC2152_g26981.t1
VX153_g15961.t1
VX151_g6254.t1
NIC2143_g6093.t1
0.000
0.002
0.004
0.006
0.008
HDR
No
Yes
Unknown
Species
C. briggsae
C. nigoni
NIC2152_g26981.t1
NIC2150_g536.t1
ZF1220_g17856.t1
NIC2143_g6093.t1
YR106_g17500.t1
QX1796_g21241.t1
QX1410_QX1410.22818.1
QG791_g3732.t1
QG786_g10151.t1
QG2964_g12050.t1
QG2902_g18783.t1
QG2665_g1800.t1
QG2641_g1604.t1
QG1005_g2893.t1
PE887_g7703.t1
NIC248_g13051.t1
NIC1529_g17259.t1
NIC1052_g1829.t1
JU4058_g1840.t1
JU3237_g17542.t1
JU3207_g21.t1
JU2617_g17111.t1
JU2484_g17547.t1
JU1422_g4288.t1
JU1420_g14713.t1
JU1419_g1099.t1
JU1418_g28951.t1
EG6271_g637.t1
EG5268_g17973.t1
ECA2857_g24088.t1
ECA2852_g19577.t1
ECA176_g13831.t1
ECA1174_g7209.t1
ECA1657_g2663.t1
VX151_g6254.t1
VX153_g15961.t1
A.A. %ID
100
90
80
70
60
50
HDR
Yes
No
NA
Species
C. briggsae
C. nigoni
HDR
species

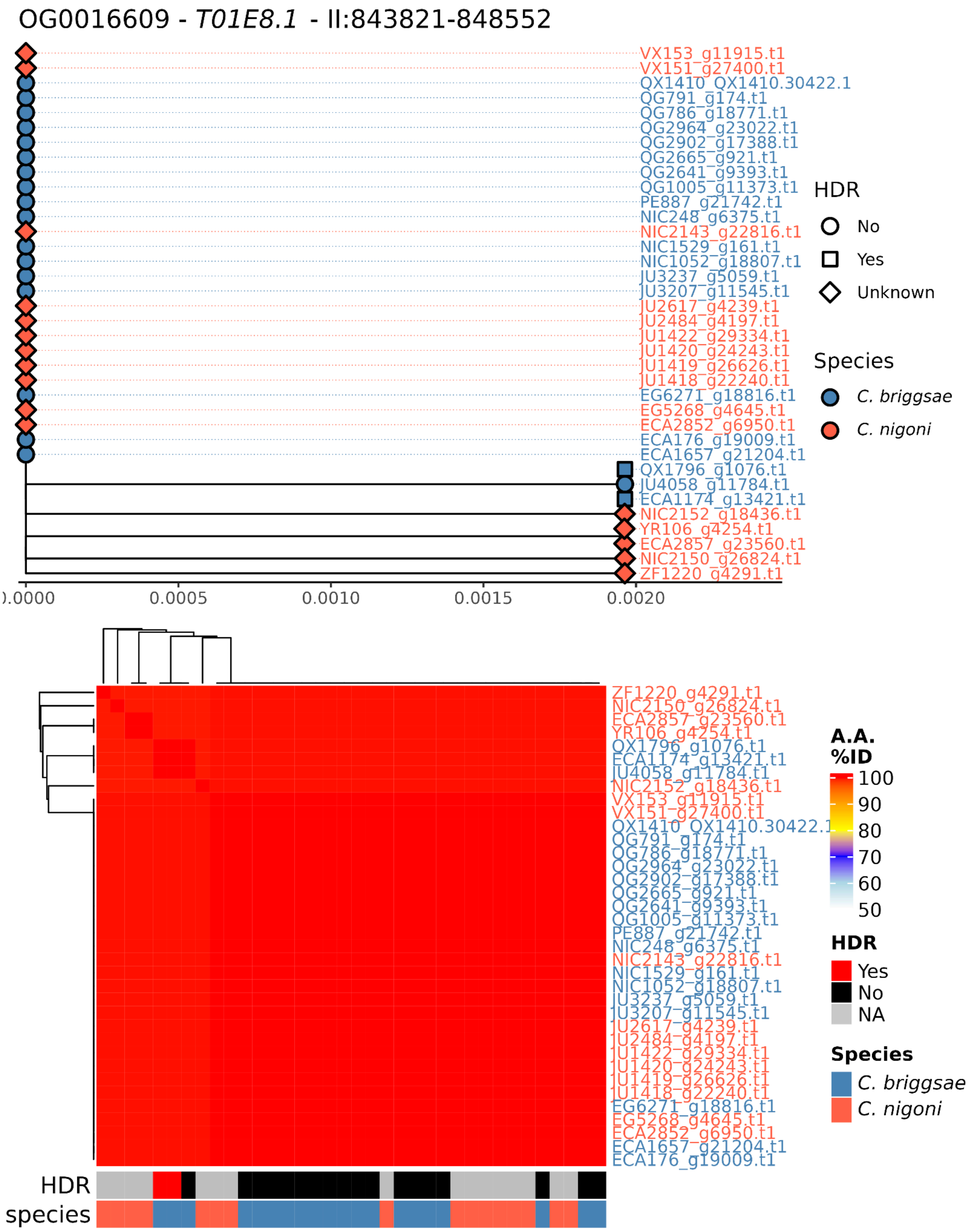
OG0016609 - T01E8.1 - II:843821-848552
VX153_g11915.t1
VX151_g27400.t1
QX1410_QX1410.30422.1
QG791_g174.t1
QG786_g18771.t1
QG2964_g23022.t1
QG2902_g17388.t1
QG2665_g921.t1
QG2641_g9393.t1
QG1005_g11373.t1
PE887_g21742.t1
NIC248_g6375.t1
NIC2143_g22816.t1
NIC1529_g161.t1
NIC1052_g18807.t1
JU3237_g5059.t1
JU3207_g11545.t1
JU2617_g4239.t1
JU2484_g4197.t1
JU1422_g29334.t1
JU1420_g24243.t1
JU1419_g26626.t1
JU1418_g22240.t1
EG6271_g18816.t1
EG5268_g4645.t1
ECA2852_g6950.t1
ECA176_g19009.t1
ECA1657_g21204.t1
QX1796_g1076.t1
JU4058_g11784.t1
ECA1174_g13421.t1
NIC2152_g18436.t1
YR106_g4254.t1
ECA2857_g23560.t1
NIC2150_g26824.t1
ZF1220_g4291.t1
0.0000
0.0005
0.0010
0.0015
0.0020
HDR
No
Yes
Unknown
Species
C. briggsae
C. nigoni
A.A. %ID
100
90
80
70
60
50
HDR
Yes
No
NA
Species
C. briggsae
C. nigoni
HDR
species

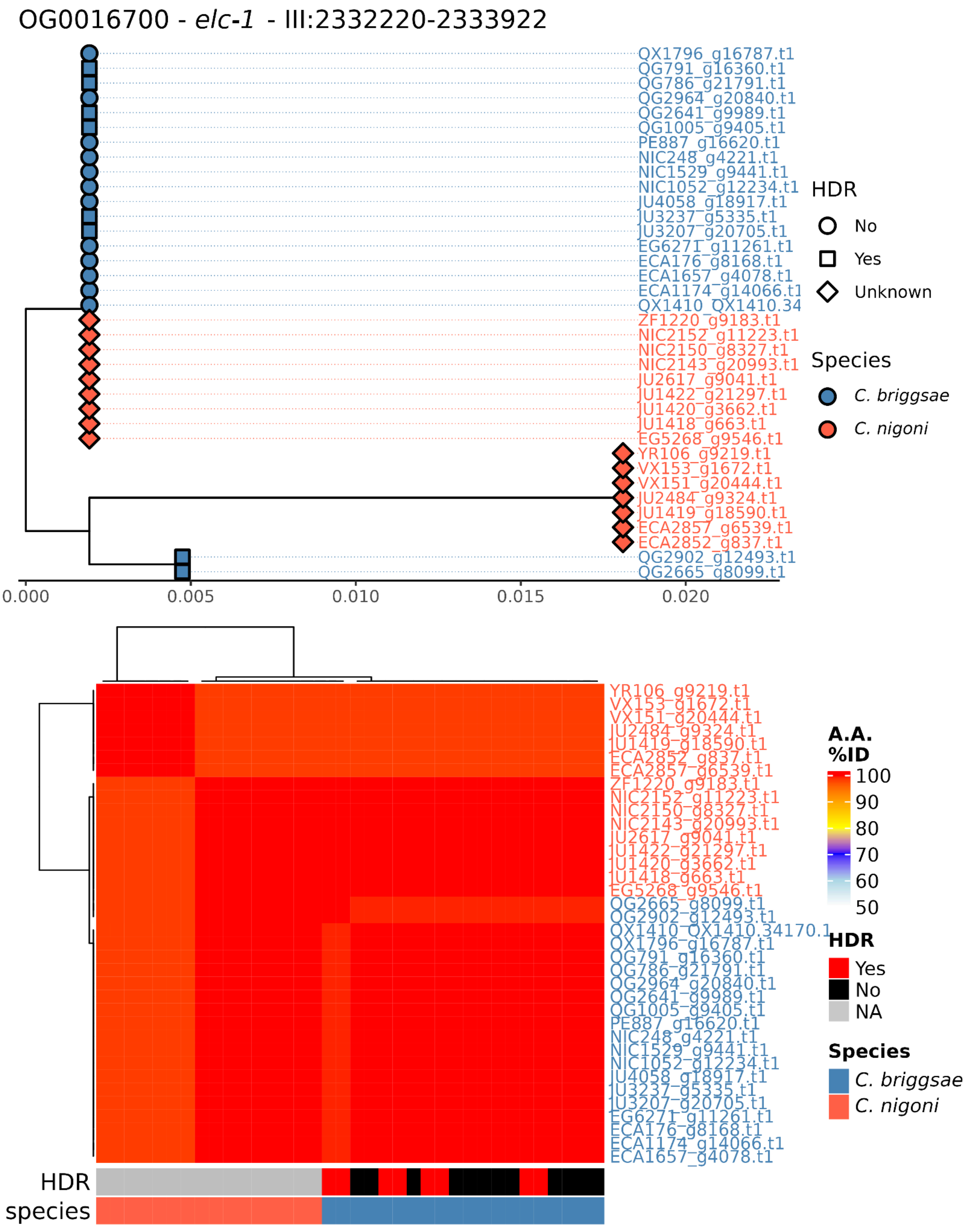

OG0016700 - elc-1 - III:2332220-2333922
QX1796_g16787.t1
QG791_g16360.t1
QG786_g21791.t1
QG2964_g20840.t1
QG2641_g9989.t1
QG1005_g9405.t1
PE887_g16620.t1
NIC248_g4221.t1
NIC1529_g9441.t1
NIC1052_g12234.t1
JU4058_g18917.t1
JU3237_g5335.t1
JU3207_g20705.t1
EG6271_g11261.t1
ECA176_g8168.t1
ECA1657_g4078.t1
ECA1174_g14066.t1
QX1410_QX1410.34
ZF1220_g9183.t1
NIC2152_g11223.t1
NIC2150_g8327.t1
NIC2143_g20993.t1
JU2617_g9041.t1
JU1422_g21297.t1
JU1420_g3662.t1
JU1418_g663.t1
EG5268_g9546.t1
YR106_g9219.t1
VX153_g1672.t1
VX151_g20444.t1
JU2484_g9324.t1
JU1419_g18590.t1
ECA2857_g6539.t1
ECA2852_g837.t1
QG2902_g12493.t1
QG2665_g8099.t1
0.000
0.005
0.010
0.015
0.020
HDR
No
Yes
Unknown
Species
C. briggsae
C. nigoni
YR106_g9219.t1
VX153_g1672.t1
VX151_g20444.t1
JU2484_g9324.t1
JU1419_g18590.t1
ECA2852_g837.t1
ECA2857_g6539.t1
ZF1220_g9183.t1
NIC2152_g11223.t1
NIC2150_g8327.t1
NIC2143_g20993.t1
JU2617_g9041.t1
JU1422_g21297.t1
JU1420_g3662.t1
JU1418_g663.t1
EG5268_g9546.t1
QG2665_g8099.t1
QG2902_g12493.t1
QX1410_QX1410.34170.1
QX1796_g16787.t1
QG791_g16360.t1
QG786_g21791.t1
QG2964_g20840.t1
QG2641_g9989.t1
QG1005_g9405.t1
PE887_g16620.t1
NIC248_g4221.t1
NIC1529_g9441.t1
NIC1052_g12234.t1
JU4058_g18917.t1
JU3237_g5335.t1
JU3207_g20705.t1
EG6271_g11261.t1
ECA176_g8168.t1
ECA1174_g14066.t1
ECA1657_g4078.t1
A.A. %ID
100
90
80
70
60
50
HDR
Yes
No
NA
Species
C. briggsae
C. nigoni
HDR
species

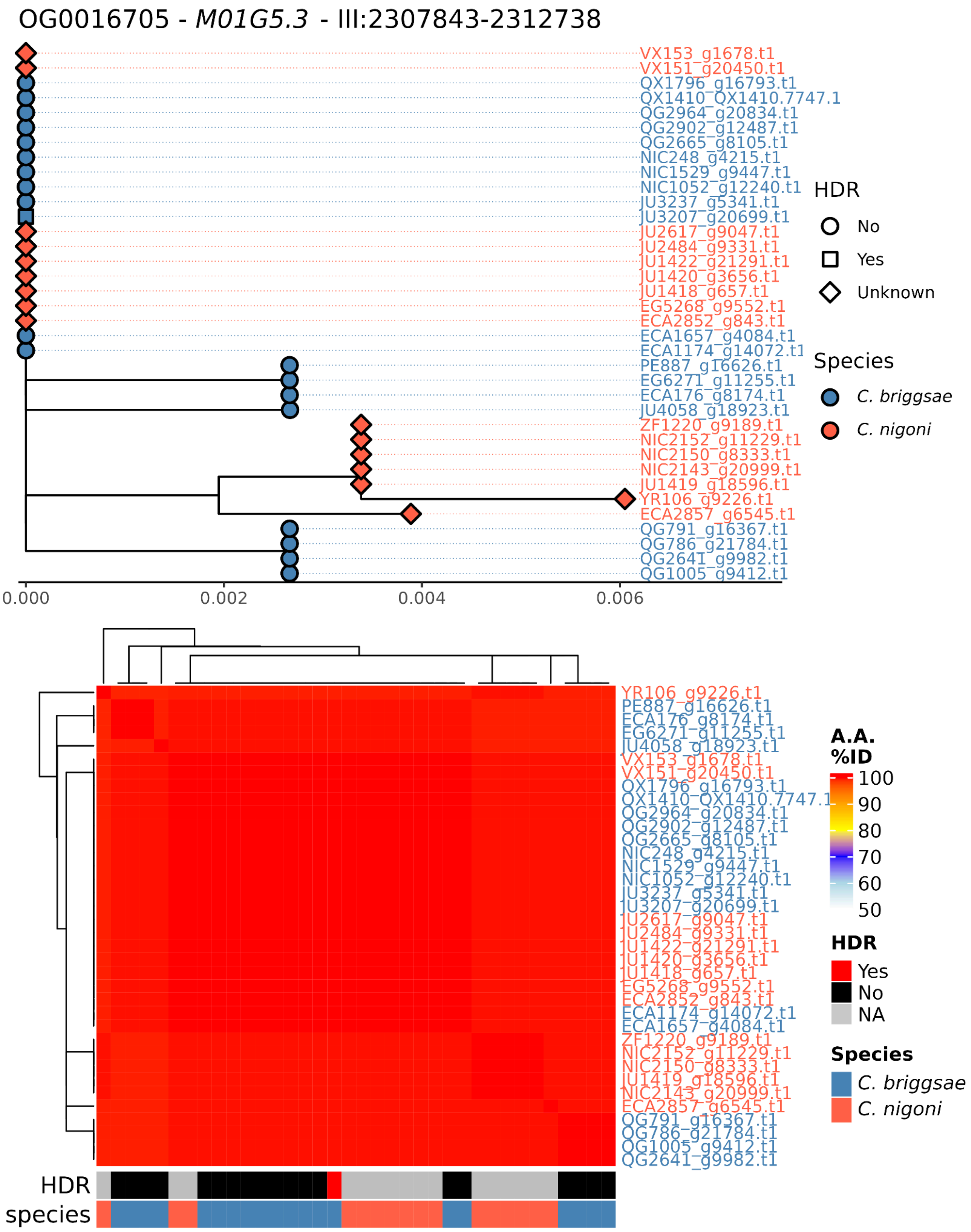

OG0016705 - *M01G5.3* - III:2307843-2312738
VX153_g1678.t1
VX151_g20450.t1
QX1796_g16793.t1
QX1410_QX1410.7747.1
QG2964_g20834.t1
QG2902_g12487.t1
QG2665_g8105.t1
NIC248_g4215.t1
NIC1529_g9447.t1
NIC1052_g12240.t1
JU3237_g5341.t1
JU3207_g20699.t1
JU2617_g9047.t1
JU2484_g9331.t1
JU1422_g21291.t1
JU1420_g3656.t1
JU1418_g657.t1
EG5268_g9552.t1
ECA2852_g843.t1
ECA1657_g4084.t1
ECA1174_g14072.t1
PE887_g16626.t1
EG6271_g11255.t1
ECA176_g8174.t1
JU4058_g18923.t1
ZF1220_g9189.t1
NIC2152_g11229.t1
NIC2150_g8333.t1
NIC2143_g20999.t1
JU1419_g18596.t1
YR106_g9226.t1
ECA2857_g6545.t1
QG791_g16367.t1
QG786_g21784.t1
QG2641_g9982.t1
QG1005_g9412.t1
0.000
0.002
0.004
0.006
HDR
No
Yes
Unknown
Species
*C. briggsae*
*C. nigoni*
A.A. %ID
100
90
80
70
60
50
HDR
Yes
No
NA
Species
*C. briggsae*
*C. nigoni*
HDR
species

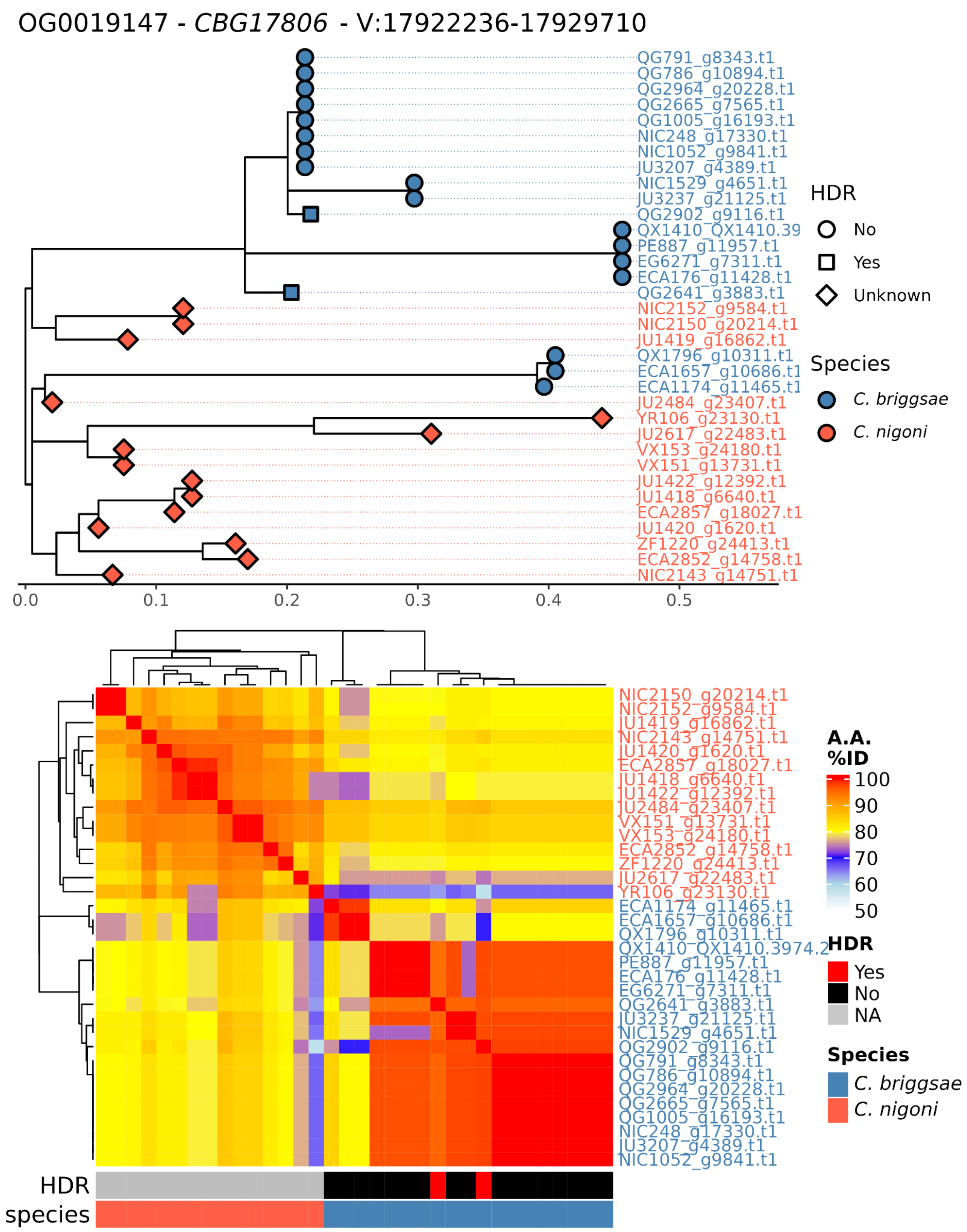
OG0019147 - CBG17806 - V:17922236-17929710
QG791_g8343.t1
QG786_g10894.t1
QG2964_g20228.t1
QG2665_g7565.t1
QG1005_g16193.t1
NIC248_g17330.t1
NIC1052_g9841.t1
JU3207_g4389.t1
NIC1529_g4651.t1
JU3237_g21125.t1
QG2902_g9116.t1
QX1410_QX1410.39
PE887_g11957.t1
EG6271_g7311.t1
ECA176_g11428.t1
QG2641_g3883.t1
NIC2152_g9584.t1
NIC2150_g20214.t1
JU1419_g16862.t1
QX1796_g10311.t1
ECA1657_g10686.t1
ECA1174_g11465.t1
JU2484_g23407.t1
YR106_g23130.t1
JU2617_g22483.t1
VX153_g24180.t1
VX151_g13731.t1
JU1422_g12392.t1
JU1418_g6640.t1
ECA2857_g18027.t1
JU1420_g1620.t1
ZF1220_g24413.t1
ECA2852_g14758.t1
NIC2143_g14751.t1
0.0
0.1
0.2
0.3
0.4
0.5
HDR
No
Yes
Unknown
Species
C. briggsae
C. nigoni
NIC2150_g20214.t1
NIC2152_g9584.t1
JU1419_g16862.t1
NIC2143_g14751.t1
JU1420_g1620.t1
ECA2857_g18027.t1
JU1418_g6640.t1
JU1422_g12392.t1
JU2484_g23407.t1
VX151_g13731.t1
VX153_g24180.t1
ECA2852_g14758.t1
ZF1220_g24413.t1
JU2617_g22483.t1
YR106_g23130.t1
ECA1174_g11465.t1
ECA1657_g10686.t1
QX1796_g10311.t1
QX1410_QX1410.3974.2
PE887_g11957.t1
ECA176_g11428.t1
EG6271_g7311.t1
QG2641_g3883.t1
JU3237_g21125.t1
NIC1529_g4651.t1
QG2902_g9116.t1
QG791_g8343.t1
QG786_g10894.t1
QG2964_g20228.t1
QG2665_g7565.t1
QG1005_g16193.t1
NIC248_g17330.t1
JU3207_g4389.t1
NIC1052_g9841.t1
HDR
species
A.A. %ID
100
90
80
70
60
50
HDR
Yes
No
NA
Species
C. briggsae
C. nigoni

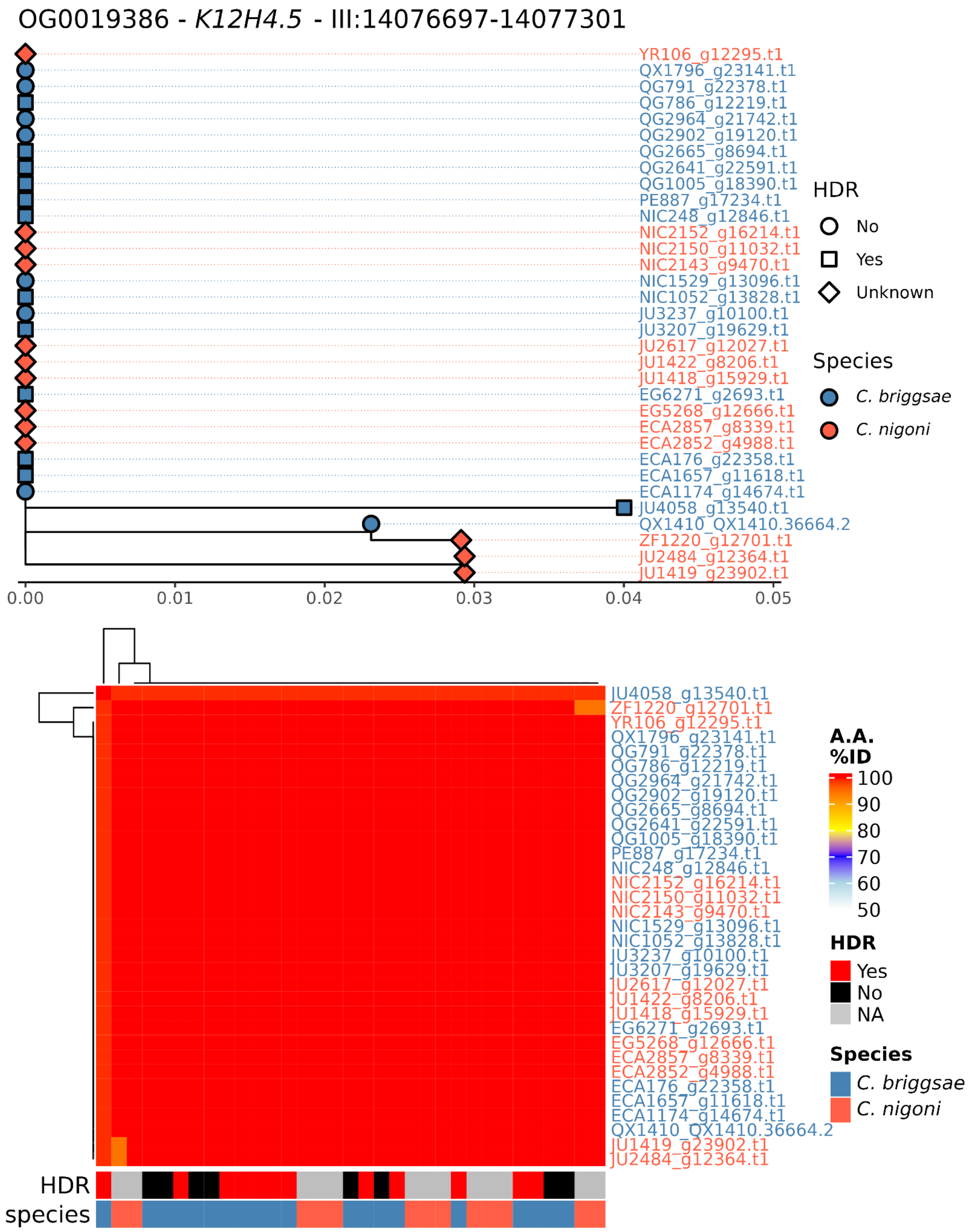

OG0019386 - *K12H4.5* - III:14076697-14077301
YR106_g12295.t1
QX1796_g23141.t1
QG791_g22378.t1
QG786_g12219.t1
QG2964_g21742.t1
QG2902_g19120.t1
QG2665_g8694.t1
QG2641_g22591.t1
QG1005_g18390.t1
PE887_g17234.t1
NIC248_g12846.t1
NIC2152_g16214.t1
NIC2150_g11032.t1
NIC2143_g9470.t1
NIC1529_g13096.t1
NIC1052_g13828.t1
JU3237_g10100.t1
JU3207_g19629.t1
JU2617_g12027.t1
JU1422_g8206.t1
JU1418_g15929.t1
EG6271_g2693.t1
EG5268_g12666.t1
ECA2857_g8339.t1
ECA2852_g4988.t1
ECA176_g22358.t1
ECA1657_g11618.t1
ECA1174_g14674.t1
JU4058_g13540.t1
QX1410_QX1410.36664.2
ZF1220_g12701.t1
JU2484_g12364.t1
JU1419_g23902.t1
0.00
0.01
0.02
0.03
0.04
0.05
HDR
No
Yes
Unknown
Species
*C. briggsae*
*C. nigoni*
JU4058_g13540.t1
ZF1220_g12701.t1
YR106_g12295.t1
QX1796_g23141.t1
QG791_g22378.t1
QG786_g12219.t1
QG2964_g21742.t1
QG2902_g19120.t1
QG2665_g8694.t1
QG2641_g22591.t1
QG1005_g18390.t1
PE887_g17234.t1
NIC248_g12846.t1
NIC2152_g16214.t1
NIC2150_g11032.t1
NIC2143_g9470.t1
NIC1529_g13096.t1
NIC1052_g13828.t1
JU3237_g10100.t1
JU3207_g19629.t1
JU2617_g12027.t1
JU1422_g8206.t1
JU1418_g15929.t1
EG6271_g2693.t1
EG5268_g12666.t1
ECA2857_g8339.t1
ECA2852_g4988.t1
ECA176_g22358.t1
ECA1657_g11618.t1
ECA1174_g14674.t1
QX1410_QX1410.36664.2
JU1419_g23902.t1
JU2484_g12364.t1
A.A. %ID
100
90
80
70
60
50
HDR
Yes
No
NA
Species
*C. briggsae*
*C. nigoni*
HDR
species

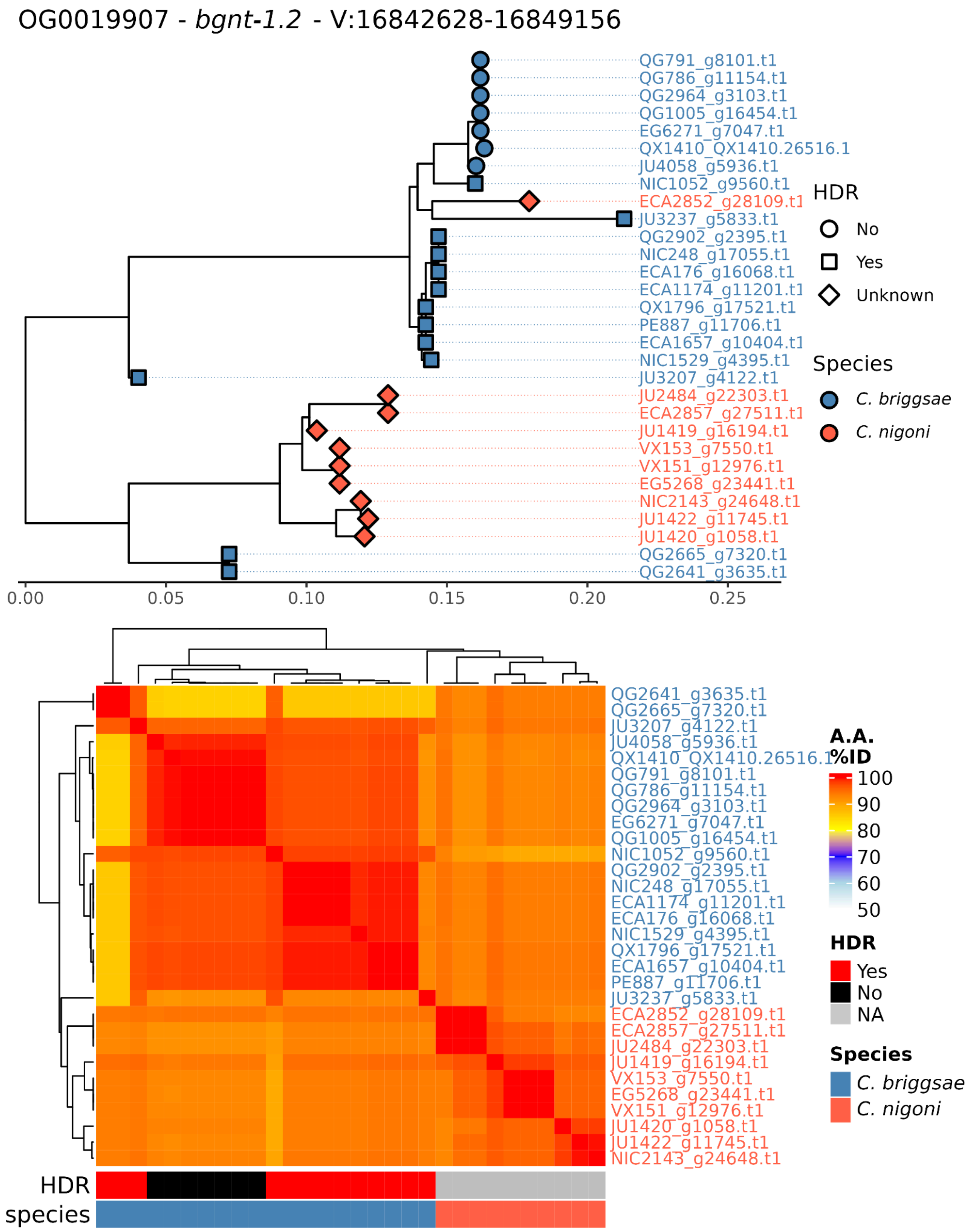
OG0019907 - bgnt-1.2 - V:16842628-16849156
QG791_g8101.t1
QG786_g11154.t1
QG2964_g3103.t1
QG1005_g16454.t1
EG6271_g7047.t1
QX1410_QX1410.26516.1
JU4058_g5936.t1
NIC1052_g9560.t1
ECA2852_g28109.t1
JU3237_g5833.t1
QG2902_g2395.t1
NIC248_g17055.t1
ECA176_g16068.t1
ECA1174_g11201.t1
QX1796_g17521.t1
PE887_g11706.t1
ECA1657_g10404.t1
NIC1529_g4395.t1
JU3207_g4122.t1
JU2484_g22303.t1
ECA2857_g27511.t1
JU1419_g16194.t1
VX153_g7550.t1
VX151_g12976.t1
EG5268_g23441.t1
NIC2143_g24648.t1
JU1422_g11745.t1
JU1420_g1058.t1
QG2665_g7320.t1
QG2641_g3635.t1
HDR
No
Yes
Unknown
Species
C. briggsae
C. nigoni
0.00
0.05
0.10
0.15
0.20
0.25
QG2641_g3635.t1
QG2665_g7320.t1
JU3207_g4122.t1
JU4058_g5936.t1
QX1410_QX1410.26516.1
QG791_g8101.t1
QG786_g11154.t1
QG2964_g3103.t1
EG6271_g7047.t1
QG1005_g16454.t1
NIC1052_g9560.t1
QG2902_g2395.t1
NIC248_g17055.t1
ECA1174_g11201.t1
ECA176_g16068.t1
NIC1529_g4395.t1
QX1796_g17521.t1
ECA1657_g10404.t1
PE887_g11706.t1
JU3237_g5833.t1
ECA2852_g28109.t1
ECA2857_g27511.t1
JU2484_g22303.t1
JU1419_g16194.t1
VX153_g7550.t1
EG5268_g23441.t1
VX151_g12976.t1
JU1420_g1058.t1
JU1422_g11745.t1
NIC2143_g24648.t1
A.A. %ID
100
90
80
70
60
50
HDR
Yes
No
NA
Species
C. briggsae
C. nigoni
HDR
species

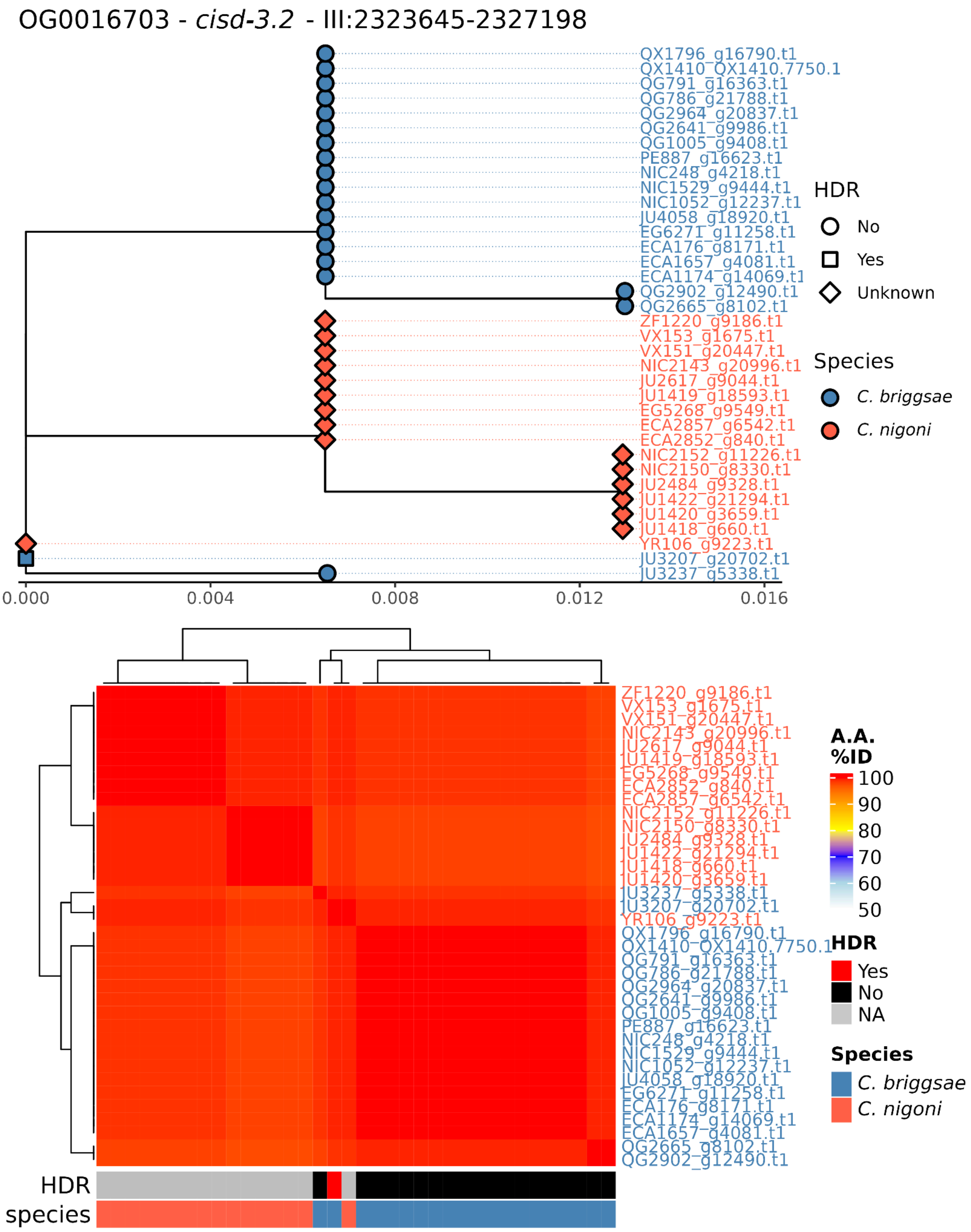
OG0016703 - *cisd-3.2* - III:2323645-2327198
QX1796_g16790.t1
QX1410_QX1410.7750.1
QG791_g16363.t1
QG786_g21788.t1
QG2964_g20837.t1
QG2641_g9986.t1
QG1005_g9408.t1
PE887_g16623.t1
NIC248_g4218.t1
NIC1529_g9444.t1
NIC1052_g12237.t1
JU4058_g18920.t1
EG6271_g11258.t1
ECA176_g8171.t1
ECA1657_g4081.t1
ECA1174_g14069.t1
QG2902_g12490.t1
QG2665_g8102.t1
ZF1220_g9186.t1
VX153_g1675.t1
VX151_g20447.t1
NIC2143_g20996.t1
JU2617_g9044.t1
JU1419_g18593.t1
EG5268_g9549.t1
ECA2857_g6542.t1
ECA2852_g840.t1
NIC2152_g11226.t1
NIC2150_g8330.t1
JU2484_g9328.t1
JU1422_g21294.t1
JU1420_g3659.t1
JU1418_g660.t1
YR106_g9223.t1
JU3207_g20702.t1
JU3237_g5338.t1
0.000
0.004
0.008
0.012
0.016
HDR
No
Yes
Unknown
Species
*C. briggsae*
*C. nigoni*
A.A. %ID
100
90
80
70
60
50
HDR
Yes
No
NA
Species
*C. briggsae*
*C. nigoni*
HDR
species

OG0015165 - *Y11D7A.10* - IV:4040191-4041333

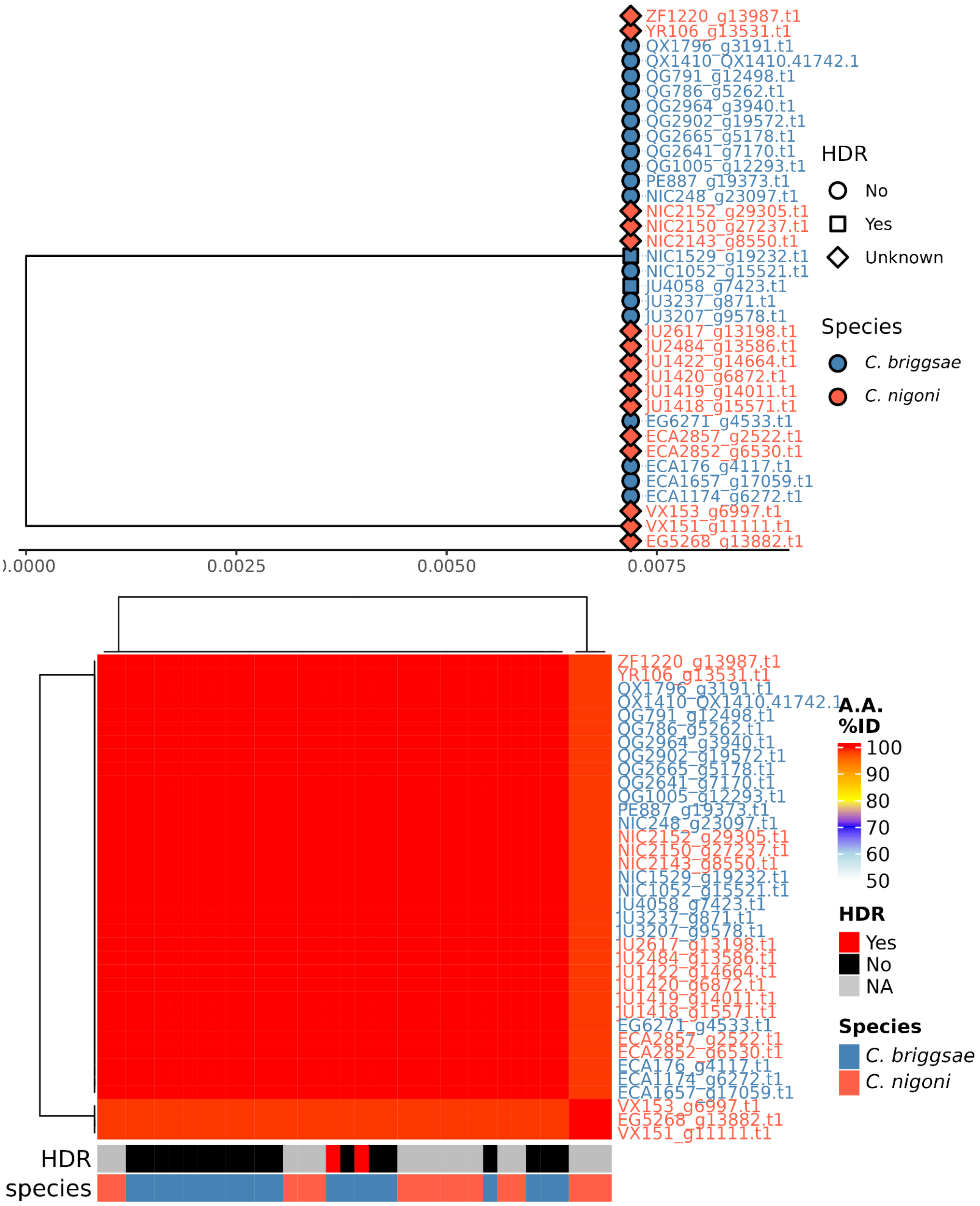

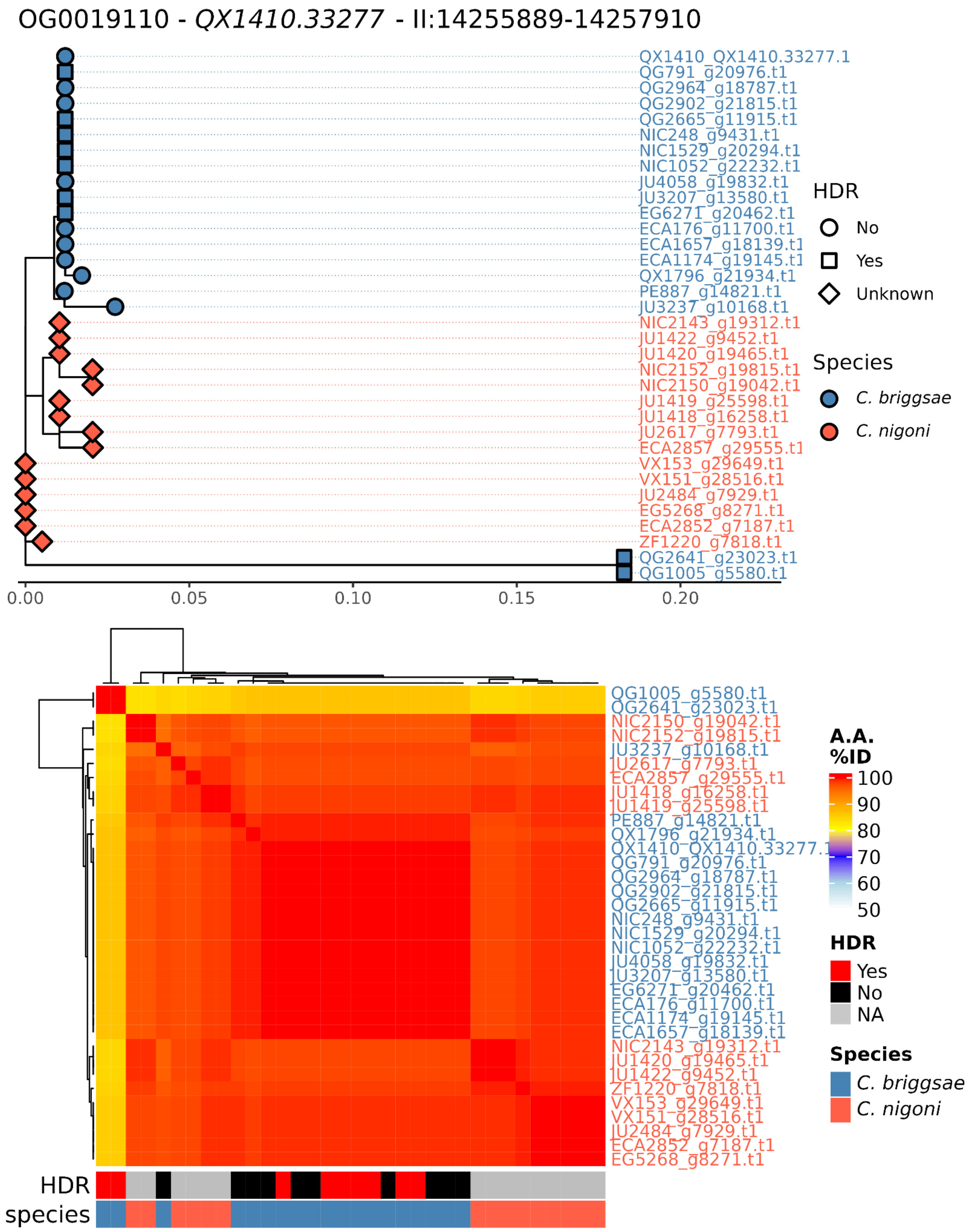
OG0019110 - QX1410.33277 - II:14255889-14257910
QX1410_QX1410.33277.1
QG791_g20976.t1
QG2964_g18787.t1
QG2902_g21815.t1
QG2665_g11915.t1
NIC248_g9431.t1
NIC1529_g20294.t1
NIC1052_g22232.t1
JU4058_g19832.t1
JU3207_g13580.t1
EG6271_g20462.t1
ECA176_g11700.t1
ECA1657_g18139.t1
ECA1174_g19145.t1
QX1796_g21934.t1
PE887_g14821.t1
JU3237_g10168.t1
NIC2143_g19312.t1
JU1422_g9452.t1
JU1420_g19465.t1
NIC2152_g19815.t1
NIC2150_g19042.t1
JU1419_g25598.t1
JU1418_g16258.t1
JU2617_g7793.t1
ECA2857_g29555.t1
VX153_g29649.t1
VX151_g28516.t1
JU2484_g7929.t1
EG5268_g8271.t1
ECA2852_g7187.t1
ZF1220_g7818.t1
QG2641_g23023.t1
QG1005_g5580.t1
0.00
0.05
0.10
0.15
0.20
HDR
No
Yes
Unknown
Species
C. briggsae
C. nigoni
A.A. %ID
100
90
80
70
60
50
HDR
Yes
No
NA
Species
C. briggsae
C. nigoni
HDR
species

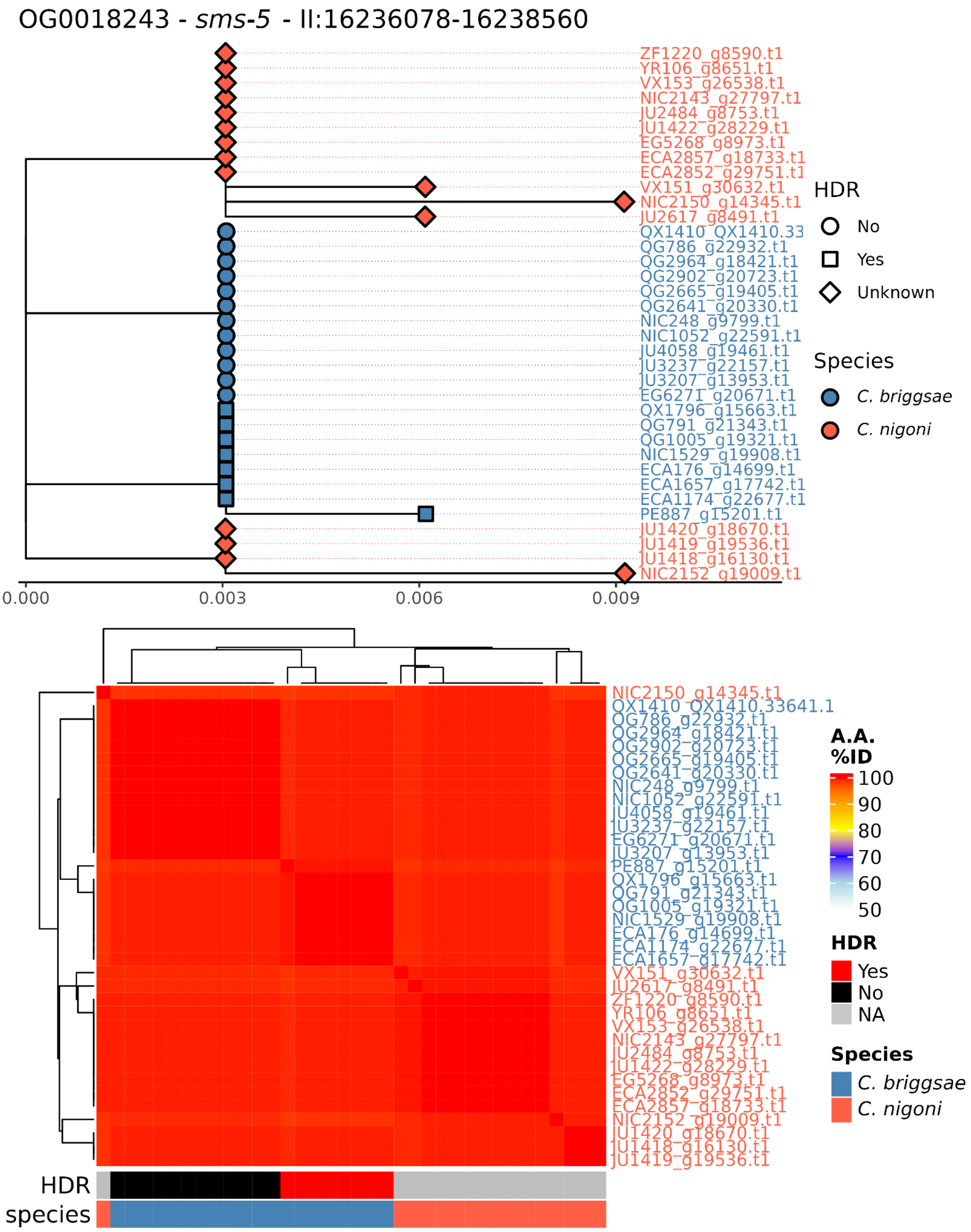
OG0018243 - *sms-5* - II:16236078-16238560
ZF1220_g8590.t1
YR106_g8651.t1
VX153_g26538.t1
NIC2143_g27797.t1
JU2484_g8753.t1
JU1422_g28229.t1
EG5268_g8973.t1
ECA2857_g18733.t1
ECA2852_g29751.t1
VX151_g30632.t1
NIC2150_g14345.t1
JU2617_g8491.t1
QX1410_QX1410.33
QG786_g22932.t1
QG2964_g18421.t1
QG2902_g20723.t1
QG2665_g19405.t1
QG2641_g20330.t1
NIC248_g9799.t1
NIC1052_g22591.t1
JU4058_g19461.t1
JU3237_g22157.t1
JU3207_g13953.t1
EG6271_g20671.t1
QX1796_g15663.t1
QG791_g21343.t1
QG1005_g19321.t1
NIC1529_g19908.t1
ECA176_g14699.t1
ECA1657_g17742.t1
ECA1174_g22677.t1
PE887_g15201.t1
JU1420_g18670.t1
JU1419_g19536.t1
JU1418_g16130.t1
NIC2152_g19009.t1
0.000
0.003
0.006
0.009
HDR
No
Yes
Unknown
Species
*C. briggsae*
*C. nigoni*
NIC2150_g14345.t1
QX1410_QX1410.33641.1
QG786_g22932.t1
QG2964_g18421.t1
QG2902_g20723.t1
QG2665_g19405.t1
QG2641_g20330.t1
NIC248_g9799.t1
NIC1052_g22591.t1
JU4058_g19461.t1
JU3237_g22157.t1
EG6271_g20671.t1
JU3207_g13953.t1
PE887_g15201.t1
QX1796_g15663.t1
QG791_g21343.t1
QG1005_g19321.t1
NIC1529_g19908.t1
ECA176_g14699.t1
ECA1174_g22677.t1
ECA1657_g17742.t1
VX151_g30632.t1
JU2617_g8491.t1
ZF1220_g8590.t1
YR106_g8651.t1
VX153_g26538.t1
NIC2143_g27797.t1
JU2484_g8753.t1
JU1422_g28229.t1
EG5268_g8973.t1
ECA2852_g29751.t1
ECA2857_g18733.t1
NIC2152_g19009.t1
JU1420_g18670.t1
JU1418_g16130.t1
JU1419_g19536.t1
HDR
species
**A.A. %ID**
100
90
80
70
60
50
**HDR**
Yes
No
NA
**Species**
*C. briggsae*
*C. nigoni*

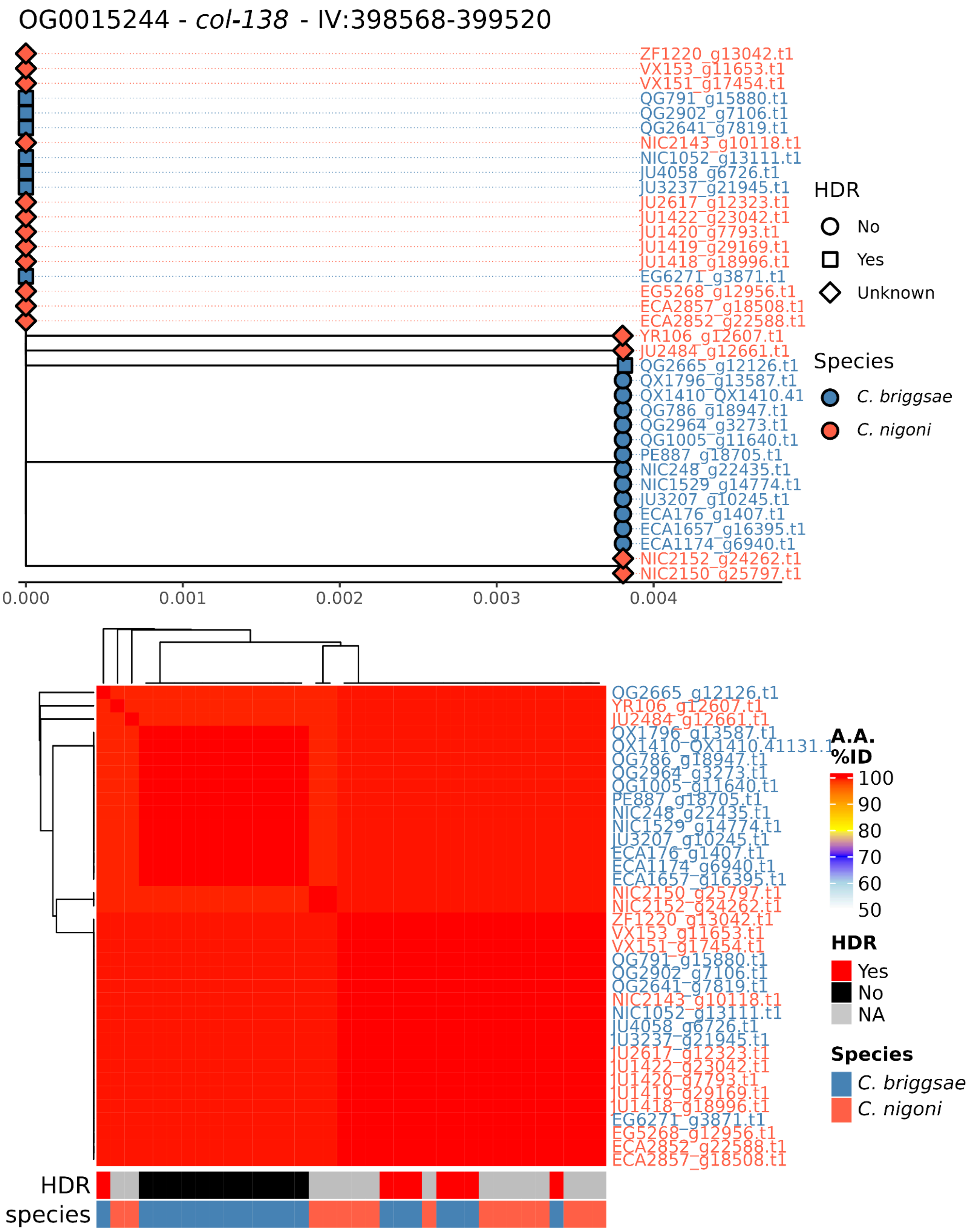

OG0015244 - *col-138* - IV:398568-399520
ZF1220_g13042.t1
VX153_g11653.t1
VX151_g17454.t1
QG791_g15880.t1
QG2902_g7106.t1
QG2641_g7819.t1
NIC2143_g10118.t1
NIC1052_g13111.t1
JU4058_g6726.t1
JU3237_g21945.t1
JU2617_g12323.t1
JU1422_g23042.t1
JU1420_g7793.t1
JU1419_g29169.t1
JU1418_g18996.t1
EG6271_g3871.t1
EG5268_g12956.t1
ECA2857_g18508.t1
ECA2852_g22588.t1
YR106_g12607.t1
JU2484_g12661.t1
QG2665_g12126.t1
QX1796_g13587.t1
QX1410_QX1410.41
QG786_g18947.t1
QG2964_g3273.t1
QG1005_g11640.t1
PE887_g18705.t1
NIC248_g22435.t1
NIC1529_g14774.t1
JU3207_g10245.t1
ECA176_g1407.t1
ECA1657_g16395.t1
ECA1174_g6940.t1
NIC2152_g24262.t1
NIC2150_g25797.t1
0.000
0.001
0.002
0.003
0.004
HDR
No
Yes
Unknown
Species
*C. briggsae*
*C. nigoni*
A.A. %ID
100
90
80
70
60
50
HDR
Yes
No
NA
Species
*C. briggsae*
*C. nigoni*
HDR
species

OG0016095 - *Y39B6A.9* - V:18262053-18263819

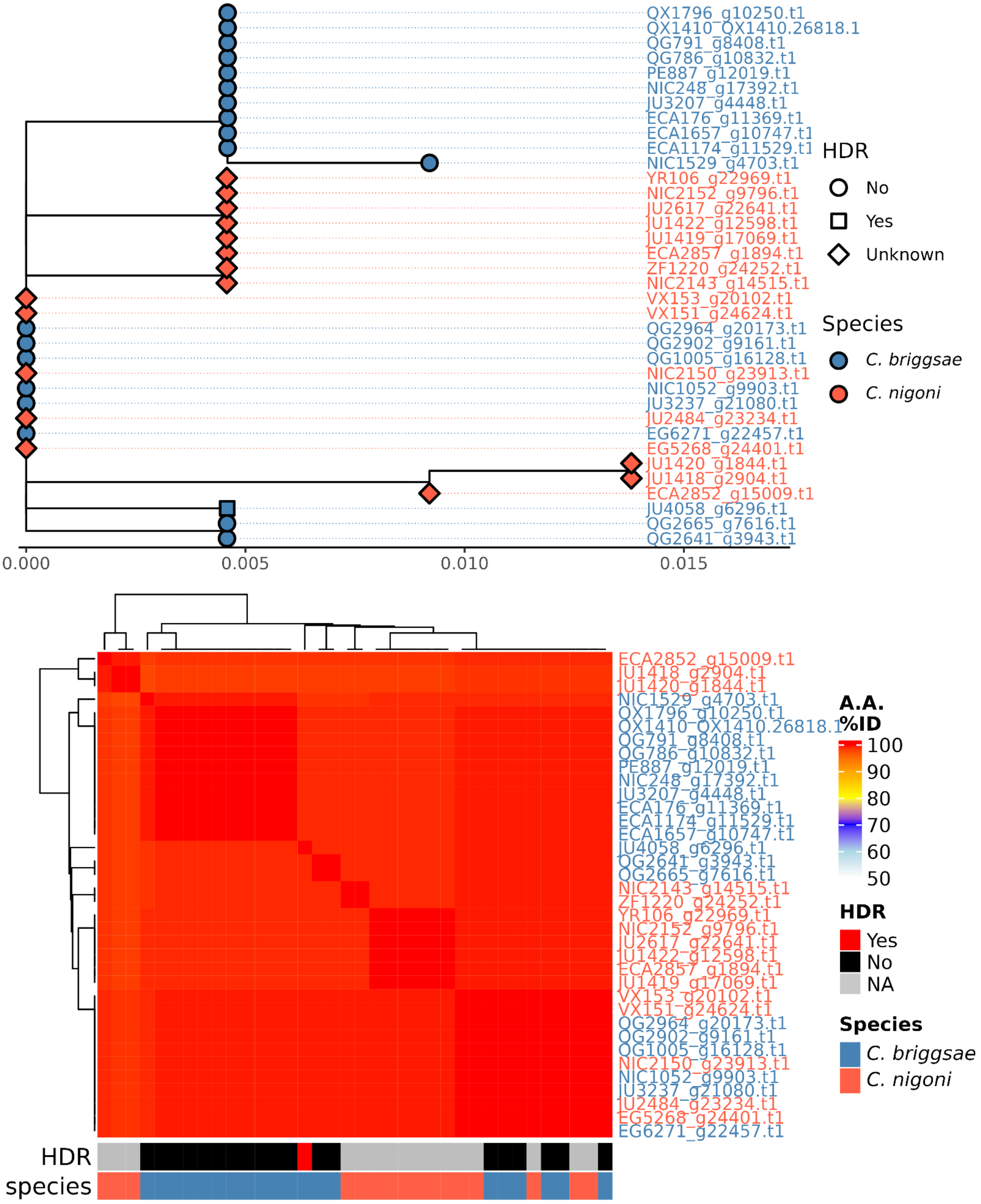

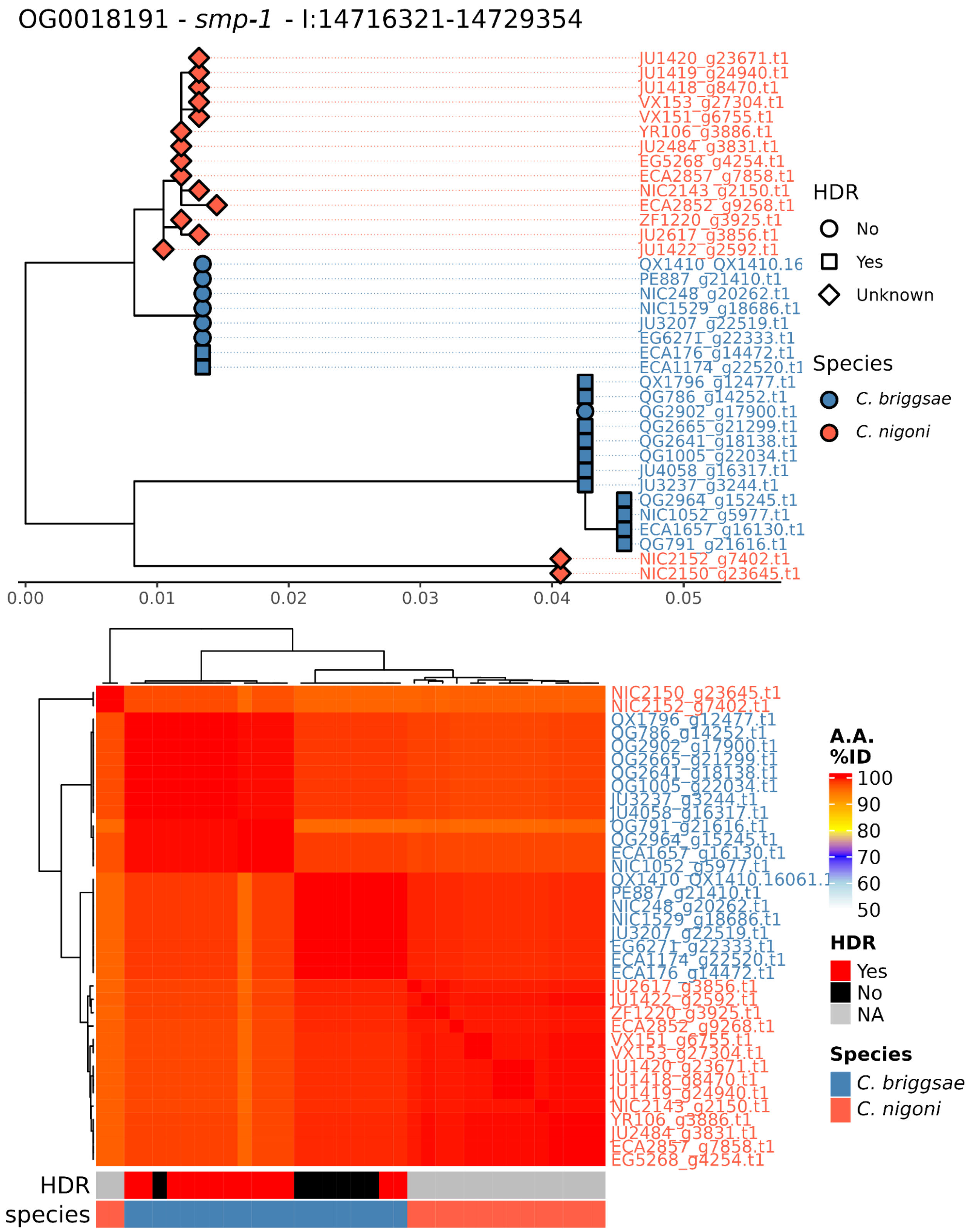
OG0018191 - smp-1 - I:14716321-14729354
JU1420_g23671.t1
JU1419_g24940.t1
JU1418_g8470.t1
VX153_g27304.t1
VX151_g6755.t1
YR106_g3886.t1
JU2484_g3831.t1
EG5268_g4254.t1
ECA2857_g7858.t1
NIC2143_g2150.t1
ECA2852_g9268.t1
ZF1220_g3925.t1
JU2617_g3856.t1
JU1422_g2592.t1
QX1410_QX1410.16
PE887_g21410.t1
NIC248_g20262.t1
NIC1529_g18686.t1
JU3207_g22519.t1
EG6271_g22333.t1
ECA176_g14472.t1
ECA1174_g22520.t1
QX1796_g12477.t1
QG786_g14252.t1
QG2902_g17900.t1
QG2665_g21299.t1
QG2641_g18138.t1
QG1005_g22034.t1
JU4058_g16317.t1
JU3237_g3244.t1
QG2964_g15245.t1
NIC1052_g5977.t1
ECA1657_g16130.t1
QG791_g21616.t1
NIC2152_g7402.t1
NIC2150_g23645.t1
0.00
0.01
0.02
0.03
0.04
0.05
HDR
No
Yes
Unknown
Species
C. briggsae
C. nigoni
NIC2150_g23645.t1
NIC2152_g7402.t1
QX1796_g12477.t1
QG786_g14252.t1
QG2902_g17900.t1
QG2665_g21299.t1
QG2641_g18138.t1
QG1005_g22034.t1
JU3237_g3244.t1
JU4058_g16317.t1
QG791_g21616.t1
QG2964_g15245.t1
ECA1657_g16130.t1
NIC1052_g5977.t1
QX1410_QX1410.16061.1
PE887_g21410.t1
NIC248_g20262.t1
NIC1529_g18686.t1
JU3207_g22519.t1
EG6271_g22333.t1
ECA1174_g22520.t1
ECA176_g14472.t1
JU2617_g3856.t1
JU1422_g2592.t1
ZF1220_g3925.t1
ECA2852_g9268.t1
VX151_g6755.t1
VX153_g27304.t1
JU1420_g23671.t1
JU1418_g8470.t1
JU1419_g24940.t1
NIC2143_g2150.t1
YR106_g3886.t1
JU2484_g3831.t1
ECA2857_g7858.t1
EG5268_g4254.t1
A.A. %ID
100
90
80
70
60
50
HDR
Yes
No
NA
Species
C. briggsae
C. nigoni
HDR
species

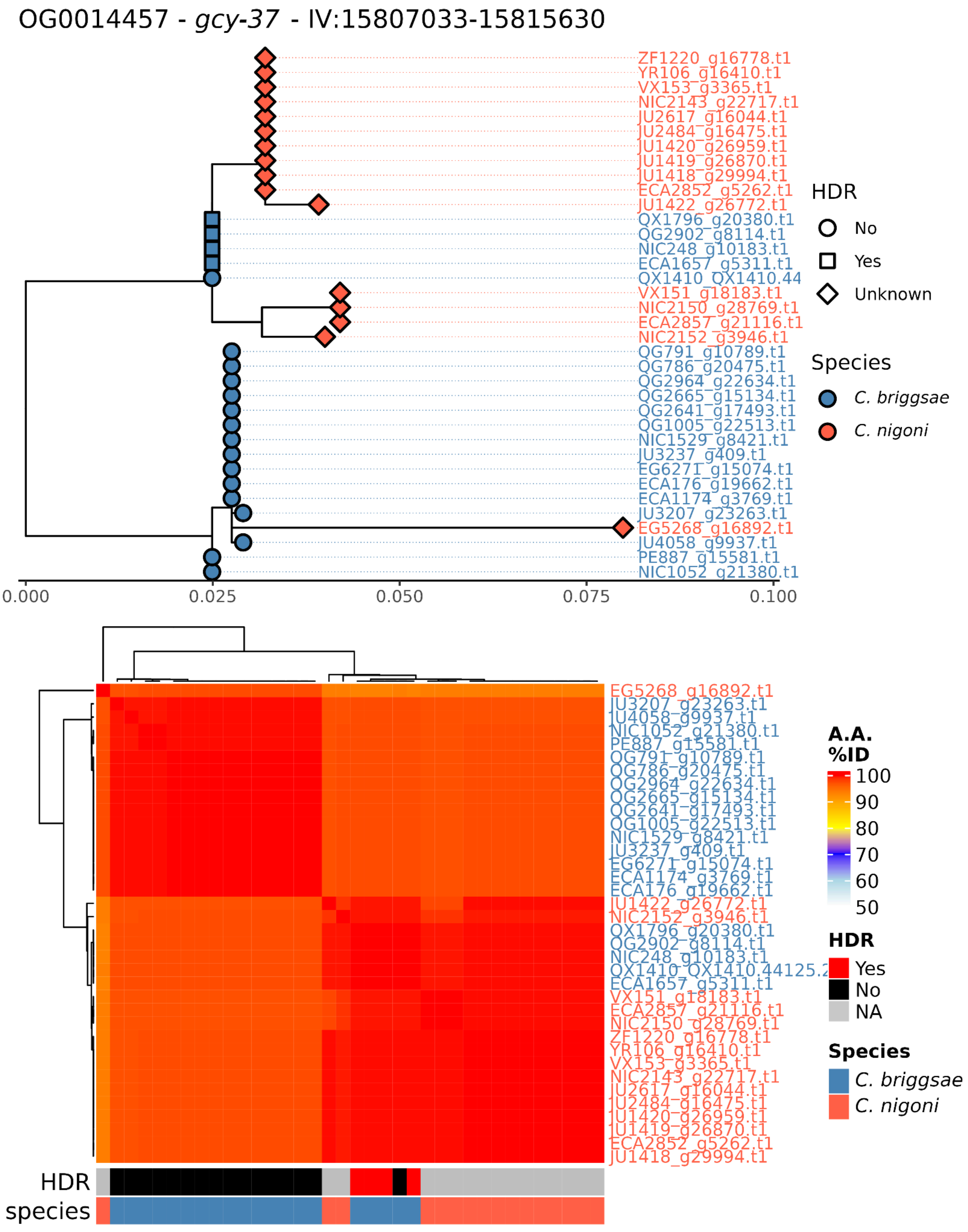
OG0014457 - *gcy-37* - IV:15807033-15815630
ZF1220_g16778.t1
YR106_g16410.t1
VX153_g3365.t1
NIC2143_g22717.t1
JU2617_g16044.t1
JU2484_g16475.t1
JU1420_g26959.t1
JU1419_g26870.t1
JU1418_g29994.t1
ECA2852_g5262.t1
JU1422_g26772.t1
QX1796_g20380.t1
QG2902_g8114.t1
NIC248_g10183.t1
ECA1657_g5311.t1
QX1410_QX1410.44
VX151_g18183.t1
NIC2150_g28769.t1
ECA2857_g21116.t1
NIC2152_g3946.t1
QG791_g10789.t1
QG786_g20475.t1
QG2964_g22634.t1
QG2665_g15134.t1
QG2641_g17493.t1
QG1005_g22513.t1
NIC1529_g8421.t1
JU3237_g409.t1
EG6271_g15074.t1
ECA176_g19662.t1
ECA1174_g3769.t1
JU3207_g23263.t1
EG5268_g16892.t1
JU4058_g9937.t1
PE887_g15581.t1
NIC1052_g21380.t1
0.000
0.025
0.050
0.075
0.100
HDR
No
Yes
Unknown
Species
*C. briggsae*
*C. nigoni*
EG5268_g16892.t1
JU3207_g23263.t1
JU4058_g9937.t1
NIC1052_g21380.t1
PE887_g15581.t1
QG791_g10789.t1
QG786_g20475.t1
QG2964_g22634.t1
QG2665_g15134.t1
QG2641_g17493.t1
QG1005_g22513.t1
NIC1529_g8421.t1
JU3237_g409.t1
EG6271_g15074.t1
ECA1174_g3769.t1
ECA176_g19662.t1
JU1422_g26772.t1
NIC2152_g3946.t1
QX1796_g20380.t1
QG2902_g8114.t1
NIC248_g10183.t1
QX1410_QX1410.44125.2
ECA1657_g5311.t1
VX151_g18183.t1
ECA2857_g21116.t1
NIC2150_g28769.t1
ZF1220_g16778.t1
YR106_g16410.t1
VX153_g3365.t1
NIC2143_g22717.t1
JU2617_g16044.t1
JU2484_g16475.t1
JU1420_g26959.t1
JU1419_g26870.t1
ECA2852_g5262.t1
JU1418_g29994.t1
HDR
species
A.A. %ID
100
90
80
70
60
50
HDR
Yes
No
NA
Species
*C. briggsae*
*C. nigoni*

OG0019038 - *F20H11.4* - IV:16350857-16353259

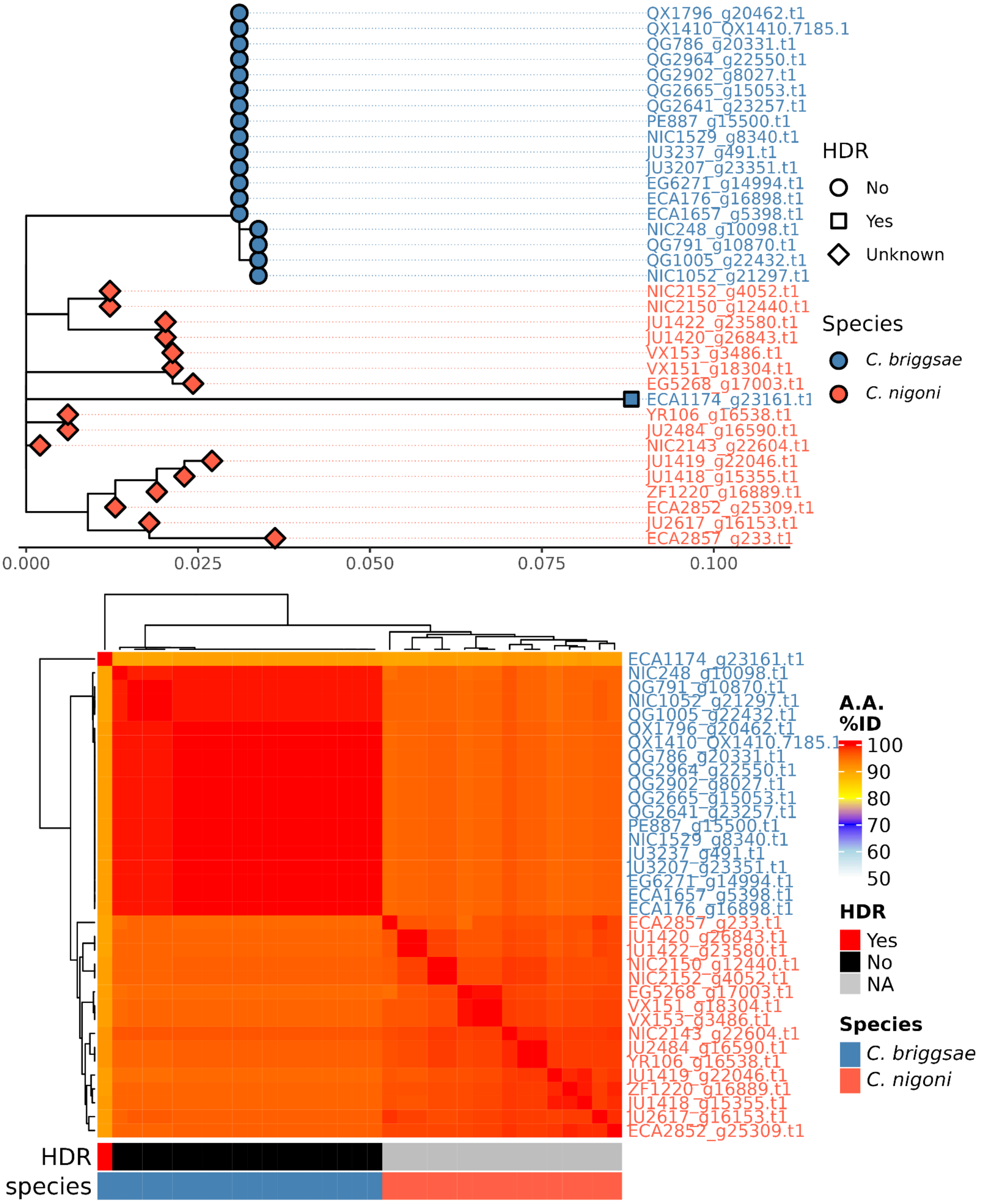

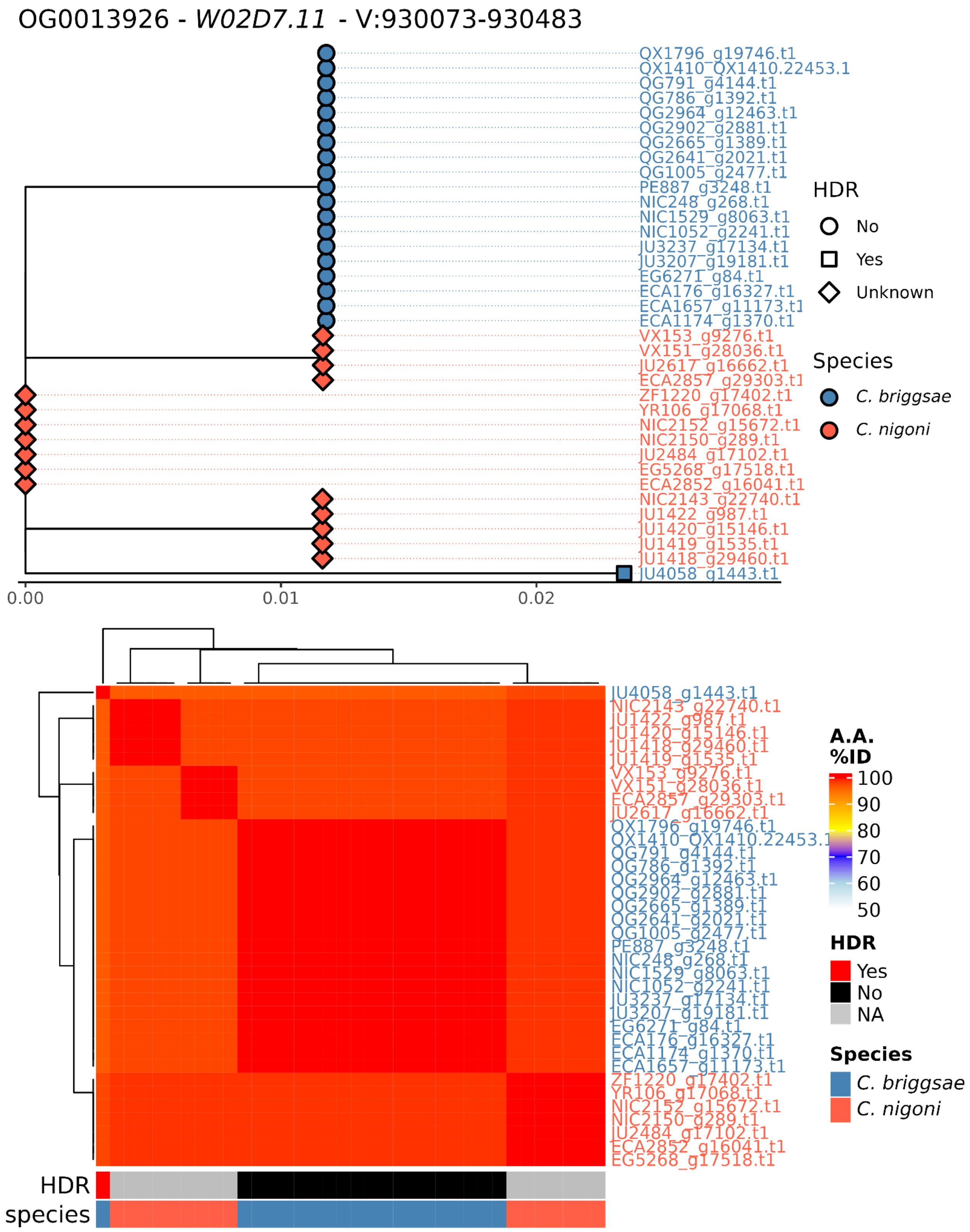
OG0013926 - W02D7.11 - V:930073-930483
QX1796_g19746.t1
QX1410_QX1410.22453.1
QG791_g4144.t1
QG786_g1392.t1
QG2964_g12463.t1
QG2902_g2881.t1
QG2665_g1389.t1
QG2641_g2021.t1
QG1005_g2477.t1
PE887_g3248.t1
NIC248_g268.t1
NIC1529_g8063.t1
NIC1052_g2241.t1
JU3237_g17134.t1
JU3207_g19181.t1
EG6271_g84.t1
ECA176_g16327.t1
ECA1657_g11173.t1
ECA1174_g1370.t1
VX153_g9276.t1
VX151_g28036.t1
JU2617_g16662.t1
ECA2857_g29303.t1
ZF1220_g17402.t1
YR106_g17068.t1
NIC2152_g15672.t1
NIC2150_g289.t1
JU2484_g17102.t1
EG5268_g17518.t1
ECA2852_g16041.t1
NIC2143_g22740.t1
JU1422_g987.t1
JU1420_g15146.t1
JU1419_g1535.t1
JU1418_g29460.t1
JU4058_g1443.t1
0.00
0.01
0.02
HDR
No
Yes
Unknown
Species
C. briggsae
C. nigoni
A.A. %ID
100
90
80
70
60
50
HDR
Yes
No
NA
Species
C. briggsae
C. nigoni
HDR
species
